# DNA methylation signatures of duplicate gene evolution in angiosperms

**DOI:** 10.1101/2020.08.31.275362

**Authors:** Sunil K. Kenchanmane Raju, S. Marshall Ledford, Chad E. Niederhuth

**Affiliations:** Department of Plant Biology, Michigan State University, East Lansing, MI 48824, U.S.A.; Vassar College, Poughkeepsie, NY 12604, U.S.A.; AgBioResearch, Michigan State University, East Lansing, MI 48824, U.S.A.

**Keywords:** Gene Duplication, Whole Genome Duplication, DNA methylation, Angiosperms

## Abstract

Gene duplication is a source of evolutionary novelty. DNA methylation may play a role in the evolution of duplicate genes through its association with gene expression. While this relationship is examined to varying extent in a few individual species, the generalizability of these results at either a broad phylogenetic scale with species of differing duplication histories or across a population, remains unknown. We apply a comparative epigenomics approach to 43 angiosperm species across the phylogeny and a population of 928 *Arabidopsis thaliana* accessions, examining the association of DNA methylation with paralog evolution. Genic DNA methylation is differentially associated with duplication type, the age of duplication, sequence evolution, and gene expression. Whole genome duplicates are typically enriched for CG-only gene-body methylated or unmethylated genes, while single-gene duplications are typically enriched for non-CG methylated or unmethylated genes. Non-CG methylation, in particular, was characteristic of more recent single-gene duplicates. Core angiosperm gene families are differentiated into those which preferentially retain paralogs and ‘duplication-resistant’ families, which convergently revert to singletons following duplication. Duplication-resistant families which still have paralogous copies are, uncharacteristically for core angiosperm genes, enriched for non-CG methylation. Non-CG methylated paralogs have higher rates of sequence evolution, higher frequency of presence-absence variation, and more limited expression. This suggests that silencing by non-CG methylation may be important to maintaining dosage following duplication and be a precursor to fractionation. Our results indicate that genic methylation marks differing evolutionary trajectories and fates between paralogous genes and have a role in maintaining dosage following duplication.

## Introduction

Gene and genome duplication increases organismal gene content, generating a repertoire for functional novelty (Bridges, 1935; Ohno, 1970; Flagel and Wendel, 2009). Whole-genome duplication (WGD) increases the entire gene content (Soltis et al., 2015) and is more pervasive in plants than in other eukaryotes (Otto and Whitton, 2000; Van de Peer et al., 2017; Cheng et al., 2018a). Small-scale and single-gene duplications (SGDs) are a continuous process with ongoing gene birth and death contributing significantly to gene content (Lynch and Conery, 2000; Maere et al., 2005; Panchy et al., 2016). The subsequent retention, divergence, or fractionation (loss) of paralogs are biased depending on the duplication type and gene function (Freeling, 2009; De Smet et al., 2013).

Factors determining the evolutionary fate of paralogs are an area of intense study, and DNA methylation is thought to be a contributing factor due to its association with gene expression (Rodin and Riggs, 2003; Wang et al., 2013b). Cytosine Methylation at CG dinucleotides is found throughout eukaryotes, while methylation of the non-CG trinucleotide CHG and CHH (H = A, T or C) contexts is limited to plants (Feng et al., 2010; Zemach et al., 2010). Plant genes have several distinct patterns of DNA methylation within coding regions (henceforth ‘genic methylation’) that are associated with gene expression (Niederhuth and Schmitz, 2017). Genes characterized by CG-only methylation in coding regions are referred to as gene-body methylated (gbM) (Tran et al., 2005; Zhang et al., 2006) and is frequently conserved between orthologous genes. GbM genes are typically broadly expressed and evolve more slowly (Takuno and Gaut, 2012; Takuno and Gaut, 2013; Niederhuth et al., 2016; Takuno et al., 2016). Some genes are methylated similar to transposable elements (TEs), having both CG and non-CG methylation within coding regions. This transposable-element like methylation (teM) is rarely conserved between orthologs and results in transcriptional silencing (Seymour et al., 2014; Niederhuth et al., 2016; El Baidouri et al., 2018). Most genes, however, are unmethylated (unM) and exhibit variable expression across tissues and conditions (Takuno and Gaut, 2012; Niederhuth et al., 2016).

DNA methylation could serve to buffer the genome against changes in gene dosage by modulating gene expression and facilitating functional divergence. Tissue-specific silencing of paralogs might lead to ‘epigenetic-complementation’, through sub-functionalization of expression and paralog retention (Adams et al., 2003; Rodin and Riggs, 2003). Alternatively, silencing may contribute to pseudogenization and subsequent fractionation (Hua et al., 2013; El Baidouri et al., 2018). Studies in plants (Hua et al., 2013; Wang et al., 2013b; Wang et al., 2014a; Kim et al., 2015; Wang et al., 2015; Wang et al., 2017; El Baidouri et al., 2018; Wang et al., 2018; Xu et al., 2018) and animals (Chang and Liao, 2012; Keller and Yi, 2014) have found that increasing DNA methylation differences between paralogs are associated with divergence in sequence evolution and expression. In *Glycine max*, gene transposition to heterochromatic regions resulted in silencing by non-CG methylation, increased sequence divergence, and likely pseudogenization (El Baidouri et al., 2018). In the highly-duplicated F-box family of *Arabidopsis thaliana*, silencing by DNA methylation and H3K27me3 was associated with increased sequence divergence and was proposed to have a role in maintaining dosage-balance (Hua et al., 2013).

Past studies of DNA methylation and gene duplication have been limited to individual species, focused primarily on WGDs, and often ignore the contextual differences of genic methylation. Lineage-specific variation in DNA methylation (Niederhuth et al., 2016), histories of gene duplication (Qiao et al., 2019), and differences in analysis have precluded an overarching understanding of the relationship between DNA methylation and paralog evolution. To address these issues, we analyze genic methylation contexts across 43 angiosperm species and a population of 928 *A. thaliana* ecotypes. We find overarching trends and relationships between genic methylation, the type and age of duplication, gene family, and paralog evolution. This work provides a broad phylogenetic and population-scale understanding of the role of DNA methylation in plant duplicate gene evolution and suggests DNA methylation may have a role in maintaining dosage prior to fractionation.

## Results

### Genic methylation across duplication types

We analyzed genic methylation and gene duplication for 43 angiosperm species (Table ST1). Genes were classified as gene-body methylated (gbM), unmethylated (unM), or transposable-element like methylated (teM) based on DNA methylation in coding regions (Figure S1, Table ST2). Gene duplicates were identified and classified (Table ST3) as either whole-genome duplicates (WGDs) or one of four types of single-gene duplicates (SGDs): tandem, proximal, translocated, or dispersed. Tandem duplicates occur through unequal crossing-over, resulting in adjacent paralogous copies (Zhang, 2003). Proximal duplicates are separated by several intervening genes and arose either through local transposition or interruption of ancient tandem duplication (Zhao et al., 1998; Freeling et al., 2008). Translocated duplicates (also known as ‘transposed’) are distally located pairs in which one of the genes is syntenic, and the other is non-syntenic (Wang et al., 2013a; Qiao et al., 2019) and can arise either by retrotransposition or DNA-based duplication (Cusack and Wolfe, 2007). Finally, dispersed duplicates are pairs that fit none of the above criteria and can arise through multiple mechanisms (Ganko et al., 2007; Freeling, 2009; Qiao et al., 2019).

We hypothesized that different duplication types would differ in genic methylation. Each duplication type was tested for enrichment or depletion of gbM, unM, and teM in each pieces (Figure 1, Table ST4). Across angiosperms, WGDs were more frequently enriched for gbM (27/43 enriched, 7/43 depleted) and unM (32/43 enriched, 5/43 depleted), and depleted for teM (3/43 enriched, 39/43 depleted). Notable exceptions include three Brassicaceae species (*Brassica oleracea, Brassica rapa*, and *Eutrema salsugineum*), three Cucurbitaceae (*Citrullus lanatus, Cucumis melo*, and *Cucumis sativus*) species, and *Solanum tuberosum*. The three Brassicaceae species are known to be depleted of gbM genome-wide (Bewick et al., 2016). No known depletion of gbM is documented in the Cucurbitaceae. While *C. melo* WGDs are depleted of gbM and enriched in TeM, *S. tuberosum* is the only species showing depletion of both gbM and unM, and enrichment of teM in WGDs. ‘Local’ tandem and proximal SGDs are more similar in enrichment/depletion to each other compared to ‘distal’ translocated and dispersed SGDs. Local SGDs are depleted of gbM (tandem & proximal – 40/43 depleted, 1/43 enriched) in all species except for the three gbM-deficient Brassicaceae species, and are enriched for unM in the majority of species (tandem – 39/43 enriched, 3/43 depleted; proximal – 31/43 enriched, 5/43 depleted). Tandem and proximal duplicates differed more in teM (tandem – 19/43 enriched, 14/43 depleted; proximal – 31/43 enriched, 1/43 depleted), with proximal duplicates showing more species enriched for teM than tandem. Like local SGDs, distal SGDs were more frequently depleted for gbM (translocated – 0/43 enriched, 27/43 depleted; dispersed – 11/43 enriched, 21/43 depleted), although, dispersed was the SGD most frequently enriched for gbM. Translocated and dispersed duplicates differ for unM; translocated duplicates have similar numbers enriched and depleted species (13/43 enriched, 14/43 depleted), while dispersed duplicates are depleted in a majority of species (0/43 enriched, 36/43 depleted). Distal SGDs are more frequently enriched for teM than local SGDs (translocated – 34/43 enriched, 4/43 depleted; dispersed – 42/43 enriched, 1/43 depleted). Increasing teM frequency from tandem to proximal to distal SGDs suggests that teM becomes more common as genes duplicate to increasingly different sequence or chromatin environments.

**Figure 1:**
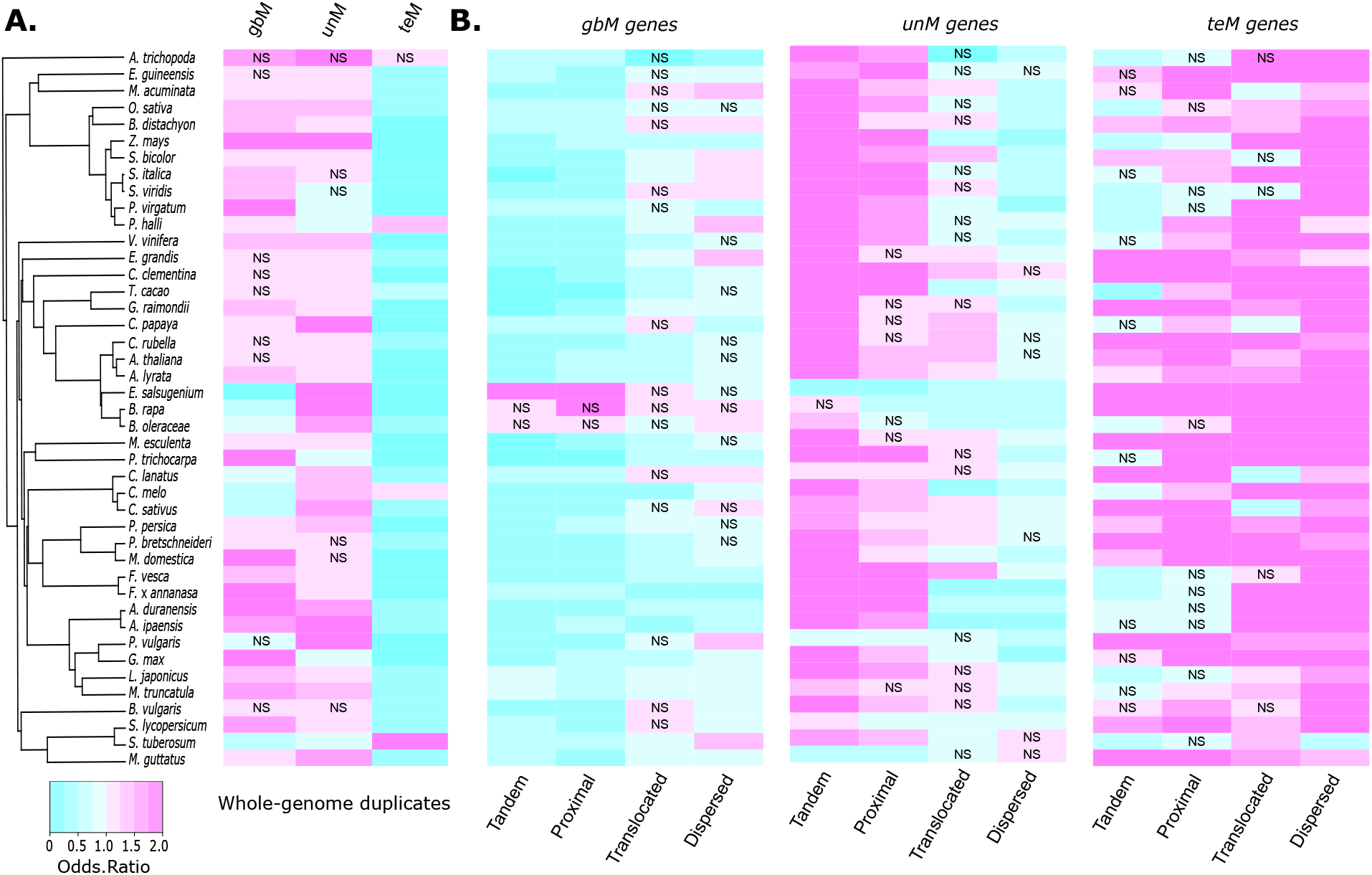
Patterns of genic methylation across different types of gene duplicates. Enrichment or depletion of each genic methylation class (gbM, unM, and teM) for whole genome duplicates **(A)** and different types of single gene duplicates **(B)**. Increasing shades of cyan indicates greater depletion, while increasing shades of magenta represents greater enrichment. Unless indicated, all associations are statistically significant at a FDR-corrected p-value < 0.05. ‘NS’ indicates no statistical significance.

### Effect of gene family on genic methylation and duplication

Gene families differ in their duplicability and retention. Past work has revealed ‘duplication-resistant’ gene families that repeatedly return to single-copy status (Paterson et al., 2006; De Smet et al., 2013; Li et al., 2016), while other gene families retain duplicates over long evolutionary timescales (Conant et al., 2014). To see how gene family composition affects the relationship between genic methylation and duplication, we identified orthogroups for the 43 species with methylome data and an additional 15 species included as outgroups. (Table ST1, Figure S2). Orthogroups showed a bimodal distribution, with the majority of orthogroups present in either a few species or conserved across most species (Li et al., 2016). Orthogroups represented in ≥ 51 species were classified as ‘core angiosperm’ genes and further divided as ‘core:single-copy’ (duplication-resistant) if represented by a single-copy in ≥ 70% species (Li et al., 2016) and the remainder as ‘core:multi-copy.’ Remaining orthogroups were classified based on increasing lineage-specificity: ‘cross family’ if present in multiple plant families, ‘family-specific’ if restricted to a single family, or ‘species/lineage specific’ if limited to a single species. Genic methylation shows more consistent enrichment and depletion across species for orthogroups than for duplication type (Figure S4, S5A, Table ST5). GbM genes are enriched in core angiosperm orthogroups (multi-copy and single-copy) and depleted in non-core orthogroups. The only exceptions were the gbM-depleted species *B. rapa* and *E. salsugineum*. TeM genes have the opposite pattern and are depleted in core angiosperm orthogroups and enriched in non-core orthogroups, suggesting a more recent evolutionary origin for gene families enriched for teM. UnM genes are variably represented across orthogroups, but are more frequently enriched in cross-family (32/43 enriched, 4/43 depleted) and depleted in core:single-copy (2/43 enriched, 41/43 depleted) orthogroups.

We next tested enrichment or depletion of duplicate types in each orthogroup category (Figure S5B, Table ST6). SGDs are more frequently enriched in cross-family and family-specific orthogroups and also in core:multi-copy genes, except for proximal duplicates, which is the only duplication type depleted in core:multi-copy genes in most species (4/43 enriched, 29/43 depleted). Every duplication type was depleted in core:single-copy orthogroups with the exception of enrichment of dispersed duplicates in *G. max* and WGDs in *F. x ananassa, G.max, M. x domestica*, and *P. virgatum*. The greatest enrichment was in the extant polyploids *F. x ananassa* (octoploid) and *P. virgatum* (tetraploid), suggesting insufficient time since WGD to revert to singletons. WGDs were enriched in core:multi-copy orthogroups and depleted in noncore orthogroups with the exception of *C. melo* WGDs, which showed enrichment in cross-family orthogroups, and *S. tuberosum* WGDs, which is enriched in cross-family, family-specific, and species/lineage-specific orthogroups. As noted in the previous section, WGDs of these two species are depleted for gbM and enriched for teM. The enrichment of WGDs in non-core orthogroups for these species may explain why WGDs differ in their enrichment/depletion of genic methylation. *S. tuberosum* is also the only extant autopolyploid in our dataset and is of relatively recent origin (Consortium and The Potato Genome Sequencing Consortium, 2011; Wang et al., 2018), which could result in overrepresentation of more lineage-specific genes that are more likely to be teM. Collectively these results indicate that gene family composition is a driving factor in the relationship between gene duplication and genic methylation.

### Methylation divergence between paralogs

Changes in genic methylation might facilitate functional divergence between paralogs and mark different evolutionary trajectories, so we determined the extent of genic methylation differences between paralogs (Table ST7). WGD pairs have the highest similarity in genic methylation (same: ~69-97%, median – 84%; different: ~3-31%, median – 16%), followed by tandem (same: ~69%-93%, median – 82%; different: ~7-31%, median – 18%), proximal (same: ~66%-90%, median – 77%; different: ~10-34%, median – 23%), and dispersed (same: ~65%-92%, median – 76%; different: ~8-35%, median – 24%). Translocated duplicates had the broadest range and the greatest proportion of pairs differing in genic methylation (same: ~57%-90%, median – 74%; different: ~10-42%, median – 25%) (Figure S6, Table ST7). *A. trichopoda* is an outlier in this analysis due to the small number of genes classified as WGD or translocated. The direction of genic methylation changes cannot typically be discerned. However, for translocated duplicates one paralog is syntenic and considered parental locus and the translocated gene the daughter locus (Wang et al., 2013a). Translocated copies had higher teM proportions in 34/43 species and lower gbM proportions in 24/43 species (Table ST8). Assuming that parental locus methylation is the original state, we can determine the directionality of methylation changes in the translocated copy. Switching to unM was the most common in 22/43 species, switching to teM in 19/43 species, and switching to gbM in only 2/43 species (Table ST9). This shows that while the majority of pairs retain the same genic methylation status, changes are not infrequent, becoming more common as duplicates move to increasingly distal locations and that changes are predominantly to unM and teM, rarely to gbM.

Core:single-copy orthogroups are also duplicated during WGD, but are quickly eliminated by fractionation and are thus depleted in WGDs in nearly every species (Figure S5B, Table ST6). These ‘duplication-resistant’ pairs are dosage-sensitive and we hypothesized that DNA methylation has a role in maintaining dosage of these genes following duplication. Core:single-copy genes are predominantly gbM or unM, with few teM genes (Figure S4). GbM and unM core:single-copy genes differ in functional enrichment (Table ST10). GbM genes are enriched for cellular response to stress and metabolic processes, such as liposaccharide metabolism; while unM are enriched in processes like photosynthesis, cell redox homeostasis, and cell cycle checkpoint signaling. Although most core:single-copy genes are singletons (SC-singletons) within a species, duplicates still persist in varying numbers across angiosperms. We termed these ‘single-copy intermediates’ (SC-intermediates). SC-intermediate genes differed in their genic methylation at a much higher frequency than both core:multi-copy pairs and duplicate pairs as a whole (Figure 2B). SC-singletons are enriched for gbM genes compared to SC-intermediates (Figure 2B), while SC-intermediates have significantly more teM genes than SC-singletons. These results support our hypothesis and indicate a potential role for DNA methylation in maintaining dosage of single-copy core angiosperm genes following duplication and prior to fractionation.

**Figure 2:**
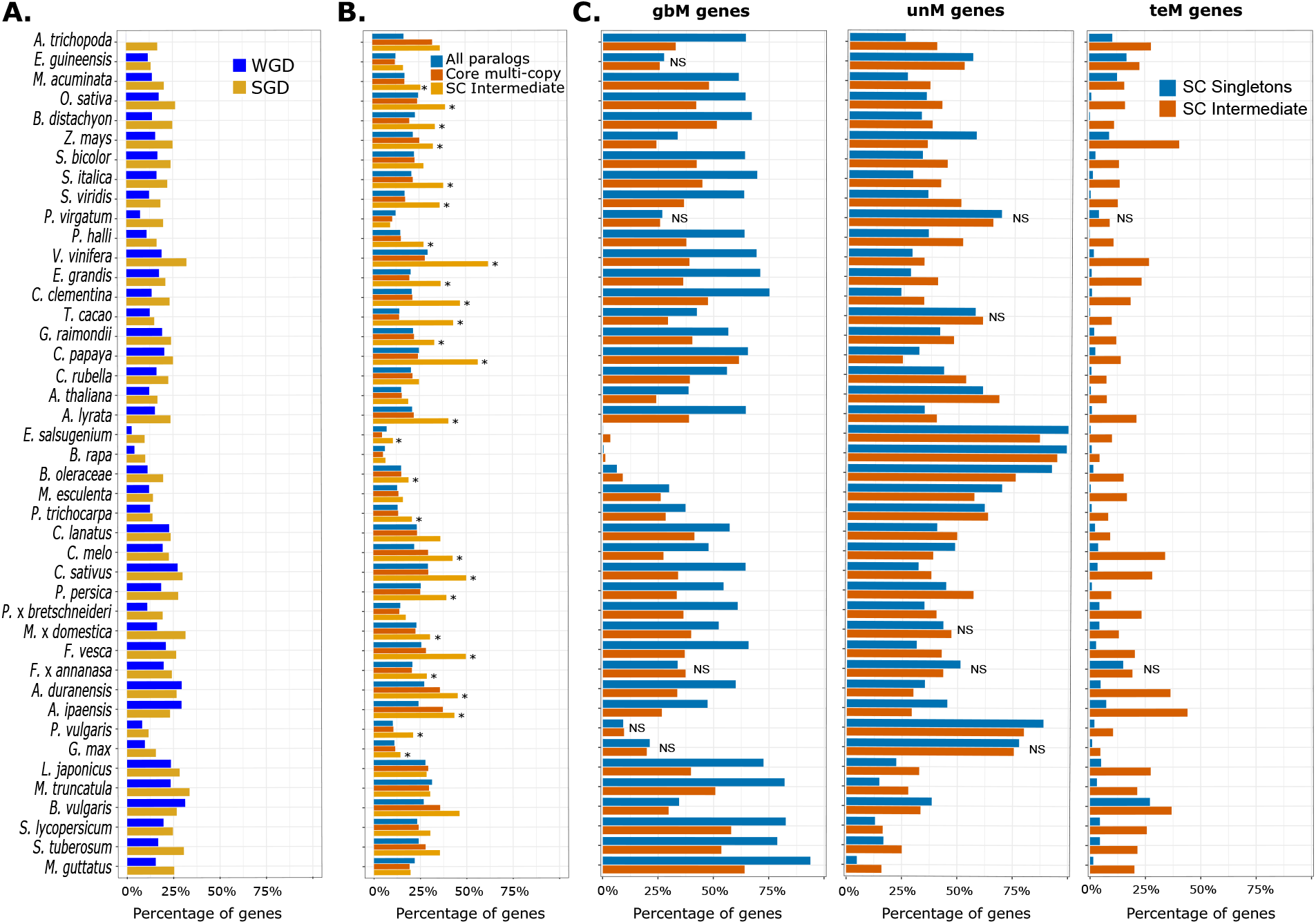
Genic DNA methylation patterns in duplicate paralogs and single-copy orthologs. **A)** Proportion of duplicate pairs with divergence in methylation between the two paralogs in whole-genome (dark blue) and single-gene duplicates (golden). **B)** Differences in proportion of duplicate paralogs with divergent methylation profiles in core: multi-copy (brown) and singlecopy intermediates (orange). The proportion of pairs with methylation divergence across all duplicate pairs are shown in light blue. Statistically significant differences are indicated by ‘*’ **C)** Proportions of gbM, unM, and teM in single-copy singletons (blue) and single-copy intermediates (dark orange). ‘NS’ represents comparision without statistically significant difference between SC-singletons and SC-intermediates.

### Genic methylation marks paralog age and sequence evolution

We next examined the relationship between genic methylation and paralog evolution. Synonymous substitutions (Ks) are assumed to accumulate neutrally with time and Ks distributions have been used to date duplication events (Lynch and Conery, 2000; Maere et al., 2005). No trend was expected for WGDs, as WGD duplicates all genes and given different WGD histories across species. However, WGD gbM-contaning pairs tended to have lower Ks values and unM-containing pairs higher Ks values (Figure 3A, Table ST11). No clear trend was observed for teM-containing WGD pairs. This may be due to the depletion and lower numbers of teM in WGDs. In contrast to WGD, SGD is a continuous process with constant gene birth and death (Lynch, 2002). SGD teM-containing pairs typically had lower Ks values than those with only gbM or unM paralogs (Figure 3B, Figure S7). This is most evident for teM-teM pairs, but gbM-teM and unM-teM also have lower Ks values. This suggests that teM paralogs tend to be evolutionarily younger. We confirmed this using a method independent of Ks for translocated genes. As the syntenic gene is assumed to be parental in translocated genes, the daughter gene can be parsed into different periods (epochs) at each node of the species tree (Table ST12) by sequential exclusion to the closest outgroup (Wang et al., 2013a). More recent translocated duplicates were enriched in teM paralogs, while more ancient translocated duplicates were enriched for gbM and unM paralogs (Figure S8, Table ST13), supporting our observations from the Ks analysis. These results also fit with the observation that evolutionarily younger lineage specific orthogroups are enriched for teM genes. We also compared Ks distributions for SC-intermediates and core:multi-copy genes, but observed no difference suggesting similar evolutionary ages (Figure S9).

**Figure 3:**
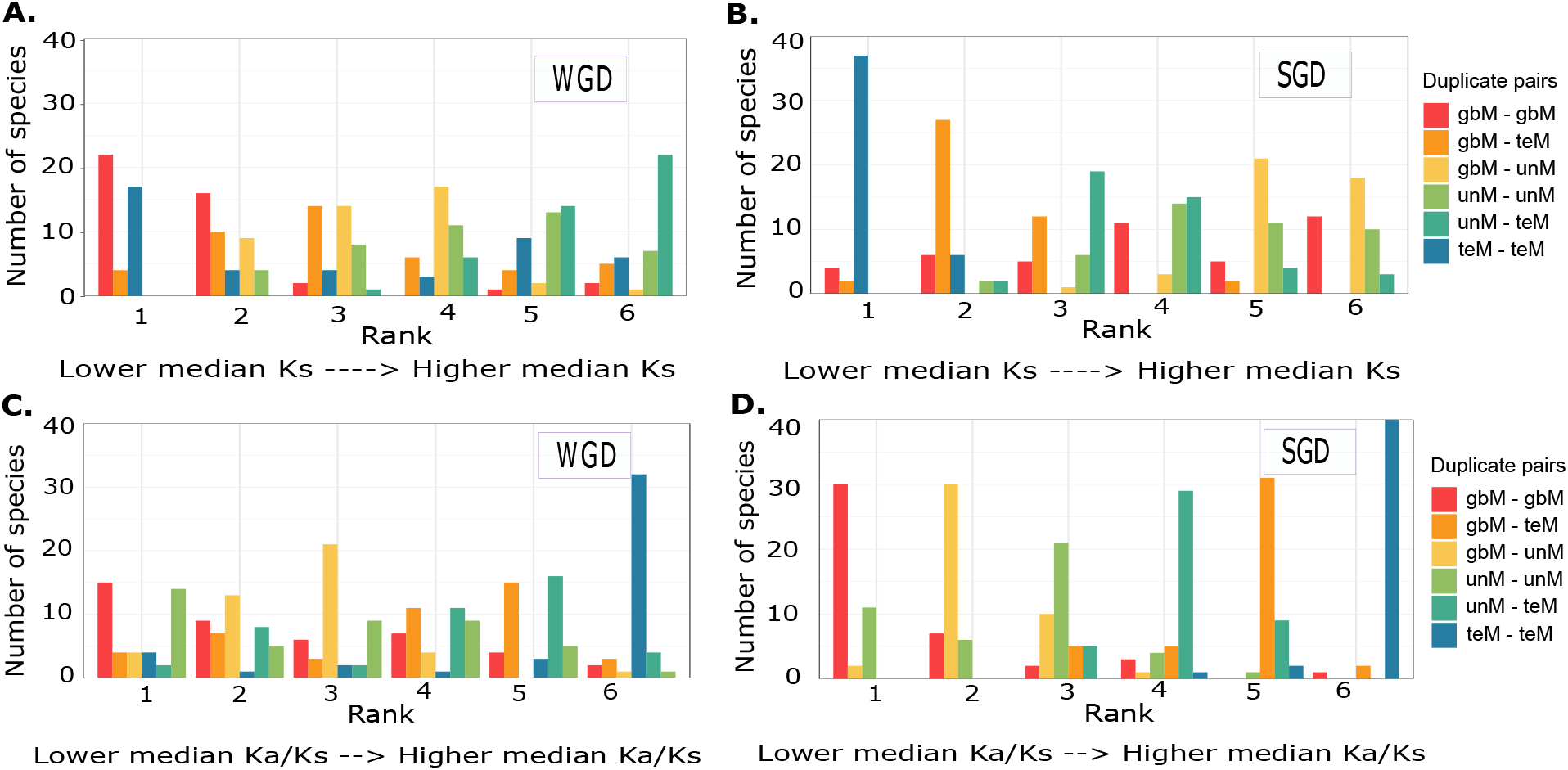
Relationship between genic methylation, age of duplication, and sequence evolution of duplicate paralogs. Bar plots showing the number of species in each of the duplicate-pair genic methylation classifications (gbM-gbM, gbM-teM, teM-teM, unM-unM, gbM-unM, and unM-teM) ranked based on median Ks values (synonymous substitutions) for whole-genome duplicates **(A)** and single-gene duplicates **(B)**. Bar plots showing the number of species in each of the duplicate-pair genic methylation classifications (gbM-gbM, gbM-teM, teM-teM, unM-unM, gbM-unM, and unM-teM) ranked based on median Ka/Ks values (ratio of Ka, non-synonymous substitutions to Ks, synonymous substitutions) for whole-genome **(C)** and single-gene duplicates **(D)**.

The ratio of non-synonymous (Ka) to synonymous (Ks) substitutions (Ka/Ks), is indicative of sequence evolution (Miyata and Yasunaga, 1980; Yang and Bielawski, 2000). A Ka/Ks < 1 is indicative of purifying selection, Ka/Ks = 0 indicates neutral selection, and Ka/Ks > 1 indicative of diversifying selection. We calculated Ka/Ks ratios for each duplicate pair and examined their distributions (Figure S10, Table ST14). The majority of pairs have a Ka/Ks < 1, regardless of duplication type or genic methylation, indicating purifying selection. However, there are differences in the distribution based on genic methylation. For both WGD and SGD genes (Figure 3C,D) gbM-containing pairs have lower Ka/Ks, teM-containing pairs have higher Ka/Ks, while unM-containing pairs are intermediate in distribution. This suggests that teM paralogs are under relaxed selective constraints compared to gbM and unM paralogs. A number of pairs had a Ka/Ks > 1 indicating diversifying selection. These were enriched for SGD and depleted for WGD in almost every species except *S. tuberosum* (Table ST15), and enriched in teM-containing pairs, in particular teM-teM pairs (Table ST16). We hypothesized that as SC-intermediates would be under relaxed selective constraints as these are enriched for teM. Indeed, SC-intermediates have higher Ka/Ks values compared to core:multi-copy pairs (Figure S11). Increased non-synonymous substitutions in SC-intermediates could lead to their pseudogenization and facilitate fractionation to singleton status.

Ongoing gene duplication and differential fractionation within a species can create presence-absence variation (PAVs). We used published lists of PAVs in *Brassica oleraceae, Zea mays, Solanum lycopersicum*, and *Solanum tuberosum (Hirsch et al., 2014; Golicz et al., 2016; Hardigan et al., 2016; Gao et al., 2019)* to examine the relationship between PAVs, genic methylation, and gene duplication (Figure S12; Table ST17). PAVs are enriched for teM genes in all four species and depleted for gbM in three species, except gbM-deficient *B. oleracea*. PAVs are depleted for unM genes in three species and enriched for unM in *S. lycopersicum*.

Results are identical whether tested for all genes or duplicated genes only. Association between PAVs and teM could result from targeting of lineage-specific SGDs or incomplete fractionation of teM duplicates in the population. An example of the latter would be fractionation of core:single-copy orthogroups following WGD. In all four species SC-intermediates had higher frequencies of PAV compared to core:multi-copy orthogroups and a higher frequency compared to all genes in *Z. mays* (Figure S13), supporting the hypothesis that teM silencing is an intermediate to fractionation for single-copy core angiosperm genes.

### Genic methylation and paralog expression

Divergent expression between paralogs is proposed to be the first step in functional diversification, enabling paralogs to subfunctionalize in expression, and increasing the odds of retention (Ohno, 1970; Ferris and Whitt, 1979; Li et al., 2005). We used gene expression atlases in *A. thaliana, G. max, Phaseolus vulgaris* and *Sorghum bicolor* (O’Rourke et al., 2014; Klepikova et al., 2016; McCormick et al., 2018; Wang et al., 2019) to explore the relationship between genic methylation and paralog expression. For each gene we calculated the expression specificity (τ), a measure of the number of conditions in which a gene is expressed (Yanai et al., 2005). The value of τ ranges from ‘0’ (broad expression) to ‘1’ (narrow expression). Typically, gbM genes have the broadest expression, teM the narrowest, while unM genes have a wide range expression specificities (Figure 4A, S14). Core angiosperm genes are more broadly expressed, expression specificity becoming narrower with increasing lineage specificity (Figure S15). This trend persists when broken down by genic methylation. GbM, unM, and teM genes become more narrowly expressed with increasing lineage specificity. By duplication type, WGDs have broader expression and SGDs narrower expression (Figure S16). Distal SGDs have broader expression than local SGDs, despite the fact that distal SGDs can place duplicates in new chromatin contexts and are more frequently enriched for teM. Similar to what was observed for orthogroups, differences between duplication types persist even when comparing genes of like genic methylation (Figure S16). WGD teM genes had overall broader expression than distal teM SGD teM genes, which had broader expression than local SGD teM genes. This same trend was observed for gbM and unM. Both orthogroup and duplication type therefore exerts an effect beyond the genic methylation and global differences between both orthogroups and duplication types are not fully explained by differential enrichment of genic methylation.

**Figure 4:**
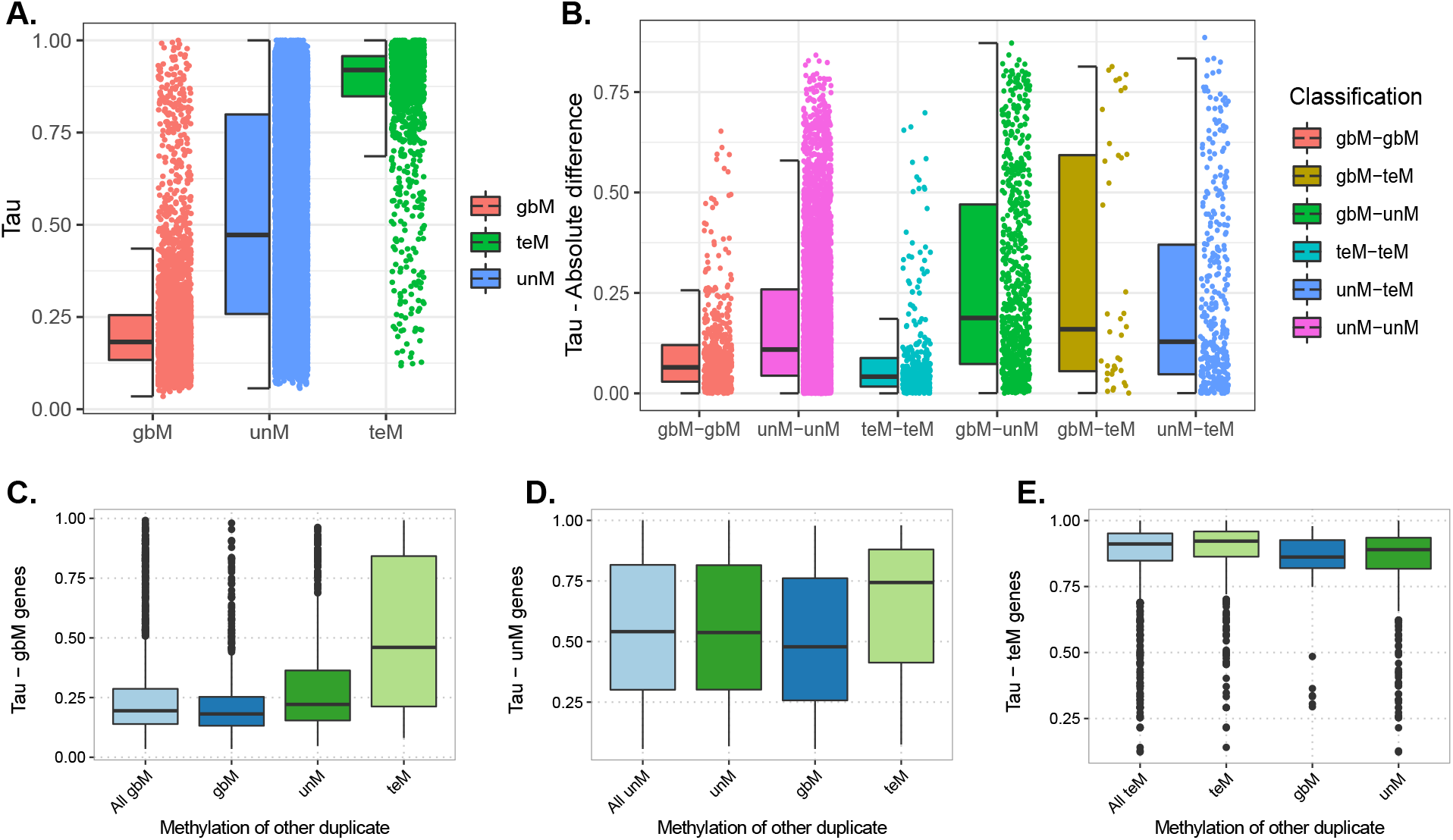
Gene expression specificity of A. thaliana duplicate gene pairs. Tissue-specificity index, Tau (τ), ranges from 0 (broadly expressed) to 1 (narrowly expressed). **(A)** Tissue-specificity of genes based on genic methylation classification (gbM, unM, and teM). **(B)** Absolute difference in tissue-specificity index (τ) between pairs of duplicate genes with similar or divergent methylation. Differences in Tau specificity of gbM, unM, and teM genes (**C, D,** and **E,** respectively) when the other duplicate pair has the same or a different genic methylation status. For example, for gbM genes, the tau specificity was plotted for all gbM genes and the gbM paralog in gbM-gbM, gbM-teM, and gbM-unM pairs. For unM genes, the tau of only the unM paralog is shown and similarly for teM genes, only the tau of the teM paralog is shown.

We next examined expression divergence between duplicate pairs, first calculating the expression correlation of each pair (Figure S17). Pairs with the same genic methylation had higher correlation than pairs differing in genic methylation, gbM-gbM pairs having the highest overall correlation. Fitting this, duplicate pairs differing in genic methylation have a greater overall absolute difference in expression specificity than pairs with the same genic methylation (Figure 4B, S18, Table ST18). Surprisingly, the expression specificity differed for genes of the same genic methylation based on the methylation of its duplicate pair (Figure 4C-E, S19). GbM genes that are part of gbM-gbM pairs tend to have broader expression specificity compared to the gbM genes in gbM-unM and gbM-teM pairs. For unM genes, those in unM-gbM pairs had broader expression specificity than those in unM-unM pairs and those in unM-teM pairs narrower expression specificity than either. Finally, teM genes in teM-gbM pairs had broader expression specificity than those in teM-unM pairs, which in turn had broader expression specificity than those in teM-teM pairs. This suggests a potential relationship between the parental locus expression and the expression of the duplicate copy. Perhaps certain genes may be predisposed to genic methylation changes by their expression.

### Transposons and chromatin environment associations

Non-CG methylation is often associated with TEs and TEs can alter the gene chromatin and expression (Hirsch and Springer, 2017; Raju et al., 2019). We identified TEs in or within 1 kb (Figure 5A, Table ST19) for each paralog. TeM paralogs are enriched (36/43 enriched, 4/43 depleted) and unM paralogs depleted (3/43 enriched, 34/43 depleted) for TEs in the majority of species. GbM was enriched (15/43) and depleted (15/43) for TEs in an equal number of species. Examining duplication type (Figure 5B, Table ST20), WGDs are depleted (2/43 enriched, 37/43 depleted) and all four SGDs enriched (Tandem: 30/43 enriched, 3/43 depleted; Proximal: 33/43 enriched, 2/43 depleted; Translocated: 21/43 enriched, 2/43 depleted; Dispersed: 27/43 enriched, 4/43 depleted) for TEs in the majority of species. Enrichment of TEs in SGDs may partly explain enrichment of teM in SGDs. We compared TE presence/absence for duplicate pairs differing in genic methylation (Table ST21), hypothesizing that the teM paralog would associate with TEs more frequently than its unM or gbM pair. This was true for *A. thaliana* and *A. lyrata*, but for most species both paralogs are associated with a TE in gbM-teM and unM-teM pairs, suggesting a more complex relationship than simple TE presence/absence in switching of genic methylation states. Differences in TE location or the TE family may be relevant, as was shown with TEs and heterochromatin spreading in *Z. mays* (Eichten et al., 2012).

**Figure 5:**
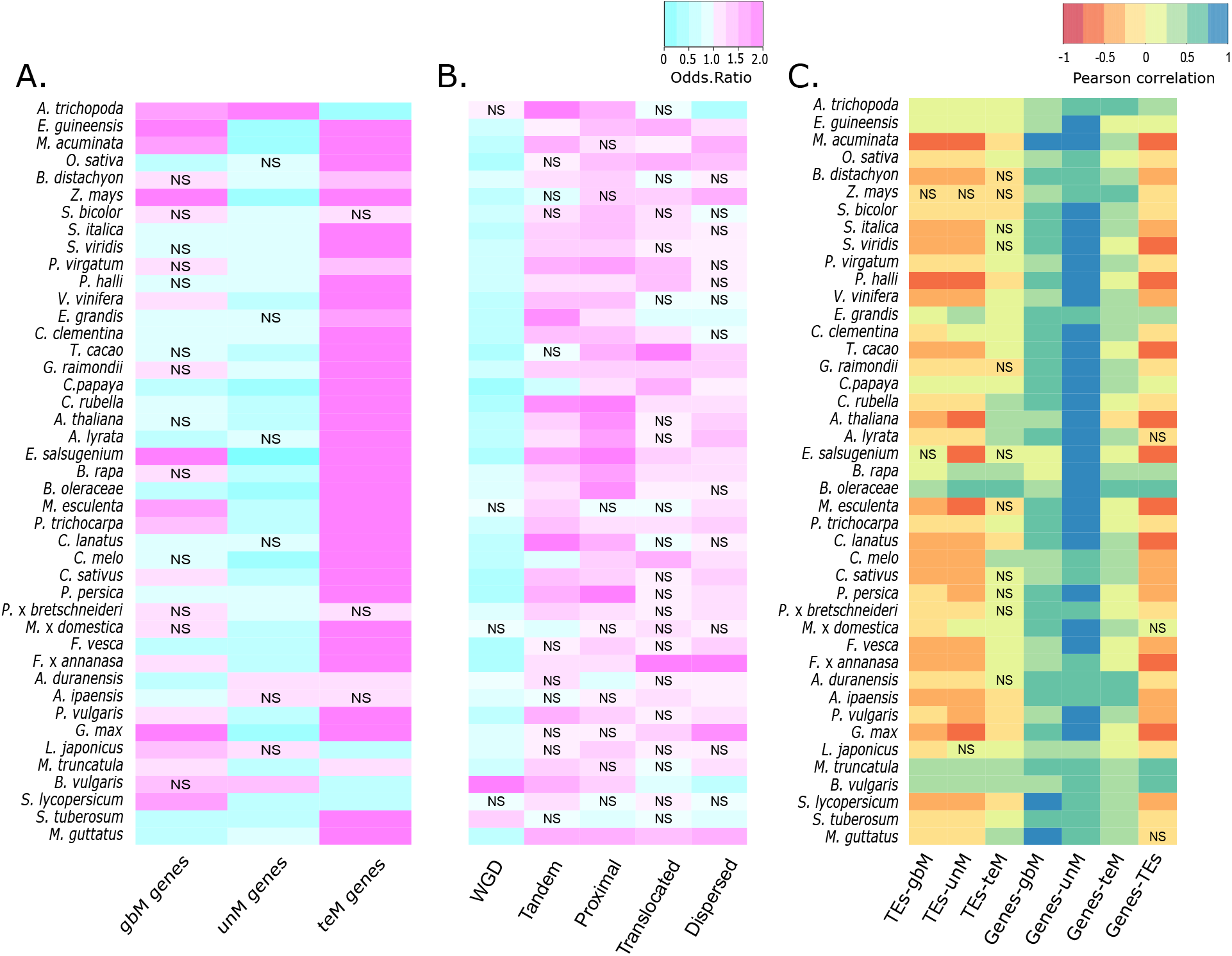
Local and genome-wide transposon and chromatin environment associations of duplicate genes. Enrichment and depletion of transposable elements (TEs) with gbM, teM, and unM paralogs **(A)** and different types of duplication **(B)** in each species. TEs within 1 kb upstream, downstream or within the gene body were considered associated with that gene. Fisher Exact test odds ratio of less than 1 represents depletion (represented in shades of cyan), greater than 1 indicates enrichment (represented in shades of magenta). Unless indicated, all associations are statistically significant at a FDR-corrected p-value < 0.05. ‘NS’ indicates no statistical significance. **C)** Genomic features such as number of genes and number of TEs were calculated in 100kb sliding windows with a 50kb step size. Increasing shades of blue indicate positive correlation, while increasing shades of red represent negative correlations. Unless indicated, all associations are statistically significant at a FDR-corrected p-value < 0.05. ‘NS’ indicates no statistical significance.

Location and chromatin environment can also affect genic methylation of duplicate genes. In *G. max*, translocation of paralogs to TE-rich pericentromeric regions often resulted in teM acquisition (El Baidouri et al., 2018). We used gene number, TE number, and fraction of TE-base pairs in sliding windows across the genome as a proxy for regions of euchromatin and heterochromatin and correlated these with the number of gbM/unM/teM paralogs (Figure 5C, Table ST22). GbM, unM, and teM duplicates are positively correlated with gene number, except *A. thaliana*, where teM has a very weak negative correlation (Pearson’s r = −0.05, FDR corrected p-value = 0.004). This may be due to *A. thaliana* genomic organization, which has the smallest genome and strongest negative correlation between total gene number and TEs (Pearson’s r = −0.71, FDR corrected p-value < 0.001) in our data. In most species the distribution of gbM and unM genes is negatively correlated (gbM: 8/43 positive, 33/43 negative; unM: 10/43 positive, 31/43 negative) and teM genes positively correlated with TEs (24/43 positive, 8/43 negative). This supports the hypothesis that duplication of genes to heterochromatic regions can lead to teM acquisition, however, this does not explain cases such as SC-intermediates.

### Epiallele frequency and paralog evolution within a population

DNA methylation varies across a population (Becker and Weigel, 2012). The relationship between this variation and paralog evolution is unknown. To address this, genes were classified based on genic methylation and binned according to the frequency of gbM/unM/teM across 928 *A. thaliana* accessions (Kawakatsu et al., 2016). We examined the proportion of duplication type (Figure 6A) and orthogroup (Figure S19) for each bin, predicting that there would be a corresponding change according to the frequency of genic methylation. This was true only in some instances. GbM showed a slight decrease in SGDs at higher frequencies, this being most evident for tandem SGDs, while tandem SGDs continually increased with unM frequency. WGD decreased and proximal SGDs increased with increasing teM frequency, while tandem SGDs peaked at ~25-50% before declining. As observed across species, a stronger association was observed for orthogroups. As the gbM frequency increases, the frequency of core:multi-copy orthogroups increases, and the frequency of cross-family, family-specific, and species/lineage-specific orthogroups decreases. We observed an opposite trend with increasing teM frequency. As the unM frequency increases in the population, we observe an increase in cross-family orthogroups and a decrease in the frequency of core single-copy genes.

**Figure 6:**
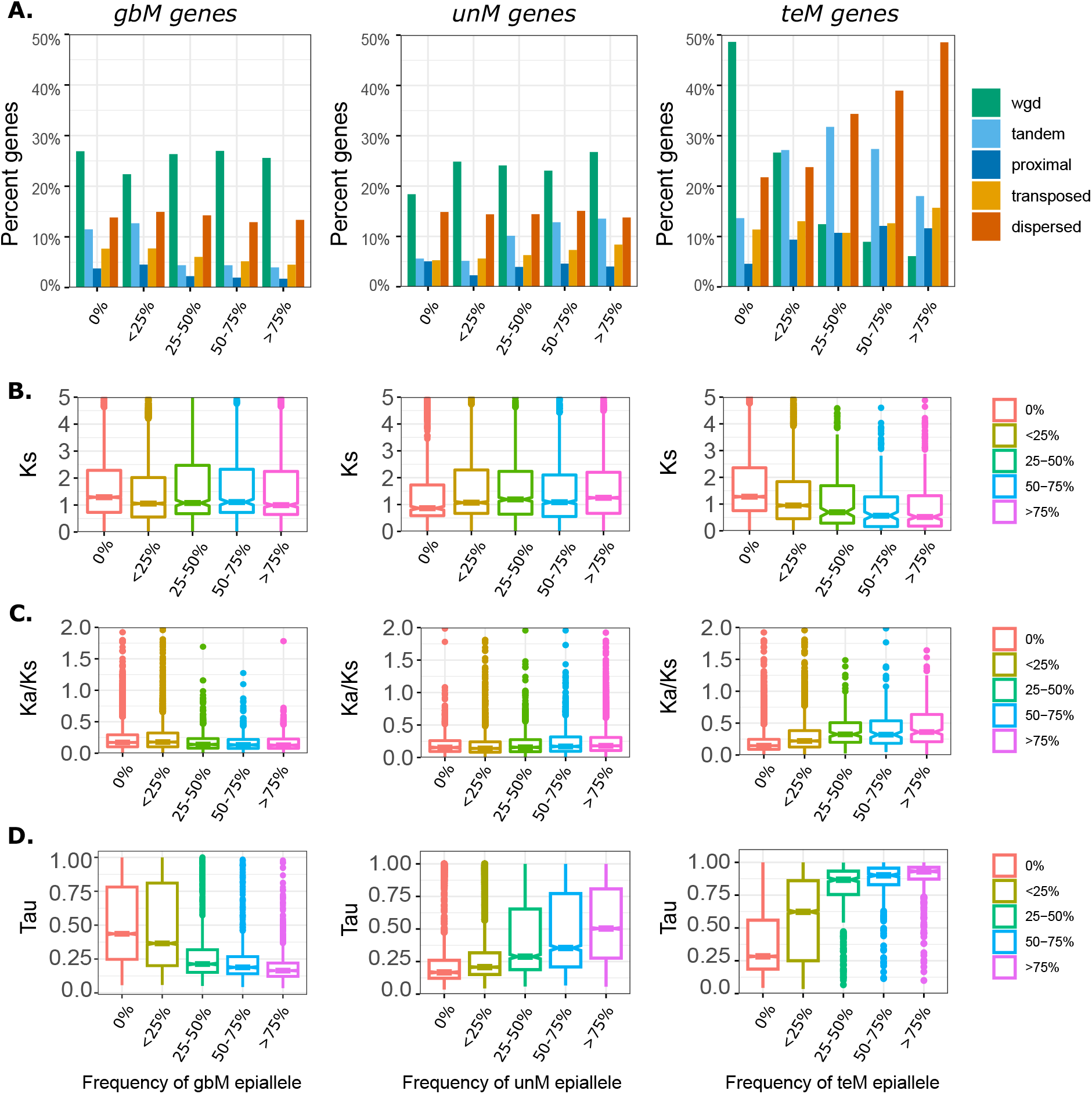
Genic methylation frequency within a population is associated with duplication type, age, sequence evolution, and gene expression specificity. **A)** Bar plots showing the the proportion of each type of duplication across different genic methylation frequency of gbM, unM, and teM (0%, <25%, 25%-50%, 50%-75%, >75%) in 928 A. thaliana acessions. **B-C)** Boxplots showing the relationshsip between genic methylation frequency of gbM, unM, and teM, age of duplication (Ks values (synonymous substitutions)) and Ka/Ks values (ratio of Ka, non-synonymous substitutions to Ks, synonymous substitutions). **D)** Boxplot showing expression specificities of genes in A. thaliana Col-0 at different frequencies of gbM, unM, and teM across 928 A.thaliana accessions.

We examined Ks (Figure 6B) and Ka/Ks (Figure 6C) at different epiallele frequencies. Both gbM and unM show little variation in Ks at different frequencies, while Ks steadily decreases with increasing teM frequency. This suggests a higher frequency of teM in evolutionarily more recent paralogs. Ka/Ks decreased with increasing gbM frequency and increased with increasing teM frequency, fitting the expectation of gbM genes being under greater purifying selection and teM genes being under relaxed selective constraints. Ka/Ks increased slightly with higher unM frequency. We hypothesized that this may be related to gene expression. Supporting this hypothesis, we observed that τ increased (more tissue specific) at higher population frequencies of both unM and teM, and decreased with increasing frequency of gbM (Figure 6D).

## Discussion

DNA methylation has been proposed to have a role in paralog evolution (Rodin and Riggs, 2003; Wang et al., 2013b; Keller and Yi, 2014; Wang et al., 2014a). However, this has not been examined at either a broad phylogenetic level or within a population, leaving the generalizability of results from individual species unresolved. To address this, we examined DNA methylation and paralog evolution across 43 angiosperms and a population of 928 *A. thaliana* accessions. Across the phylogeny WGDs are broadly enriched for gbM and unM genes, and depleted for teM genes. There is further differentiation between ‘local’ SGDs (tandem and proximal) and ‘distal’ SGDs (translocated and dispersed). Both are more frequently depleted in gbM, local duplicates are more frequently enriched for unM, while there is increasing frequency of teM from tandem to proximal to translocated to dispersed SGDs. There are notable exceptions to these trends. For three Brassicaceae species (*B. oleracea, B. rapa*, and *E. salsugineum*) divergence from these patterns is explained by a known depletion of gbM (Bewick et al., 2016). The Cucurbitaceae and *S. tuberosum* were also depleted of WGD gbM, despite no known depletion of gbM in these species and needs further investigation. We observe an even more consistent association of genic methylation with different types of orthogroups and this may drive patterns observed between genic methylation and gene duplication. This appears to be the case for *S. tuberosum*, a relatively recent autopolyploid (Consortium and The Potato Genome Sequencing Consortium, 2011). Unlike other species, *S. tuberosum* WGDs are enriched for increasingly lineage-specific orthogroups which explains the depletion of gbM and unM and enrichment of teM in WGDs.

GbM is characteristic of evolutionarily conserved genes (Takuno and Gaut, 2012; Takuno and Gaut, 2013; Bewick et al., 2016; Takuno et al., 2016). Fitting this, gbM is enriched in core angiosperm genes and gbM-gbM duplicate pairs are more conserved in sequence and expression. In contrast, teM genes are evolutionarily younger, increasing in enrichment with greater lineage-specificity, and predominantly found in recent SGDs. TeM paralogs have narrower expression, higher Ka/Ks ratios, and enrichment in PAV, suggesting that most are on the path to pseudogenization. This process would lead to their depletion in more ancient WGDs, while continual duplication in SGDs would result in their enrichment. UnM genes are seemingly intermediate between gbM and teM in most aspects. UnM might be considered the ‘default’ state and spans from more gbM-like to more teM-like genes. In gbM-depleted species, the gbM ortholog is unM (Bewick et al., 2016). UnM is the largest group and broadly represented across both core angiosperm orthogroups and more lineage-specific orthogroups. Many transcription factors and kinases are retained following WGD and have tissue-specific expression characteristic of unM (Pophaly and Tellier, 2015). At the same time, many tandem and proximal duplications are associated with environmental adaptation (Freeling, 2009). This would favor retention of unM in both WGDs and local SGDs. Unexpectedly, unM-containing WGD pairs typically have a higher Ks than gbM-containing pairs. We speculate that this could result from unM-containing pairs being derived from more ancient WGD events or differences in the mutation rates of gbM and unM genes.

Within a population, paralog evolution associates with genic methylation frequency. This is especially true for teM genes, where increasing teM frequency associates with evolutionary younger genes, narrower expression, and greater sequence divergence. Differences are also observed in gbM and unM genes and appear to be driven by expression as more narrowly expressed unM genes have a higher Ka/Ks ratio. To achieve high-frequency in a population, the simplest explanation is that a genic methylation state was established early following duplication and spread with expansion of the population. Low frequency states would reflect either relict populations (Kawakatsu et al., 2016) or cases of recent acquisition. A deeper analysis taking account the structure and relationship accessions in the population will provide further insight into the establishment of genic methylation states and paralog evolution.

Gene family is also an important factor, as indicated by our orthogroup analyses and should be accounted for when trying to understand the role of DNA methylation in paralog evolution. Gene families differ in susceptibility to fractionation, some being preferentially retained, while others convergently revert to singletons. Both are thought to be dosage sensitive. The former retains duplicate copies to maintain relative dosage to other genes in the genome as explained by the gene balance hypothesis (Birchler and Veitia, 2010) and are characteristic of what we termed ‘core:multi-copy’ genes. The latter ‘duplication-resistant’ genes we have termed ‘core:single-copy’ and are thought to be under selective pressure to maintain singleton status (Paterson et al., 2006; De Smet et al., 2013; Li et al., 2016). GbM and unM mark different functional sets of core:single-copy genes and likely reflect differences in expression. While core:single-copy genes are predominantly singletons across angiosperms, in each species some duplicate copies still persist (SC-intermediates). By contrasting these SC-intermediates to core:multi-copy genes, we found that SC-intermediates have more frequent differences in genic methylation between duplicate pairs and a higher frequency of teM compared to core:multi-copy genes and often duplicates pairs as a whole. SC-intermediates are predominantly syntenic genes resulting from WGD and do not differ in evolutionary age from core:multi-copy genes. As such, they are unlikely to have been silenced prior to WGD and the gain of teM would not have occurred via movement to heterochromatic regions. Despite being from conserved gene families, SC-intermediates have higher Ka/Ks ratios indicating relaxed selection and are associated with presence-absence variation. We propose that SC-intermediates are in the process of fractionation and that silencing by DNA methylation has a role in maintaining dosage and is a first step to their fractionation. However, it is unclear how these conserved genes would become silenced. Experimental approaches, including use of resynthesized or synthetic polyploids (Edger et al., 2017) may be necessary to capture this process.

It has been proposed that silencing by DNA methylation can result in retention of paralogs and their functional divergence (e.g. epigenetic complementation) (Adams et al., 2003; Rodin and Riggs, 2003; Chang and Liao, 2012). Alternatively, silencing may lead to pseudogenization and gene loss (Hua et al., 2013; El Baidouri et al., 2018). Neither hypothesis is necessarily wrong and cases of both likely exist within a genome. Our results suggest that pseudogenization and loss is the predominant consequence. This is most evident for SGDs and core:single-copy angiosperm genes. However, there is suggestive evidence for epigenetic complementation in the divergence of expression states and many teM genes are still expressed under limited conditions. Epigenetic complementation could also occur through silencing by other chromatin marks, like H3K27me3. Furthermore, many teM containing duplicates have a Ka/Ks > 1, possibly indicating positive selection. Rapid functional divergence of SGDs was observed in grasses; many of these have characteristics similar to teM SGDs (Jiang and Assis, 2019), but this will require a further analysis. Our data shows the genic DNA methylation marks differing evolutionary fates of duplicate genes and may have a role in maintaining dosage following gene duplication.

## Methods

### Genome and Methylome data

We used genomes and annotations for 58 angiosperm species (Tuskan et al., 2006; Jaillon et al., 2007; Ming et al., 2008; Sato et al., 2008; Initiative and The International Brachypodium Initiative, 2010; Schmutz et al., 2010; Hu et al., 2011; Bennetzen et al., 2012; D’Hont et al., 2012; Garcia-Mas et al., 2012; Lamesch et al., 2012; Paterson et al., 2012; Amborella Genome Project, 2013; Guo et al., 2013; Hellsten et al., 2013; Kawahara et al., 2013; Ming et al., 2013; Motamayor et al., 2013; Sharma et al., 2013; Singh et al., 2013; Slotte et al., 2013; Yang et al., 2013; Dohm et al., 2014; Liu et al., 2014; Parkin et al., 2014; Schmutz et al., 2014; Tang et al., 2014; Wang et al., 2014b; Bartholomé et al., 2015; VanBuren et al., 2015; Bertioli et al., 2016; Bombarely et al., 2016; Bredeson et al., 2016; Cheng et al., 2017; Daccord et al., 2017; Harkess et al., 2017; Jiao et al., 2017; Verde et al., 2017; Xu et al., 2017; Edger et al., 2018; Filiault et al., 2018; Hibrand Saint-Oyant et al., 2018; Hulse-Kemp et al., 2018; Lovell et al., 2018; McCormick et al., 2018; VanBuren et al., 2018; Wu et al., 2018; Xue et al., 2018; Barchi et al., 2019; Colle et al., 2019; Edger et al., 2019; Li et al., 2019; Valliyodan et al., 2019; Lovell et al., 2021; Hosmani et al.; Mamidi et al.), including 43 species (Table S1) with whole-genome bisulfite sequencing (WGBS) data (Amborella Genome Project, 2013; Seymour et al., 2014; Kim et al., 2015; Ong-Abdullah et al., 2015; Secco et al., 2015; Bertioli et al., 2016; Bewick et al., 2016; Niederhuth et al., 2016; Daccord et al., 2017; Dong et al., 2017; Picard and Gehring, 2017; Song et al., 2017; Turco et al., 2017; Lü et al., 2018; Wang et al., 2018; Cheng et al., 2018b; Noshay et al., 2019; Yang et al., 2019) and an additional 15 species included as outgroups (Table S1). Genes were filtered to remove putative mis-annotated transposons as previously described (Bowman et al., 2017) with slight modifications. First, genes were searched against Pfam-A using hmmscan (Potter et al., 2018) filtering genes matching a curated list of TE domains (https://github.com/Childs-Lab/GC_specific_MAKER) with an e-value < 1e-5. Next, genes were searched against a set of transposase sequences (www.hrt.msu.edu/uploads/535/78637/Tpases020812.gz) using DIAMOND blastp (Buchfink et al., 2015) and hits with a e-value < 1e-10 removed.

### DNA methylation analyses

Whole-genome bisulfite sequencing (WGBS) from 43 angiosperm species (Table S1) were mapped to their respective genomes using methylpy v1.2.9 (Schultz et al., 2015). Genes were classified as gene-body methylated (gbM), TE-like methylated (teM), and unmethylated (unM) as previously done with slight modification (Takuno and Gaut, 2012; Niederhuth et al., 2016). First, a background rate was calculated for CG, CHG, CHH, and non-CG (combined CHG & CHH) methylation by averaging the percentage of methylated sites in that context across primary transcript coding regions (CDS feature) of all species. Each gene was tested for enrichment of CG, CHG, CHH, or non-CG in its primary transcript coding region against this background rate using a binomial test and p-values corrected for false discovery rate (FDR) by the Benjamini-Hochberg (BH) procedure (Benjamini and Hochberg, 1995). Genes enriched for CG methylation with ≥ 10 CG sites and non-significant CHG or CHH methylation were classified as gbM. Genes enriched for CHG, CHH, or non-CG and ≥ 10 sites in that context were classified as teM. Genes with ≤ 1 methylated site in any context or a weighted methylation (Schultz et al., 2012) ≤ 2% for all contexts (CG, CHG, or CHH) were classified as unM. Genes lacking DNA methylation data were classified ‘missing’ and those with intermediate DNA methylation levels not fitting the above criteria ‘unclassified’.

### Gene duplication classification

Each species was blasted against itself and *A. trichopoda* (outgroup) using DIAMOND blastp (Buchfink et al., 2015). *A. thaliana* was used as the outgroup for *A. trichopoda*. Hits from the same orthogroup and an e-value cutoff < 1e-5. were retained. Paralogs were classified by *DupGen_finder-unique (Qiao et al., 2019)*, requiring ≥ 5 genes for collinearity and ≤ 10 intervening genes to classify as ‘proximal’ duplicates. *MCScanX-transposed (Wang et al., 2013a)* was used to detect translocated duplicates at different epochs since species divergence (Table S12). Genic methylation enrichment in duplication types was determined by a two-sided Fisher’s exact test (Fisher, 1934) with FDR-correction by BH and plotted using *heatmap.2* in *gplots* (2020). The phylogenetic tree was generated using ‘V.PhyloMaker’ (Jin and Qian, 2019) and ‘phytools’ (Revell, 2012). To avoid overcounting, if a gene had more than one potential paralog, we retained the pair with the lowest e-value.

### Orthogroup analyses

Orthogroups were identified for protein sequences of 58 angiosperm species (Table S1) using Orthofinder v2.5.2 (Emms and Kelly, 2015; Emms and Kelly, 2019), with the options ‘-M dendroblast -S diamond_ultra_sens, -I 1.3’. Orthogroups represented in ≥51 species (~87.9%, Figure S3) were classified as “core angiosperm” orthogroups. This accounts for missing annotations and is equivalent to cutoffs in past work (Li et al., 2016). Following Li et al., we classified core angiosperm orthogroups as core:single-copy if represented by a single gene in ≥ 70% species and the remainder as core:multi-copy. Remaining orthogroups were classified based on increasing lineage-specificity: ‘cross family’ if present in multiple plant families, ‘family-specific’ if found in a single plant family, or ‘species/lineage specific’ if limited to a single species. Within each species, a subset of core:single-copy orthogroups still retained duplicate copies and were classified as single-copy intermediates (SC-intermediates) and those represented by only a single gene as single-copy singletons (SC-singletons). A two-proportion Z-test was used to test differences in genic methylation between SC-intermediates and SC-singletons.

### Sequence evolution

The calculate_Ka_Ks_pipeline.pl (Qiao et al., 2019) was used to determine nonsynonymous (Ka) and synonymous substitutions (Ks) for duplicate pairs. Protein sequences are aligned by MAFFT (v7.402) (Katoh and Standley, 2013), converted to a codon alignment with PAL2NAL (Suyama et al., 2006), and KaKs_Calculator 2.0 used to calculate Ka, Ks, and Ka/Ks with the γ-MYN method (Wang et al., 2010; Qiao et al., 2019). PAV variants were downloaded for *B. oleracea (Golicz et al., 2016), S. lycopersicum (Gao et al., 2019), S. tuberosum (Hardigan et al., 2016)*, and *Z. mays (Hirsch et al., 2014)*. For *S. tuberosum* and *Z. mays*, genes with an average read coverage of < 0.2 in ≥ 1 accession were considered PAV. Enrichment was tested using a twosided Fisher’s Exact test with FDR-correction by BH.

### Gene expression

Expression data for *A. thaliana, G. max, P. vulgaris*, and *S.bicolor* are from published expression atlases (O’Rourke et al., 2014; Klepikova et al., 2016; McCormick et al., 2018; Wang et al., 2019). *A. thaliana* reads were downloaded from NCBI SRA (PRJNA314076 and PRJNA324514), mapped with STAR (Dobin et al., 2013), and normalized for library size in DESeq2 (Love et al., 2014). *G. max, P. vulgaris*, and *S.bicolor* normalized data was downloaded from Phytozome (Goodstein et al., 2012). The tissue-specificity index (τ) was calculated in R for each gene as previously described (Yanai et al., 2005). Genes not expressed under any conditions were excluded as τ could not be calculated. Pearson correlation coefficients were calculated for each duplicate pair in R.

### Transposons and genomic distribution

TEs were annotated *de novo* for all species using EDTA (Ou et al., 2019). We calculated the total number of genes, genes belonging to each of the genic methylation classes, the number of TEs, and number of TE base pairs in 100 kb sliding windows with 50 kb steps. Pearson correlation coefficients were calculated using the *‘rcorr’* function in *‘corrplot’* (Wei and Viliam).

### Arabidopsis diversity

WGBS data for 928 *Arabidopsis thaliana* accessions, previously aligned by methylpy (Kawakatsu et al., 2016), was downloaded from the Gene Expression Omnibus (GEO Accession GSE43857). Genes were classified as before and the frequency of gbM/unM/teM in the population calculated for each gene.

## Data availability and research reproducibility

Raw data sources are listed in Table-S1. Formatted genomes and data are available at DataDryad XX. Code is available at: https://github.com/niederhuth/DNA-methylation-signatures-of-duplicate-gene-evolution-in-angiosperms.

## Acknowledgements

We thank Dr. Patrick Edger for the unpublished *C. violaceae* genome and Dr. Leslie Kollar for comments. This work was supported by Michigan State University, the USDA National Institute of Food and Agriculture Hatch Funds (MICL02572), and the National Science Foundation (gIOS-2029959). S. Marshall Ledford was supported by the Plant Genomics@MSU REU (DBI-1757043).

## Author contributions

S.K.K.R and C.E.N designed the work and analyses. S.K.K.R, C.E.N. and S.M.L performed data analysis. S.K.K.R and C.E.N wrote and edited the manuscript. All authors read and approved the final manuscript.

## References

Adams KL, Cronn R, Percifield R, Wendel JF (2003) Genes duplicated by polyploidy show unequal contributions to the transcriptome and organ-specific reciprocal silencing. Proceedings of the National Academy of Sciences 100: 4649–4654

Amborella Genome Project (2013) The Amborella genome and the evolution of flowering plants. Science 342: 1241089

Barchi L, Pietrella M, Venturini L, Minio A, Toppino L, Acquadro A, Andolfo G, Aprea G, Avanzato C, Bassolino L, et al (2019) A chromosome-anchored eggplant genome sequence reveals key events in Solanaceae evolution. Sci Rep 9: 11769

Bartholomé J, Mandrou E, Mabiala A, Jenkins J, Nabihoudine I, Klopp C, Schmutz J, Plomion C, Gion J-M (2015) High-resolution genetic maps of Eucalyptus improve Eucalyptus grandis genome assembly. New Phytol 206: 1283–1296

Becker C, Weigel D (2012) Epigenetic variation: origin and transgenerational inheritance. Curr Opin Plant Biol 15: 562–567

Benjamini Y, Hochberg Y (1995) Controlling the False Discovery Rate: A Practical and Powerful Approach to Multiple Testing. Journal of the Royal Statistical Society: Series B (Methodological) 57: 289–300

Bennetzen JL, Schmutz J, Wang H, Percifield R, Hawkins J, Pontaroli AC, Estep M, Feng L, Vaughn JN, Grimwood J, et al (2012) Reference genome sequence of the model plant Setaria. Nat Biotechnol 30: 555–561

Bertioli DJ, Cannon SB, Froenicke L, Huang G, Farmer AD, Cannon EKS, Liu X, Gao D, Clevenger J, Dash S, et al (2016) The genome sequences of Arachis duranensis and Arachis ipaensis, the diploid ancestors of cultivated peanut. Nat Genet 48: 438–446

Bewick AJ, Ji L, Niederhuth CE, Willing E-M, Hofmeister BT, Shi X, Wang L, Lu Z, Rohr NA, Hartwig B, et al (2016) On the origin and evolutionary consequences of gene body DNA methylation. Proc Natl Acad Sci U S A 113: 9111–9116

Birchler JA, Veitia RA (2010) The gene balance hypothesis: implications for gene regulation, quantitative traits and evolution. New Phytol 186: 54–62

Bombarely A, Moser M, Amrad A, Bao M, Bapaume L, Barry CS, Bliek M, Boersma MR, Borghi L, Bruggmann R, et al (2016) Insight into the evolution of the Solanaceae from the parental genomes of Petunia hybrida. Nat Plants 2: 16074

Bowman MJ, Pulman JA, Liu TL, Childs KL (2017) A modified GC-specific MAKER gene annotation method reveals improved and novel gene predictions of high and low GC content in Oryza sativa. BMC Bioinformatics 18: 522

Bredeson JV, Lyons JB, Prochnik SE, Wu GA, Ha CM, Edsinger-Gonzales E, Grimwood J, Schmutz J, Rabbi IY, Egesi C, et al (2016) Sequencing wild and cultivated cassava and related species reveals extensive interspecific hybridization and genetic diversity. Nat Biotechnol 34: 562–570

Bridges CB (1935) SALIVARY CHROMOSOME MAPS. Journal of Heredity 26: 60–64

Buchfink B, Xie C, Huson DH (2015) Fast and sensitive protein alignment using DIAMOND. Nature Methods 12: 59–60

Chang AY-F, Liao B-Y (2012) DNA methylation rebalances gene dosage after mammalian gene duplications. Mol Biol Evol 29: 133–144

Cheng C-Y, Krishnakumar V, Chan AP, Thibaud-Nissen F, Schobel S, Town CD (2017) Araport11: a complete reannotation of the Arabidopsis thaliana reference genome. Plant J 89:789–804

Cheng F, Wu J, Cai X, Liang J, Freeling M, Wang X (2018a) Gene retention, fractionation and subgenome differences in polyploid plants. Nat Plants 4: 258–268

Cheng J, Niu Q, Zhang B, Chen K, Yang R, Zhu J-K, Zhang Y, Lang Z (2018b) Downregulation of RdDM during strawberry fruit ripening. Genome Biol 19: 212

Colle M, Leisner CP, Wai CM, Ou S, Bird KA, Wang J, Wisecaver JH, Yocca AE, Alger EI, Tang H, et al (2019) Haplotype-phased genome and evolution of phytonutrient pathways of tetraploid blueberry. Gigascience. doi: 10.1093/gigascience/giz012

Conant GC, Birchler JA, Pires JC (2014) Dosage, duplication, and diploidization: clarifying the interplay of multiple models for duplicate gene evolution over time. Curr Opin Plant Biol 19: 91–98

Consortium TPGS, The Potato Genome Sequencing Consortium (2011) Genome sequence and analysis of the tuber crop potato. Nature 475: 189–195

Cusack BP, Wolfe KH (2007) Not born equal: increased rate asymmetry in relocated and retrotransposed rodent gene duplicates. Mol Biol Evol 24: 679–686

Daccord N, Celton J-M, Linsmith G, Becker C, Choisne N, Schijlen E, van de Geest H, Bianco L, Micheletti D, Velasco R, et al (2017) High-quality de novo assembly of the apple genome and methylome dynamics of early fruit development. Nat Genet 49: 1099–1106

De Smet R, Adams KL, Vandepoele K, Van Montagu MCE, Maere S, Van de Peer Y (2013) Convergent gene loss following gene and genome duplications creates single-copy families in flowering plants. Proc Natl Acad Sci U S A 110: 2898–2903

D’Hont A, Denoeud F, Aury J-M, Baurens F-C, Carreel F, Garsmeur O, Noel B, Bocs S, Droc G, Rouard M, et al (2012) The banana (Musa acuminata) genome and the evolution of monocotyledonous plants. Nature 488: 213–217

Dobin A, Davis CA, Schlesinger F, Drenkow J, Zaleski C, Jha S, Batut P, Chaisson M, Gingeras TR (2013) STAR: ultrafast universal RNA-seq aligner. Bioinformatics 29: 15–21

Dohm JC, Minoche AE, Holtgräwe D, Capella-Gutiérrez S, Zakrzewski F, Tafer H, Rupp O, Sörensen TR, Stracke R, Reinhardt R, et al (2014) The genome of the recently domesticated crop plant sugar beet (Beta vulgaris). Nature 505: 546–549

Dong P, Tu X, Chu P-Y, Lü P, Zhu N, Grierson D, Du B, Li P, Zhong S (2017) 3D Chromatin Architecture of Large Plant Genomes Determined by Local A/B Compartments. Molecular Plant 10: 1497–1509

Edger PP, Poorten TJ, VanBuren R, Hardigan MA, Colle M, McKain MR, Smith RD, Teresi SJ, Nelson ADL, Wai CM, et al (2019) Origin and evolution of the octoploid strawberry genome. Nat Genet 51: 541–547

Edger PP, Smith R, McKain MR, Cooley AM, Vallejo-Marin M, Yuan Y, Bewick AJ, Ji L, Platts AE, Bowman MJ, et al (2017) Subgenome Dominance in an Interspecific Hybrid, Synthetic Allopolyploid, and a 140-Year-Old Naturally Established Neo-Allopolyploid Monkeyflower. The Plant Cell 29: 2150–2167

Edger PP, VanBuren R, Colle M, Poorten TJ, Wai CM, Niederhuth CE, Alger EI, Ou S, Acharya CB, Wang J, et al (2018) Single-molecule sequencing and optical mapping yields an improved genome of woodland strawberry (Fragaria vesca) with chromosome-scale contiguity. Gigascience 7: 1–7

Eichten SR, Ellis NA, Makarevitch I, Yeh C-T, Gent JI, Guo L, McGinnis KM, Zhang X, Schnable PS, Vaughn MW, et al (2012) Spreading of heterochromatin is limited to specific families of maize retrotransposons. PLoS Genet 8: e1003127

El Baidouri M, Kim KD, Abernathy B, Li Y-H, Qiu L-J, Jackson SA (2018) Genic C-Methylation in Soybean Is Associated with Gene Paralogs Relocated to Transposable Element-Rich Pericentromeres. Mol Plant 11: 485–495

Emms DM, Kelly S (2019) OrthoFinder: phylogenetic orthology inference for comparative genomics. Genome Biol 20: 238

Emms DM, Kelly S (2015) OrthoFinder: solving fundamental biases in whole genome comparisons dramatically improves orthogroup inference accuracy. Genome Biol 16: 157

Feng S, Cokus SJ, Zhang X, Chen P-Y, Bostick M, Goll MG, Hetzel J, Jain J, Strauss SH, Halpern ME, et al (2010) Conservation and divergence of methylation patterning in plants and animals. Proc Natl Acad Sci U S A 107: 8689–8694

Ferris SD, Whitt GS (1979) Evolution of the differential regulation of duplicate genes after polyploidization. J Mol Evol 12: 267–317

Filiault DL, Ballerini ES, Mandáková T, Aköz G, Derieg NJ, Schmutz J, Jenkins J, Grimwood J, Shu S, Hayes RD, et al (2018) The genome provides insight into adaptive radiation and reveals an extraordinarily polymorphic chromosome with a unique history. Elife. doi: 10.7554/eLife.36426

Fisher SRA (1934) Statistical Methods for Research Workers.

Flagel LE, Wendel JF (2009) Gene duplication and evolutionary novelty in plants. New Phytologist 183: 557–564

Freeling M (2009) Bias in Plant Gene Content Following Different Sorts of Duplication: Tandem, Whole-Genome, Segmental, or by Transposition. Annual Review of Plant Biology 60: 433–453

Freeling M, Lyons E, Pedersen B, Alam M, Ming R, Lisch D (2008) Many or most genes in Arabidopsis transposed after the origin of the order Brassicales. Genome Research 18: 1924–1937

Ganko EW, Meyers BC, Vision TJ (2007) Divergence in expression between duplicated genes in Arabidopsis. Mol Biol Evol 24: 2298–2309

Gao L, Gonda I, Sun H, Ma Q, Bao K, Tieman DM, Burzynski-Chang EA, Fish TL, Stromberg KA, Sacks GL, et al (2019) The tomato pan-genome uncovers new genes and a rare allele regulating fruit flavor. Nat Genet 51: 1044–1051

Garcia-Mas J, Benjak A, Sanseverino W, Bourgeois M, Mir G, González VM, Hénaff E, Câmara F, Cozzuto L, Lowy E, et al (2012) The genome of melon (Cucumis melo L.). Proc Natl Acad Sci U S A 109: 11872–11877

Golicz AA, Bayer PE, Barker GC, Edger PP, Kim H, Martinez PA, Chan CKK, Severn-Ellis A, McCombie WR, Parkin IAP, et al (2016) The pangenome of an agronomically important crop plant Brassica oleracea. Nat Commun 7: 13390

Goodstein DM, Shu S, Howson R, Neupane R, Hayes RD, Fazo J, Mitros T, Dirks W, Hellsten U, Putnam N, et al (2012) Phytozome: a comparative platform for green plant genomics. Nucleic Acids Res 40: D1178–86

Guo S, Zhang J, Sun H, Salse J, Lucas WJ, Zhang H, Zheng Y, Mao L, Ren Y, Wang Z, et al (2013) The draft genome of watermelon (Citrullus lanatus) and resequencing of 20 diverse accessions. Nat Genet 45: 51–58

Hardigan MA, Crisovan E, Hamilton JP, Kim J, Laimbeer P, Leisner CP, Manrique-Carpintero NC, Newton L, Pham GM, Vaillancourt B, et al (2016) Genome Reduction Uncovers a Large Dispensable Genome and Adaptive Role for Copy Number Variation in Asexually Propagated Solanum tuberosum. Plant Cell 28: 388–405

Harkess A, Zhou J, Xu C, Bowers JE, Van der Hulst R, Ayyampalayam S, Mercati F, Riccardi P, McKain MR, Kakrana A, et al (2017) The asparagus genome sheds light on the origin and evolution of a young Y chromosome. Nature Communications. doi: 10.1038/s41467-017-01064-8

Hellsten U, Wright KM, Jenkins J, Shu S, Yuan Y, Wessler SR, Schmutz J, Willis JH, Rokhsar DS (2013) Fine-scale variation in meiotic recombination in Mimulus inferred from population shotgun sequencing. Proc Natl Acad Sci U S A 110: 19478–19482

Hibrand Saint-Oyant L, Ruttink T, Hamama L, Kirov I, Lakhwani D, Zhou NN, Bourke PM, Daccord N, Leus L, Schulz D, et al (2018) A high-quality genome sequence of Rosa chinensis to elucidate ornamental traits. Nat Plants 4: 473–484

Hirsch CD, Springer NM (2017) Transposable element influences on gene expression in plants. Biochim Biophys Acta Gene Regul Mech 1860: 157–165

Hirsch CN, Foerster JM, Johnson JM, Sekhon RS, Muttoni G, Vaillancourt B, Peñagaricano F, Lindquist E, Pedraza MA, Barry K, et al (2014) Insights into the maize pan-genome and pan-transcriptome. Plant Cell 26: 121–135

Hosmani PS, Flores-Gonzalez M, van de Geest H, Maumus F, Bakker LV, Schijlen E, van Haarst J, Cordewener J, Sanchez-Perez G, Peters S, et al An improved de novo assembly and annotation of the tomato reference genome using single-molecule sequencing, Hi-C proximity ligation and optical maps. doi: 10.1101/767764

Hua Z, Pool JE, Schmitz RJ, Schultz MD, Shiu S-H, Ecker JR, Vierstra RD (2013) Epigenomic programming contributes to the genomic drift evolution of the F-Box protein superfamily in Arabidopsis. Proc Natl Acad Sci U S A 110: 16927–16932

Hulse-Kemp AM, Maheshwari S, Stoffel K, Hill TA, Jaffe D, Williams SR, Weisenfeld N, Ramakrishnan S, Kumar V, Shah P, et al (2018) Reference quality assembly of the 3.5-Gb genome of Capsicum annuum from a single linked-read library. Horticulture Research. doi: 10.1038/s41438-017-0011-0

Hu TT, Pattyn P, Bakker EG, Cao J, Cheng J-F, Clark RM, Fahlgren N, Fawcett JA, Grimwood J, Gundlach H, et al (2011) The Arabidopsis lyrata genome sequence and the basis of rapid genome size change. Nat Genet 43: 476–481

Initiative TIB, The International Brachypodium Initiative (2010) Genome sequencing and analysis of the model grass Brachypodium distachyon. Nature 463: 763–768

Jaillon O, Aury J-M, Noel B, Policriti A, Clepet C, Casagrande A, Choisne N, Aubourg S, Vitulo N, Jubin C, et al (2007) The grapevine genome sequence suggests ancestral hexaploidization in major angiosperm phyla. Nature 449: 463–467

Jiang X, Assis R (2019) Rapid functional divergence after small-scale gene duplication in grasses. BMC Evol Biol 19: 97

Jiao Y, Peluso P, Shi J, Liang T, Stitzer MC, Wang B, Campbell MS, Stein JC, Wei X, Chin C-S, et al (2017) Improved maize reference genome with single-molecule technologies. Nature 546: 524–527

Jin Y, Qian H (2019) V.PhyloMaker: an R package that can generate very large phylogenies for vascular plants. Ecography 42: 1353–1359

Katoh K, Standley DM (2013) MAFFT multiple sequence alignment software version 7: improvements in performance and usability. Mol Biol Evol 30: 772–780

Kawahara Y, de la Bastide M, Hamilton JP, Kanamori H, McCombie WR, Ouyang S, Schwartz DC, Tanaka T, Wu J, Zhou S, et al (2013) Improvement of the Oryza sativa Nipponbare reference genome using next generation sequence and optical map data. Rice 6: 4

Kawakatsu T, Huang S-SC, Jupe F, Sasaki E, Schmitz RJ, Urich MA, Castanon R, Nery JR, Barragan C, He Y, et al (2016) Epigenomic Diversity in a Global Collection of Arabidopsis thaliana Accessions. Cell 166: 492–505

Keller TE, Yi SV (2014) DNA methylation and evolution of duplicate genes. Proceedings of the National Academy of Sciences 111: 5932–5937

Kim KD, El Baidouri M, Abernathy B, Iwata-Otsubo A, Chavarro C, Gonzales M, Libault M, Grimwood J, Jackson SA (2015) A Comparative Epigenomic Analysis of Polyploidy-Derived Genes in Soybean and Common Bean. Plant Physiol 168: 1433–1447

Klepikova AV, Kasianov AS, Gerasimov ES, Logacheva MD, Penin AA (2016) A high resolution map of the Arabidopsis thaliana developmental transcriptome based on RNA-seq profiling. Plant J 88: 1058–1070

Lamesch P, Berardini TZ, Li D, Swarbreck D, Wilks C, Sasidharan R, Muller R, Dreher K, Alexander DL, Garcia-Hernandez M, et al (2012) The Arabidopsis Information Resource (TAIR): improved gene annotation and new tools. Nucleic Acids Res 40: D1202–10

Li Q, Li H, Huang W, Xu Y, Zhou Q, Wang S, Ruan J, Huang S, Zhang Z (2019) A chromosome-scale genome assembly of cucumber (Cucumis sativus L.). Gigascience. doi: 10.1093/gigascience/giz072

Liu M-J, Zhao J, Cai Q-L, Liu G-C, Wang J-R, Zhao Z-H, Liu P, Dai L, Yan G, Wang W-J, et al (2014) The complex jujube genome provides insights into fruit tree biology. Nat Commun 5: 5315

Li W-H, Yang J, Gu X (2005) Expression divergence between duplicate genes. Trends Genet 21:602–607

Li Z, Defoort J, Tasdighian S, Maere S, Van de Peer Y, De Smet R (2016) Gene Duplicability of Core Genes Is Highly Consistent across All Angiosperms. Plant Cell 28: 326–344

Lovell JT, Jenkins J, Lowry DB, Mamidi S, Sreedasyam A, Weng X, Barry K, Bonnette J, Campitelli B, Daum C, et al (2018) The genomic landscape of molecular responses to natural drought stress in Panicum hallii. Nat Commun 9: 5213

Lovell JT, MacQueen AH, Mamidi S, Bonnette J, Jenkins J, Napier JD, Sreedasyam A, Healey A, Session A, Shu S, et al (2021) Genomic mechanisms of climate adaptation in polyploid bioenergy switchgrass. Nature 590: 438–444

Love MI, Huber W, Anders S (2014) Moderated estimation of fold change and dispersion for RNA-seq data with DESeq2. Genome Biol 15: 550

Lü P, Yu S, Zhu N, Chen Y-R, Zhou B, Pan Y, Tzeng D, Fabi JP, Argyris J, Garcia-Mas J, et al (2018) Genome encode analyses reveal the basis of convergent evolution of fleshy fruit ripening. Nat Plants 4: 784–791

Lynch M (2002) Genomics. Gene duplication and evolution. Science 297: 945–947

Lynch M, Conery JS (2000) The evolutionary fate and consequences of duplicate genes. Science 290: 1151–1155

Maere S, De Bodt S, Raes J, Casneuf T, Van Montagu M, Kuiper M, Van de Peer Y (2005) Modeling gene and genome duplications in eukaryotes. Proceedings of the National Academy of Sciences 102: 5454–5459

Mamidi S, Healey A, Huang P, Grimwood J, Jenkins J, Barry K, Sreedasyam A, Shu S, Lovell JT, Feldman M, et al The Setaria viridis genome and diversity panel enables discovery of a novel domestication gene. doi: 10.1101/744557

McCormick RF, Truong SK, Sreedasyam A, Jenkins J, Shu S, Sims D, Kennedy M, Amirebrahimi M, Weers BD, McKinley B, et al (2018) The Sorghum bicolor reference genome: improved assembly, gene annotations, a transcriptome atlas, and signatures of genome organization. Plant J 93: 338–354

Ming R, Hou S, Feng Y, Yu Q, Dionne-Laporte A, Saw JH, Senin P, Wang W, Ly BV, Lewis KLT, et al (2008) The draft genome of the transgenic tropical fruit tree papaya (Carica papaya Linnaeus). Nature 452: 991–996

Ming R, VanBuren R, Liu Y, Yang M, Han Y, Li L-T, Zhang Q, Kim M-J, Schatz MC, Campbell M, et al (2013) Genome of the long-living sacred lotus (Nelumbo nucifera Gaertn.). Genome Biol 14: R41

Miyata T, Yasunaga T (1980) Molecular evolution of mRNA: a method for estimating evolutionary rates of synonymous and amino acid substitutions from homologous nucleotide sequences and its application. J Mol Evol 16: 23–36

Motamayor JC, Mockaitis K, Schmutz J, Haiminen N, Livingstone D 3rd, Cornejo O, Findley SD, Zheng P, Utro F, Royaert S, et al (2013) The genome sequence of the most widely cultivated cacao type and its use to identify candidate genes regulating pod color. Genome Biol 14: r53

Niederhuth CE, Bewick AJ, Ji L, Alabady MS, Kim KD, Li Q, Rohr NA, Rambani A, Burke JM, Udall JA, et al (2016) Widespread natural variation of DNA methylation within angiosperms. Genome Biol 17: 194

Niederhuth CE, Schmitz RJ (2017) Putting DNA methylation in context: from genomes to gene expression in plants. Biochim Biophys Acta Gene Regul Mech 1860: 149–156

Noshay JM, Anderson SN, Zhou P, Ji L, Ricci W, Lu Z, Stitzer MC, Crisp PA, Hirsch CN, Zhang X, et al (2019) Monitoring the interplay between transposable element families and DNA methylation in maize. PLoS Genet 15: e1008291

Ohno S (1970) Evolution by Gene Duplication. doi: 10.1007/978-3-642-86659-3

Ong-Abdullah M, Ordway JM, Jiang N, Ooi S-E, Kok S-Y, Sarpan N, Azimi N, Hashim AT, Ishak Z, Rosli SK, et al (2015) Loss of Karma transposon methylation underlies the mantled somaclonal variant of oil palm. Nature 525: 533–537

O’Rourke JA, Iniguez LP, Fu F, Bucciarelli B, Miller SS, Jackson SA, McClean PE, Li J, Dai X, Zhao PX, et al (2014) An RNA-Seq based gene expression atlas of the common bean. BMC Genomics 15: 866

Otto SP, Whitton J (2000) Polyploid Incidence and Evolution. Annual Review of Genetics 34: 401–437

Ou S, Su W, Liao Y, Chougule K, Agda JRA, Hellinga AJ, Lugo CSB, Elliott TA, Ware D, Peterson T, et al (2019) Benchmarking transposable element annotation methods for creation of a streamlined, comprehensive pipeline. Genome Biol 20: 275

Panchy N, Lehti-Shiu MD, Shiu S-H (2016) Evolution of gene duplication in plants. Plant Physiology 00523.2016

Parkin IAP, Koh C, Tang H, Robinson SJ, Kagale S, Clarke WE, Town CD, Nixon J, Krishnakumar V, Bidwell SL, et al (2014) Transcriptome and methylome profiling reveals relics of genome dominance in the mesopolyploid Brassica oleracea. Genome Biol 15: R77

Paterson AH, Chapman BA, Kissinger JC, Bowers JE, Feltus FA, Estill JC (2006) Many gene and domain families have convergent fates following independent whole-genome duplication events in Arabidopsis, Oryza, Saccharomyces and Tetraodon. Trends Genet 22: 597–602

Paterson AH, Wendel JF, Gundlach H, Guo H, Jenkins J, Jin D, Llewellyn D, Showmaker KC, Shu S, Udall J, et al (2012) Repeated polyploidization of Gossypium genomes and the evolution of spinnable cotton fibres. Nature 492: 423–427

Picard CL, Gehring M (2017) Proximal methylation features associated with nonrandom changes in gene body methylation. Genome Biol 18: 73

Pophaly SD, Tellier A (2015) Population Level Purifying Selection and Gene Expression Shape Subgenome Evolution in Maize. Mol Biol Evol 32: 3226–3235

Potter SC, Luciani A, Eddy SR, Park Y, Lopez R, Finn RD (2018) HMMER web server: 2018 update. Nucleic Acids Res 46: W200–W204

Qiao X, Li Q, Yin H, Qi K, Li L, Wang R, Zhang S, Paterson AH (2019) Gene duplication and evolution in recurring polyploidization-diploidization cycles in plants. Genome Biol 20: 38

Raju SKK, Ritter EJ, Niederhuth CE (2019) Establishment, maintenance, and biological roles of non-CG methylation in plants. Essays in Biochemistry 63: 743–755

Revell LJ (2012) phytools: an R package for phylogenetic comparative biology (and other things). Methods in Ecology and Evolution 3: 217–223

Rodin SN, Riggs AD (2003) Epigenetic silencing may aid evolution by gene duplication. J Mol Evol 56: 718–729

Sato S, Nakamura Y, Kaneko T, Asamizu E, Kato T, Nakao M, Sasamoto S, Watanabe A, Ono A, Kawashima K, et al (2008) Genome structure of the legume, Lotus japonicus. DNA Res 15: 227–239

Schmutz J, Cannon SB, Schlueter J, Ma J, Mitros T, Nelson W, Hyten DL, Song Q, Thelen JJ, Cheng J, et al (2010) Genome sequence of the palaeopolyploid soybean. Nature 463: 178–183

Schmutz J, McClean PE, Mamidi S, Wu GA, Cannon SB, Grimwood J, Jenkins J, Shu S, Song Q, Chavarro C, et al (2014) A reference genome for common bean and genomewide analysis of dual domestications. Nat Genet 46: 707–713

Schultz MD, He Y, Whitaker JW, Hariharan M, Mukamel EA, Leung D, Rajagopal N, Nery JR, Urich MA, Chen H, et al (2015) Human body epigenome maps reveal noncanonical DNA methylation variation. Nature 523: 212–216

Schultz MD, Schmitz RJ, Ecker JR (2012) “Leveling” the playing field for analyses of singlebase resolution DNA methylomes. Trends in Genetics 28: 583–585

Secco D, Wang C, Shou H, Schultz MD, Chiarenza S, Nussaume L, Ecker JR, Whelan J, Lister R (2015) Stress induced gene expression drives transient DNA methylation changes at adjacent repetitive elements. Elife. doi: 10.7554/eLife.09343

Seymour DK, Koenig D, Hagmann J, Becker C, Weigel D (2014) Evolution of DNA Methylation Patterns in the Brassicaceae is Driven by Differences in Genome Organization. PLoS Genetics 10: e1004785

Sharma SK, Bolser D, de Boer J, Sønderkær M, Amoros W, Carboni MF, D’Ambrosio JM, de la Cruz G, Di Genova A, Douches DS, et al (2013) Construction of reference chromosome-scale pseudomolecules for potato: integrating the potato genome with genetic and physical maps. G3 3: 2031–2047

Singh R, Ong-Abdullah M, Low E-TL, Manaf MAA, Rosli R, Nookiah R, Ooi LC-L, Ooi S-E, Chan K-L, Halim MA, et al (2013) Oil palm genome sequence reveals divergence of interfertile species in Old and New worlds. Nature 500: 335–339

Slotte T, Hazzouri KM, Ågren JA, Koenig D, Maumus F, Guo Y-L, Steige K, Platts AE, Escobar JS, Newman LK, et al (2013) The Capsella rubella genome and the genomic consequences of rapid mating system evolution. Nat Genet 45: 831–835

Soltis PS, Marchant DB, Van de Peer Y, Soltis DE (2015) Polyploidy and genome evolution in plants. Curr Opin Genet Dev 35: 119–125

Song Q, Zhang T, Stelly DM, Chen ZJ (2017) Epigenomic and functional analyses reveal roles of epialleles in the loss of photoperiod sensitivity during domestication of allotetraploid cottons. Genome Biol 18: 99

Suyama M, Torrents D, Bork P (2006) PAL2NAL: robust conversion of protein sequence alignments into the corresponding codon alignments. Nucleic Acids Res 34: W609–12

Takuno S, Gaut BS (2013) Gene body methylation is conserved between plant orthologs and is of evolutionary consequence. Proc Natl Acad Sci U S A 110: 1797–1802

Takuno S, Gaut BS (2012) Body-methylated genes in Arabidopsis thaliana are functionally important and evolve slowly. Mol Biol Evol 29: 219–227

Takuno S, Ran J-H, Gaut BS (2016) Evolutionary patterns of genic DNA methylation vary across land plants. Nat Plants 2: 15222

Tang H, Krishnakumar V, Bidwell S, Rosen B, Chan A, Zhou S, Gentzbittel L, Childs KL, Yandell M, Gundlach H, et al (2014) An improved genome release (version Mt4.0) for the model legume Medicago truncatula. BMC Genomics 15: 312

Tran RK, Henikoff JG, Zilberman D, Ditt RF, Jacobsen SE, Henikoff S (2005) DNA methylation profiling identifies CG methylation clusters in Arabidopsis genes. Curr Biol 15: 154–159

Turco GM, Kajala K, Kunde-Ramamoorthy G, Ngan C-Y, Olson A, Deshphande S, Tolkunov D, Waring B, Stelpflug S, Klein P, et al (2017) DNA methylation and gene expression regulation associated with vascularization in Sorghum bicolor. New Phytol 214: 1213–1229

Tuskan GA, Difazio S, Jansson S, Bohlmann J, Grigoriev I, Hellsten U, Putnam N, Ralph S, Rombauts S, Salamov A, et al (2006) The genome of black cottonwood, Populus trichocarpa (Torr. & Gray). Science 313: 1596–1604

Valliyodan B, Cannon SB, Bayer PE, Shu S, Brown AV, Ren L, Jenkins J, Chung CY-L, Chan T-F, Daum CG, et al (2019) Construction and comparison of three reference-quality genome assemblies for soybean. Plant J 100: 1066–1082

VanBuren R, Bryant D, Edger PP, Tang H, Burgess D, Challabathula D, Spittle K, Hall R, Gu J, Lyons E, et al (2015) Single-molecule sequencing of the desiccation-tolerant grass Oropetium thomaeum. Nature 527: 508–511

VanBuren R, Wai CM, Colle M, Wang J, Sullivan S, Bushakra JM, Liachko I, Vining KJ, Dossett M, Finn CE, et al (2018) A near complete, chromosome-scale assembly of the black raspberry (Rubus occidentalis) genome. Gigascience. doi: 10.1093/gigascience/giy094

Van de Peer Y, Mizrachi E, Marchal K (2017) The evolutionary significance of polyploidy. Nat Rev Genet 18: 411–424

Verde I, Jenkins J, Dondini L, Micali S, Pagliarani G, Vendramin E, Paris R, Aramini V, Gazza L, Rossini L, et al (2017) The Peach v2.0 release: high-resolution linkage mapping and deep resequencing improve chromosome-scale assembly and contiguity. BMC Genomics 18: 225

Wang D, Zhang Y, Zhang Z, Zhu J, Yu J (2010) KaKs_Calculator 2.0: a toolkit incorporating gamma-series methods and sliding window strategies. Genomics Proteomics Bioinformatics 8: 77–80

Wang H, Beyene G, Zhai J, Feng S, Fahlgren N, Taylor NJ, Bart R, Carrington JC, Jacobsen SE, Ausin I (2015) CG gene body DNA methylation changes and evolution of duplicated genes in cassava. Proc Natl Acad Sci U S A 112: 13729–13734

Wang J, Hossain MS, Lyu Z, Schmutz J, Stacey G, Xu D, Joshi T (2019) SoyCSN: Soybean context specific network analysis and prediction based on tissue specific transcriptome data. Plant Direct. doi: 10.1002/pld3.167

Wang J, Marowsky NC, Fan C (2014a) Divergence of gene body DNA methylation and evolution of plant duplicate genes. PLoS One 9: e110357

Wang L, Xie J, Hu J, Lan B, You C, Li F, Wang Z, Wang H (2018) Comparative epigenomics reveals evolution of duplicated genes in potato and tomato. Plant J 93: 460–471

Wang W, Haberer G, Gundlach H, Gläßer C, Nussbaumer T, Luo MC, Lomsadze A, Borodovsky M, Kerstetter RA, Shanklin J, et al (2014b) The Spirodela polyrhiza genome reveals insights into its neotenous reduction fast growth and aquatic lifestyle. Nat Commun 5: 3311

Wang X, Zhang Z, Fu T, Hu L, Xu C, Gong L, Wendel JF, Liu B (2017) Gene-body CG methylation and divergent expression of duplicate genes in rice. Scientific Reports. doi: 10.1038/s41598-017-02860-4

Wang Y, Li J, Paterson AH (2013a) MCScanX-transposed: detecting transposed gene duplications based on multiple colinearity scans. Bioinformatics 29: 1458–1460

Wang Y, Wang X, Lee T-H, Mansoor S, Paterson AH (2013b) Gene body methylation shows distinct patterns associated with different gene origins and duplication modes and has a heterogeneous relationship with gene expression inOryza sativa(rice). New Phytologist 198: 274–283

Wei T, Viliam S R package “corrplot”: Visualization of a Correlation Matrix (Version 0.84). R package “corrplot”: Visualization of a Correlation Matrix (Version 0.84). Available from https://github.com/taiyun/corrplot

Wu GA, Terol J, Ibanez V, López-García A, Pérez-Román E, Borredá C, Domingo C, Tadeo FR, Carbonell-Caballero J, Alonso R, et al (2018) Genomics of the origin and evolution of Citrus. Nature 554: 311–316

Xu C, Nadon BD, Kim KD, Jackson SA (2018) Genetic and epigenetic divergence of duplicate genes in two legume species. Plant Cell Environ 41: 2033–2044

Xue H, Wang S, Yao J-L, Deng CH, Wang L, Su Y, Zhang H, Zhou H, Sun M, Li X, et al (2018) Chromosome level high-density integrated genetic maps improve the Pyrus bretschneideri “DangshanSuli” v1.0 genome. BMC Genomics. doi: 10.1186/s12864-018-5224-6

Xu S, Brockmöller T, Navarro-Quezada A, Kuhl H, Gase K, Ling Z, Zhou W, Kreitzer C, Stanke M, Tang H, et al (2017) Wild tobacco genomes reveal the evolution of nicotine biosynthesis. Proc Natl Acad Sci U S A 114: 6133–6138

Yanai I, Benjamin H, Shmoish M, Chalifa-Caspi V, Shklar M, Ophir R, Bar-Even A, Horn-Saban S, Safran M, Domany E, et al (2005) Genome-wide midrange transcription profiles reveal expression level relationships in human tissue specification. Bioinformatics 21: 650–659

Yang R, Jarvis DE, Chen H, Beilstein MA, Grimwood J, Jenkins J, Shu S, Prochnik S, Xin M, Ma C, et al (2013) The Reference Genome of the Halophytic Plant Eutrema salsugineum. Front Plant Sci 4: 46

Yang Y, Tang K, Datsenka TU, Liu W, Lv S, Lang Z, Wang X, Gao J, Wang W, Nie W, et al (2019) Critical function of DNA methyltransferase 1 in tomato development and regulation of the DNA methylome and transcriptome. J Integr Plant Biol 61: 1224–1242

Yang Z, Bielawski JP (2000) Statistical methods for detecting molecular adaptation. Trends in Ecology & Evolution 15: 496–503

Zemach A, McDaniel IE, Silva P, Zilberman D (2010) Genome-wide evolutionary analysis of eukaryotic DNA methylation. Science 328: 916–919

Zhang J (2003) Evolution by gene duplication: an update. Trends in Ecology & Evolution 18: 292–298

Zhang X, Yazaki J, Sundaresan A, Cokus S, Chan SW-L, Chen H, Henderson IR, Shinn P, Pellegrini M, Jacobsen SE, et al (2006) Genome-wide high-resolution mapping and functional analysis of DNA methylation in arabidopsis. Cell 126: 1189–1201

Zhao XP, Si Y, Hanson RE, Crane CF, Price HJ, Stelly DM, Wendel JF, Paterson AH (1998) Dispersed repetitive DNA has spread to new genomes since polyploid formation in cotton. Genome Res 8: 479–492

