## Supplemental Figures 1-20 for "DNA methylation signatures of duplicate gene evolution in angiosperms"

**Figure S1: Schematic representation of genic methylation classification.**

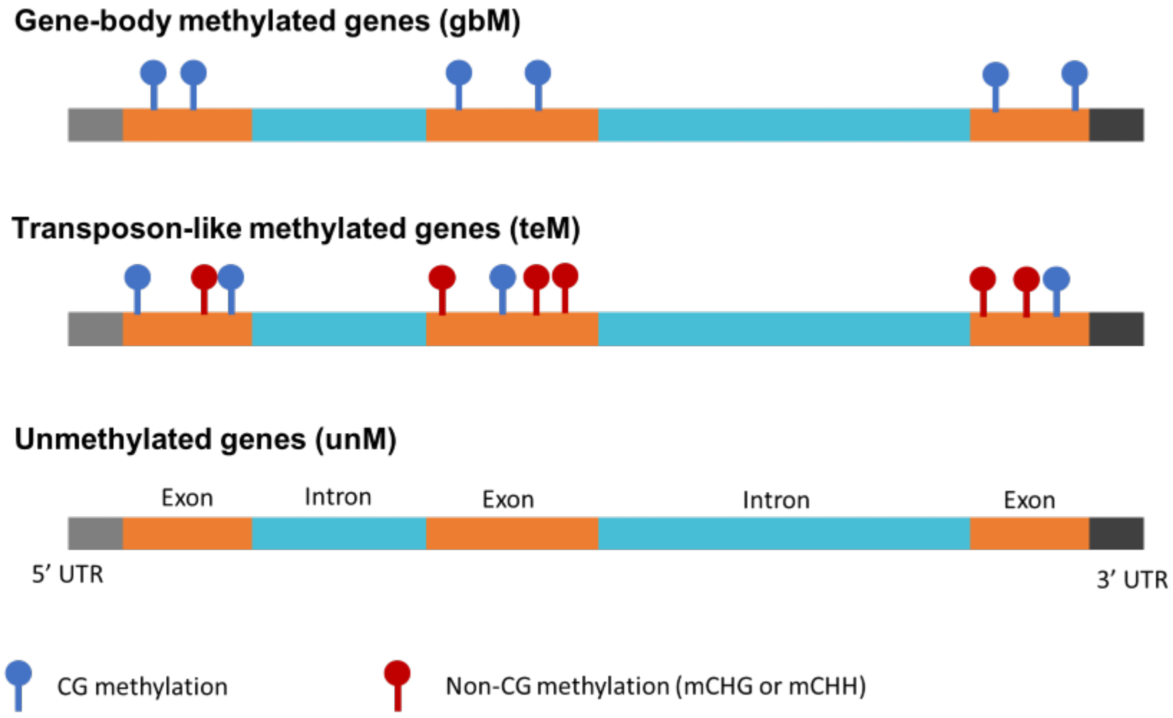

**Figure S2: Schematic representation of different orthogroup classification.**

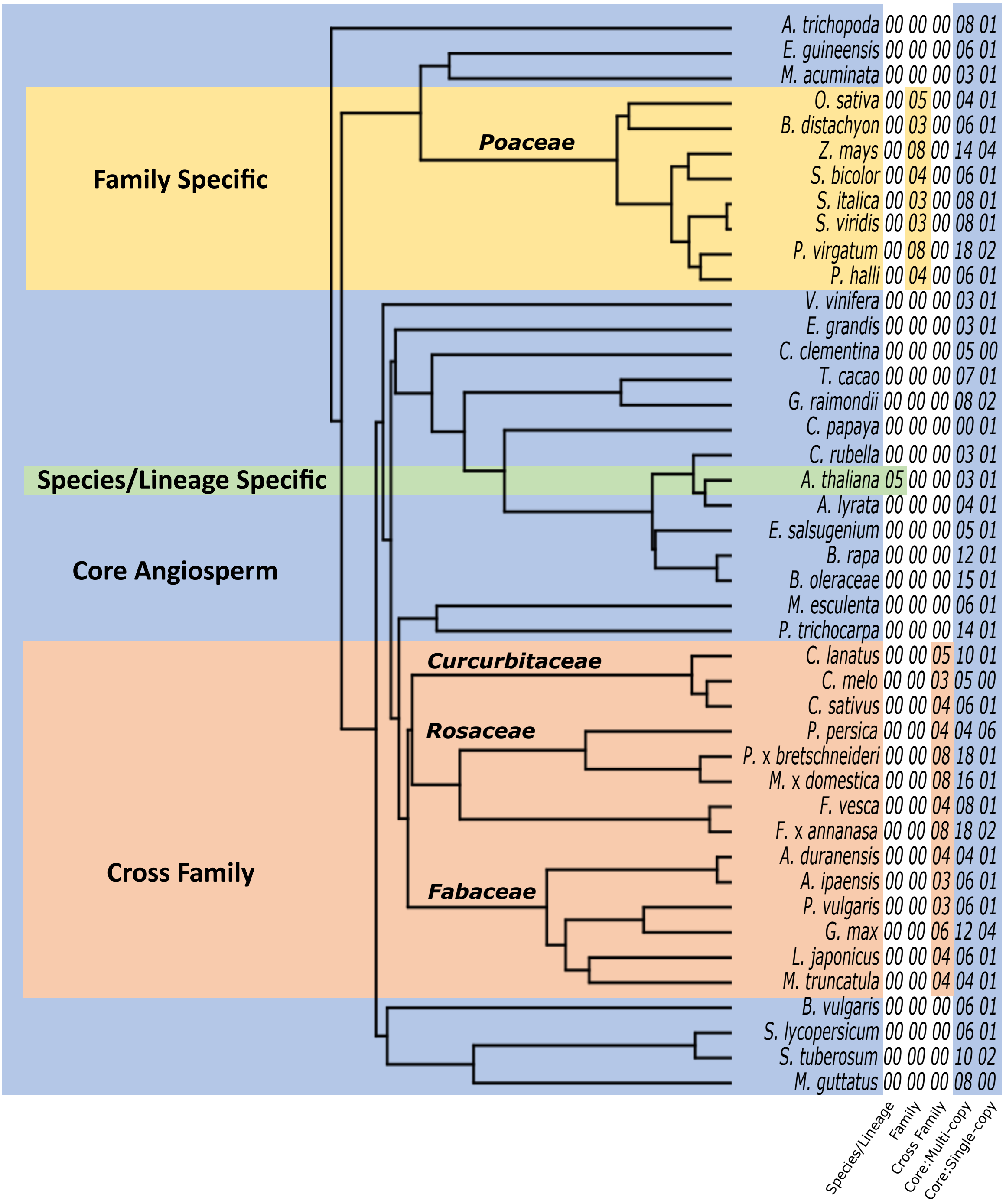

**Figure S3: Distribution of orthogroups across 58 angiosperm species.**
Histogram showing the number of orthogroups represented in 1 to 58 species (A) and the same plot zoomed into species with 2 to 58 species (B). Orange colored bars represent those orthogroups classified as 'core angiosperm'.

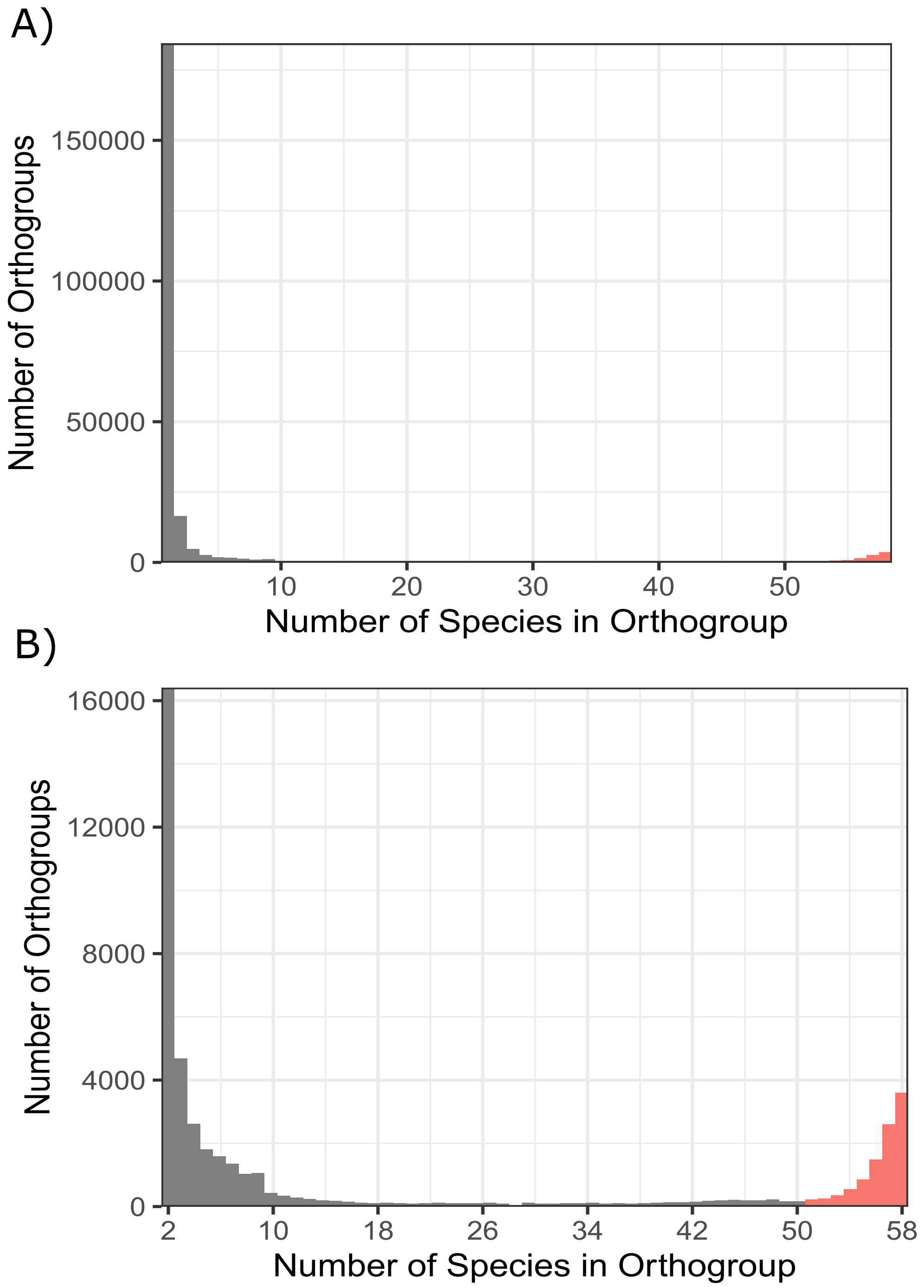

**Figure S4: Distribution of orthogroups in genic methylation classes.**
**A)** For each species, the percentage of genes classified into different orthogroup categories (core:single-copy, core: multi-copy, cross-family, family-specific, and species/lineage-specific) in each of the three genic methylation classification (gbM, teM, and unM genes).
**B)** Distribution of genes classified as gbM, unM, teM, unclassified, and 'missing methylation data' across different orthogroup classifications (Core:single copy, core:multi-copy, cross-family, family-specific, and species/lineage specific)

**A)**

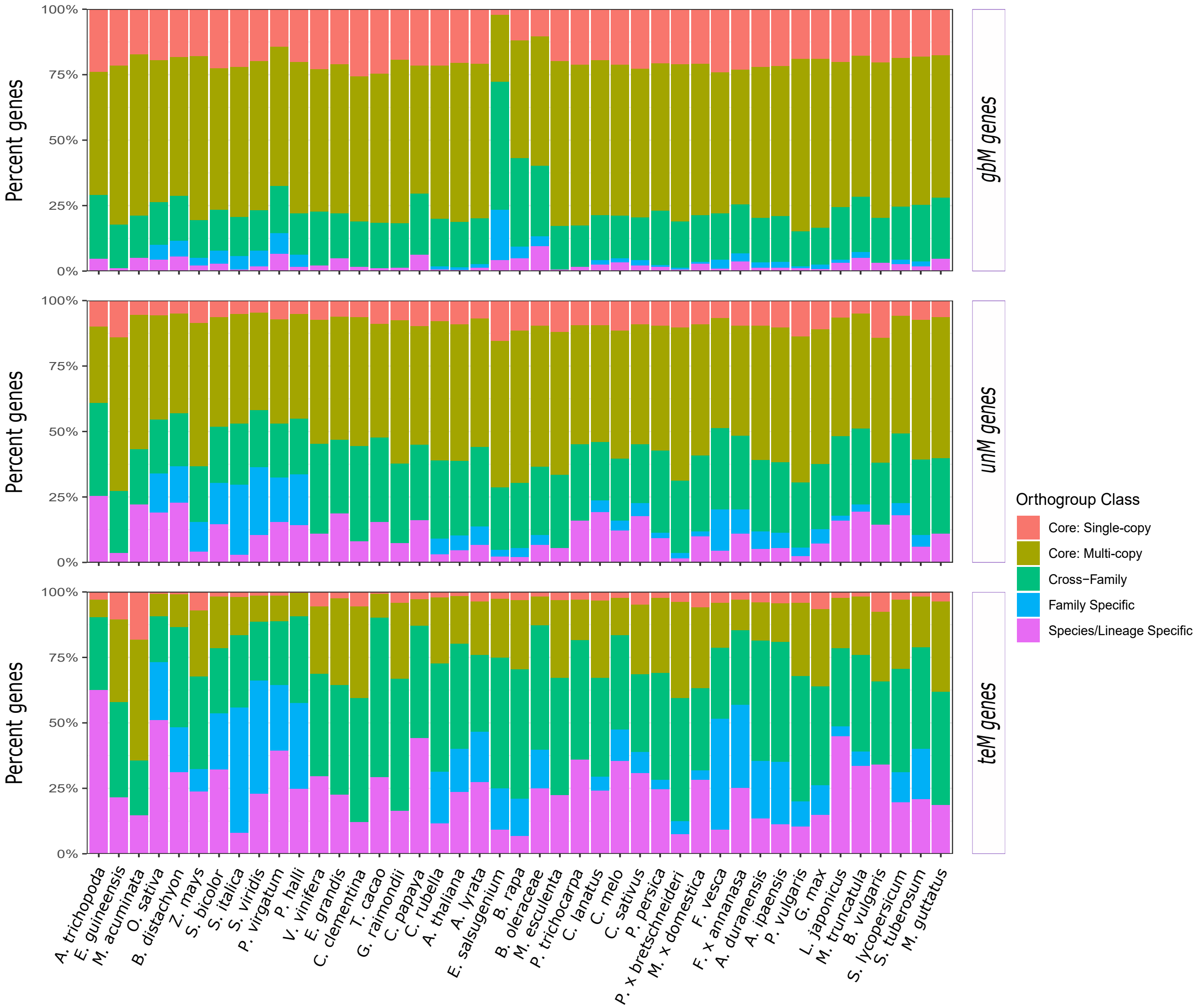

**Figure S4: Distribution of orthogroups in genic methylation classes.**
**A)** For each species, the percentage of genes classified into different orthogroup categories (core:single-copy, core: multi-copy, cross-family, family-specific, and species/lineage-specific) in each of the three genic methylation classification (gbM, teM, and unM genes).
**B)** Distribution of genes classified as gbM, unM, teM, unclassified, and 'missing methylation data' across different orthogroup classifications (Core:single copy, core:multi-copy, cross-family, family-specific, and species/lineage specific)

**B)**

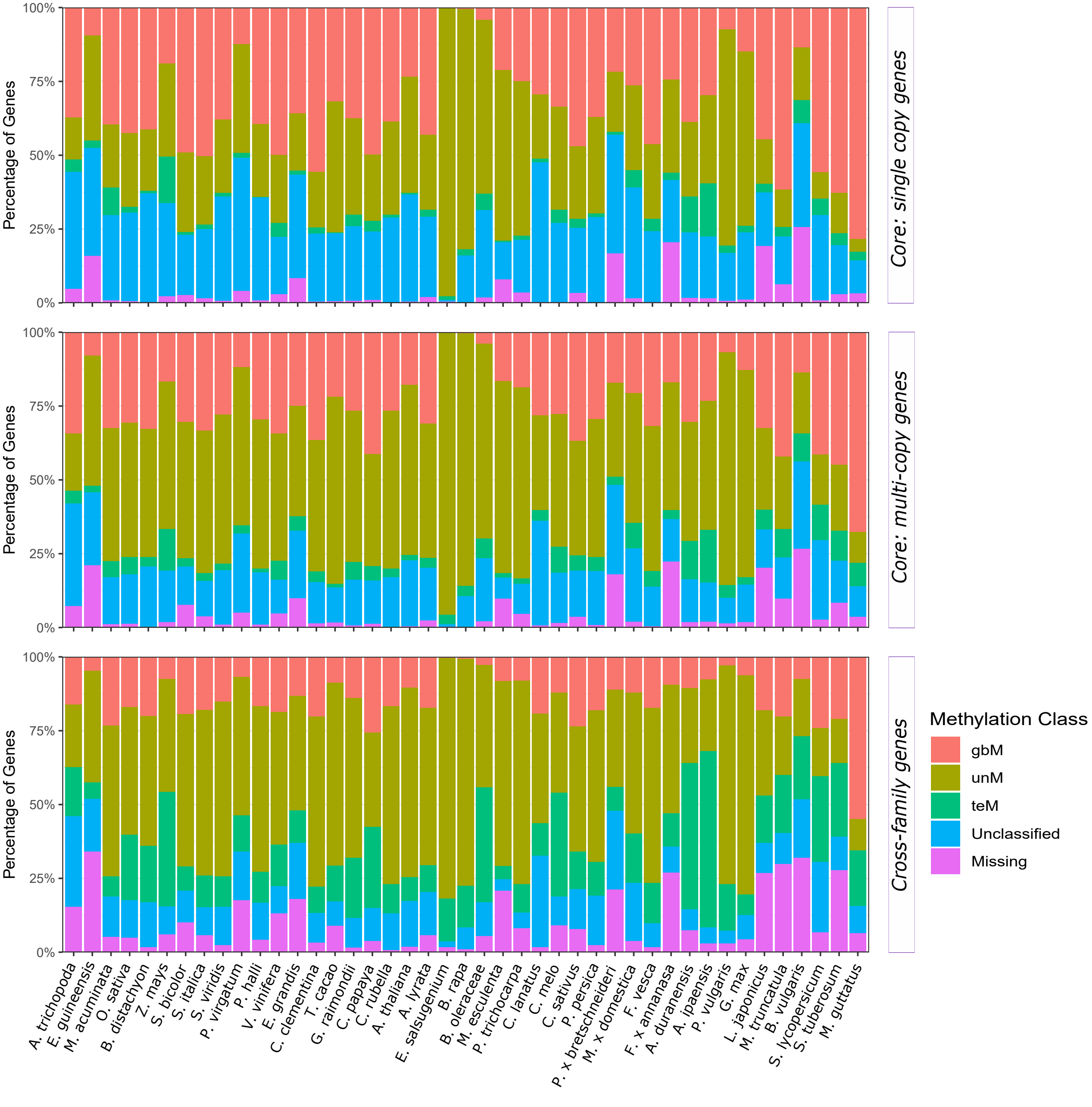

**Figure S4: Distribution of orthogroups in genic methylation classes.**
**A)** For each species, the percentage of genes classified into different orthogroup categories (core:single-copy, core: multi-copy, cross-family, family-specific, and species/lineage-specific) in each of the three genic methylation classification (gbM, teM, and unM genes).
**B)** Distribution of genes classified as gbM, unM, teM, unclassified, and 'missing methylation data' across different orthogroup classifications (Core:single copy, core:multi-copy, cross-family, family-specific, and species/lineage specific)

**B)**

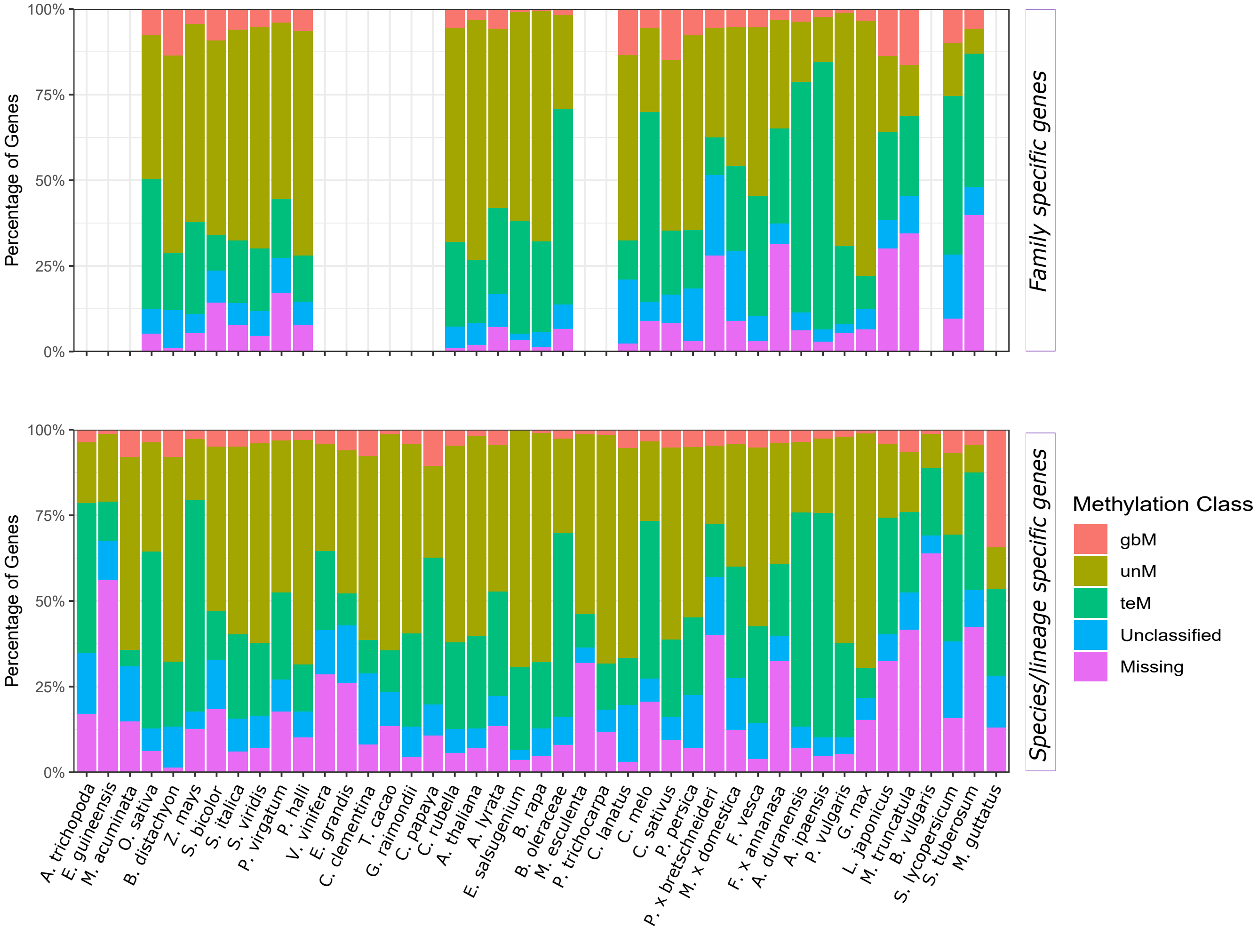

**Figure S5**: Enrichment or depletion of different genic methylation classes (gbM, unM, and teM) for each orthogroup category (core:multi-copy, core:single-copy, cross-family, family-specific, and lineage/species-specific). Increasing shades of cyan indicates greater depletion, while increasing shades of magenta represents greater enrichment. Unless indicated, all associations are statistically significant at a FDR-corrected p-value < 0.05. ‘NS’ indicates no statistical significance.

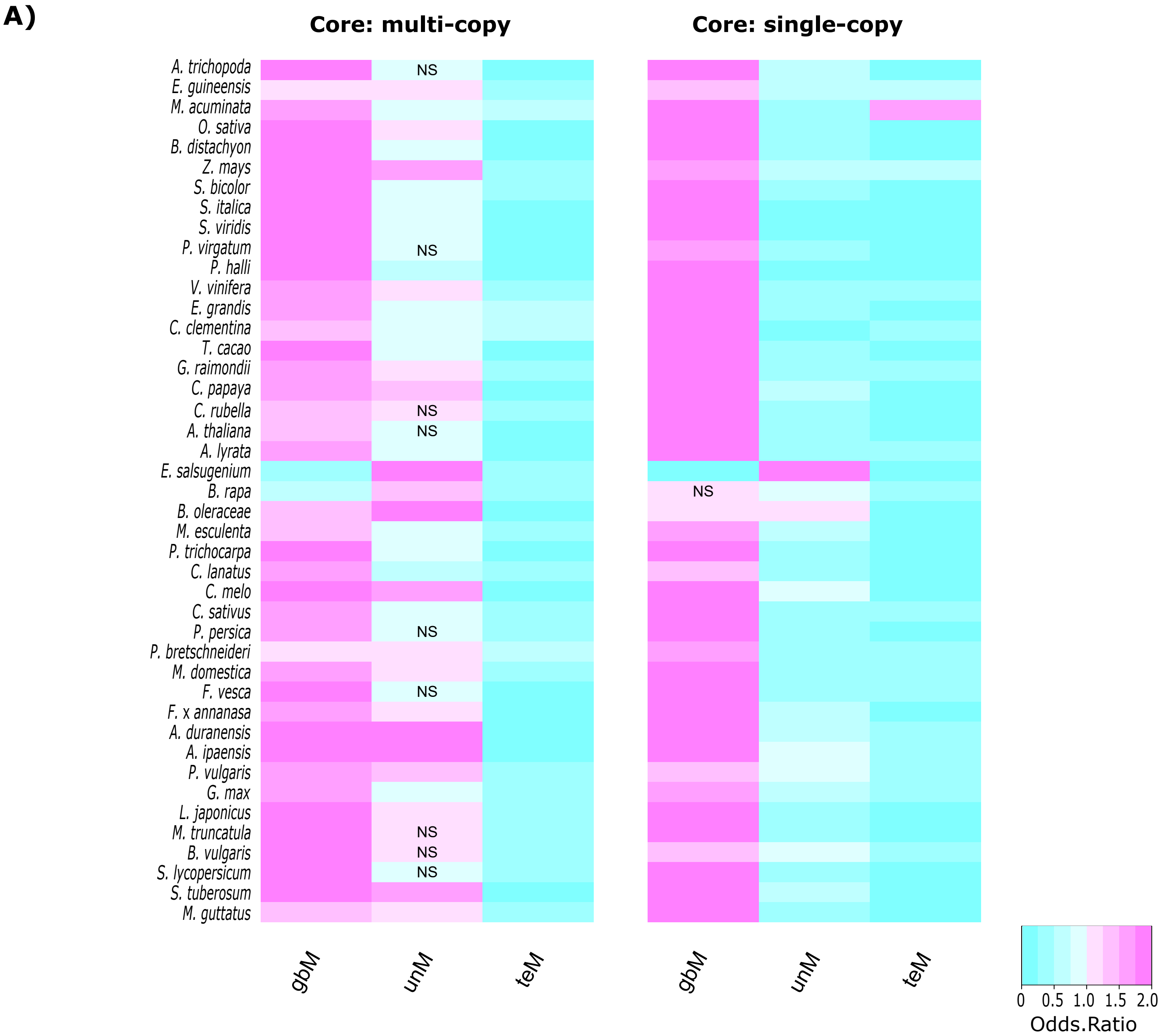

**Figure S5**: Enrichment or depletion of different genic methylation classes (gbM, unM, and teM) for each orthogroup category (core:multi-copy, core:single-copy, cross-family, family-specific, and lineage/species-specific). Increasing shades of cyan indicates greater depletion, while increasing shades of magenta represents greater enrichment. Unless indicated, all associations are statistically significant at a FDR-corrected p-value < 0.05. 'NS' indicates no statistical significance.

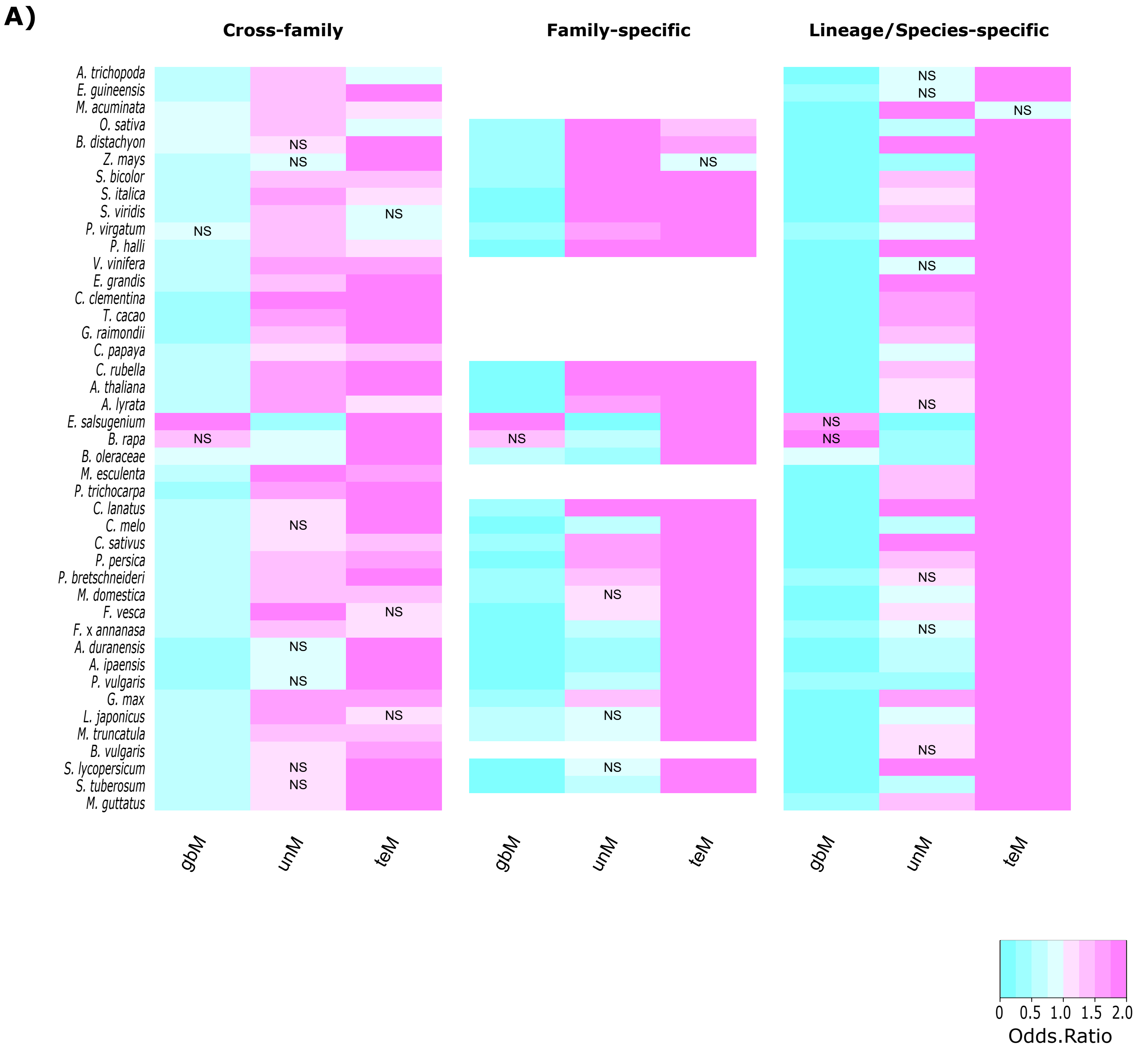

**Figure S5**: Enrichment or depletion of different types of duplicates (whole-genome, and different types of single gene duplicates) for each orthogroup category (core:multi-copy, core:single-copy, cross-family, family-specific, and lineage/species-specific) for . Increasing shades of cyan indicates greater depletion, while increasing shades of magenta represents greater enrichment. Unless indicated, all associations are statistically significant at a FDR-corrected p-value < 0.05. 'NS' indicates no statistical significance.

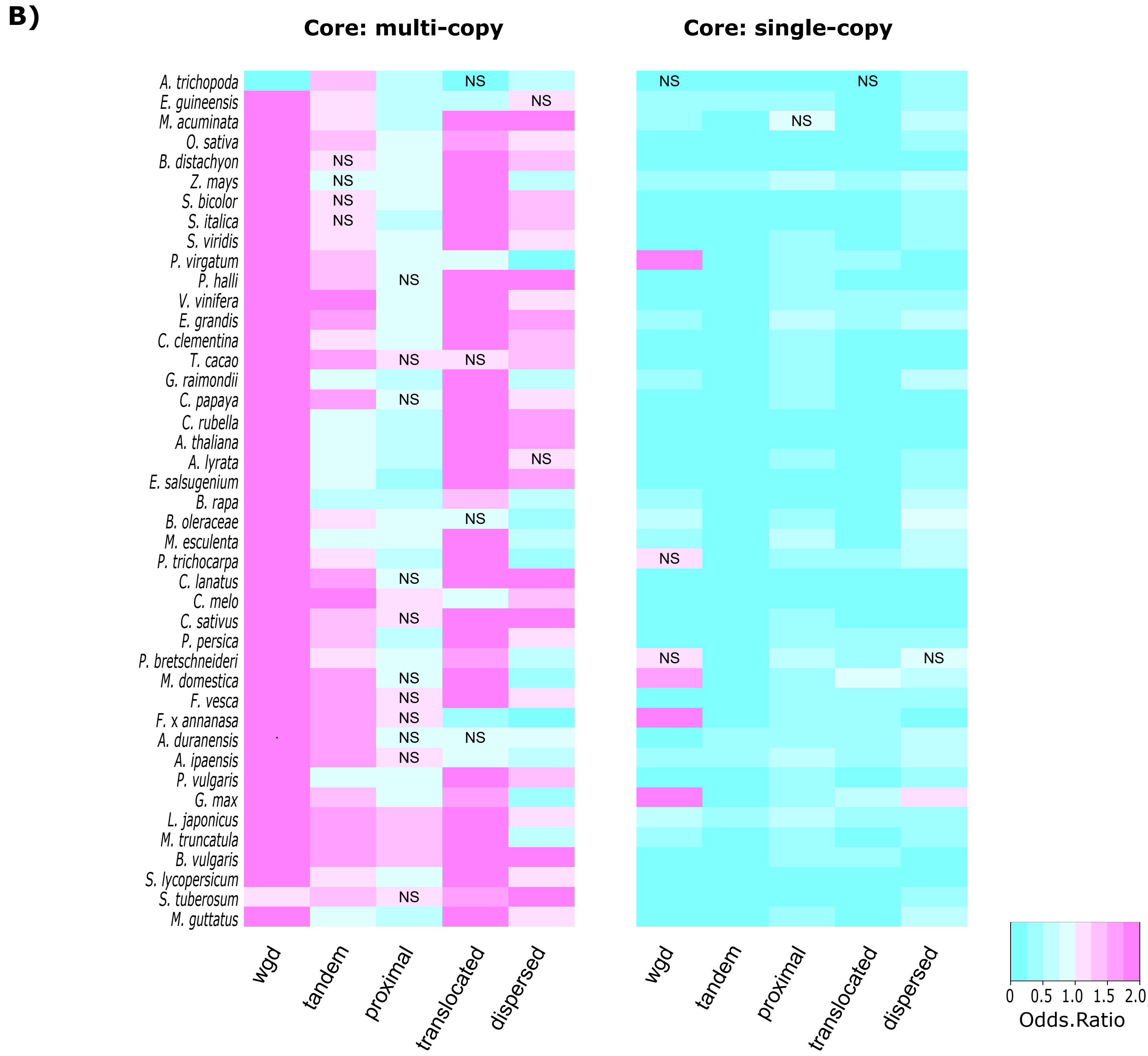

**Figure S5**: Enrichment or depletion of different types of duplicates (whole-genome, and different types of single gene duplicates) for each orthogroup category (core:multi-copy, core:single-copy, cross-family, family-specific, and lineage/species-specific) for . Increasing shades of cyan indicates greater depletion, while increasing shades of magenta represents greater enrichment. Unless indicated, all associations are statistically significant at a FDR-corrected p-value < 0.05. ‘NS’ indicates no statistical significance.

**B)**

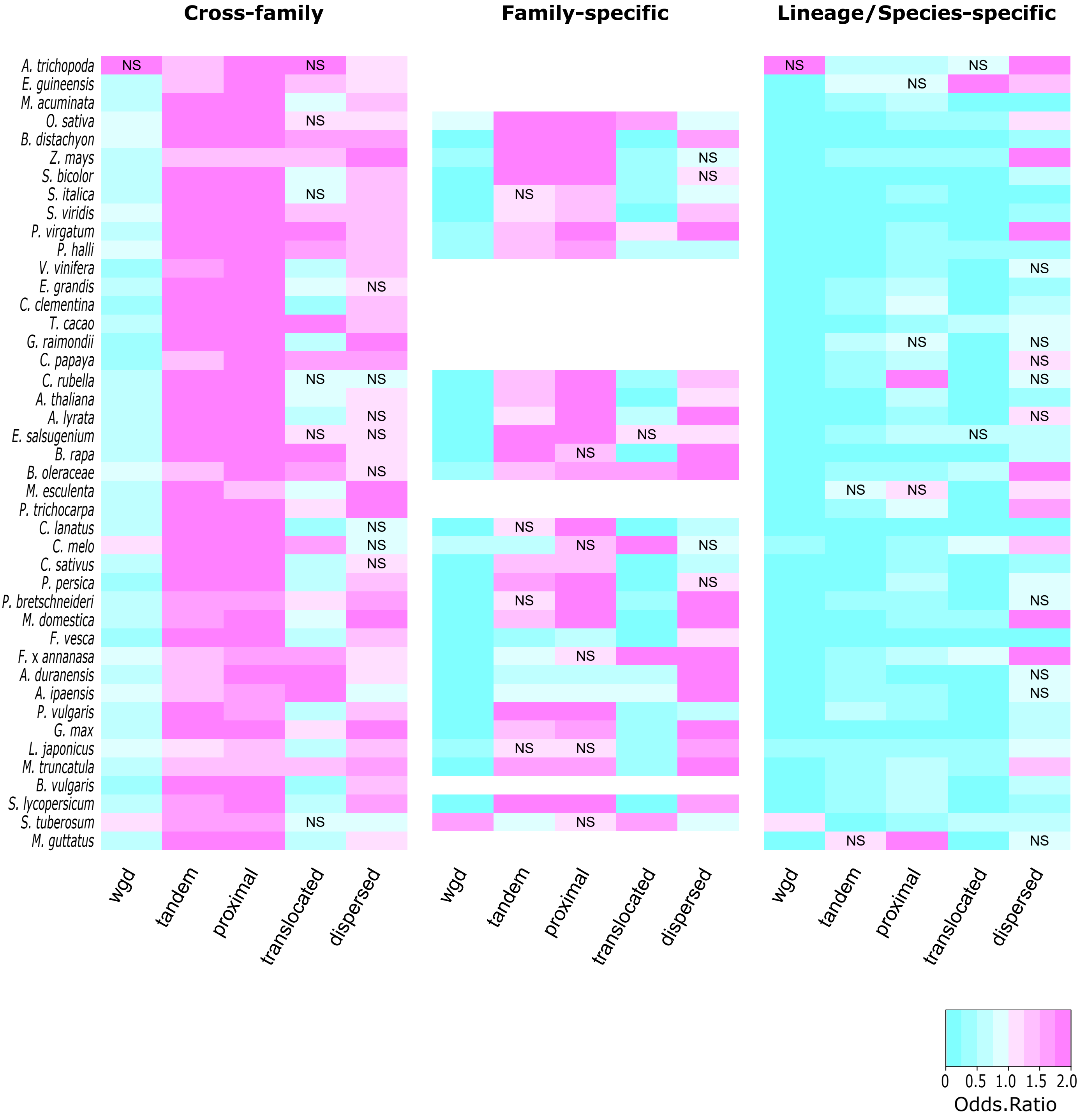

**Figure S6**: **Proportion of paralogs with similar and divergent DNA methylation profiles.**
The proportion of duplicate pairs with similar DNA methylation profiles among different types of duplicate genes (Whole-genome duplicates - WGD, Single-gene duplicates - tandem, proximal, translocated, and dispersed) are shown in Blue. Yellow bars represent proportion of duplicate pairs with divergent DNA methylation profiles. Grey bars represent cases where DNA methylation status of at least one of the duplicate pairs was 'undetermined'.

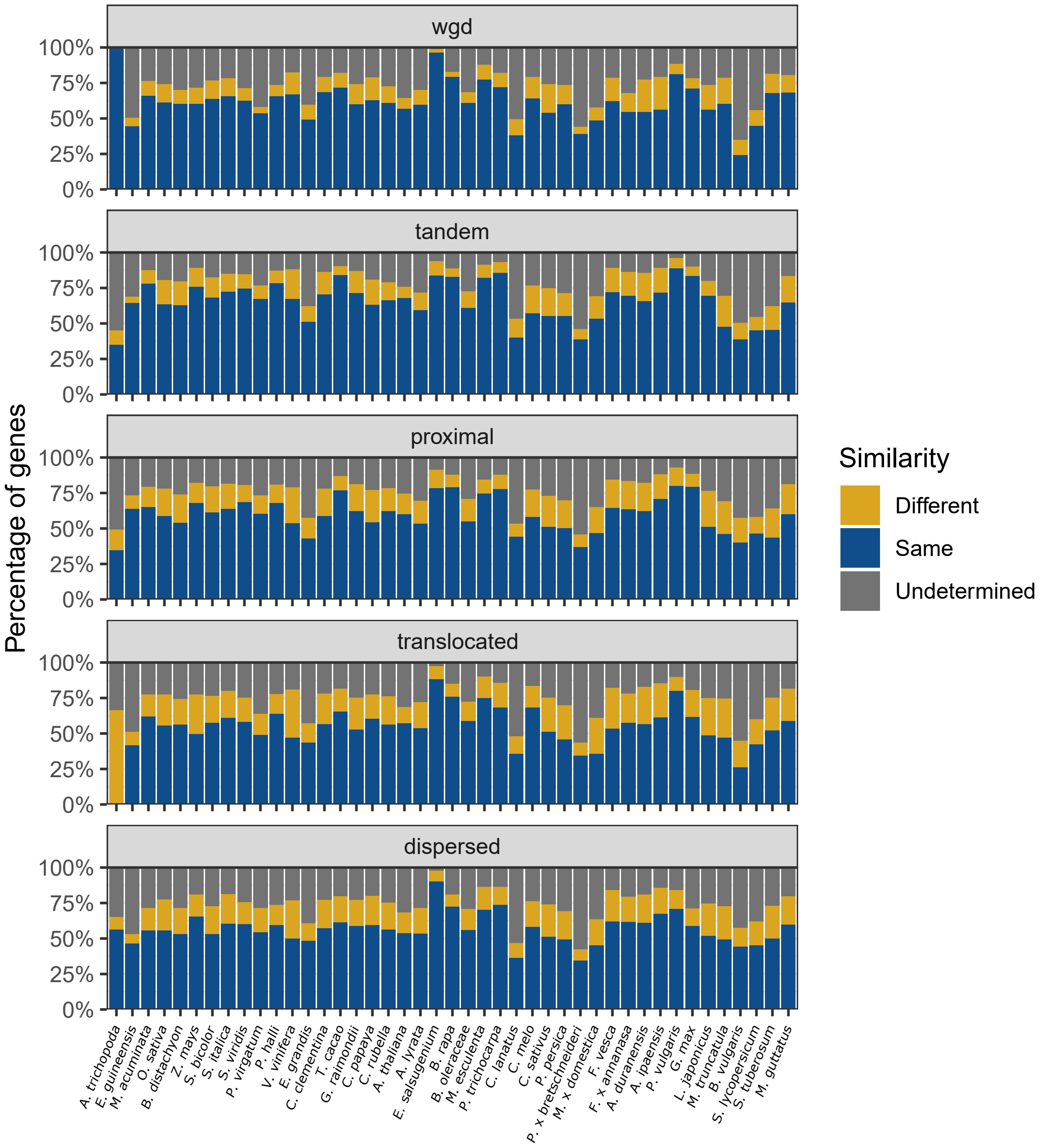

**Figure S7**: Distribution of genic methylation classified genes based on synonymous substitution (Ks) across different types of gene duplicate pairs. Whole-genome duplicates - WGD, Single-gene duplicates - SGD (combined data from tandem, proximal, translocated, and dispersed duplicates).

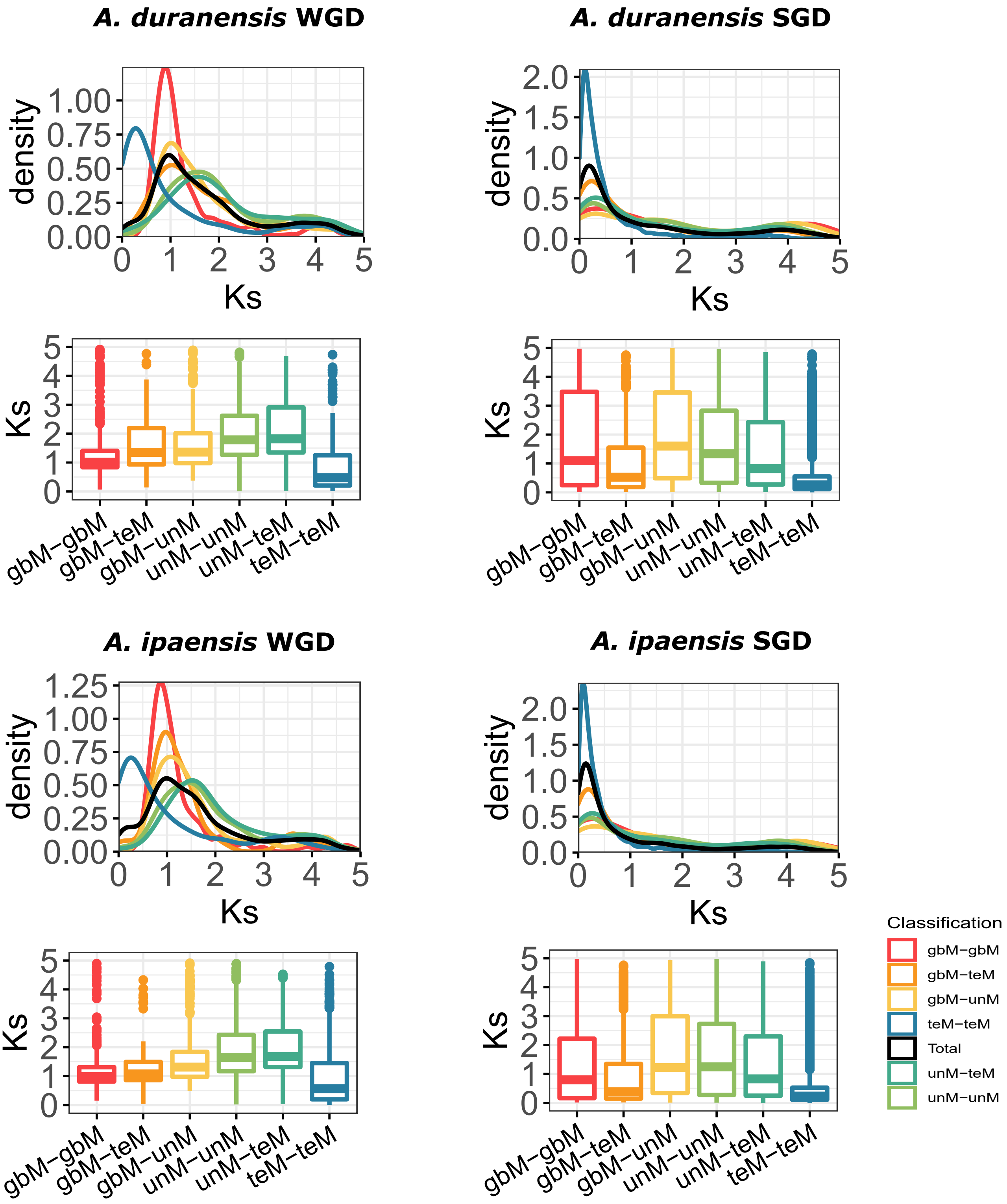

**Figure S7**: Distribution of genic methylation classified genes based on synonymous substitution (Ks) across different types of gene duplicate pairs. Whole-genome duplicates - WGD, Single-gene duplicates - SGD (combined data from tandem, proximal, translocated, and dispersed duplicates).

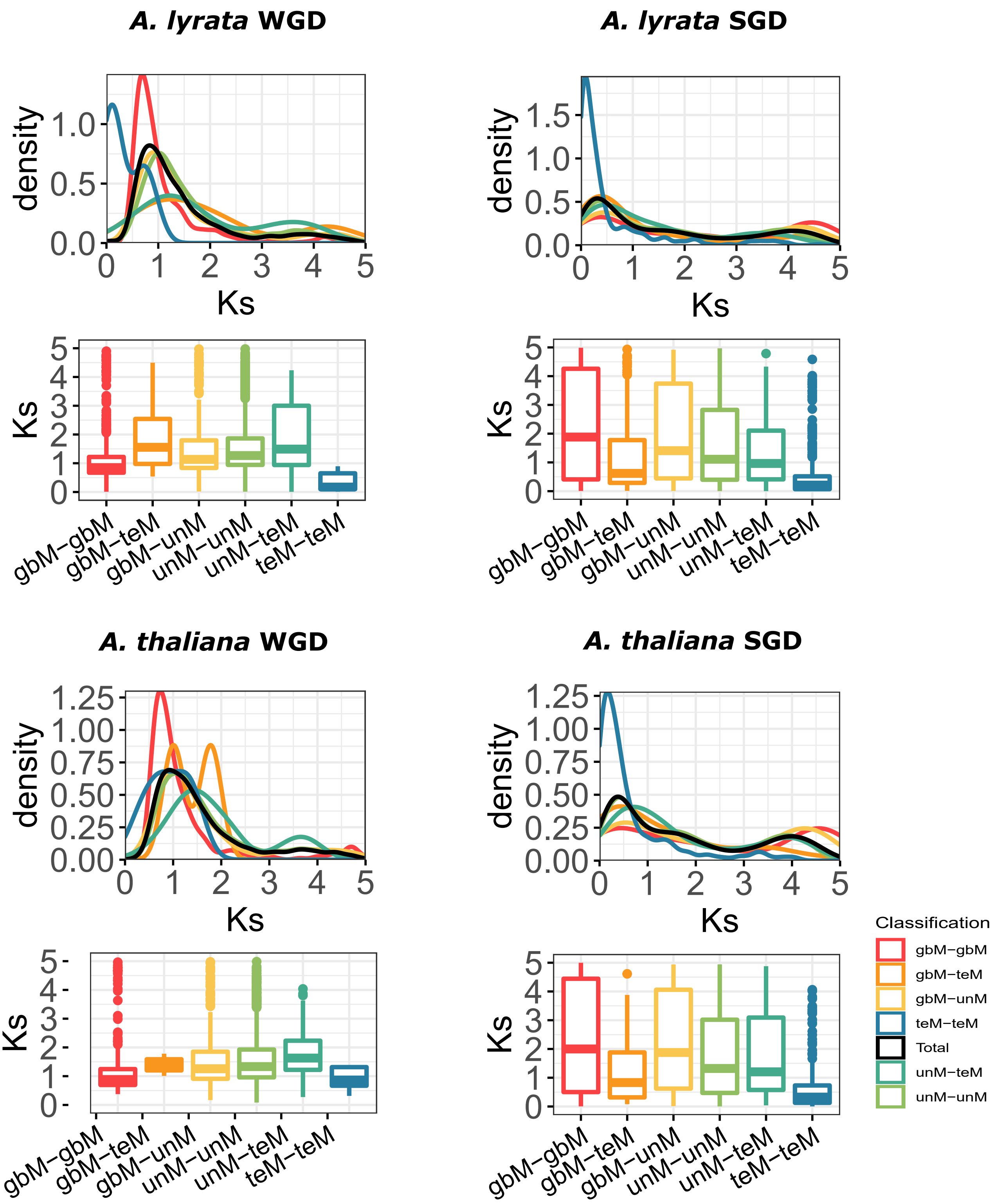

**Figure S7**: Distribution of genic methylation classified genes based on synonymous substitution (Ks) across different types of gene duplicate pairs. Whole-genome duplicates - WGD, Single-gene duplicates - SGD (combined data from tandem, proximal, translocated, and dispersed duplicates).

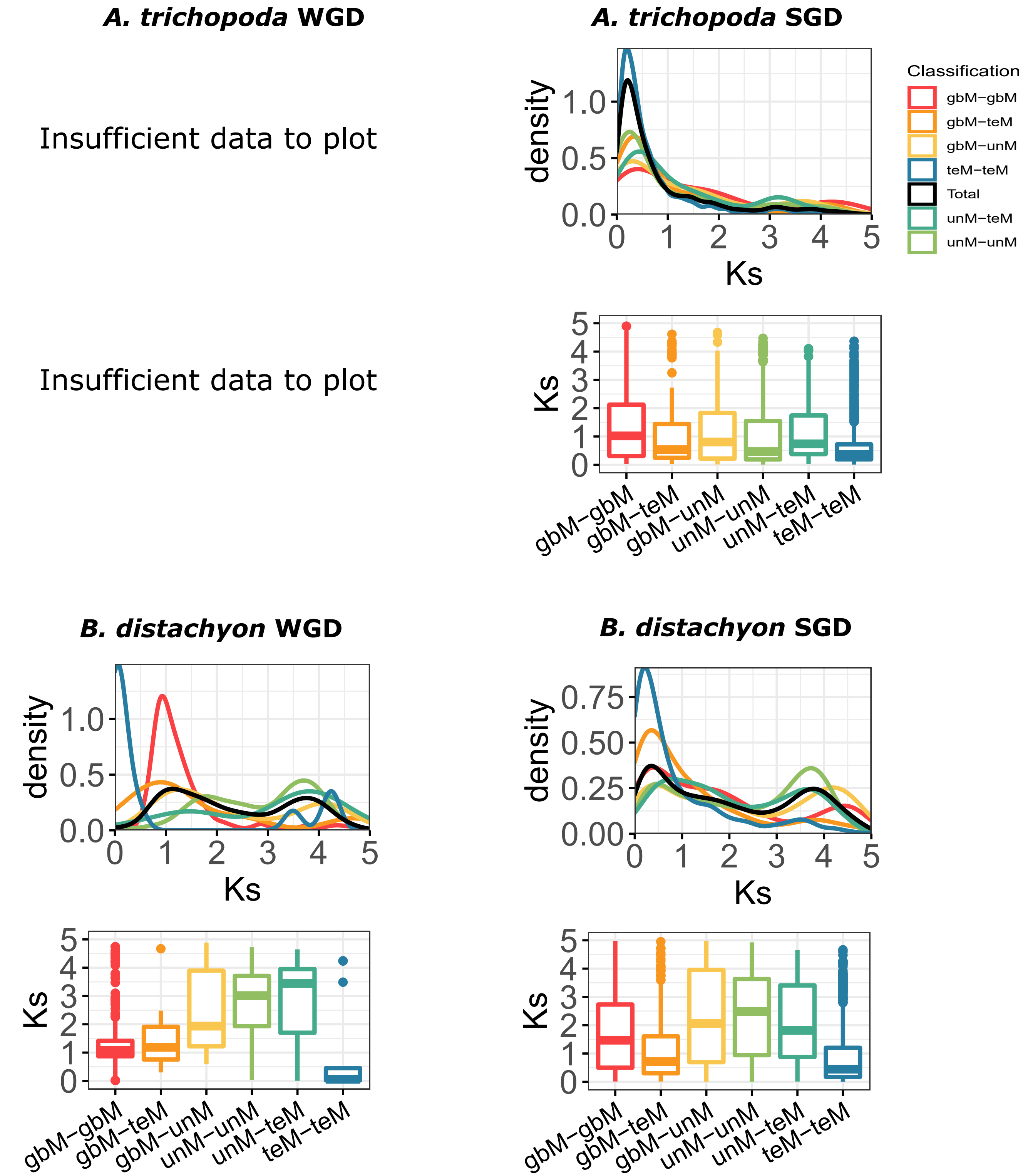

**Figure S7**: Distribution of genic methylation classified genes based on synonymous substitution (Ks) across different types of gene duplicate pairs. Whole-genome duplicates - WGD, Single-gene duplicates - SGD (combined data from tandem, proximal, translocated, and dispersed duplicates).

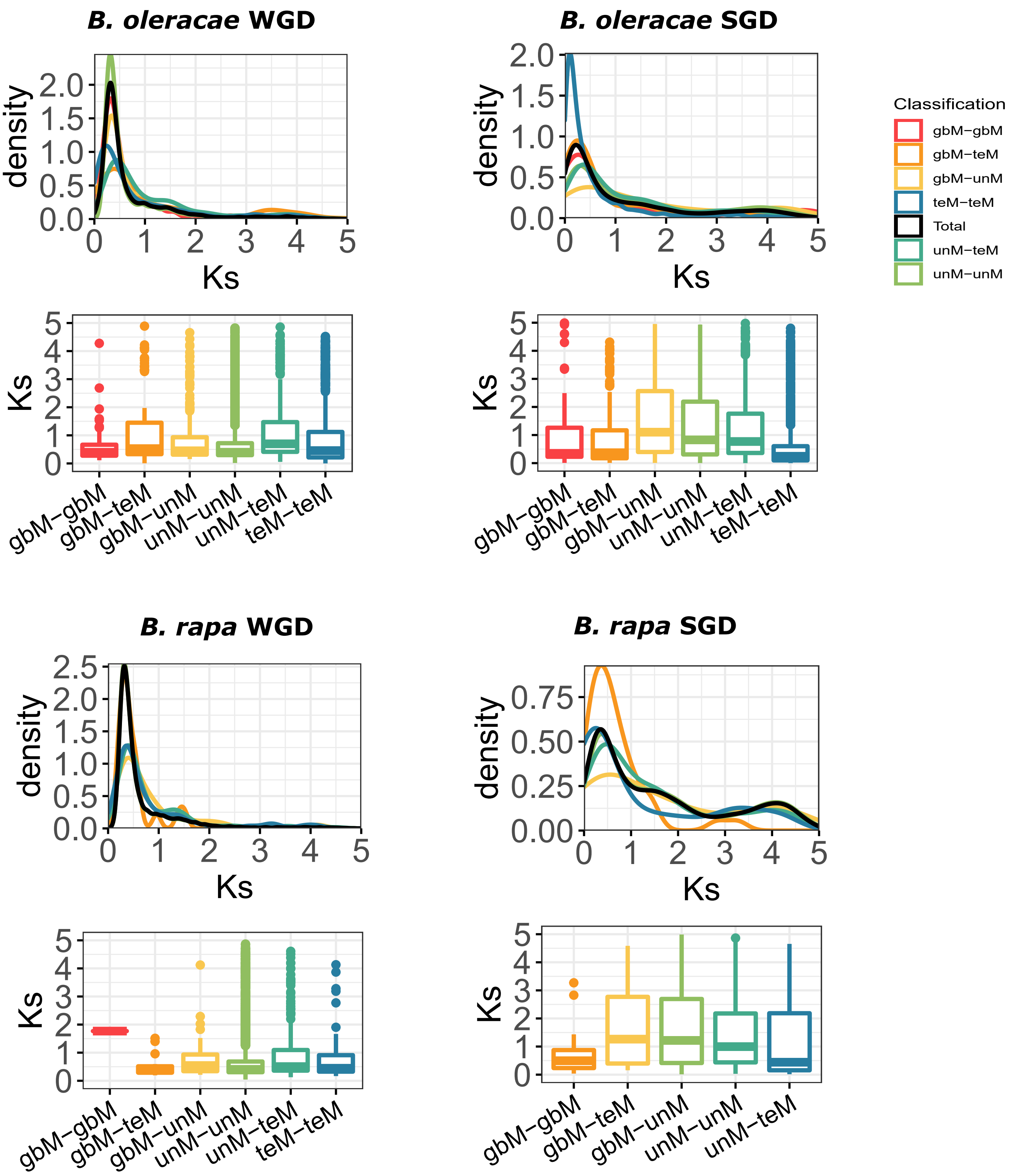

**Figure S7**: Distribution of genic methylation classified genes based on synonymous substitution (Ks) across different types of gene duplicate pairs. Whole-genome duplicates - WGD, Single-gene duplicates - SGD (combined data from tandem, proximal, translocated, and dispersed duplicates).

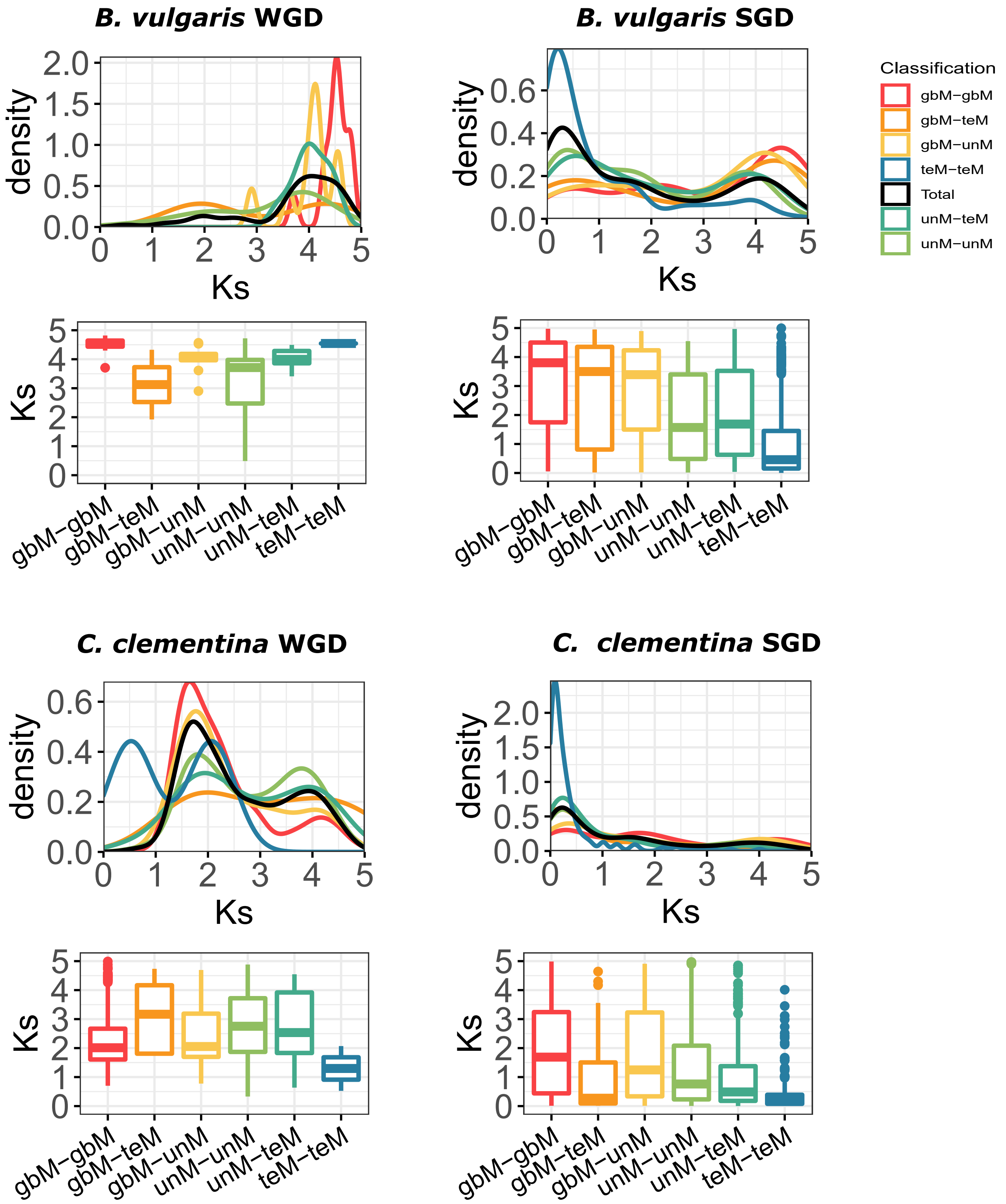

**Figure S7**: Distribution of genic methylation classified genes based on synonymous substitution (Ks) across different types of gene duplicate pairs. Whole-genome duplicates - WGD, Single-gene duplicates - SGD (combined data from tandem, proximal, translocated, and dispersed duplicates).

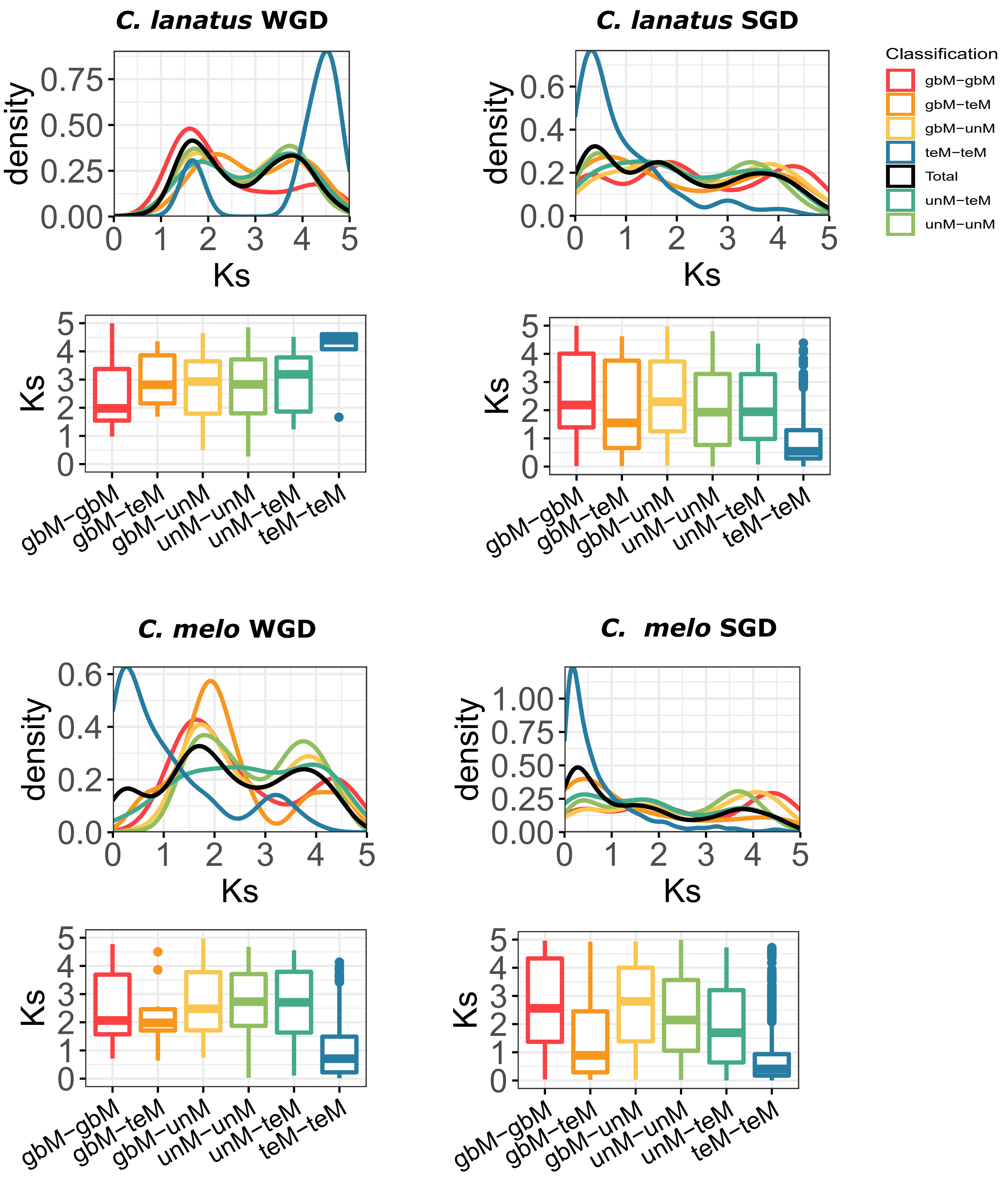

**Figure S7**: Distribution of genic methylation classified genes based on synonymous substitution (Ks) across different types of gene duplicate pairs. Whole-genome duplicates - WGD, Single-gene duplicates - SGD (combined data from tandem, proximal, translocated, and dispersed duplicates).

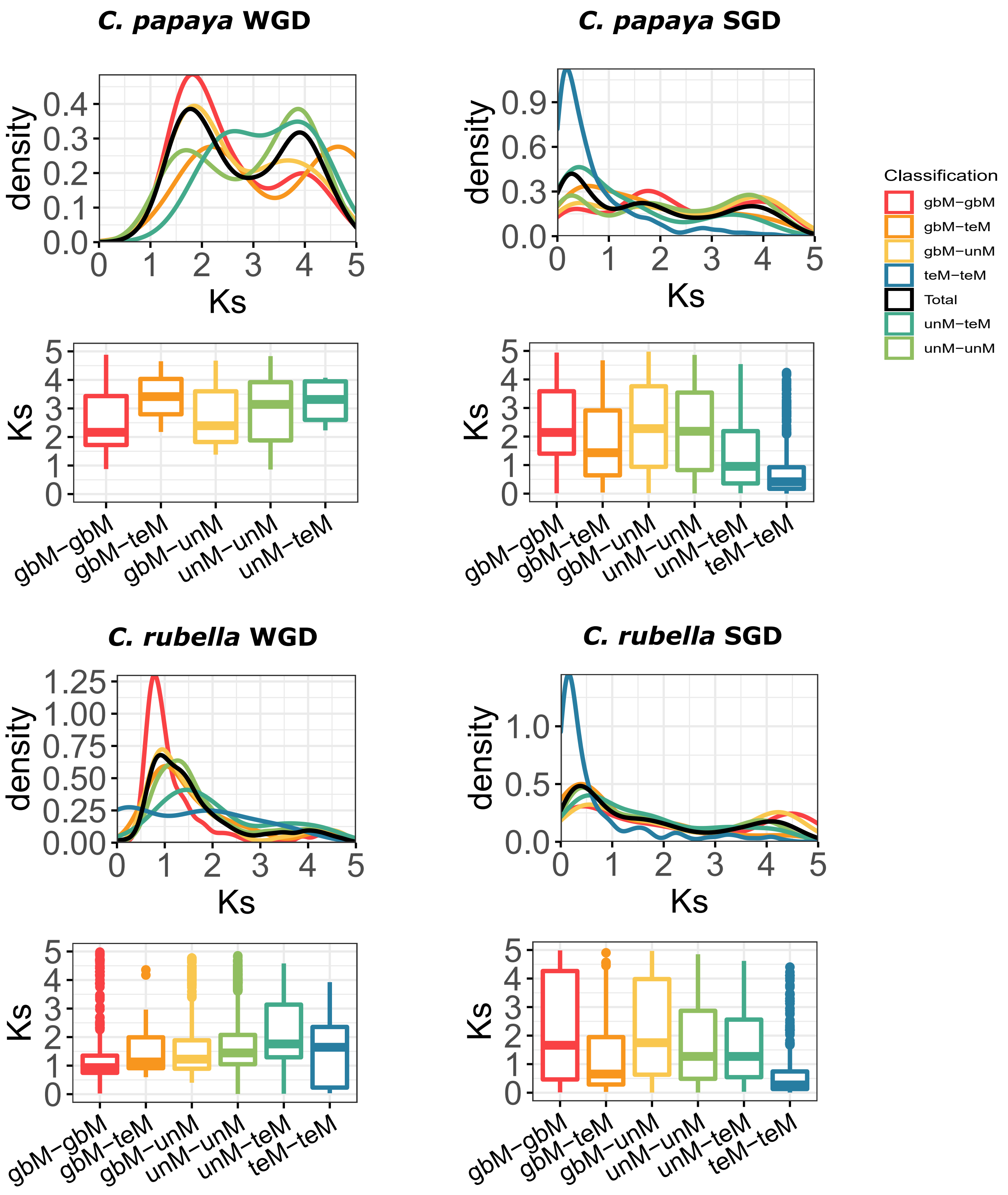

**Figure S7**: Distribution of genic methylation classified genes based on synonymous substitution (Ks) across different types of gene duplicate pairs. Whole-genome duplicates - WGD, Single-gene duplicates - SGD (combined data from tandem, proximal, translocated, and dispersed duplicates).

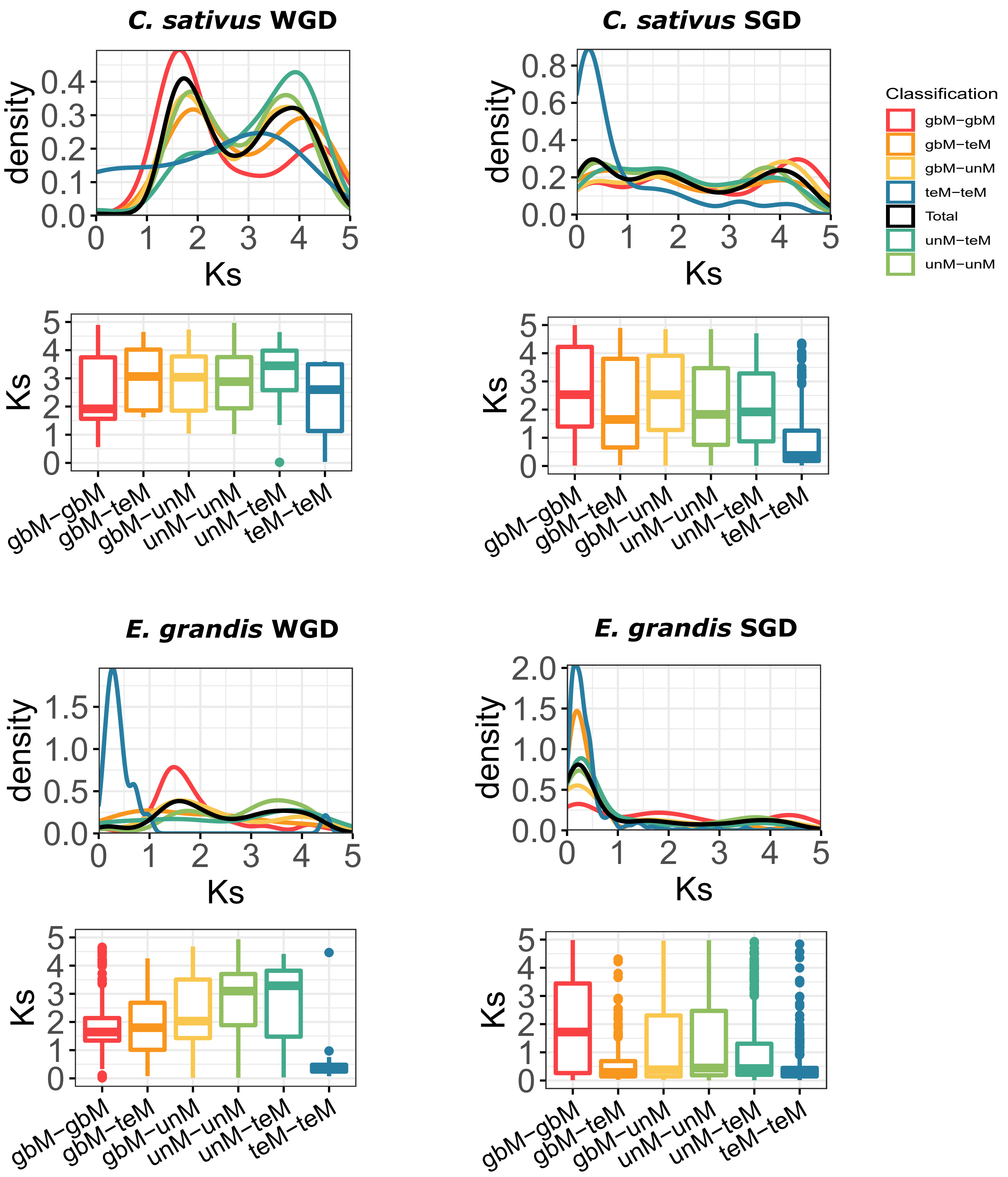

**Figure S7**: Distribution of genic methylation classified genes based on synonymous substitution (Ks) across different types of gene duplicate pairs. Whole-genome duplicates - WGD, Single-gene duplicates - SGD (combined data from tandem, proximal, translocated, and dispersed duplicates).

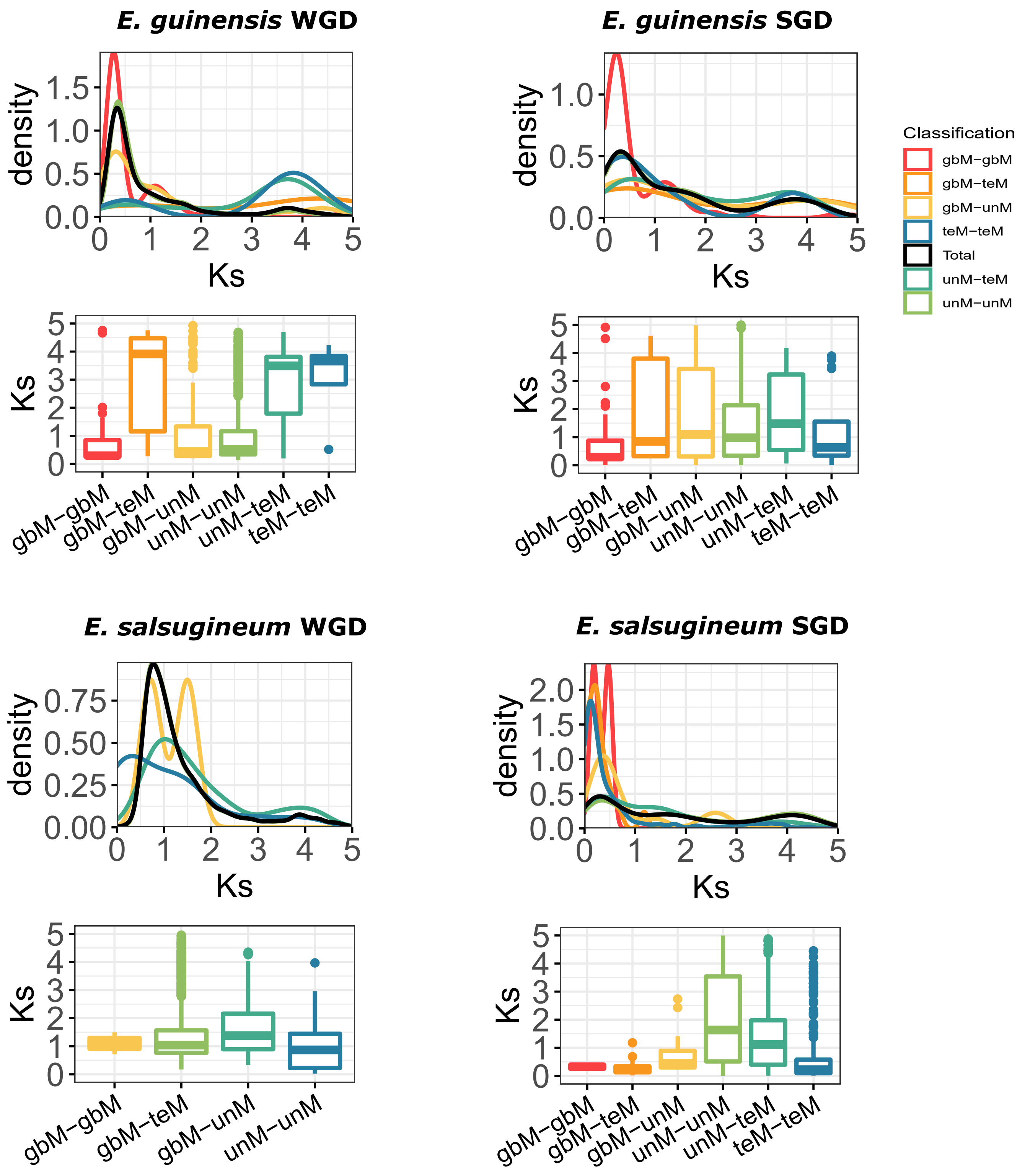

**Figure S7**: Distribution of genic methylation classified genes based on synonymous substitution (Ks) across different types of gene duplicate pairs. Whole-genome duplicates - WGD, Single-gene duplicates - SGD (combined data from tandem, proximal, translocated, and dispersed duplicates).

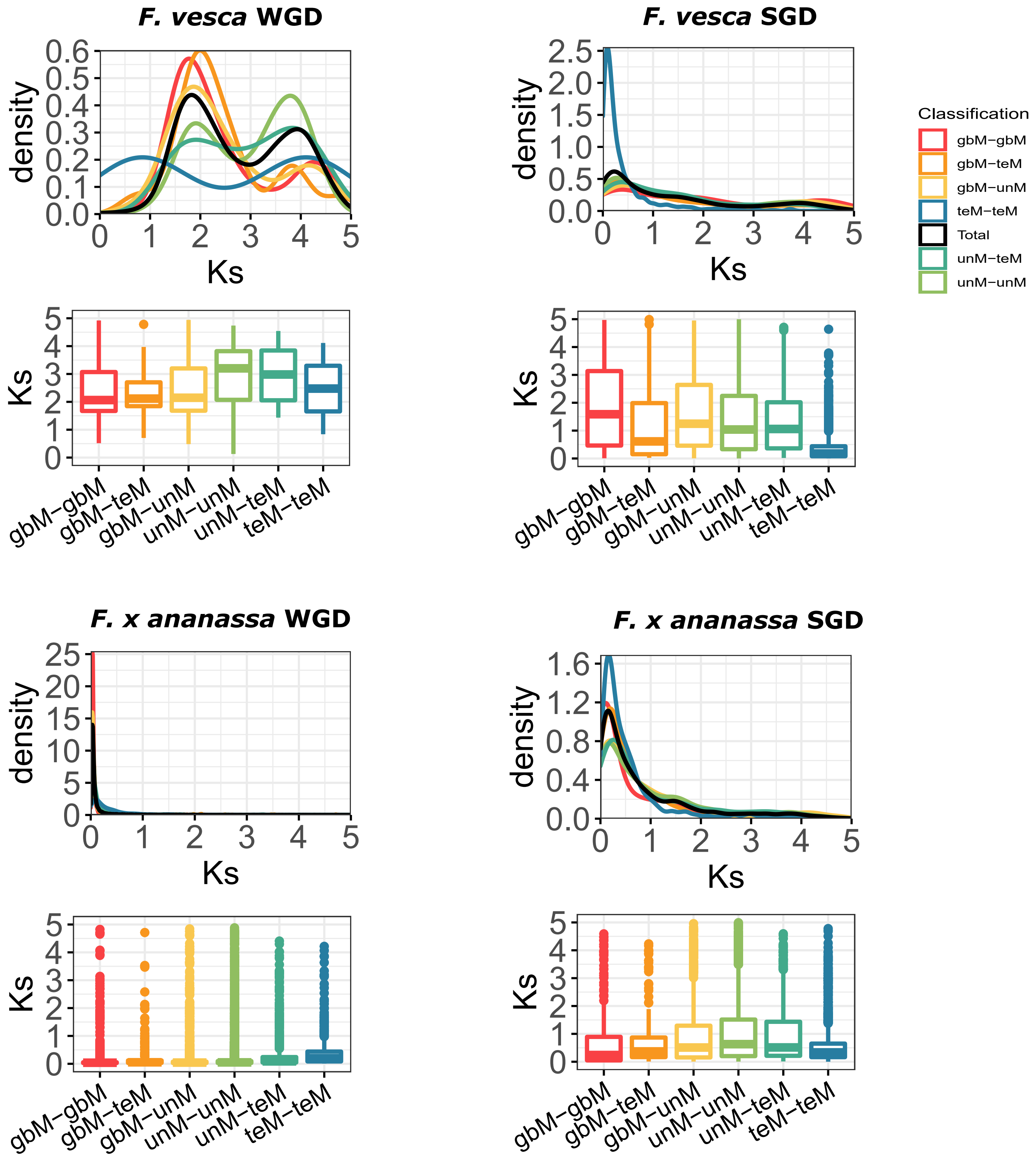

**Figure S7**: Distribution of genic methylation classified genes based on synonymous substitution (Ks) across different types of gene duplicate pairs. Whole-genome duplicates - WGD, Single-gene duplicates - SGD (combined data from tandem, proximal, translocated, and dispersed duplicates).

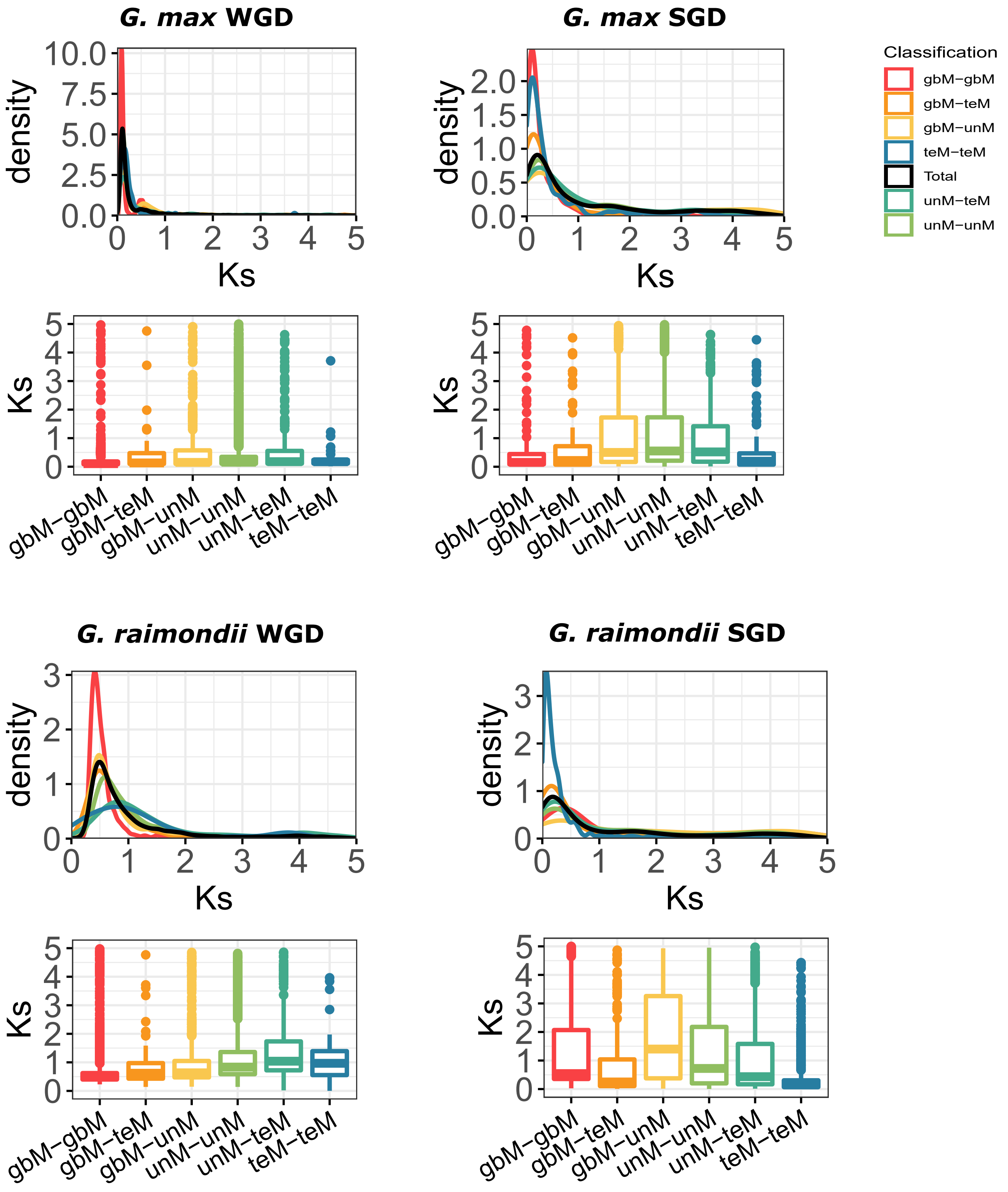

**Figure S7**: Distribution of genic methylation classified genes based on synonymous substitution (Ks) across different types of gene duplicate pairs. Whole-genome duplicates - WGD, Single-gene duplicates - SGD (combined data from tandem, proximal, translocated, and dispersed duplicates).

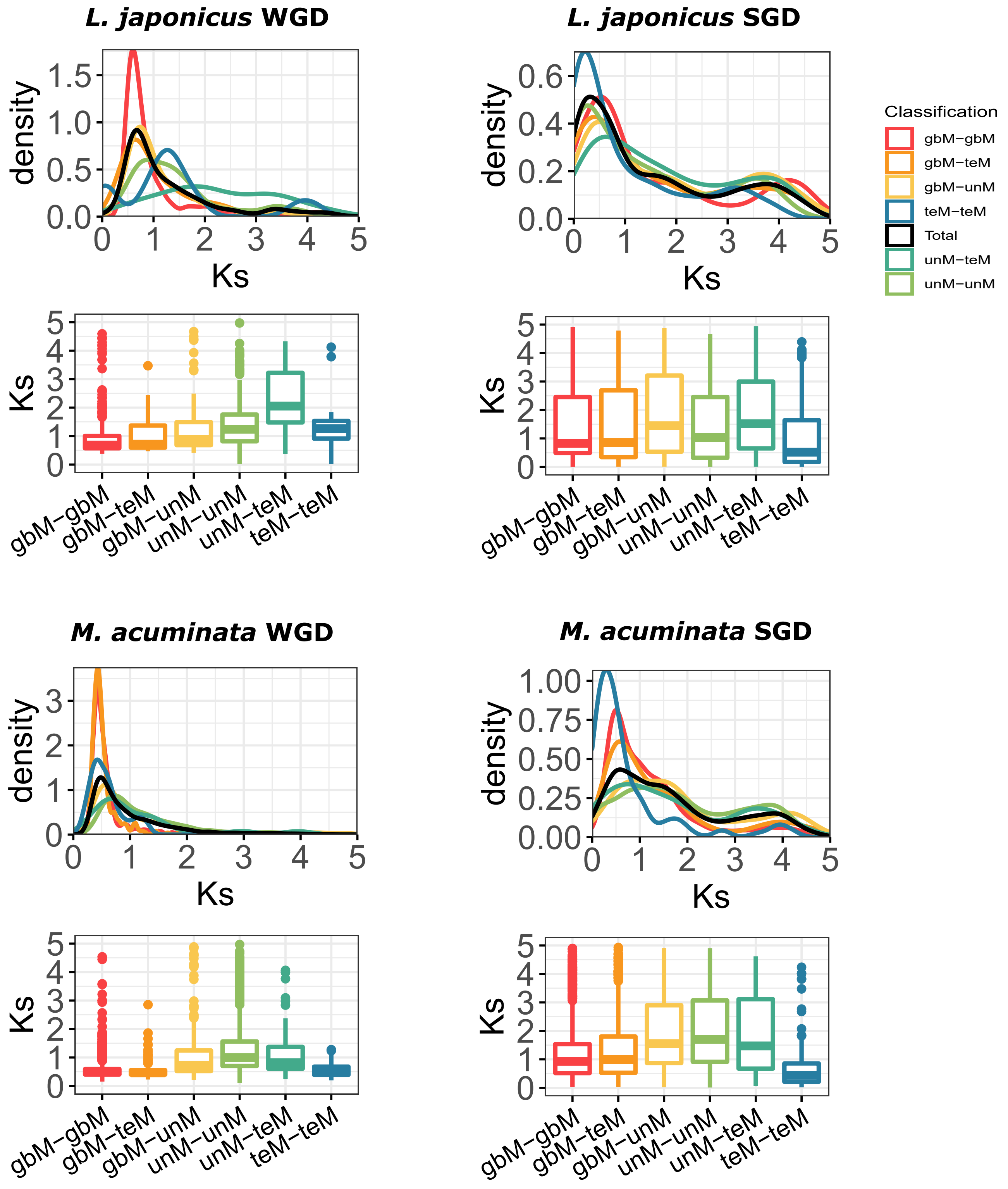

**Figure S7**: Distribution of genic methylation classified genes based on synonymous substitution (Ks) across different types of gene duplicate pairs. Whole-genome duplicates - WGD, Single-gene duplicates - SGD (combined data from tandem, proximal, translocated, and dispersed duplicates).

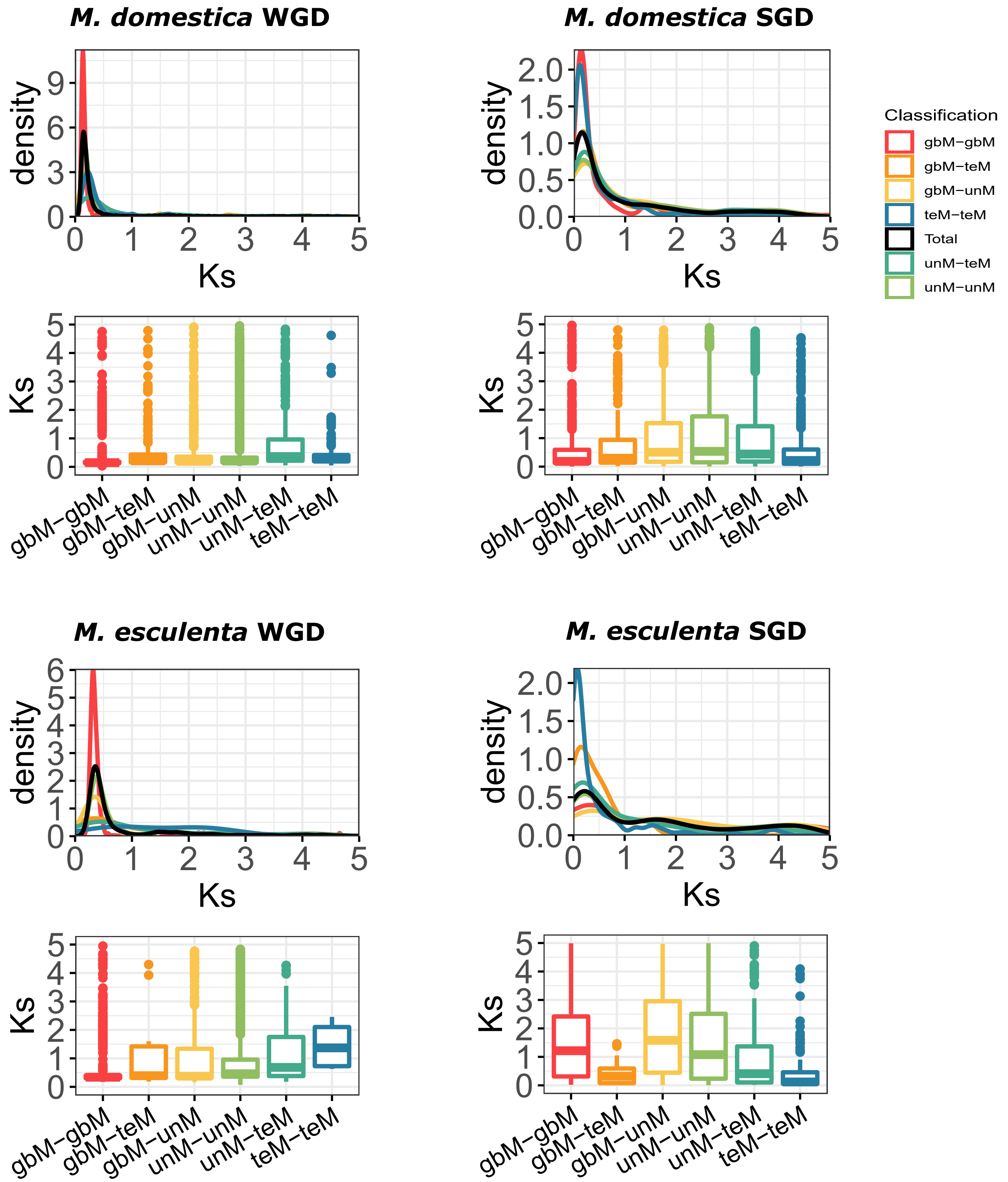

**Figure S7**: Distribution of genic methylation classified genes based on synonymous substitution (Ks) across different types of gene duplicate pairs. Whole-genome duplicates - WGD, Single-gene duplicates - SGD (combined data from tandem, proximal, translocated, and dispersed duplicates).

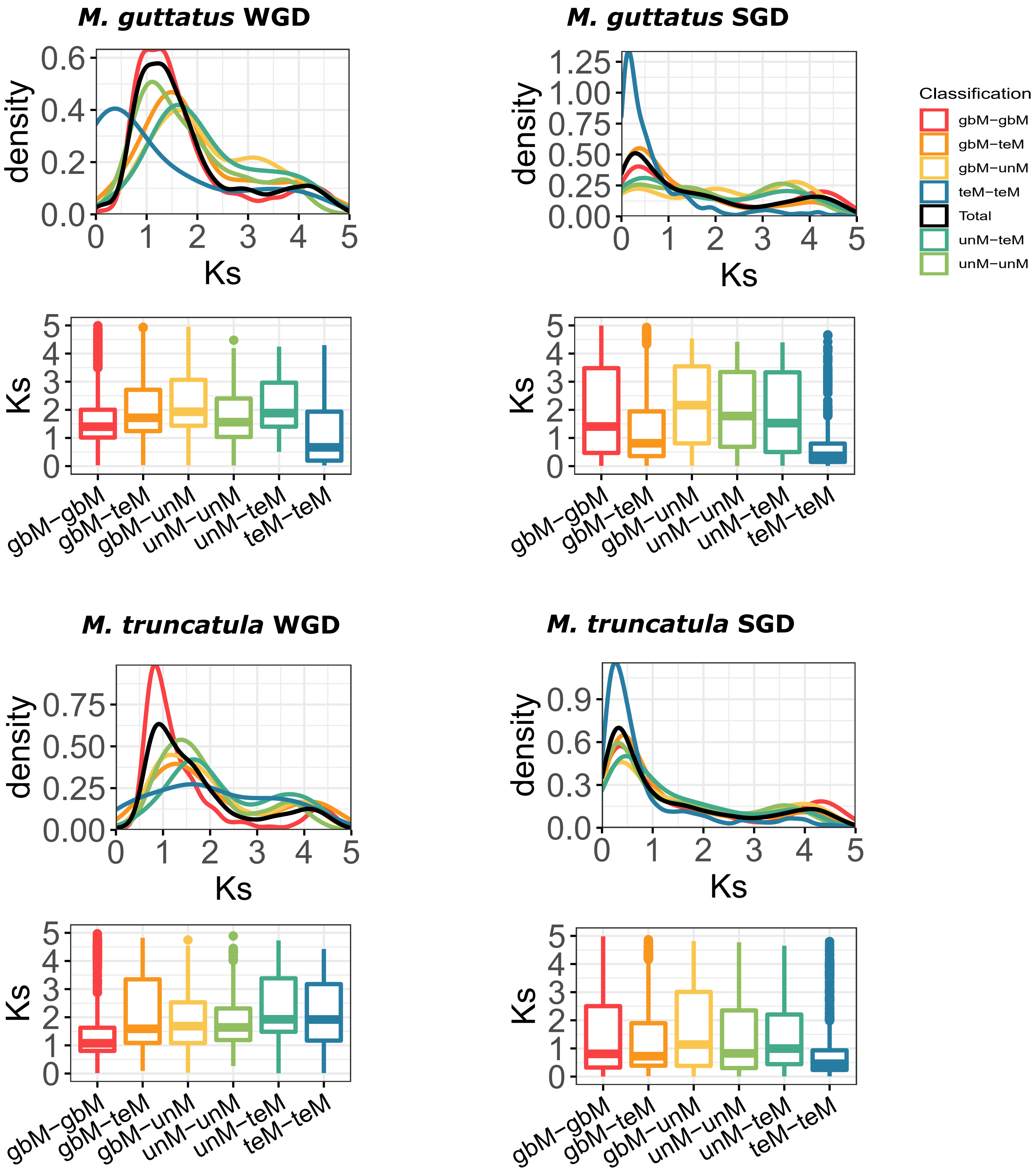

**Figure S7**: Distribution of genic methylation classified genes based on synonymous substitution (Ks) across different types of gene duplicate pairs. Whole-genome duplicates - WGD, Single-gene duplicates - SGD (combined data from tandem, proximal, translocated, and dispersed duplicates).

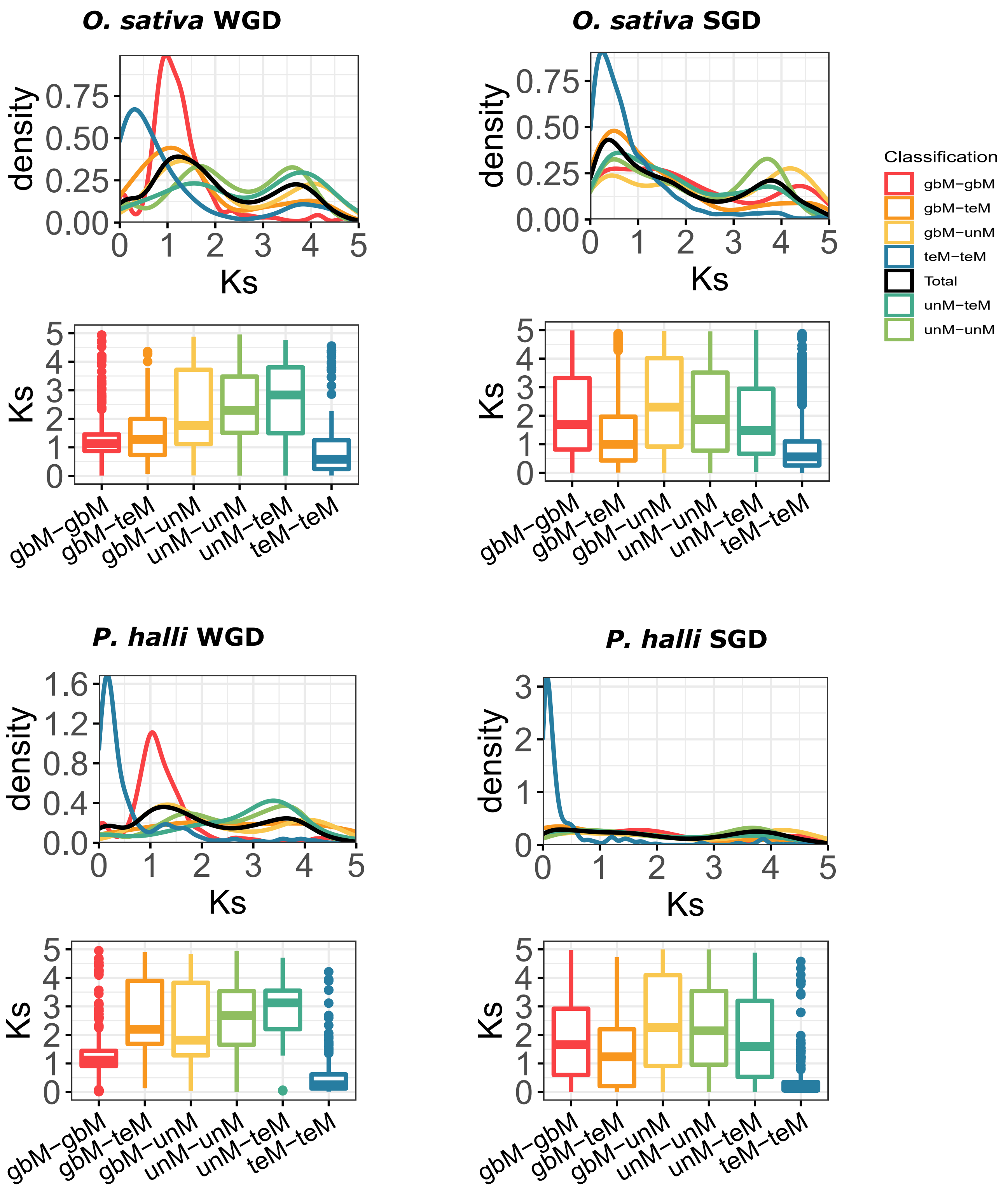

**Figure S7**: Distribution of genic methylation classified genes based on synonymous substitution (Ks) across different types of gene duplicate pairs. Whole-genome duplicates - WGD, Single-gene duplicates - SGD (combined data from tandem, proximal, translocated, and dispersed duplicates).

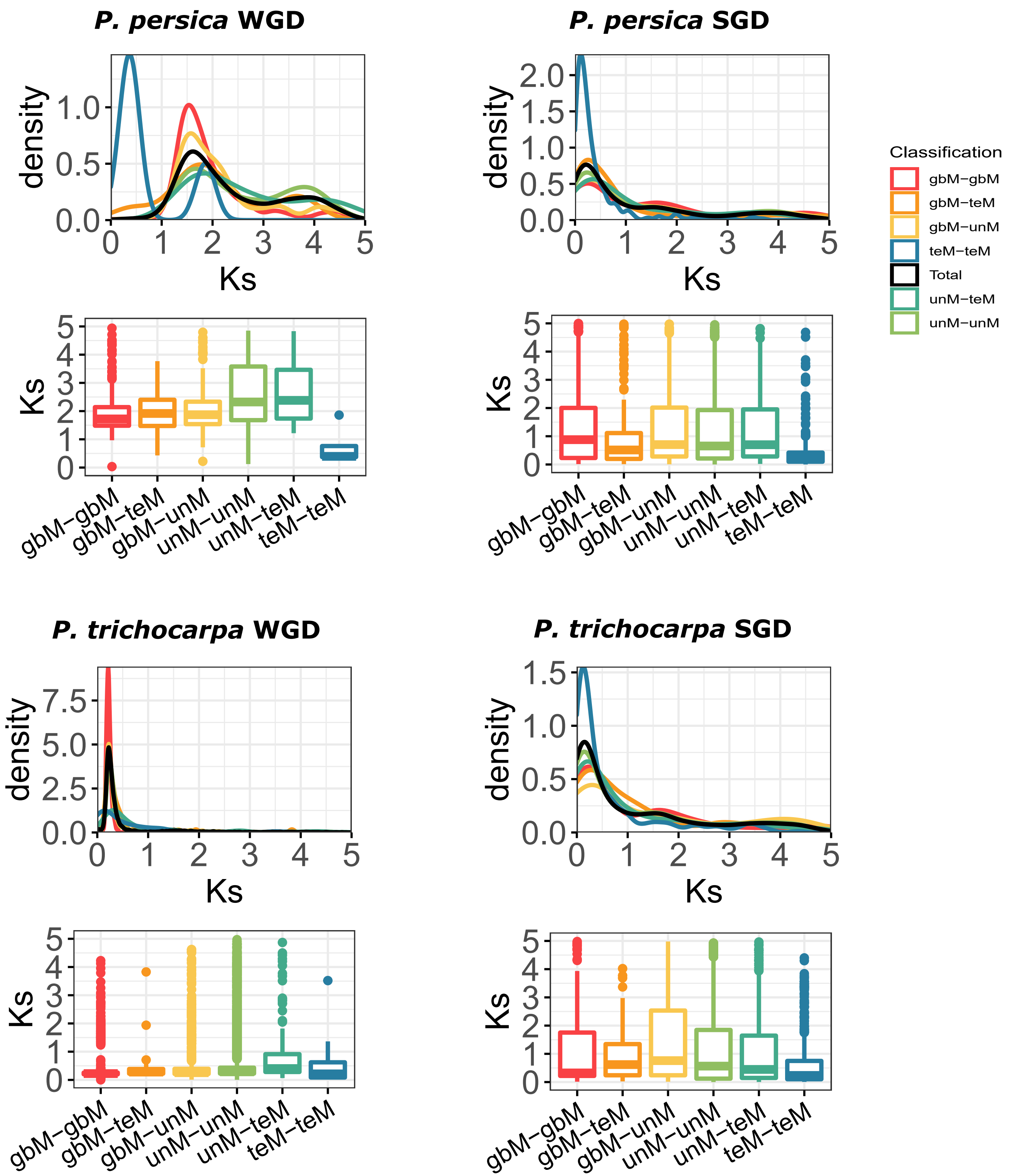

**Figure S7**: Distribution of genic methylation classified genes based on synonymous substitution (Ks) across different types of gene duplicate pairs. Whole-genome duplicates - WGD, Single-gene duplicates - SGD (combined data from tandem, proximal, translocated, and dispersed duplicates).

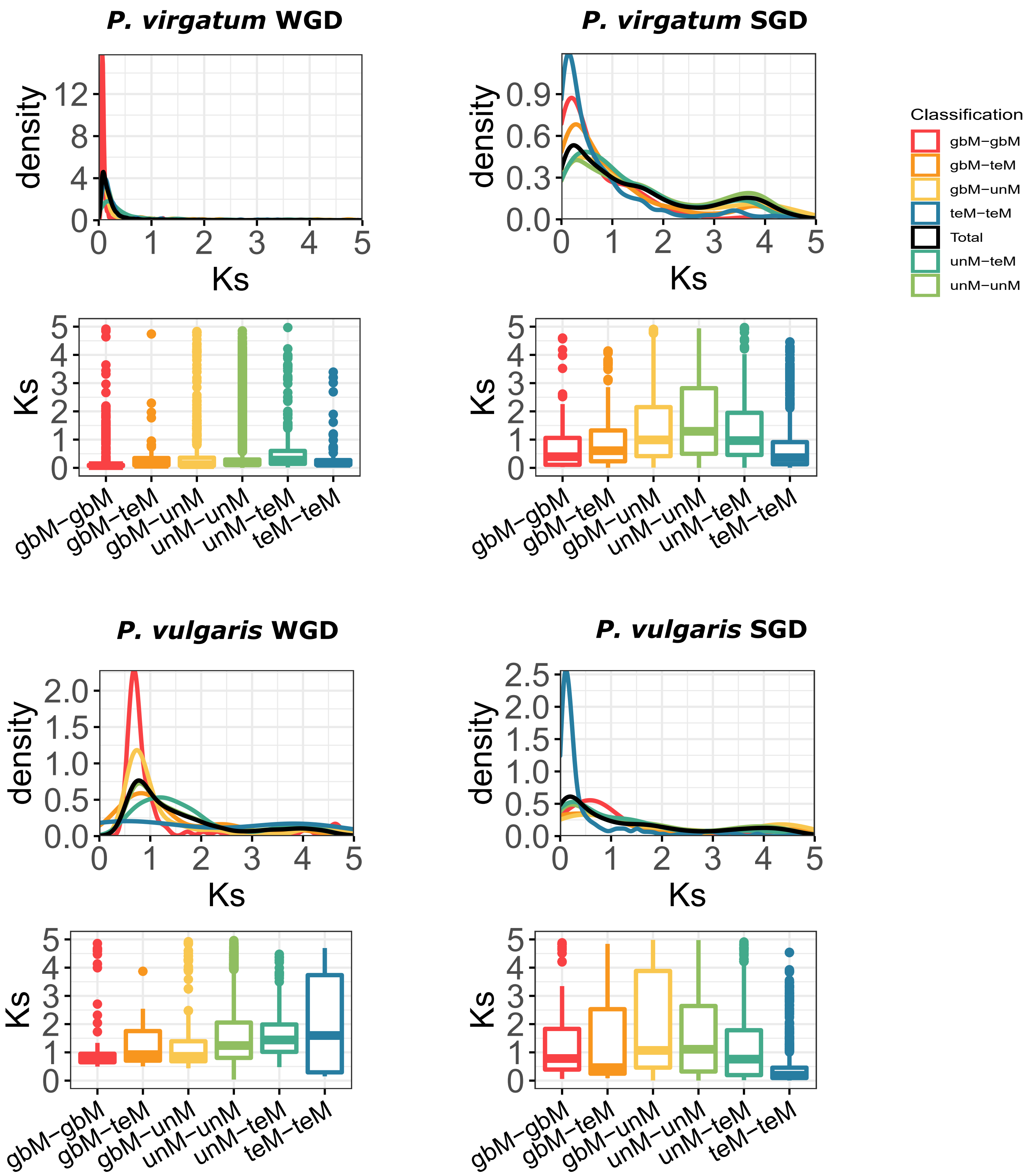

**Figure S7**: Distribution of genic methylation classified genes based on synonymous substitution (Ks) across different types of gene duplicate pairs. Whole-genome duplicates - WGD, Single-gene duplicates - SGD (combined data from tandem, proximal, translocated, and dispersed duplicates).

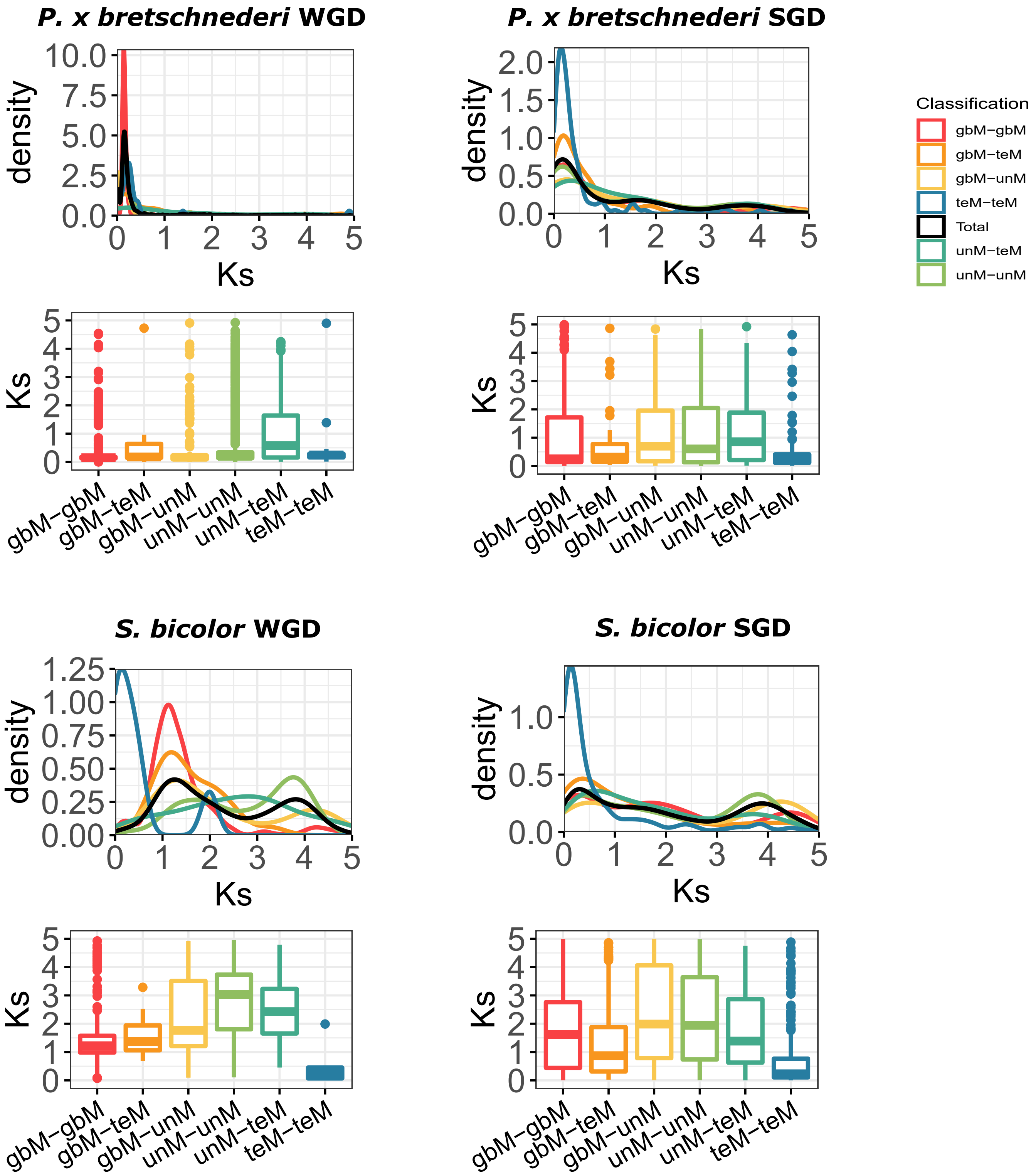

**Figure S7**: Distribution of genic methylation classified genes based on synonymous substitution (Ks) across different types of gene duplicate pairs. Whole-genome duplicates - WGD, Single-gene duplicates - SGD (combined data from tandem, proximal, translocated, and dispersed duplicates).

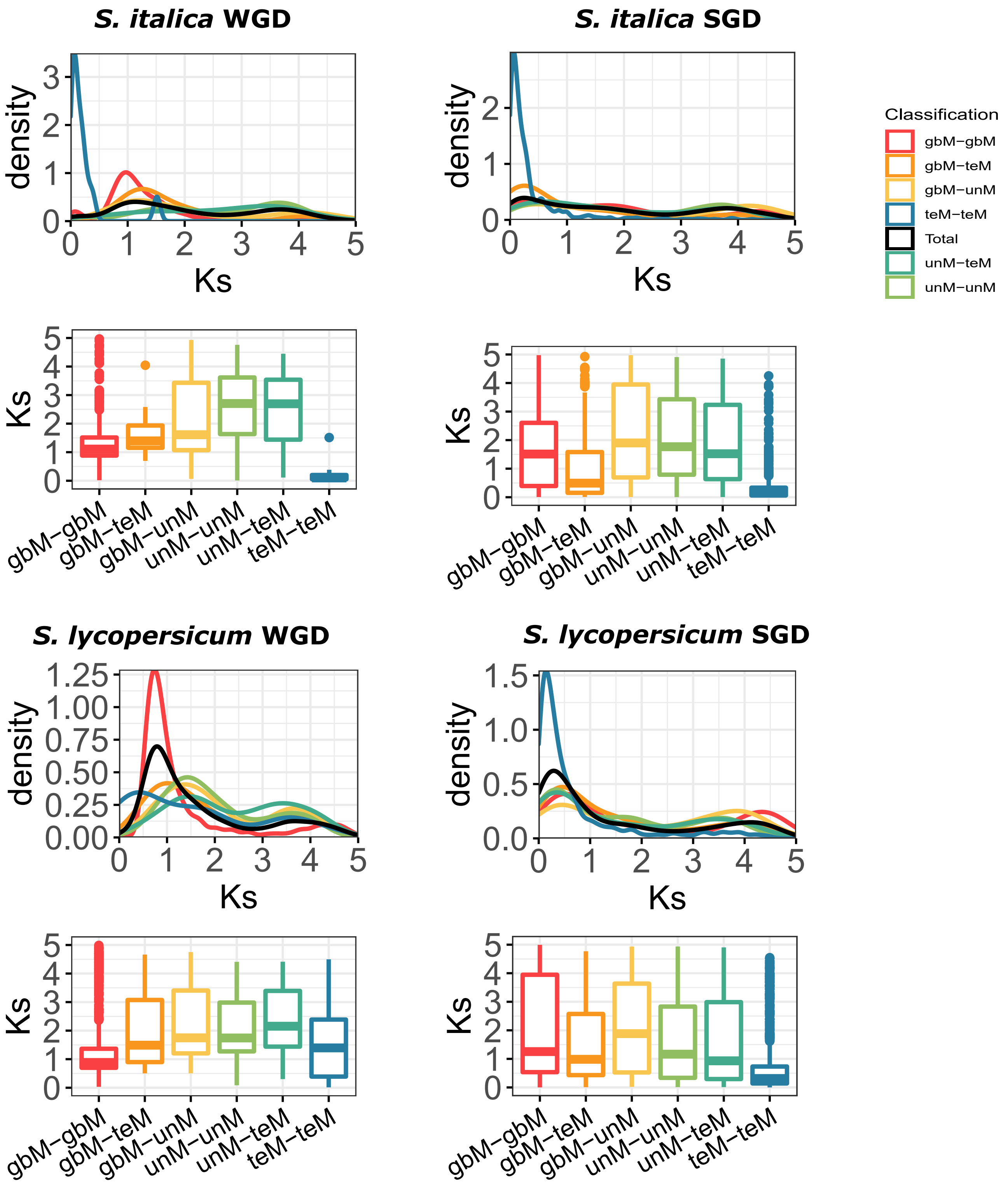

**Figure S7**: Distribution of genic methylation classified genes based on synonymous substitution (Ks) across different types of gene duplicate pairs. Whole-genome duplicates - WGD, Single-gene duplicates - SGD (combined data from tandem, proximal, translocated, and dispersed duplicates).

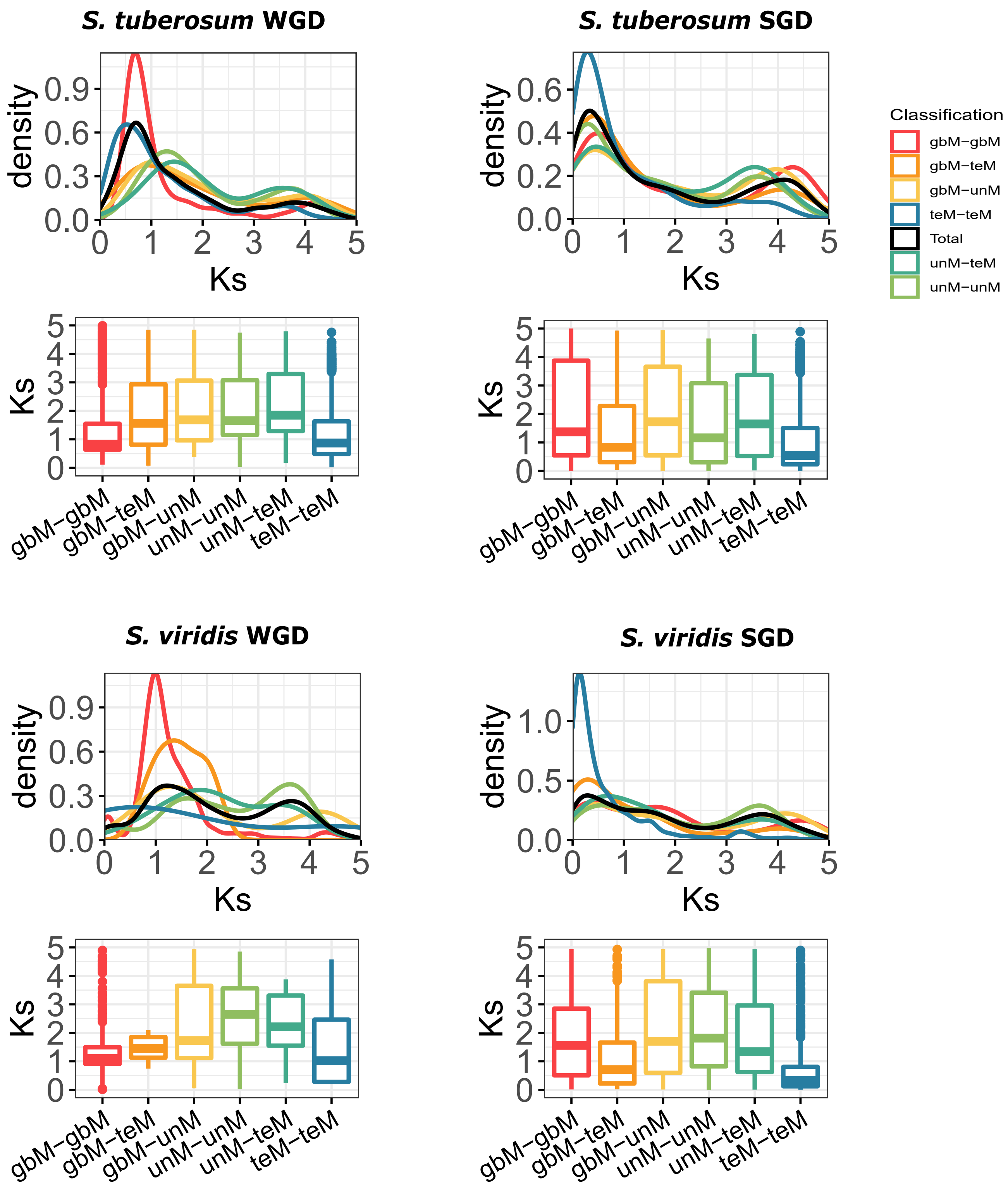

**Figure S7**: Distribution of genic methylation classified genes based on synonymous substitution (Ks) across different types of gene duplicate pairs. Whole-genome duplicates - WGD, Single-gene duplicates - SGD (combined data from tandem, proximal, translocated, and dispersed duplicates).

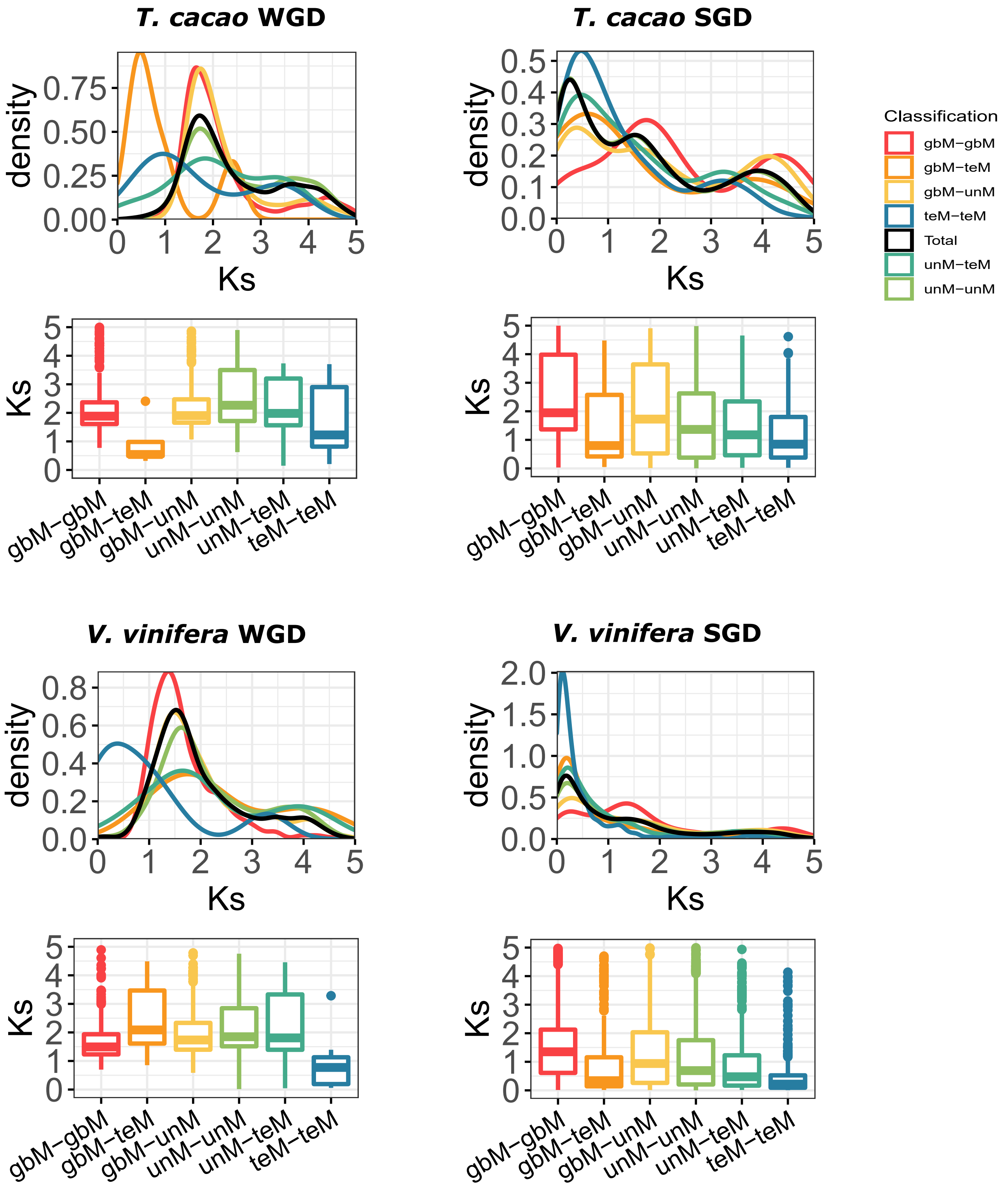

**Figure S7**: Distribution of genic methylation classified genes based on synonymous substitution (Ks) across different types of gene duplicate pairs. Whole-genome duplicates - WGD, Single-gene duplicates - SGD (combined data from tandem, proximal, translocated, and dispersed duplicates).

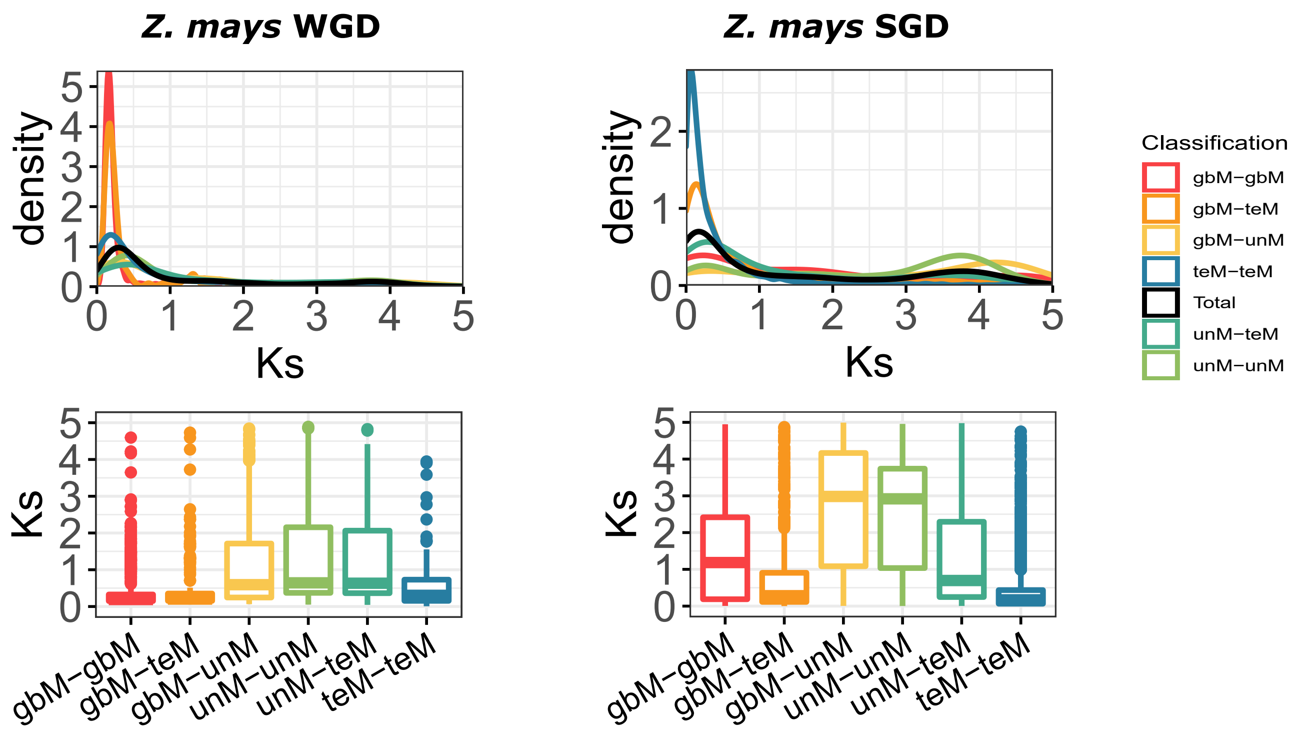

**Figure S8:** The percentage of gene copies in each genic methylation class for translocated genes that have duplicated during that 'epoch' since divergence from the species on the x-axis. For example, in *A. duranensis* translocated genes that have duplicated since *A. duranensis* diverged from *A. ipaensis* are shown on the x-axis under *A. ipaensis*. Those shown under *G. max*, duplicated in the period since the common ancestor of *A. duranensis* and *A. ipaensis* diverged from their common ancestor with *G. max*, but before the divergence of *A. duranensis* and *A. ipaensis.* Horizontal dotted lines indicate the percentage of each genic methylation class in all translocated duplicates. Bars above this line indicate enrichment, below this line depletion.

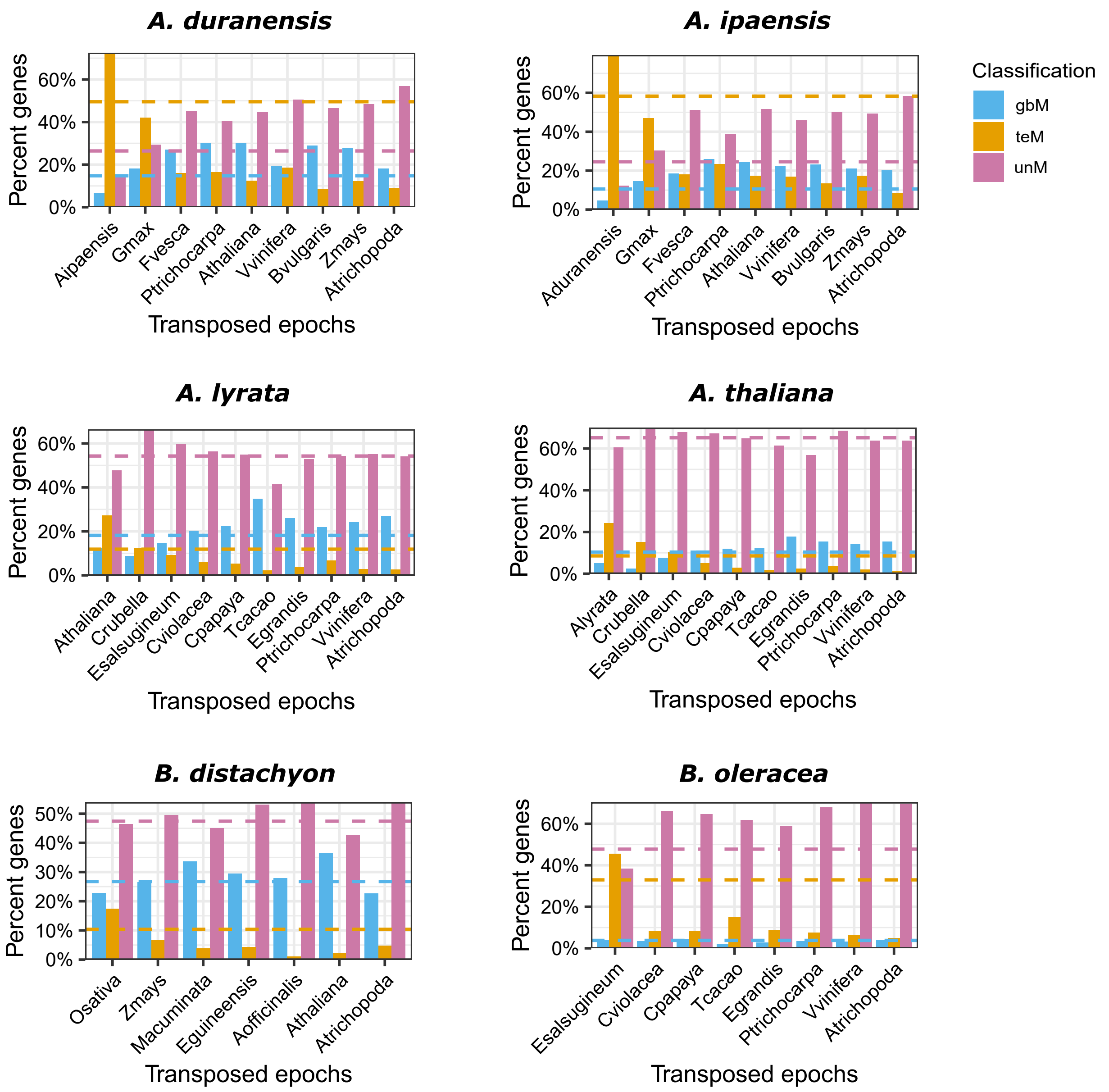

**Figure S8:** The percentage of gene copies in each genic methylation class for translocated genes that have duplicated during that 'epoch' since divergence from the species on the x-axis. For example, in *A. duranensis* translocated genes that have duplicated since *A. duranensis* diverged from *A. ipaensis* are shown on the x-axis under *A. ipaensis*. Those shown under *G. max*, duplicated in the period since the common ancestor of *A. duranensis* and *A. ipaensis* diverged from their common ancestor with *G. max*, but before the divergence of *A. duranensis* and *A. ipaensis.* Horizontal dotted lines indicate the percentage of each genic methylation class in all translocated duplicates. Bars above this line indicate enrichment, below this line depletion.

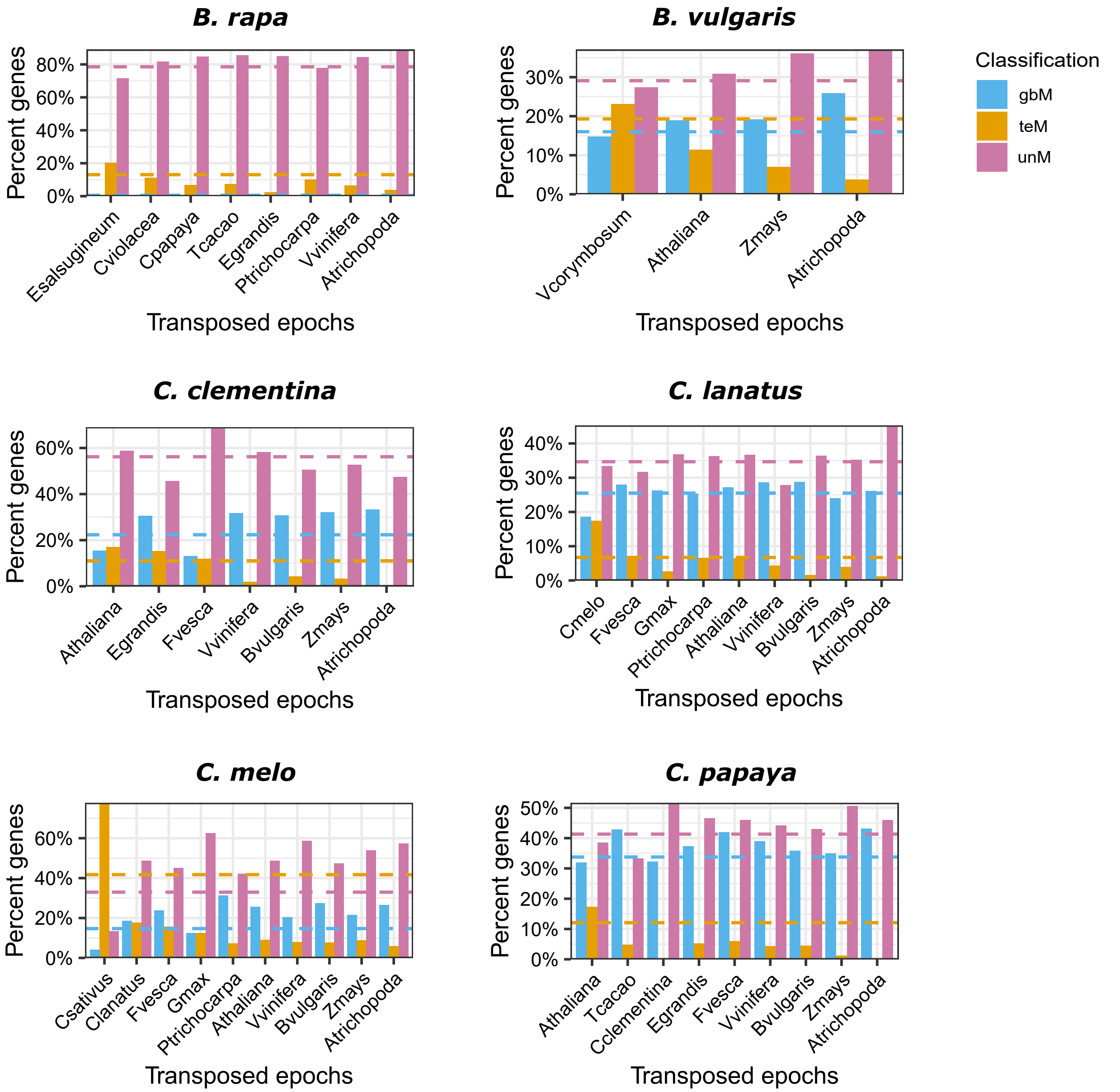

**Figure S8:** The percentage of gene copies in each genic methylation class for translocated genes that have duplicated during that ‘epoch’ since divergence from the species on the x-axis. For example, in *A. duranensis* translocated genes that have duplicated since *A. duranensis* diverged from *A. ipaensis* are shown on the x-axis under *A. ipaensis*. Those shown under *G. max*, duplicated in the period since the common ancestor of *A. duranensis* and *A. ipaensis* diverged from their common ancestor with *G. max*, but before the divergence of *A. duranensis* and *A. ipaensis.* Horizontal dotted lines indicate the percentage of each genic methylation class in all translocated duplicates. Bars above this line indicate enrichment, below this line depletion.

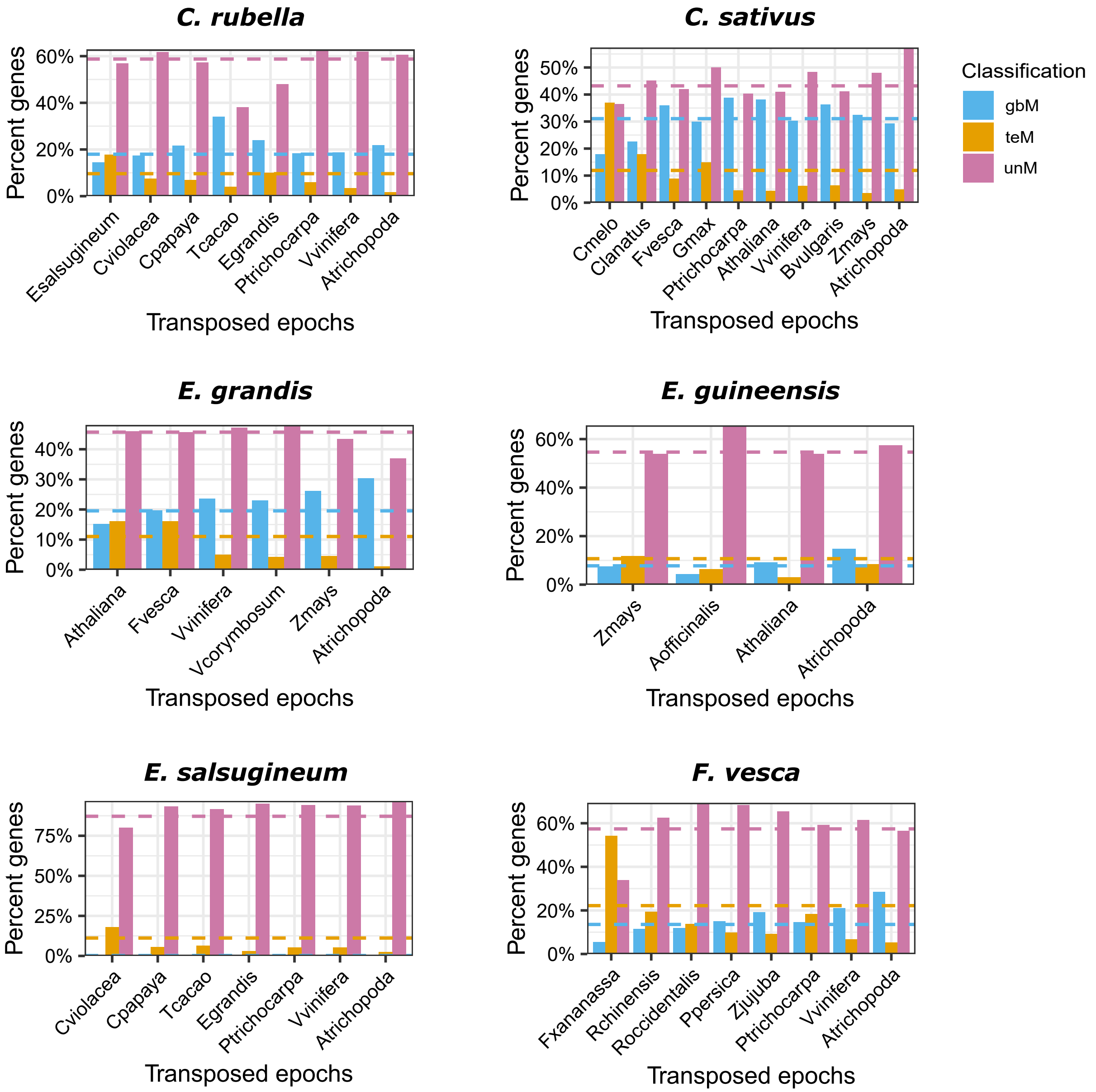

**Figure S8:** The percentage of gene copies in each genic methylation class for translocated genes that have duplicated during that ‘epoch’ since divergence from the species on the x-axis. For example, in *A. duranensis* translocated genes that have duplicated since *A. duranensis* diverged from *A. ipaensis* are shown on the x-axis under *A. ipaensis*. Those shown under *G. max*, duplicated in the period since the common ancestor of *A. duranensis* and *A. ipaensis* diverged from their common ancestor with *G. max*, but before the divergence of *A. duranensis* and *A. ipaensis.* Horizontal dotted lines indicate the percentage of each genic methylation class in all translocated duplicates. Bars above this line indicate enrichment, below this line depletion.

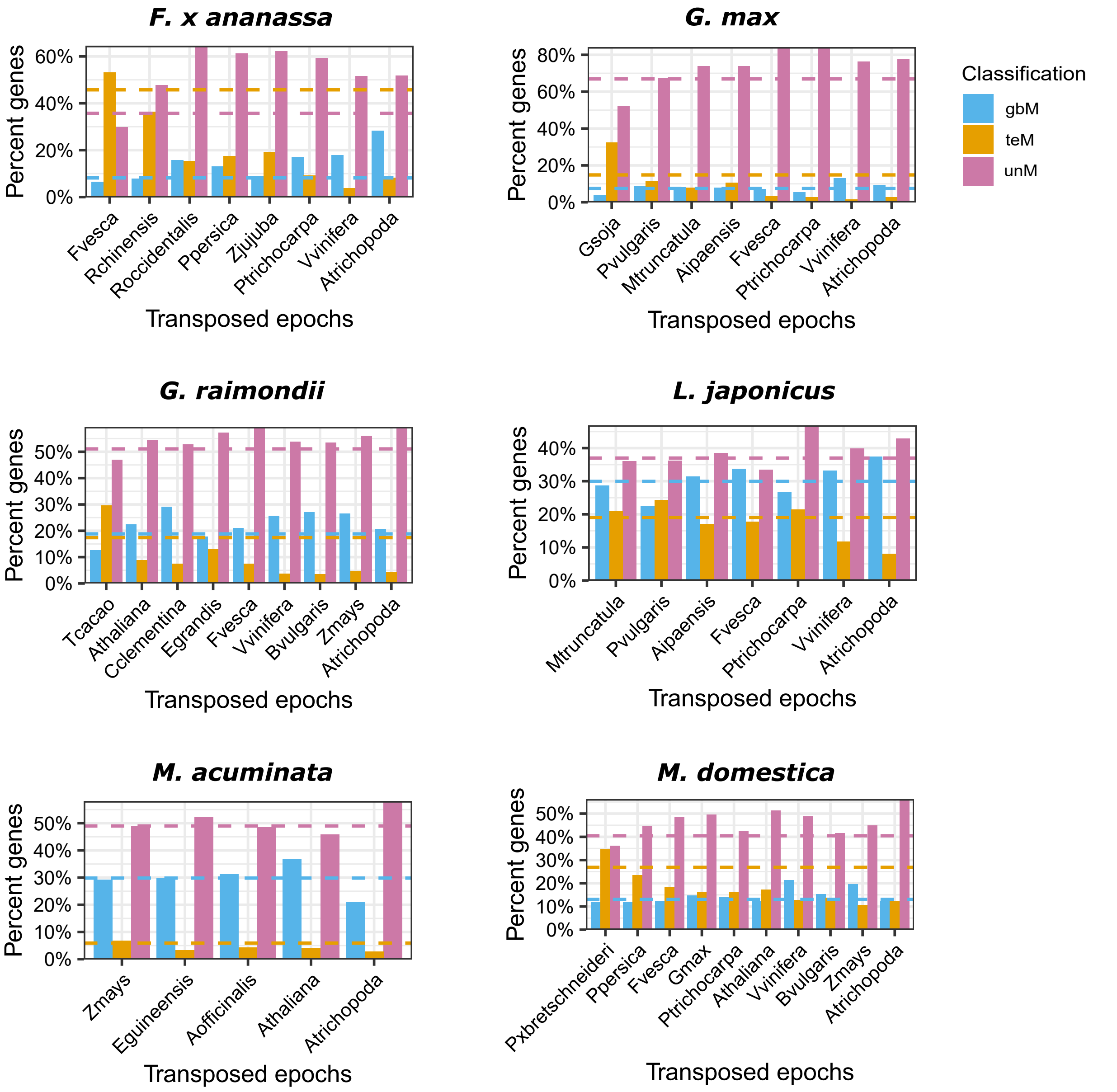

**Figure S8:** The percentage of gene copies in each genic methylation class for translocated genes that have duplicated during that 'epoch' since divergence from the species on the x-axis. For example, in *A. duranensis* translocated genes that have duplicated since *A. duranensis* diverged from *A. ipaensis* are shown on the x-axis under *A. ipaensis*. Those shown under *G. max*, duplicated in the period since the common ancestor of *A. duranensis* and *A. ipaensis* diverged from their common ancestor with *G. max*, but before the divergence of *A. duranensis* and *A. ipaensis.* Horizontal dotted lines indicate the percentage of each genic methylation class in all translocated duplicates. Bars above this line indicate enrichment, below this line depletion.

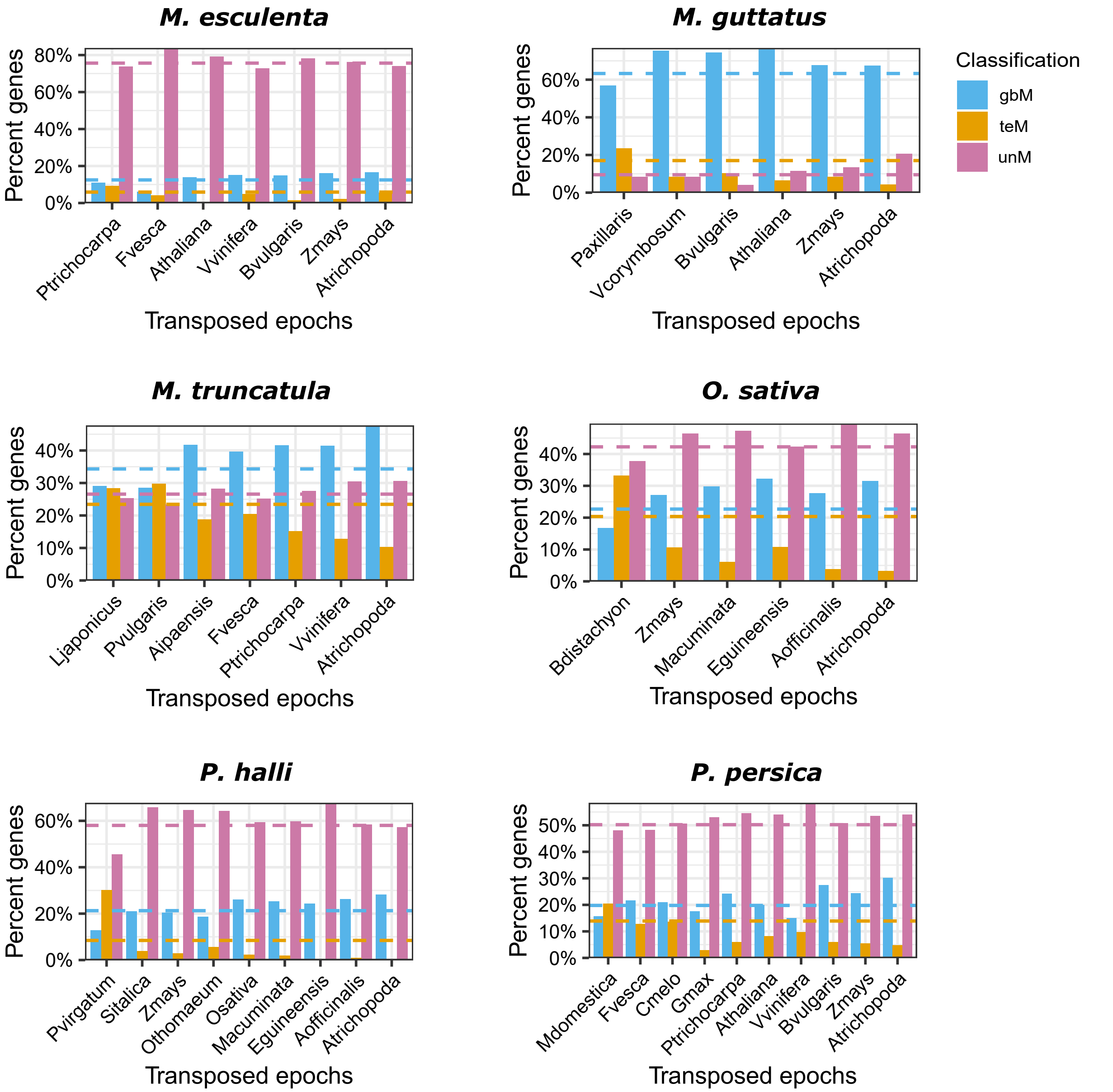

**Figure S8:** The percentage of gene copies in each genic methylation class for translocated genes that have duplicated during that 'epoch' since divergence from the species on the x-axis. For example, in *A. duranensis* translocated genes that have duplicated since *A. duranensis* diverged from *A. ipaensis* are shown on the x-axis under *A. ipaensis*. Those shown under *G. max*, duplicated in the period since the common ancestor of *A. duranensis* and *A. ipaensis* diverged from their common ancestor with *G. max*, but before the divergence of *A. duranensis* and *A. ipaensis.* Horizontal dotted lines indicate the percentage of each genic methylation class in all translocated duplicates. Bars above this line indicate enrichment, below this line depletion.

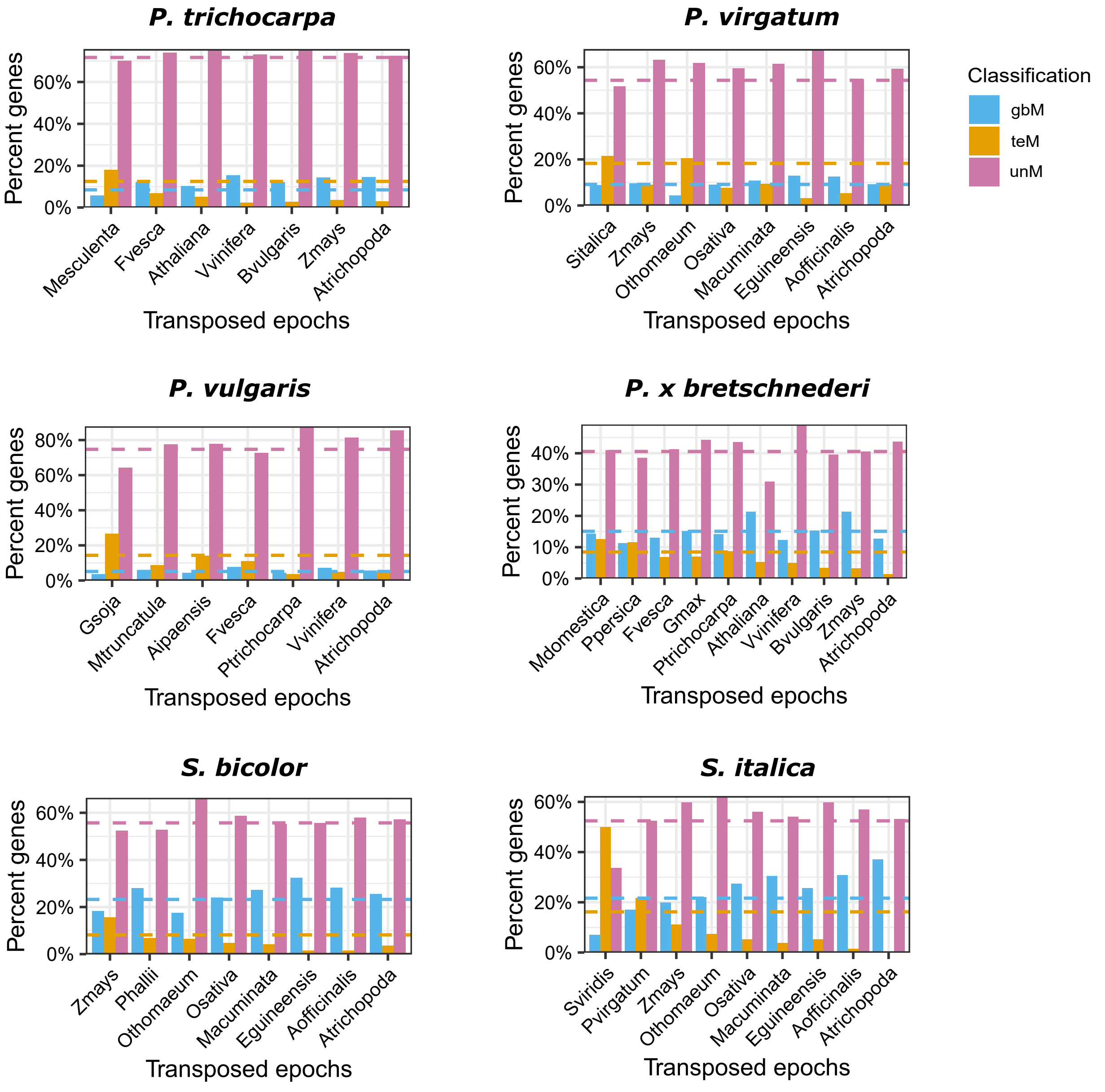

**Figure S8:** The percentage of gene copies in each genic methylation class for translocated genes that have duplicated during that 'epoch' since divergence from the species on the x-axis. For example, in *A. duranensis* translocated genes that have duplicated since *A. duranensis* diverged from *A. ipaensis* are shown on the x-axis under *A. ipaensis*. Those shown under *G. max*, duplicated in the period since the common ancestor of *A. duranensis* and *A. ipaensis* diverged from their common ancestor with *G. max*, but before the divergence of *A. duranensis* and *A. ipaensis.* Horizontal dotted lines indicate the percentage of each genic methylation class in all translocated duplicates. Bars above this line indicate enrichment, below this line depletion.

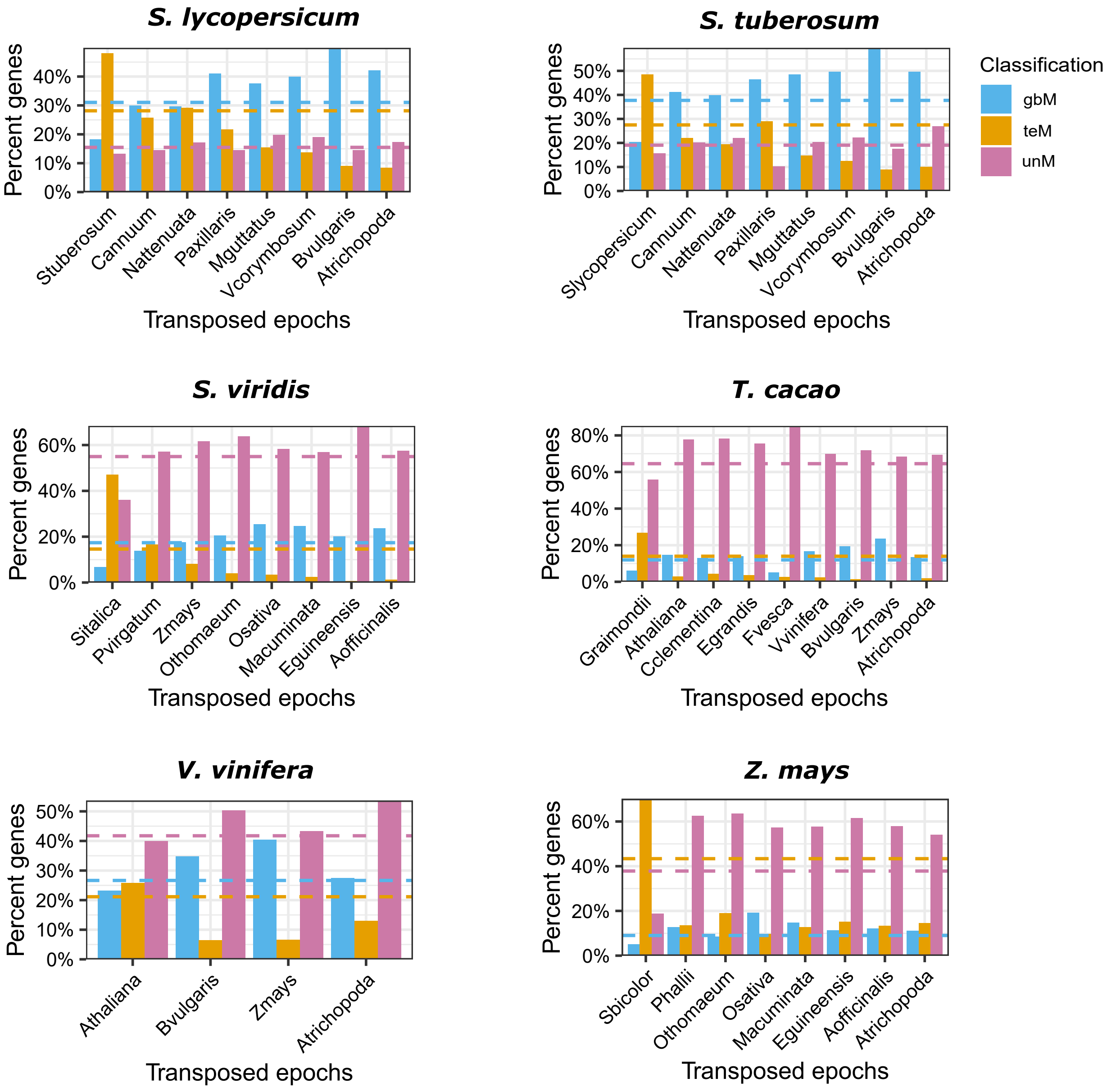

**Figure S9**: Distribution of core: multi-copy and core: single-copy (intermediate) paralogs based on synonymous substitutions (Ks). 'All pairs' represent Ks values of all duplicate gene pairs in the genome, 'Core-MC' represents duplicate pairs among core: multi-copy orthogroup, while 'SC-Int' represent duplicate pairs among the core: single-copy orthogroups.

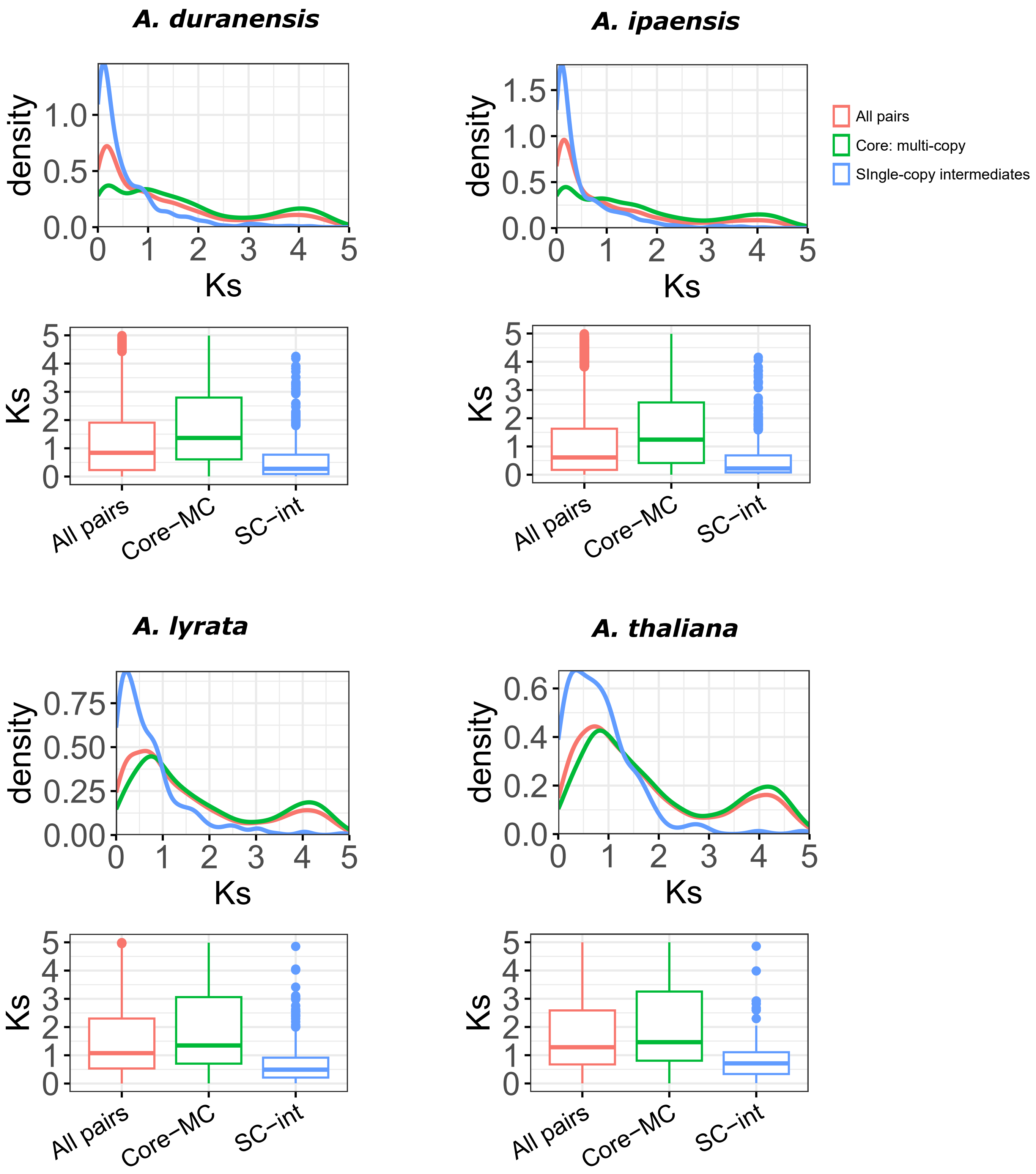

**Figure S9**: Distribution of core: multi-copy and core: single-copy (intermediate) paralogs based on synonymous substitutions (Ks). 'All pairs' represent Ks values of all duplicate gene pairs in the genome, 'Core-MC' represents duplicate pairs among core: multi-copy orthogroup, while 'SC-Int' represent duplicate pairs among the core: single-copy orthogroups.

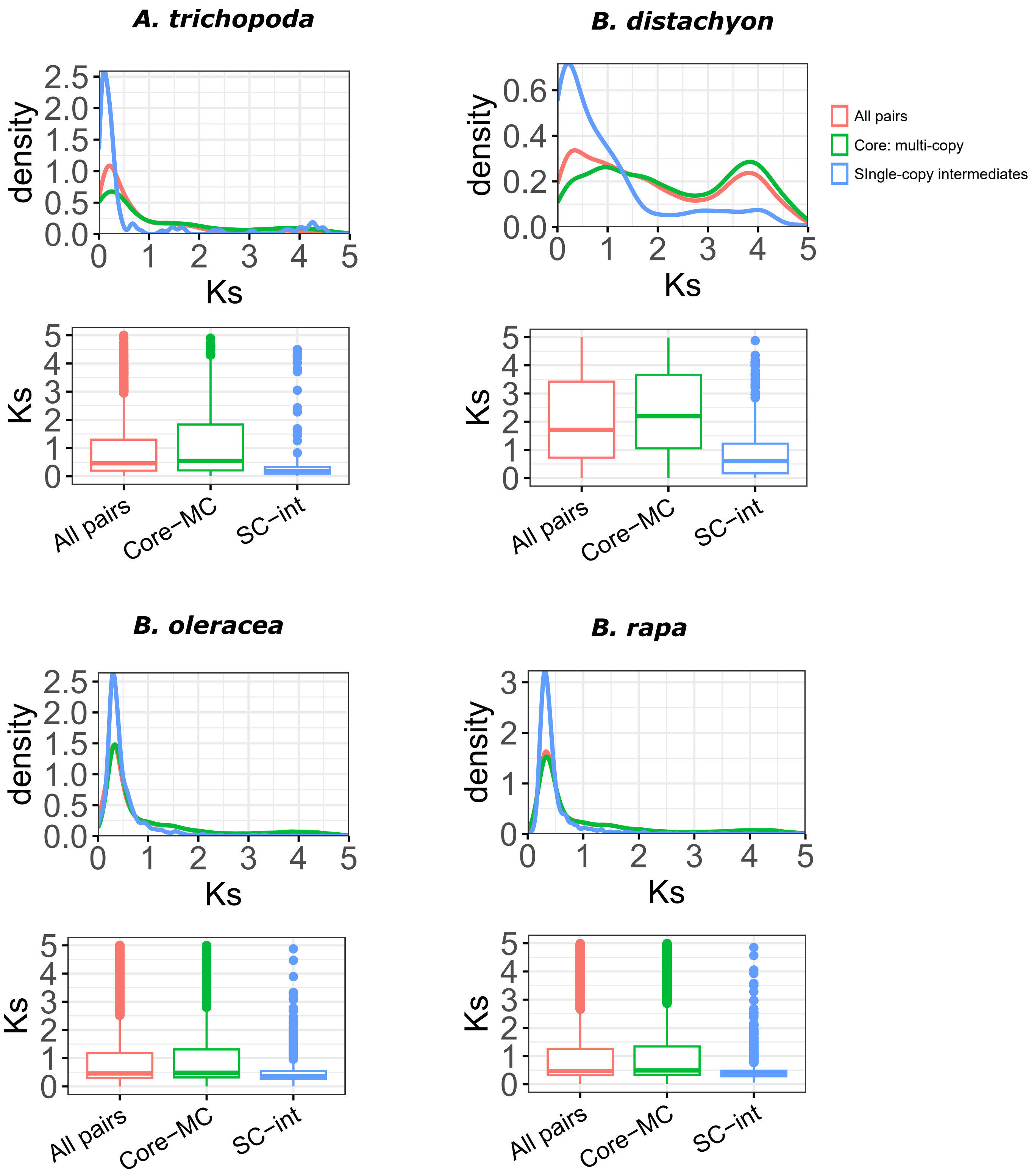

**Figure S9**: Distribution of core: multi-copy and core: single-copy (intermediate) paralogs based on synonymous substitutions (Ks). 'All pairs' represent Ks values of all duplicate gene pairs in the genome, 'Core-MC' represents duplicate pairs among core: multi-copy orthogroup, while 'SC-Int' represent duplicate pairs among the core: single-copy orthogroups.

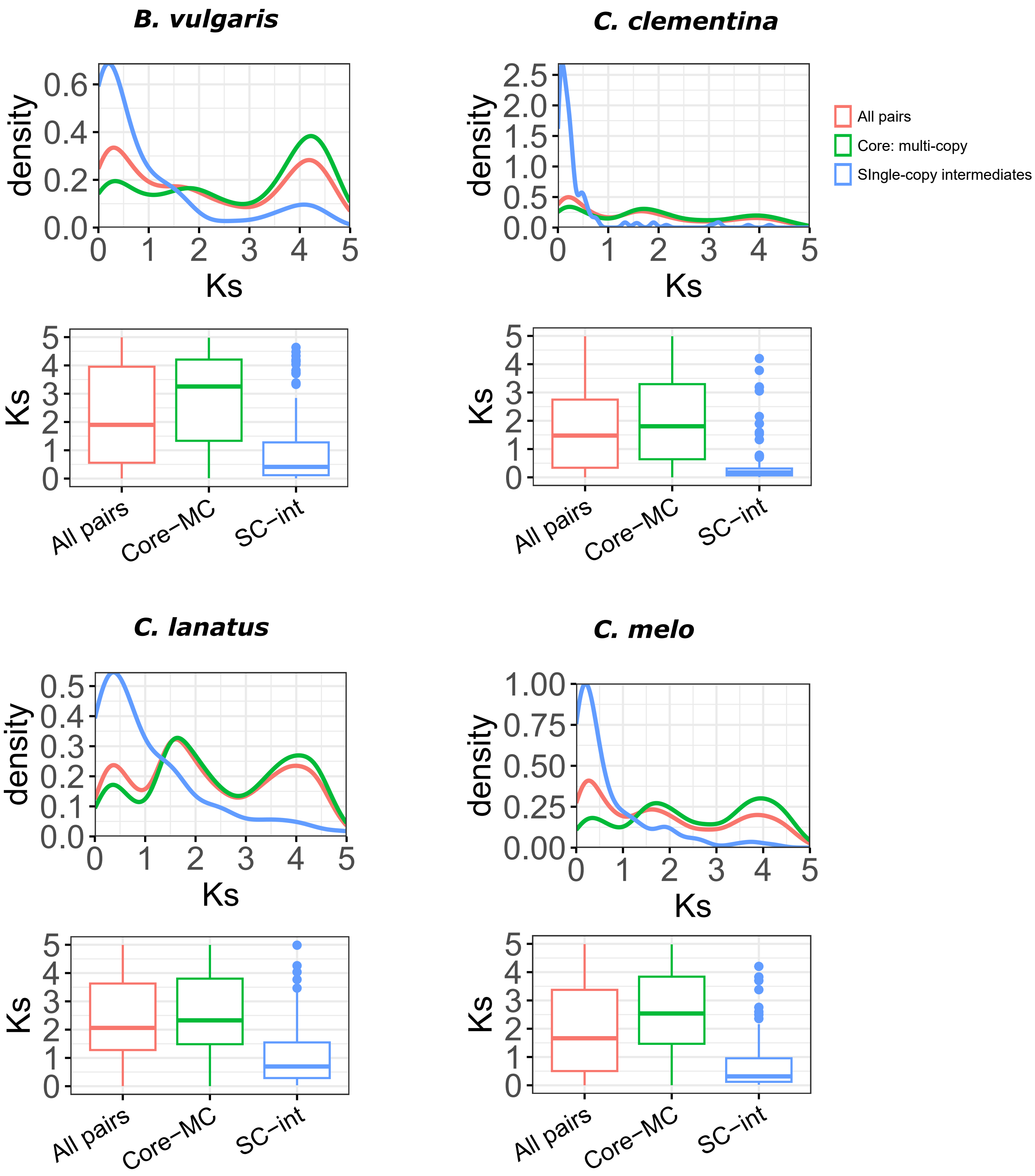

**Figure S9**: Distribution of core: multi-copy and core: single-copy (intermediate) paralogs based on synonymous substitutions (Ks). 'All pairs' represent Ks values of all duplicate gene pairs in the genome, 'Core-MC' represents duplicate pairs among core: multi-copy orthogroup, while 'SC-Int' represent duplicate pairs among the core: single-copy orthogroups.

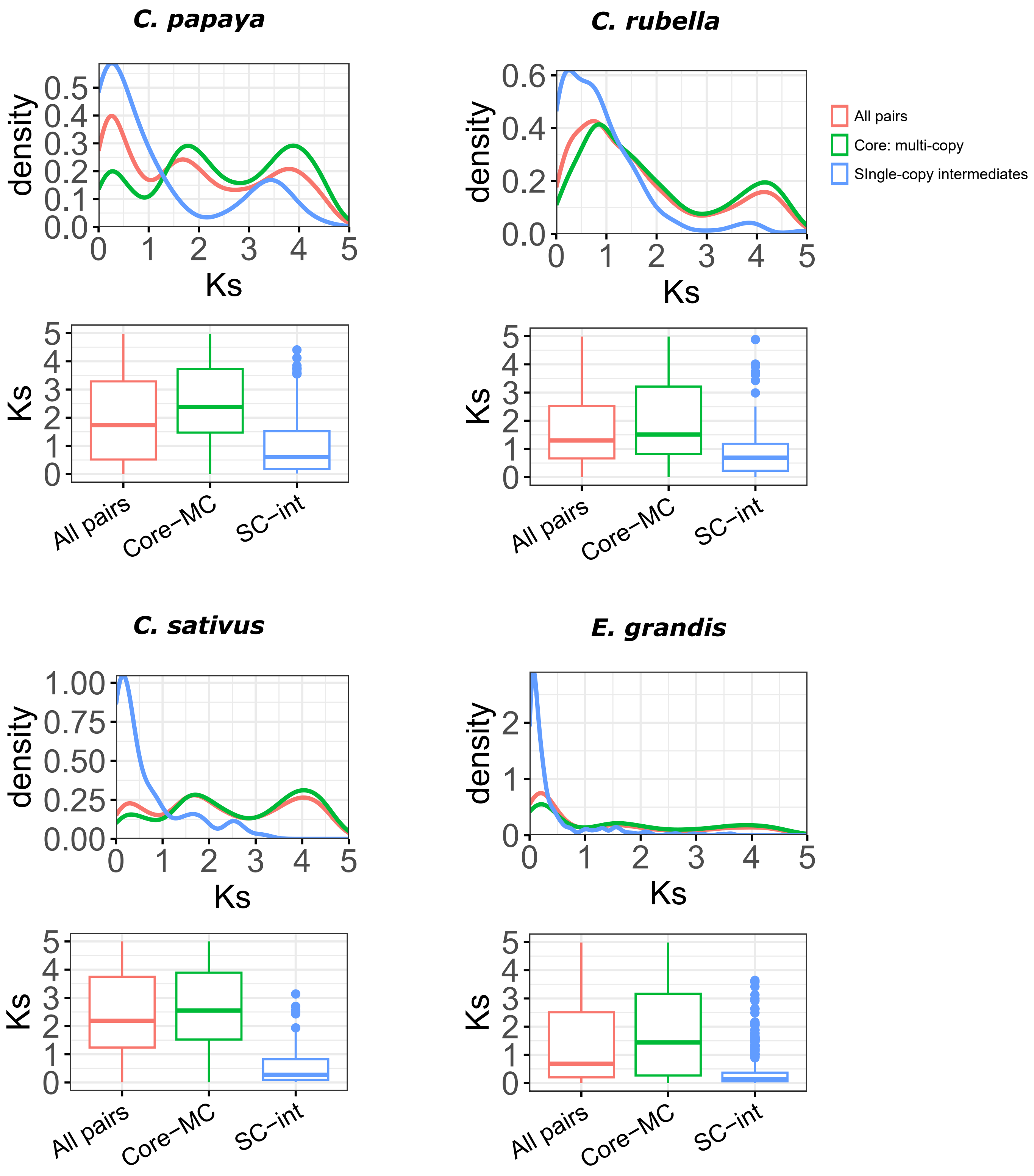

**Figure S9**: Distribution of core: multi-copy and core: single-copy (intermediate) paralogs based on synonymous substitutions (Ks). 'All pairs' represent Ks values of all duplicate gene pairs in the genome, 'Core-MC' represents duplicate pairs among core: multi-copy orthogroup,
while 'SC-Int' represent duplicate pairs among the core: single-copy orthogroups.

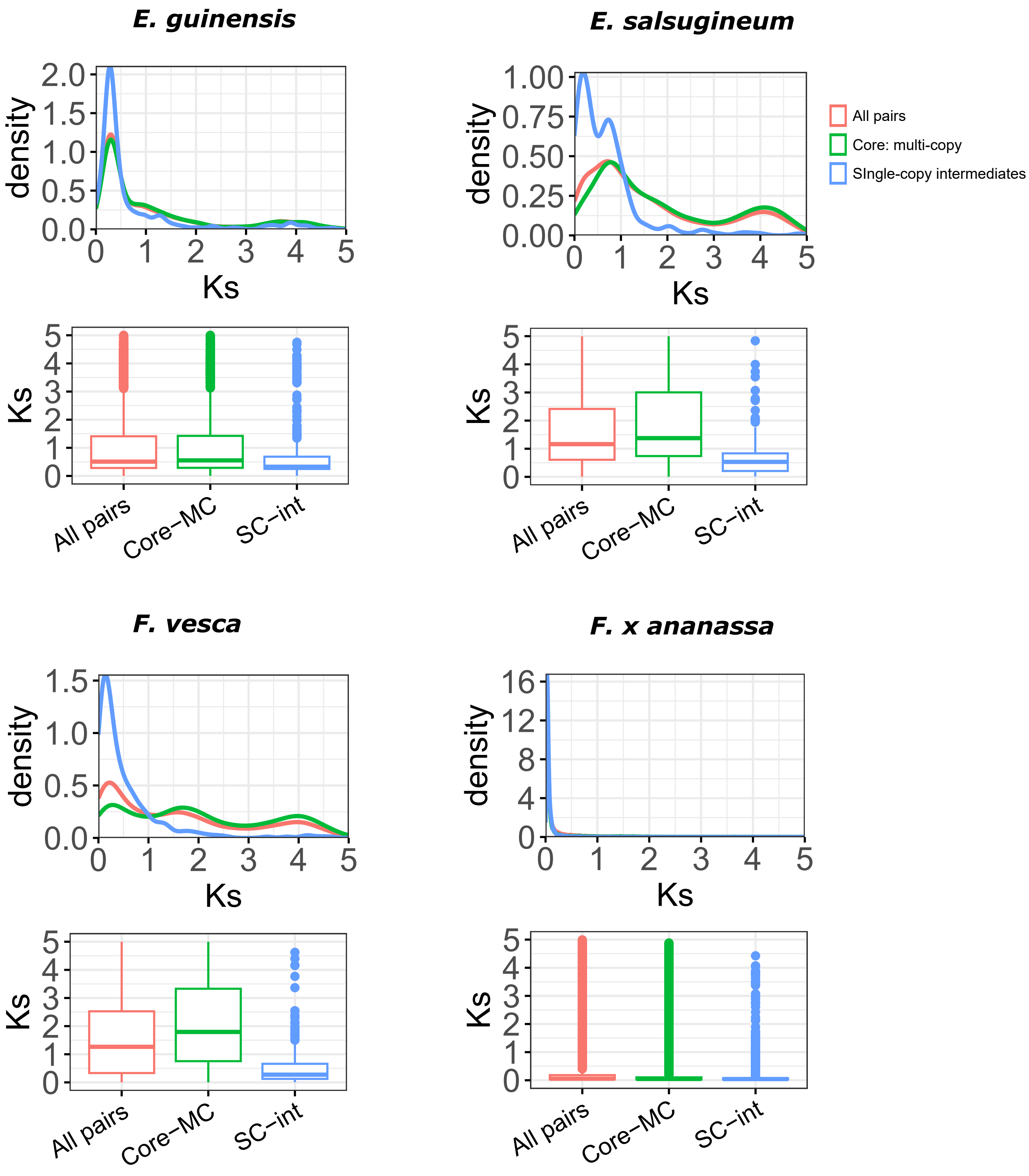

**Figure S9**: Distribution of core: multi-copy and core: single-copy (intermediate) paralogs based on synonymous substitutions (Ks). 'All pairs' represent Ks values of all duplicate gene pairs in the genome, 'Core-MC' represents duplicate pairs among core: multi-copy orthogroup, while 'SC-Int' represent duplicate pairs among the core: single-copy orthogroups.

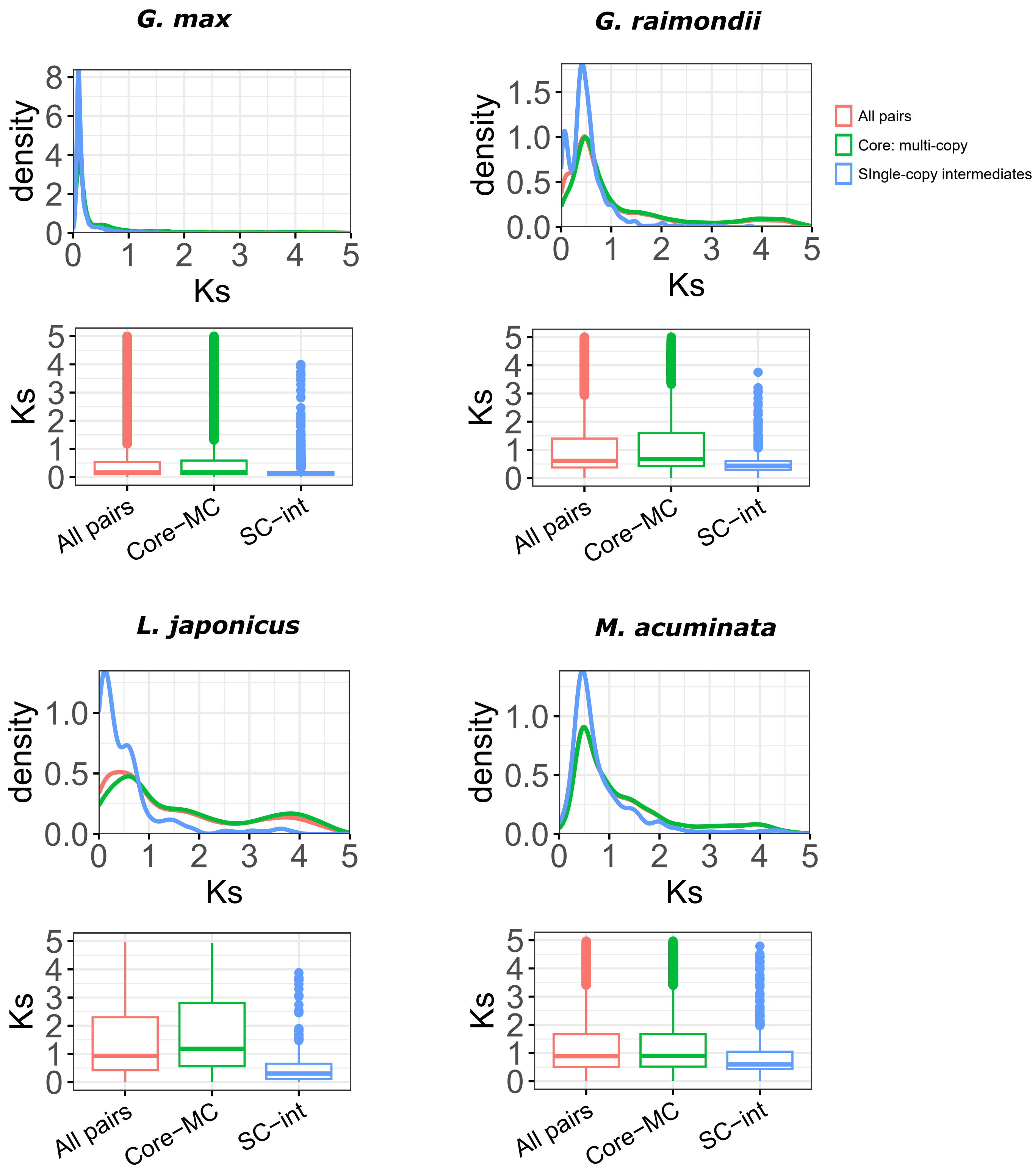

**Figure S9**: Distribution of core: multi-copy and core: single-copy (intermediate) paralogs based on synonymous substitutions (Ks). 'All pairs' represent Ks values of all duplicate gene pairs in the genome, 'Core-MC' represents duplicate pairs among core: multi-copy orthogroup, while 'SC-Int' represent duplicate pairs among the core: single-copy orthogroups.

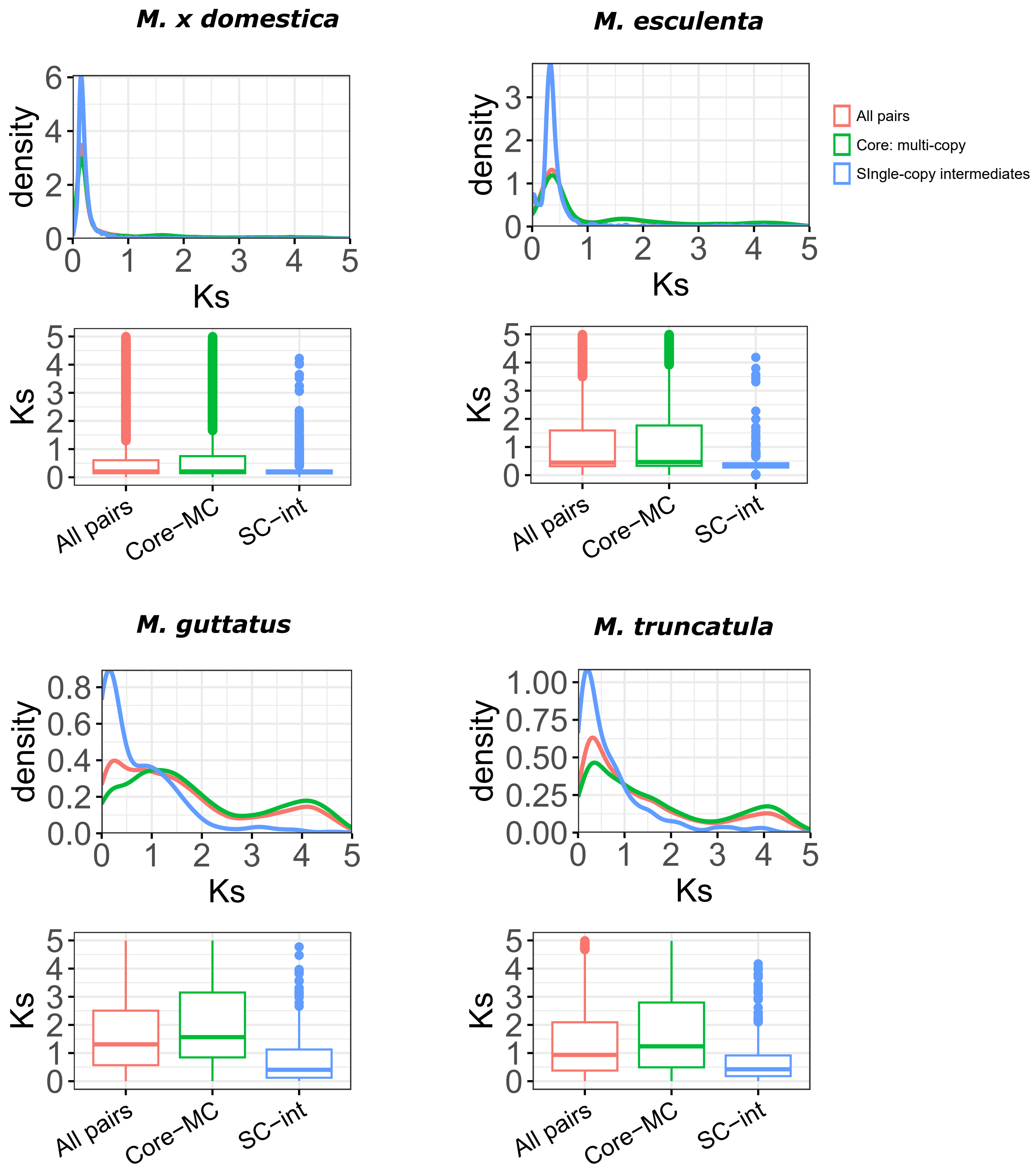

**Figure S9**: Distribution of core: multi-copy and core: single-copy (intermediate) paralogs based on synonymous substitutions (Ks). 'All pairs' represent Ks values of all duplicate gene pairs in the genome, 'Core-MC' represents duplicate pairs among core: multi-copy orthogroup, while 'SC-Int' represent duplicate pairs among the core: single-copy orthogroups.

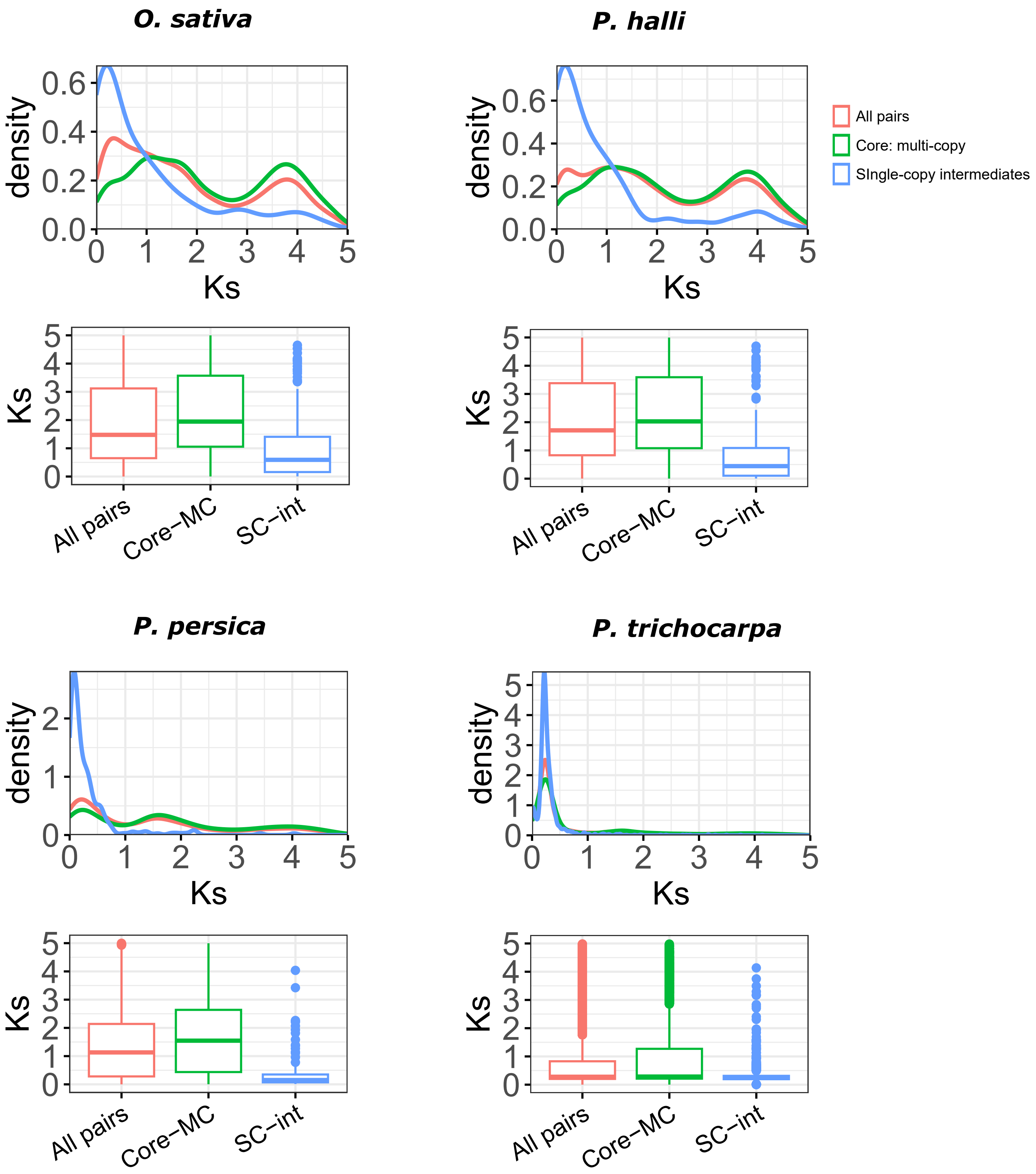

**Figure S9**: Distribution of core: multi-copy and core: single-copy (intermediate) paralogs based on synonymous substitutions (Ks). 'All pairs' represent Ks values of all duplicate gene pairs in the genome, 'Core-MC' represents duplicate pairs among core: multi-copy orthogroup, while 'SC-Int' represent duplicate pairs among the core: single-copy orthogroups.

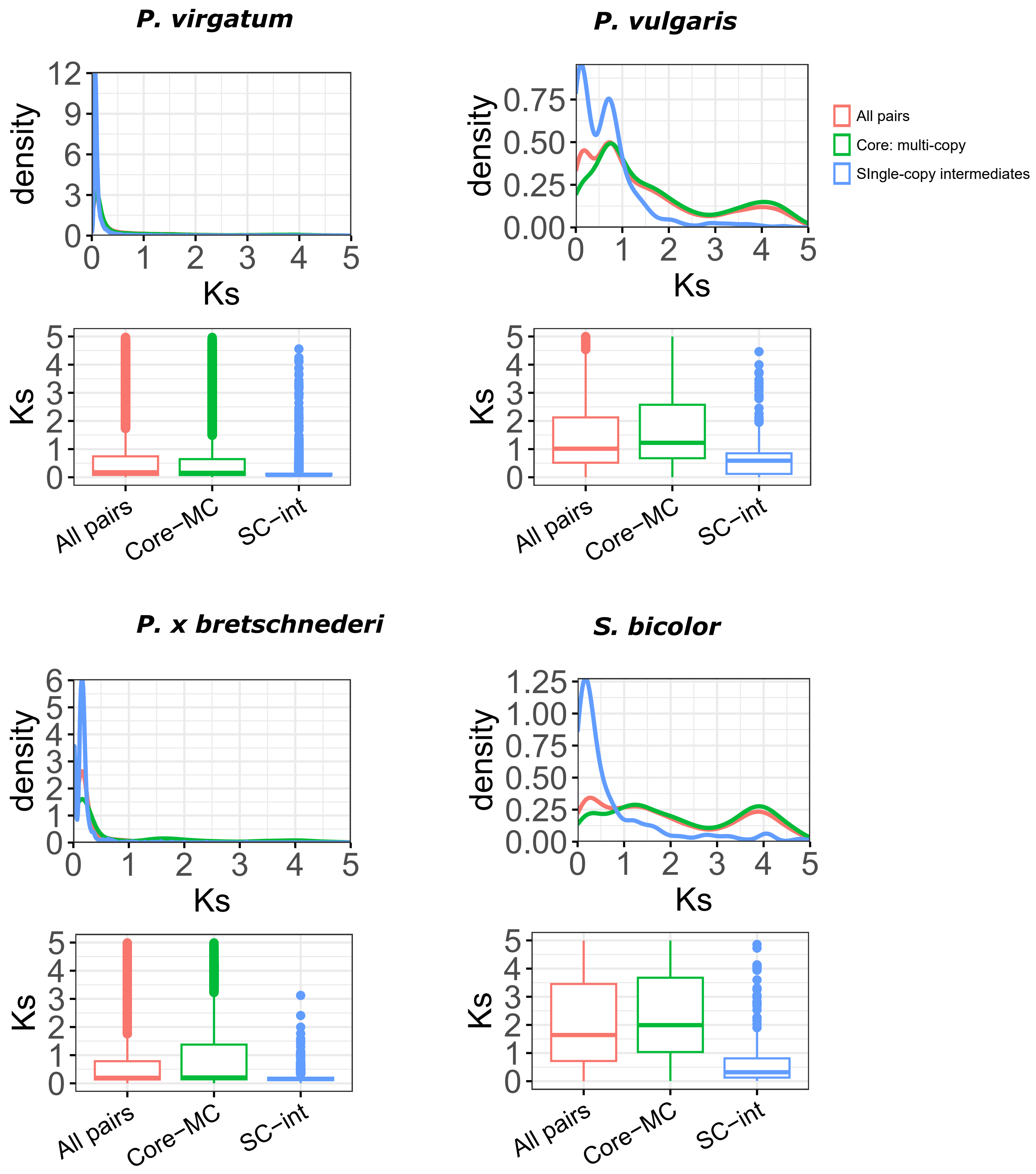

**Figure S9**: Distribution of core: multi-copy and core: single-copy (intermediate) paralogs based on synonymous substitutions (Ks). 'All pairs' represent Ks values of all duplicate gene pairs in the genome, 'Core-MC' represents duplicate pairs among core: multi-copy orthogroup, while 'SC-Int' represent duplicate pairs among the core: single-copy orthogroups.

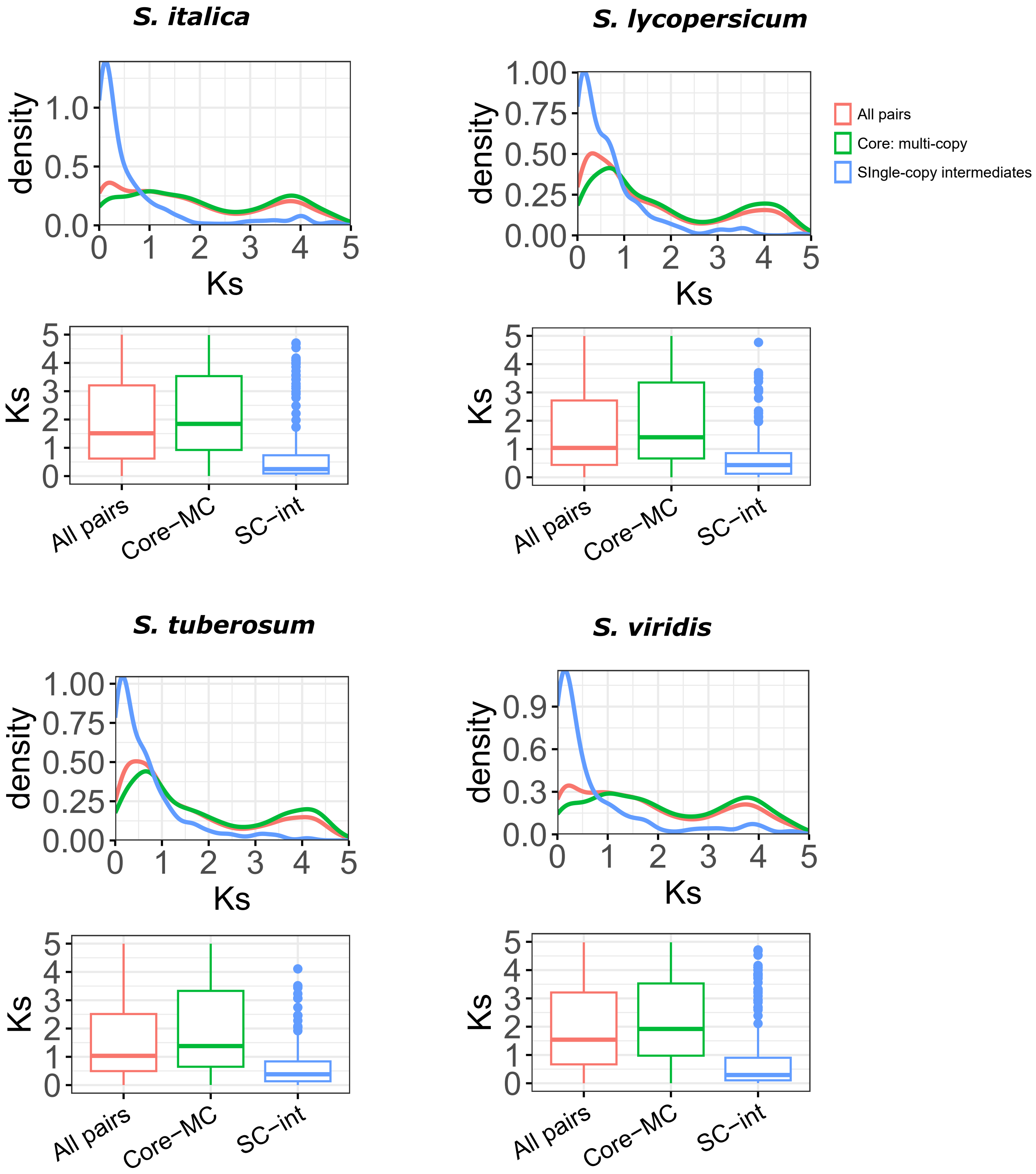

**Figure S9**: Distribution of core: multi-copy and core: single-copy (intermediate) paralogs based on synonymous substitutions (Ks). 'All pairs' represent Ks values of all duplicate gene pairs in the genome, 'Core-MC' represents duplicate pairs among core: multi-copy orthogroup, while 'SC-Int' represent duplicate pairs among the core: single-copy orthogroups.

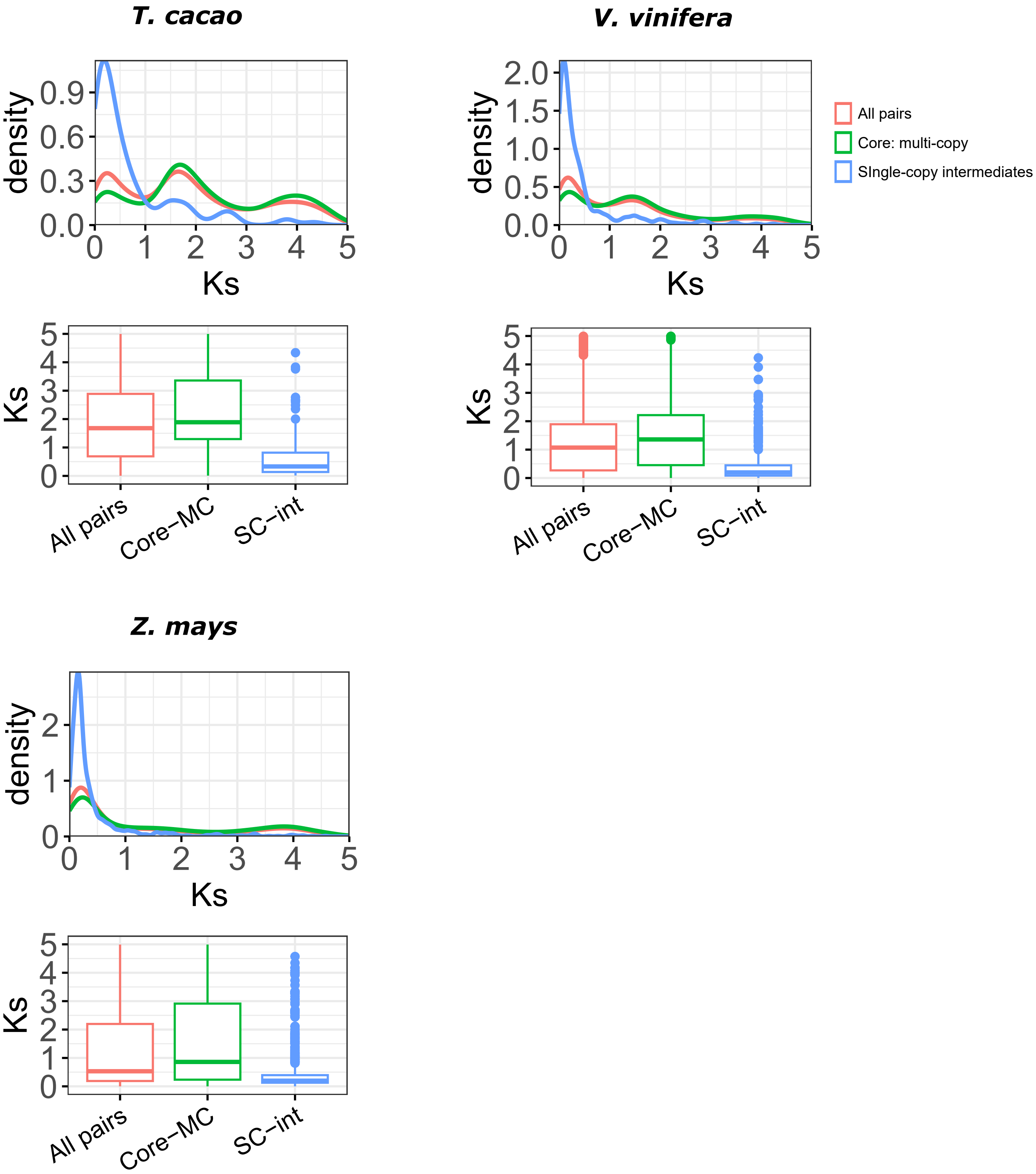

**Figure S10**: Distribution of genic methylation classified genes based on the ratio of nonsynonymous substitution (Ka), with synonymous substitutions (Ks) across different types of gene duplicate pairs.
Whole-genome duplicates - WGD, Single-gene duplicates - SGD (combined data from tandem, proximal, translocated, and dispersed duplicates).

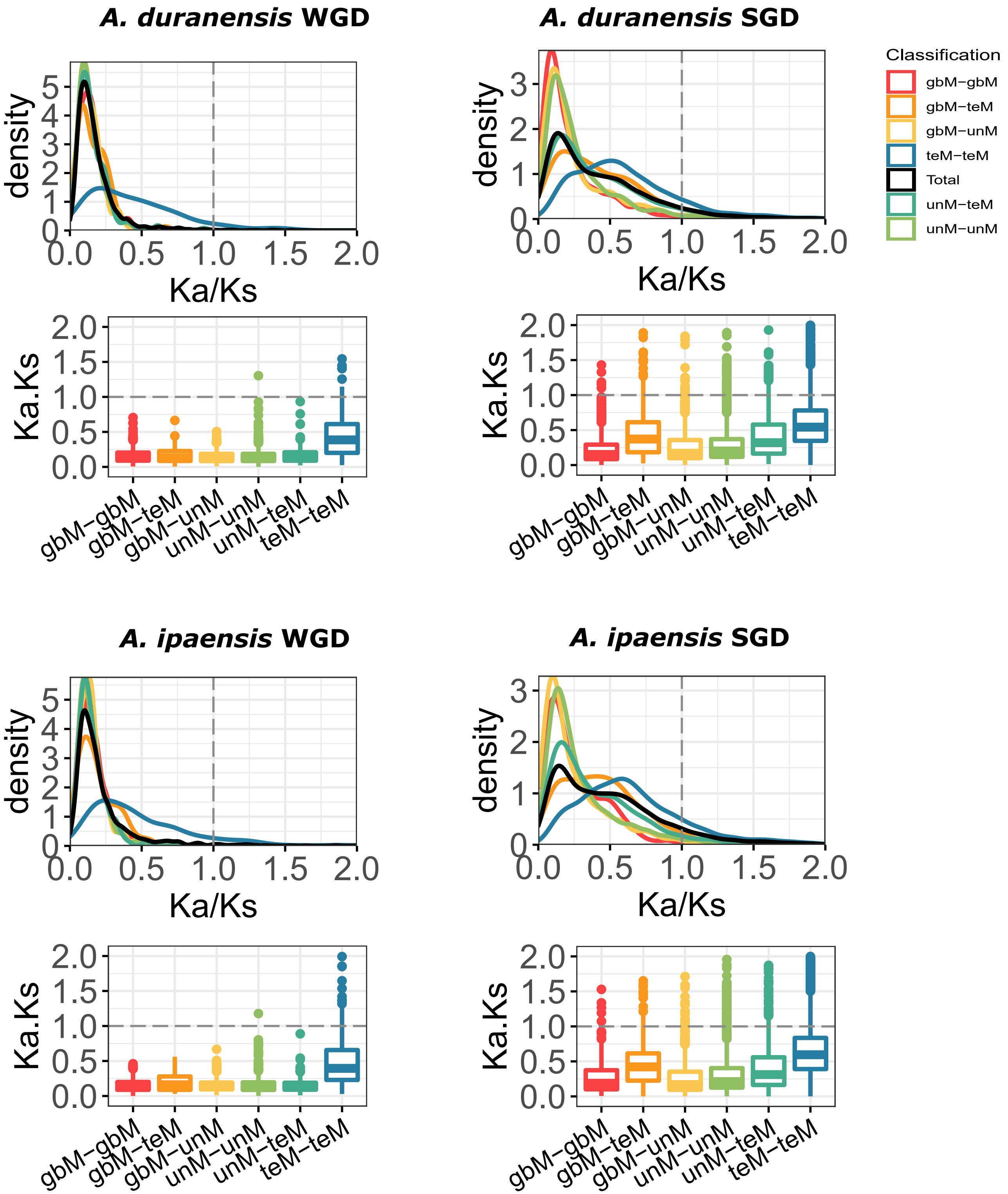

**Figure S10**: Distribution of genic methylation classified genes based on the ratio of nonsynonymous substitution (Ka), with synonymous substitutions (Ks) across different types of gene duplicate pairs.
Whole-genome duplicates - WGD, Single-gene duplicates - SGD (combined data from tandem, proximal, translocated, and dispersed duplicates).

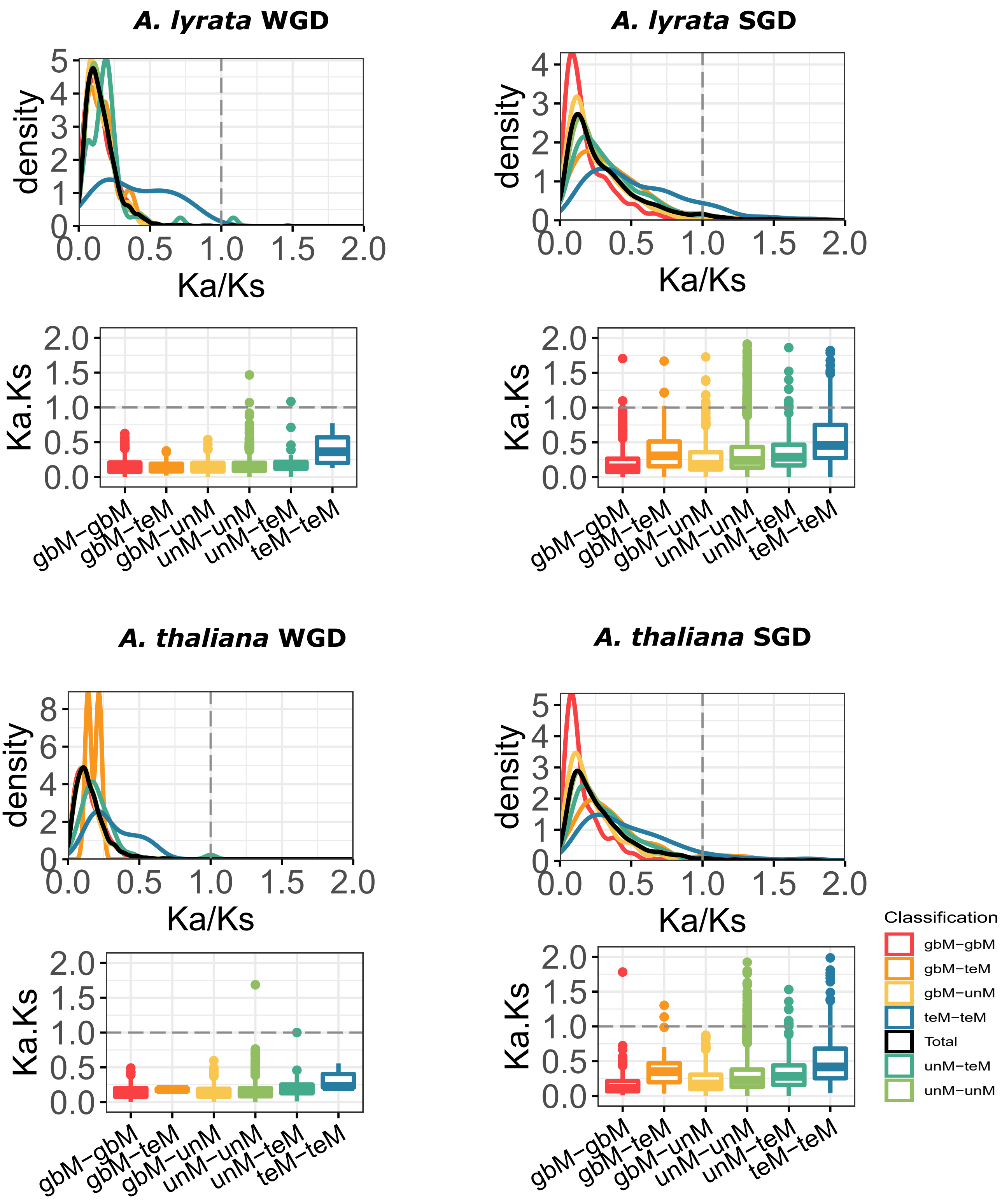

**Figure S10**: Distribution of genic methylation classified genes based on the ratio of nonsynonymous substitution (Ka), with synonymous substitutions (Ks) across different types of gene duplicate pairs.
Whole-genome duplicates - WGD, Single-gene duplicates - SGD (combined data from tandem, proximal, translocated, and dispersed duplicates).

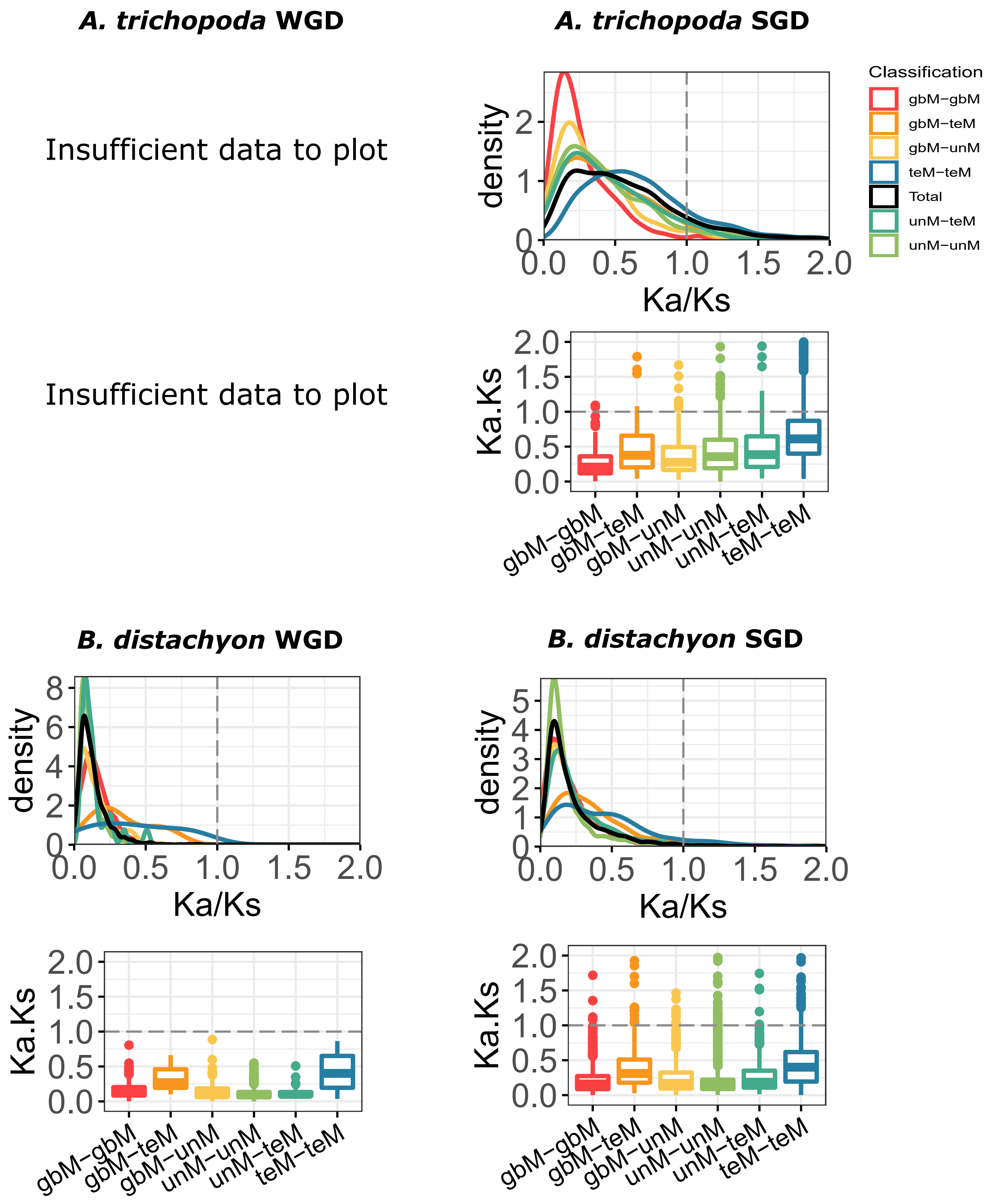

**Figure S10**: Distribution of genic methylation classified genes based on the ratio of nonsynonymous substitution (Ka), with synonymous substitutions (Ks) across different types of gene duplicate pairs.
Whole-genome duplicates - WGD, Single-gene duplicates - SGD (combined data from tandem, proximal, translocated, and dispersed duplicates).

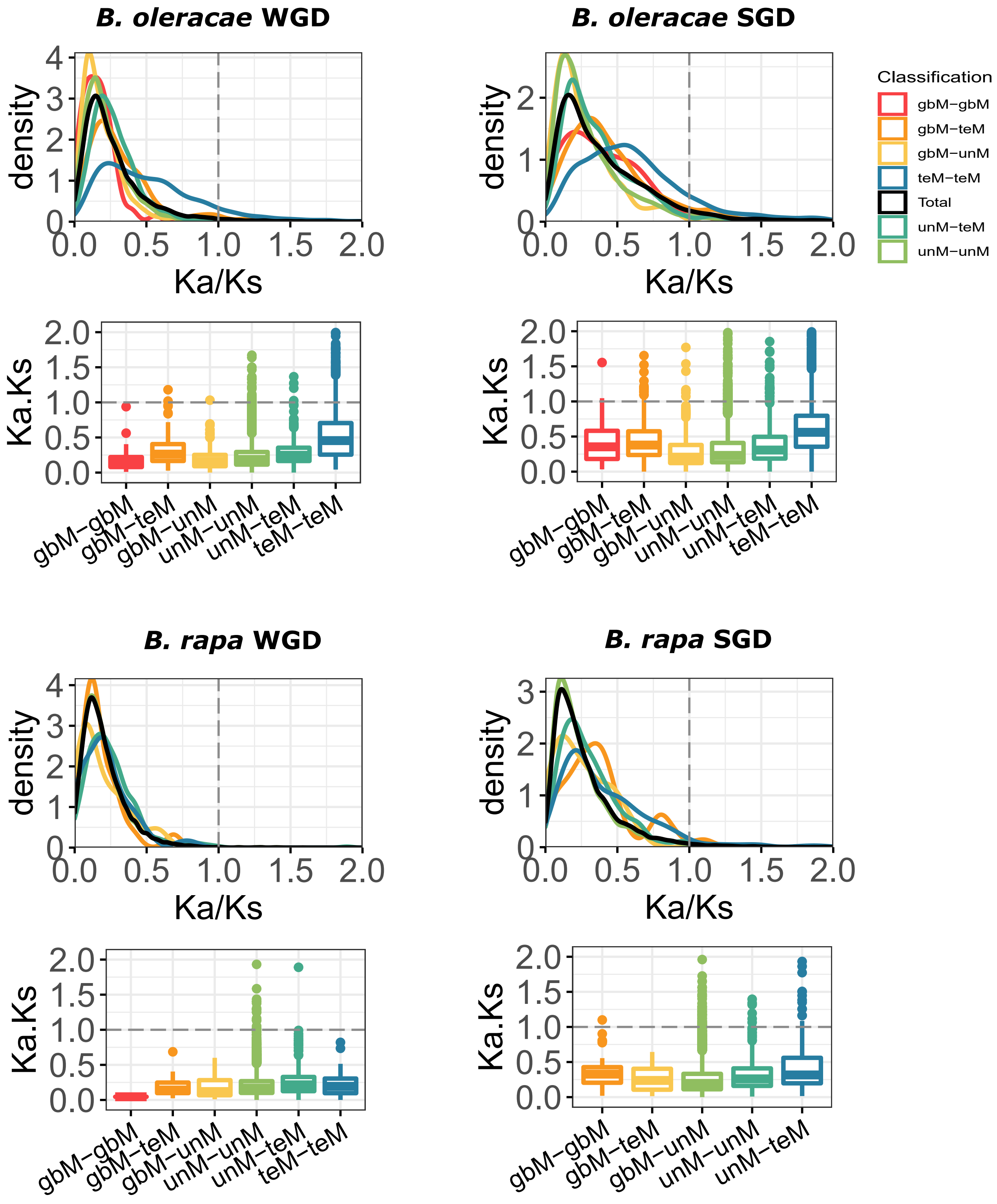

**Figure S10**: Distribution of genic methylation classified genes based on the ratio of nonsynonymous substitution (Ka), with synonymous substitutions (Ks) across different types of gene duplicate pairs.
Whole-genome duplicates - WGD, Single-gene duplicates - SGD (combined data from tandem, proximal, translocated, and dispersed duplicates).

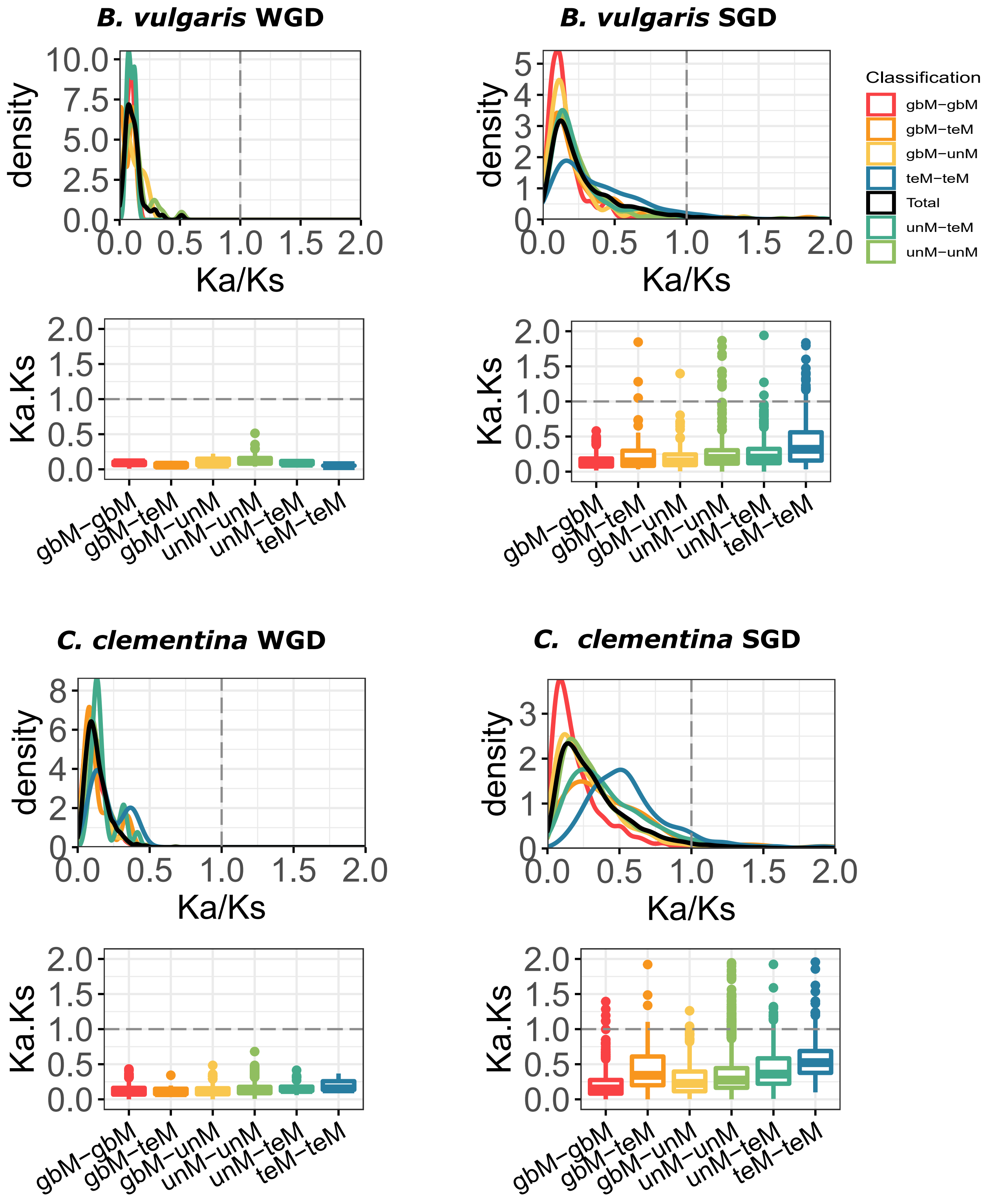

**Figure S10**: Distribution of genic methylation classified genes based on the ratio of nonsynonymous substitution (Ka), with synonymous substitutions (Ks) across different types of gene duplicate pairs.
Whole-genome duplicates - WGD, Single-gene duplicates - SGD (combined data from tandem, proximal, translocated, and dispersed duplicates).

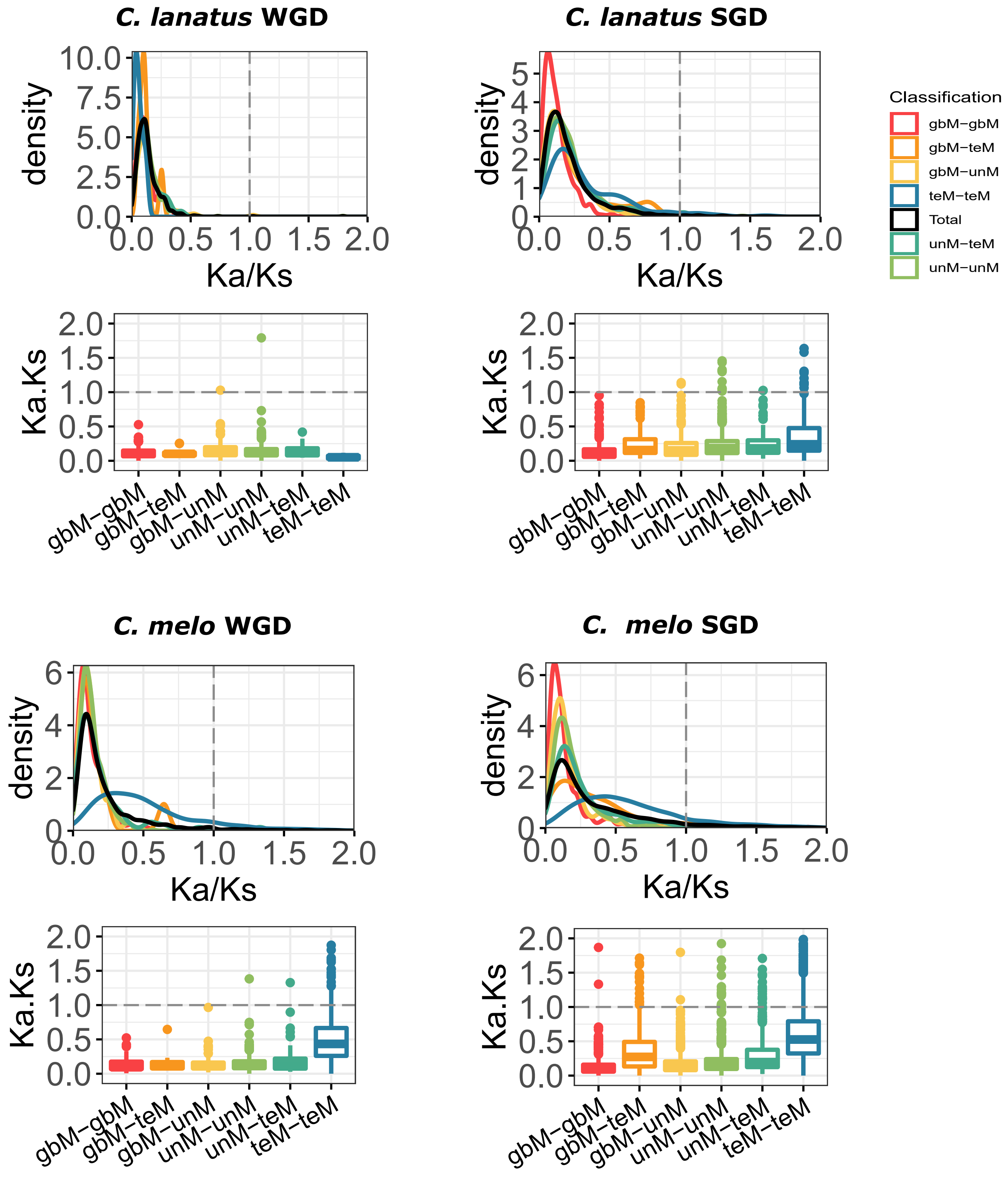

**Figure S10**: Distribution of genic methylation classified genes based on the ratio of nonsynonymous substitution (Ka), with synonymous substitutions (Ks) across different types of gene duplicate pairs.
Whole-genome duplicates - WGD, Single-gene duplicates - SGD (combined data from tandem, proximal, translocated, and dispersed duplicates).

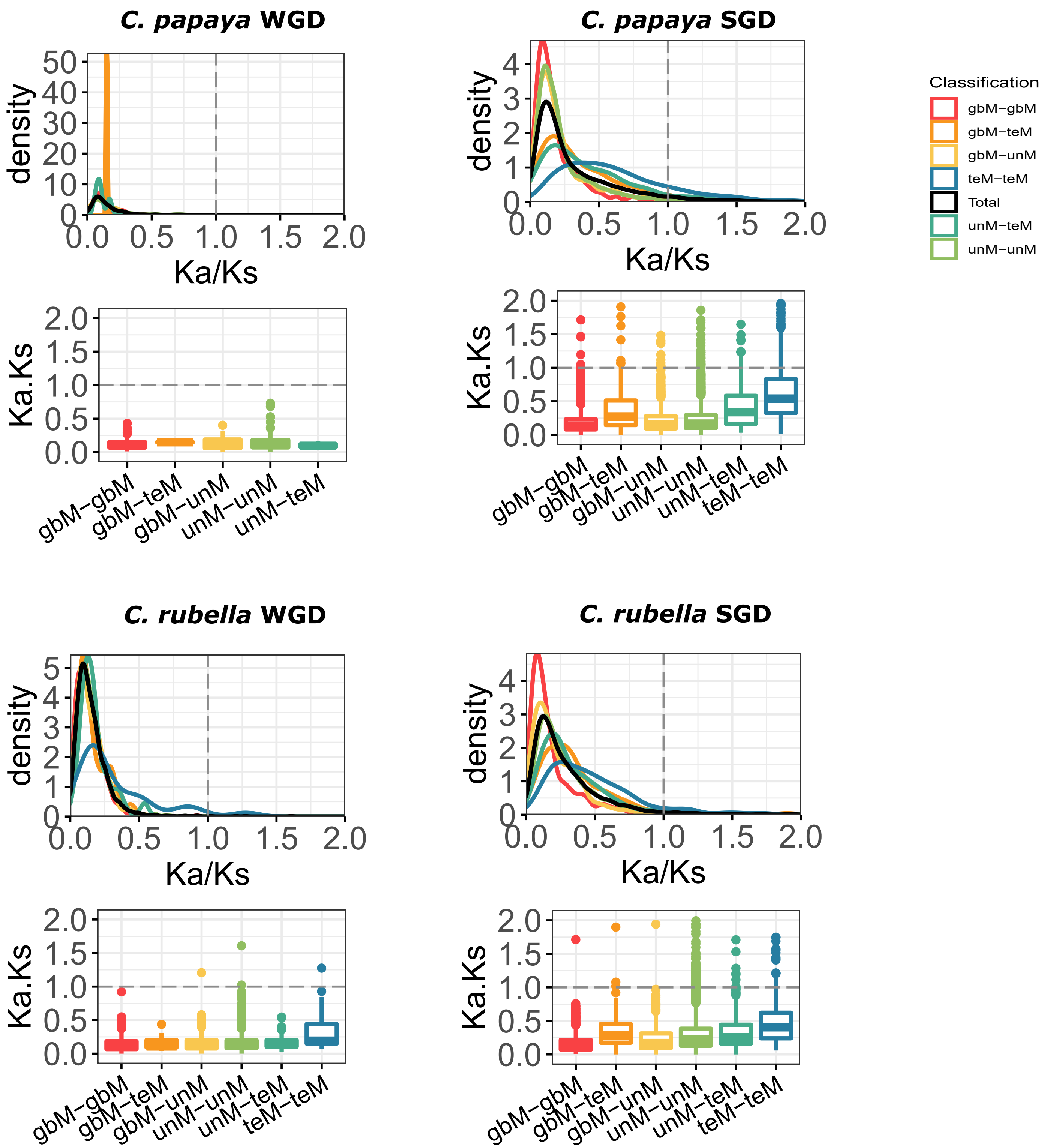

**Figure S10**: Distribution of genic methylation classified genes based on the ratio of nonsynonymous substitution (Ka), with synonymous substitutions (Ks) across different types of gene duplicate pairs.
Whole-genome duplicates - WGD, Single-gene duplicates - SGD (combined data from tandem, proximal, translocated, and dispersed duplicates).

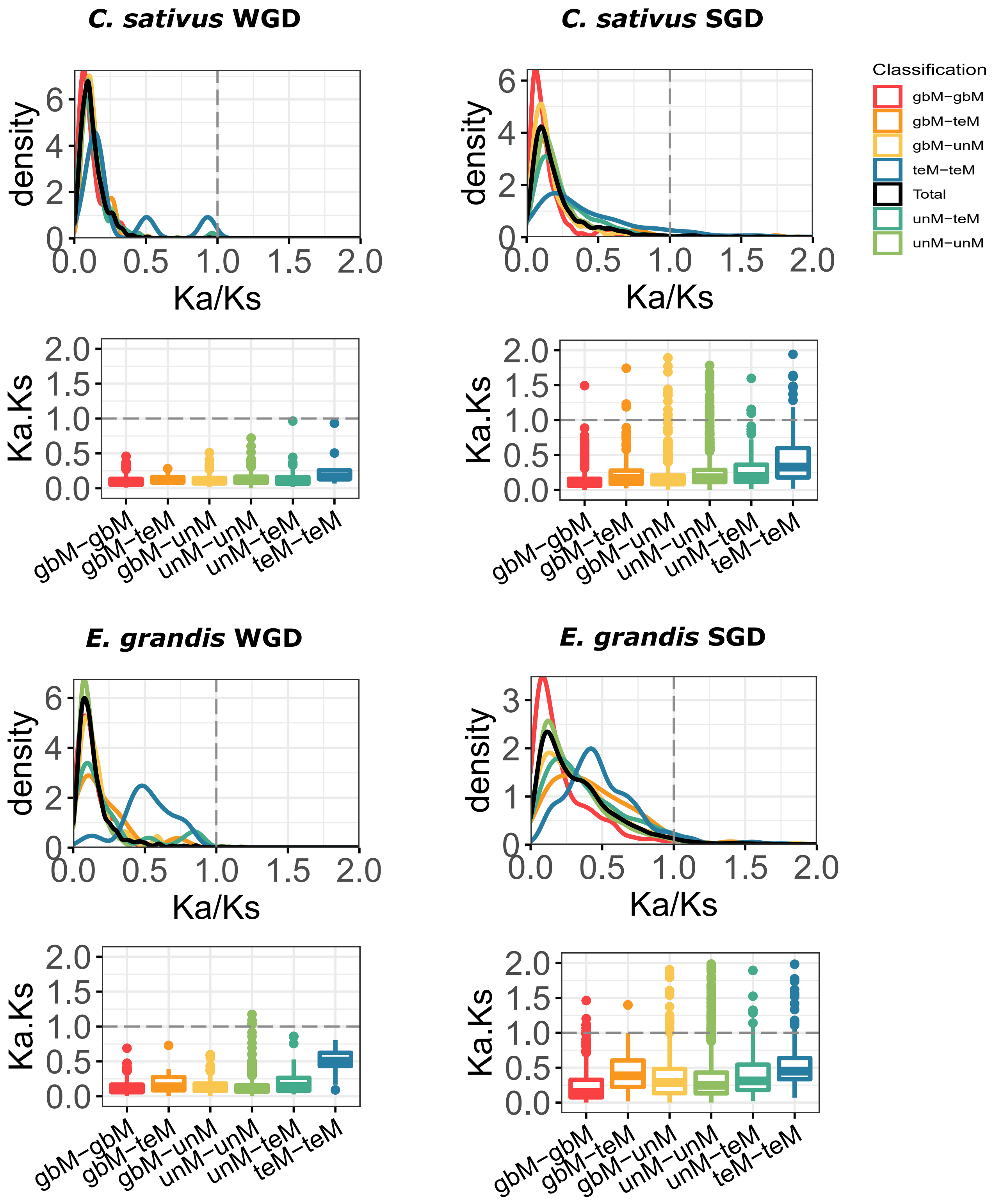

**Figure S10**: Distribution of genic methylation classified genes based on the ratio of nonsynonymous substitution (Ka), with synonymous substitutions (Ks) across different types of gene duplicate pairs.
Whole-genome duplicates - WGD, Single-gene duplicates - SGD (combined data from tandem, proximal, translocated, and dispersed duplicates).

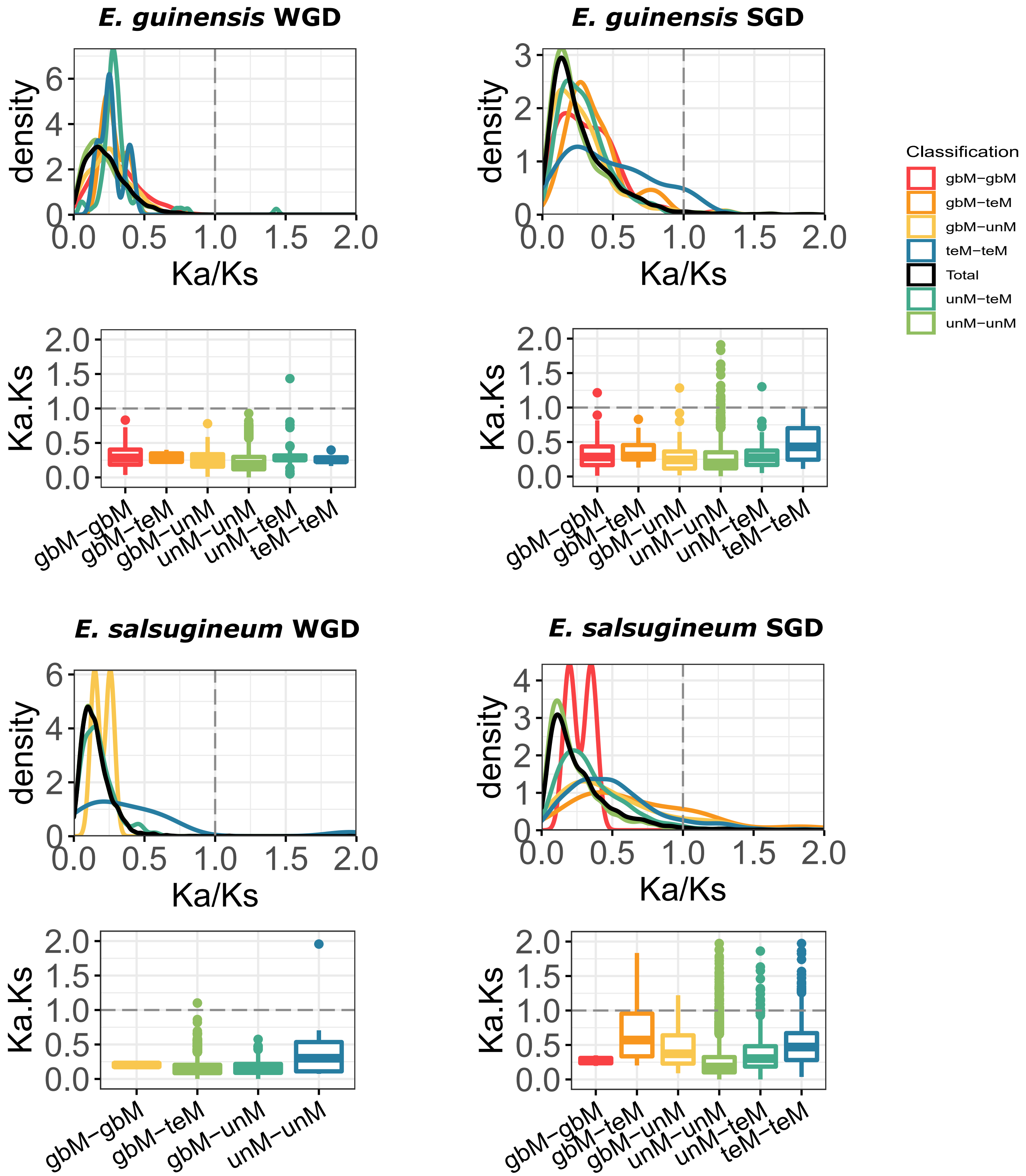

**Figure S10**: Distribution of genic methylation classified genes based on the ratio of nonsynonymous substitution (Ka), with synonymous substitutions (Ks) across different types of gene duplicate pairs.
Whole-genome duplicates - WGD, Single-gene duplicates - SGD (combined data from tandem, proximal, translocated, and dispersed duplicates).

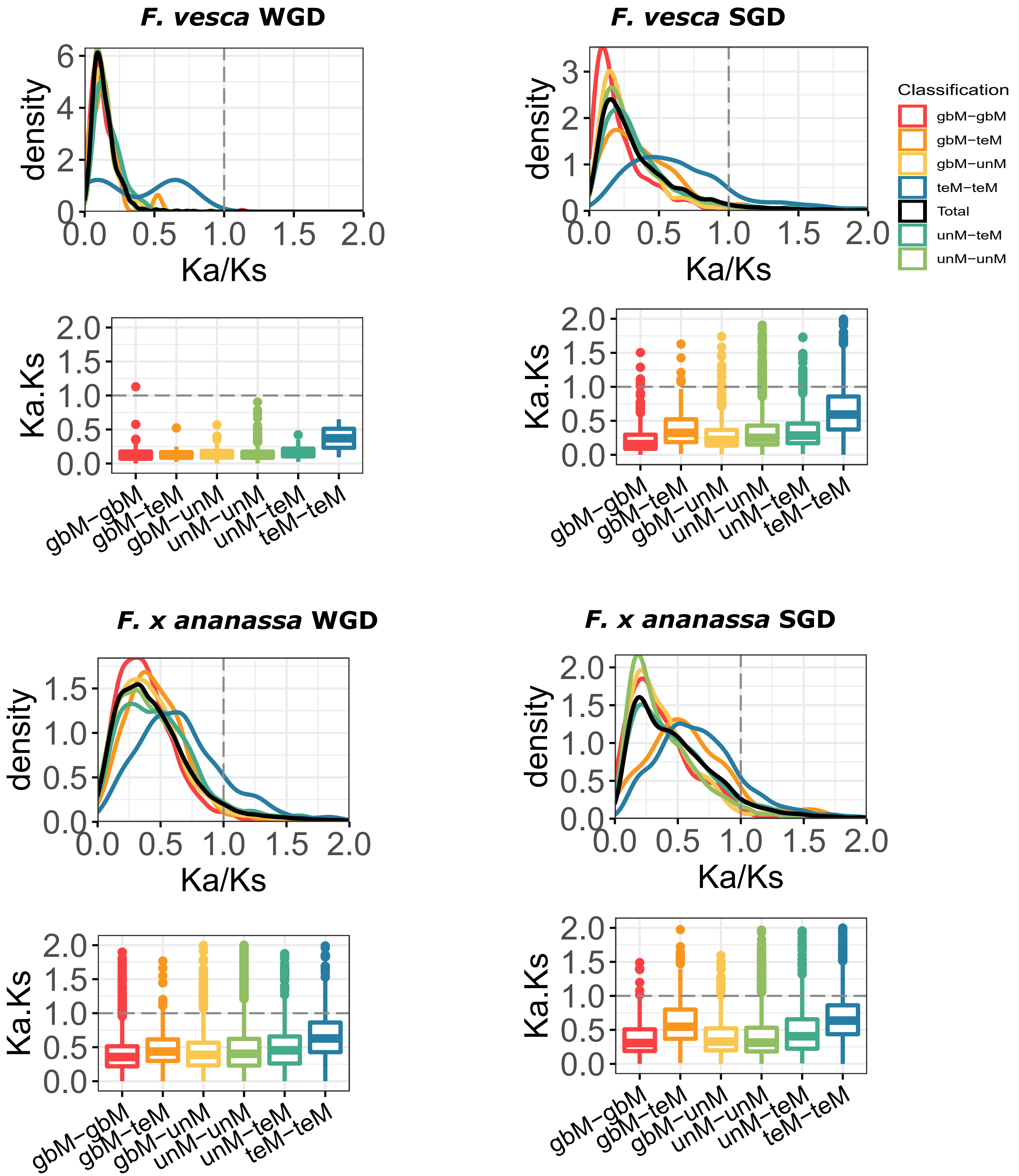

**Figure S10**: Distribution of genic methylation classified genes based on the ratio of nonsynonymous substitution (Ka), with synonymous substitutions (Ks) across different types of gene duplicate pairs. Whole-genome duplicates - WGD, Single-gene duplicates - SGD (combined data from tandem, proximal, translocated, and dispersed duplicates).

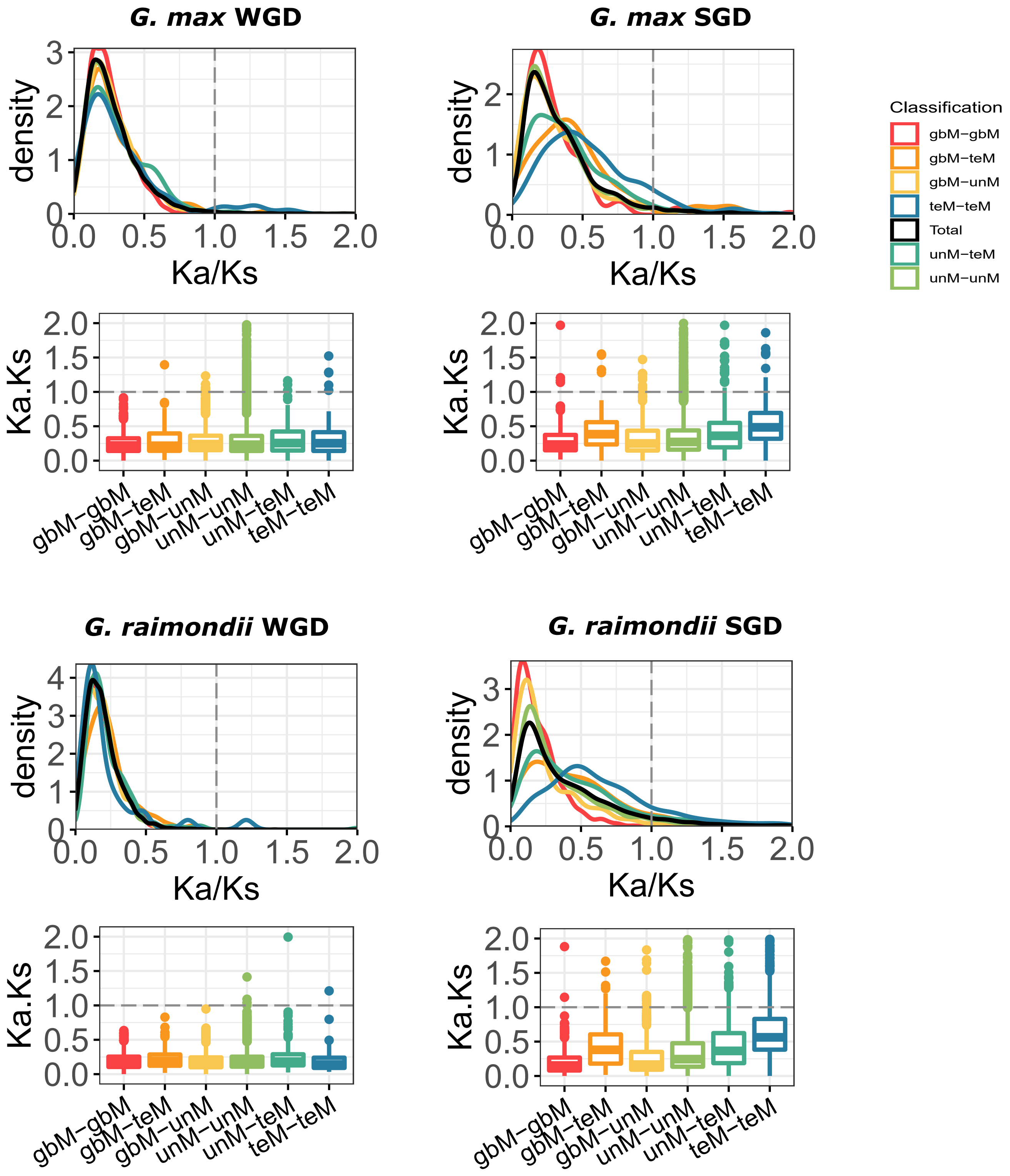

**Figure S10**: Distribution of genic methylation classified genes based on the ratio of nonsynonymous substitution (Ka), with synonymous substitutions (Ks) across different types of gene duplicate pairs.
Whole-genome duplicates - WGD, Single-gene duplicates - SGD (combined data from tandem, proximal, translocated, and dispersed duplicates).

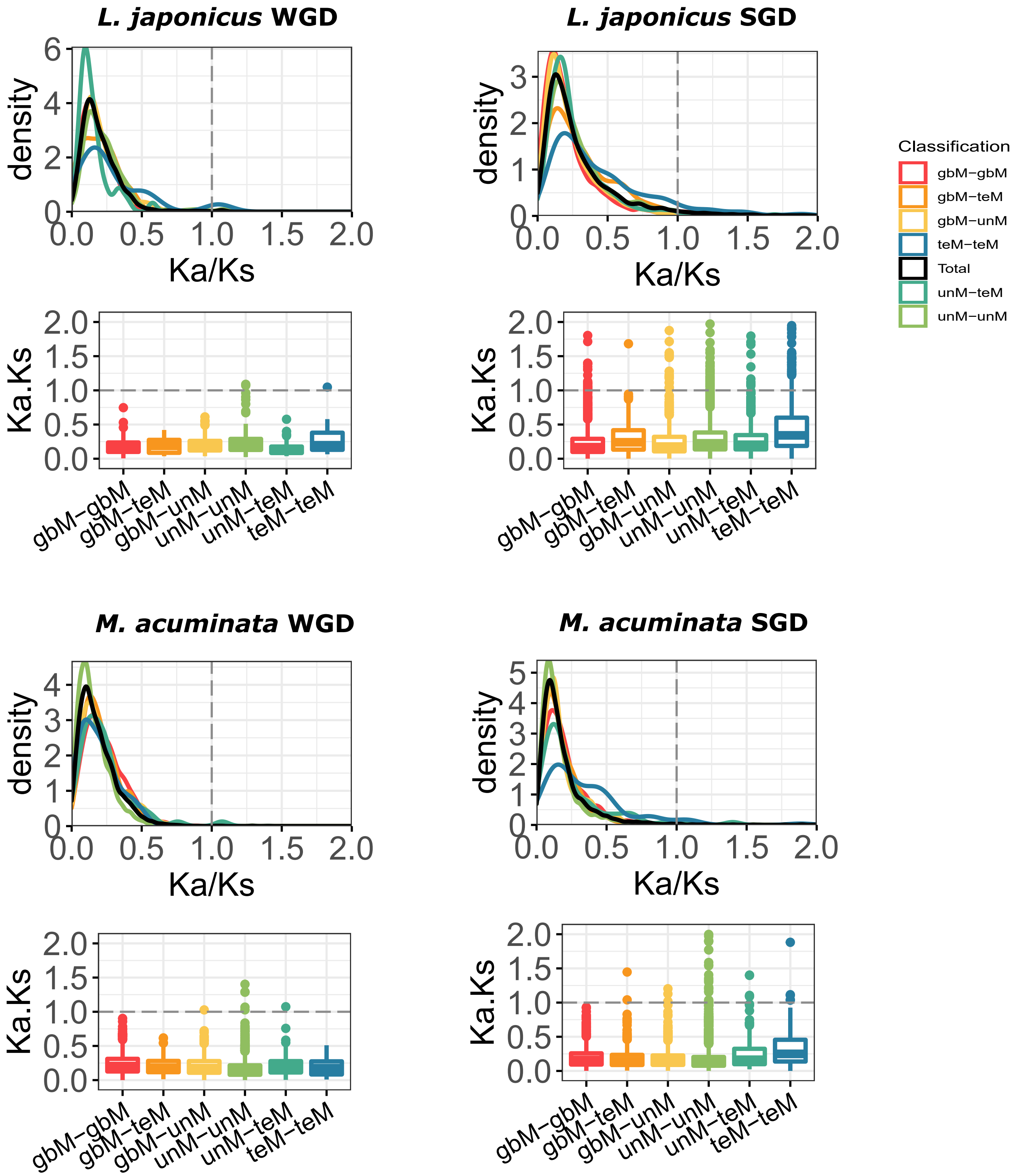

**Figure S10**: Distribution of genic methylation classified genes based on the ratio of nonsynonymous substitution (Ka), with synonymous substitutions (Ks) across different types of gene duplicate pairs.
Whole-genome duplicates - WGD, Single-gene duplicates - SGD (combined data from tandem, proximal, translocated, and dispersed duplicates).

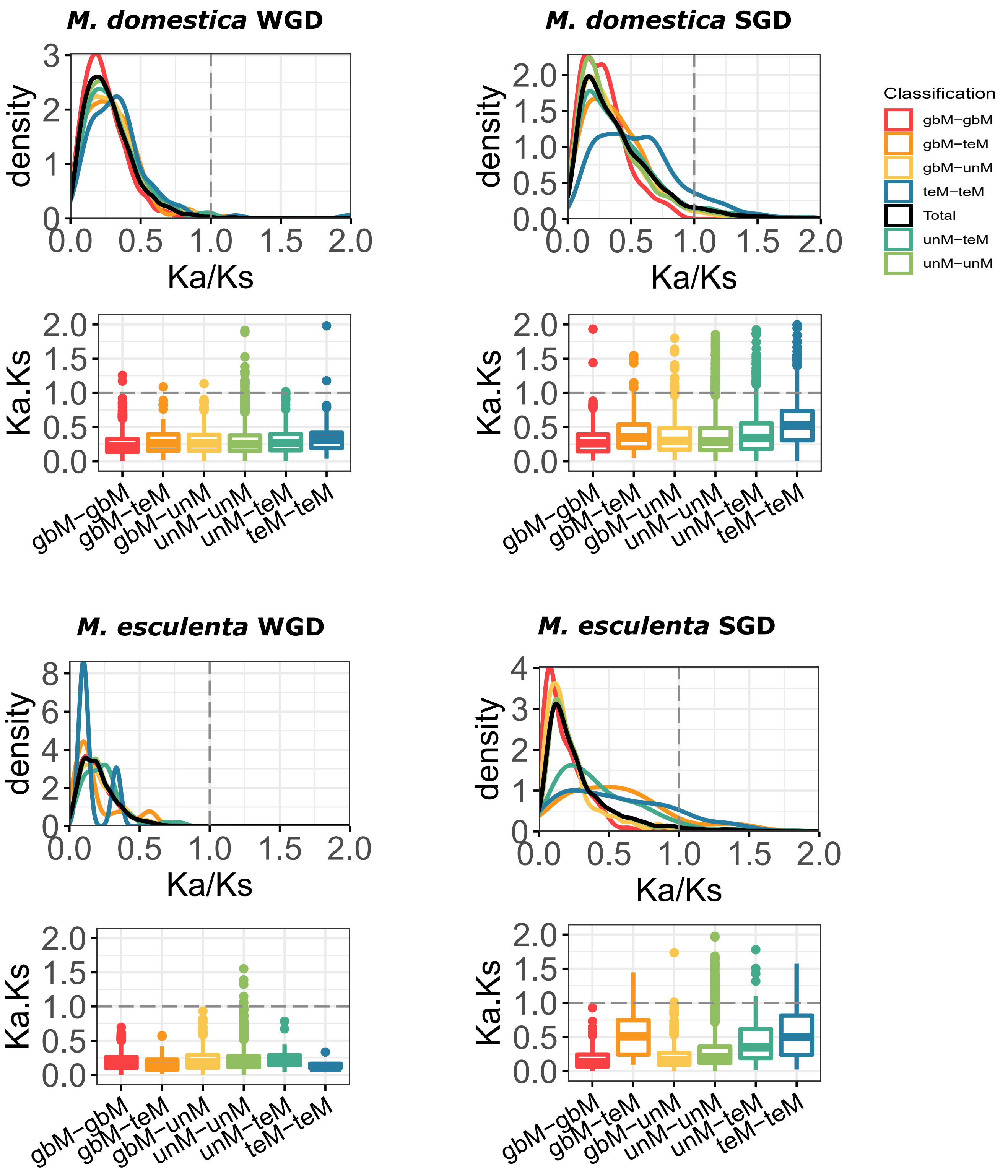

**Figure S10**: Distribution of genic methylation classified genes based on the ratio of nonsynonymous substitution (Ka), with synonymous substitutions (Ks) across different types of gene duplicate pairs.
Whole-genome duplicates - WGD, Single-gene duplicates - SGD (combined data from tandem, proximal, translocated, and dispersed duplicates).

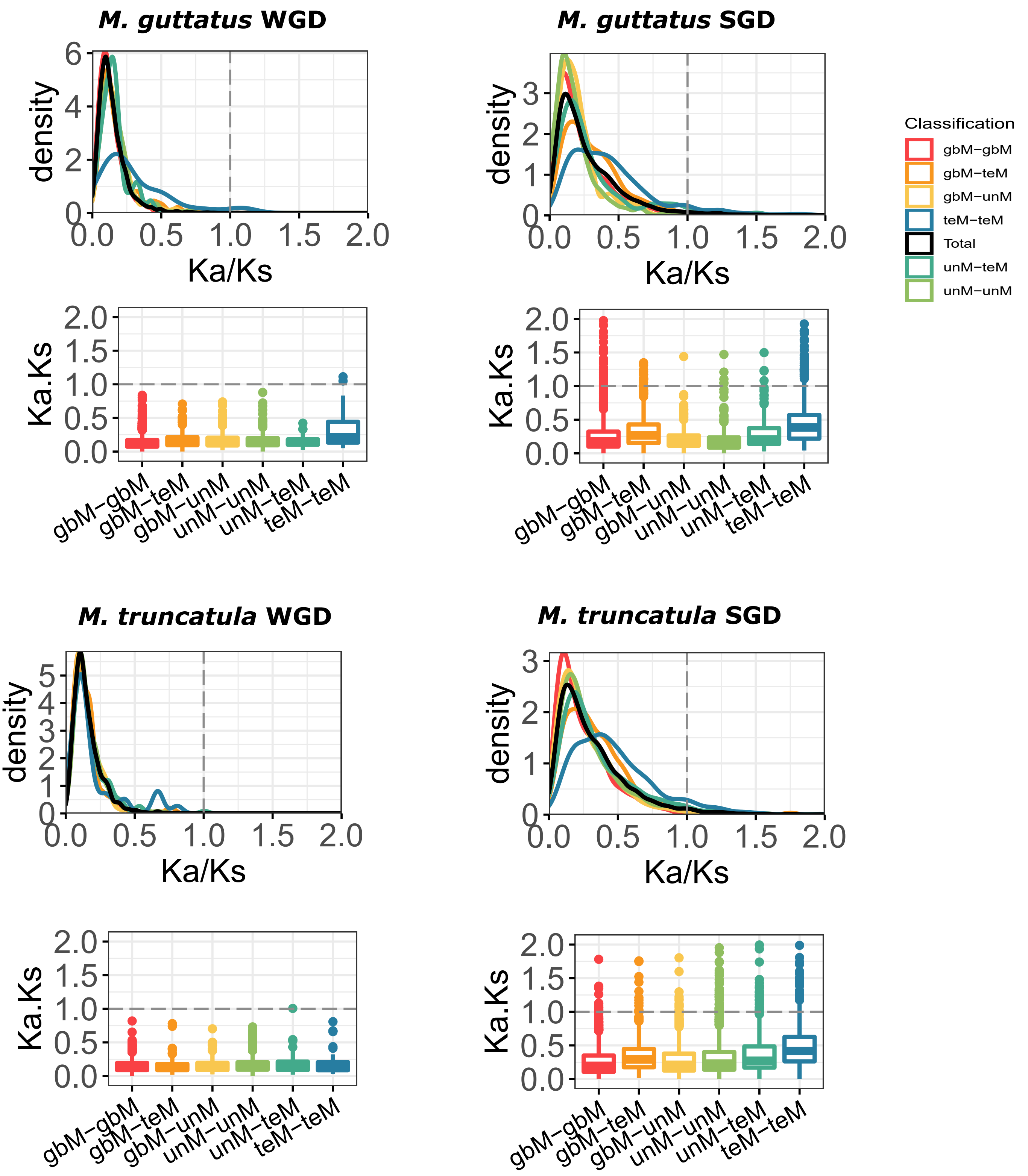

**Figure S10**: Distribution of genic methylation classified genes based on the ratio of nonsynonymous substitution (Ka), with synonymous substitutions (Ks) across different types of gene duplicate pairs.
Whole-genome duplicates - WGD, Single-gene duplicates - SGD (combined data from tandem, proximal, translocated, and dispersed duplicates).

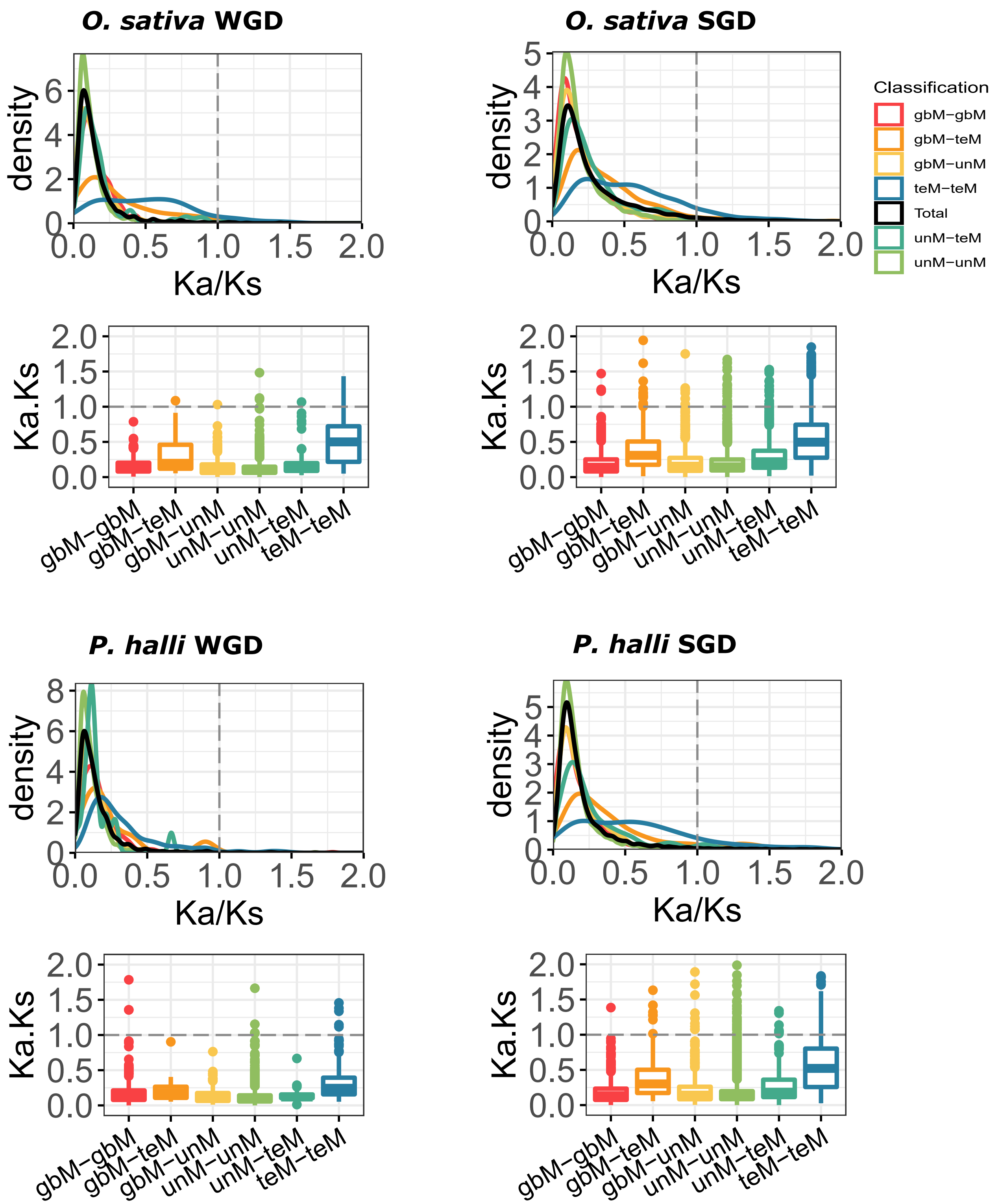

**Figure S10**: Distribution of genic methylation classified genes based on the ratio of nonsynonymous substitution (Ka), with synonymous substitutions (Ks) across different types of gene duplicate pairs.
Whole-genome duplicates - WGD, Single-gene duplicates - SGD (combined data from tandem, proximal, translocated, and dispersed duplicates).

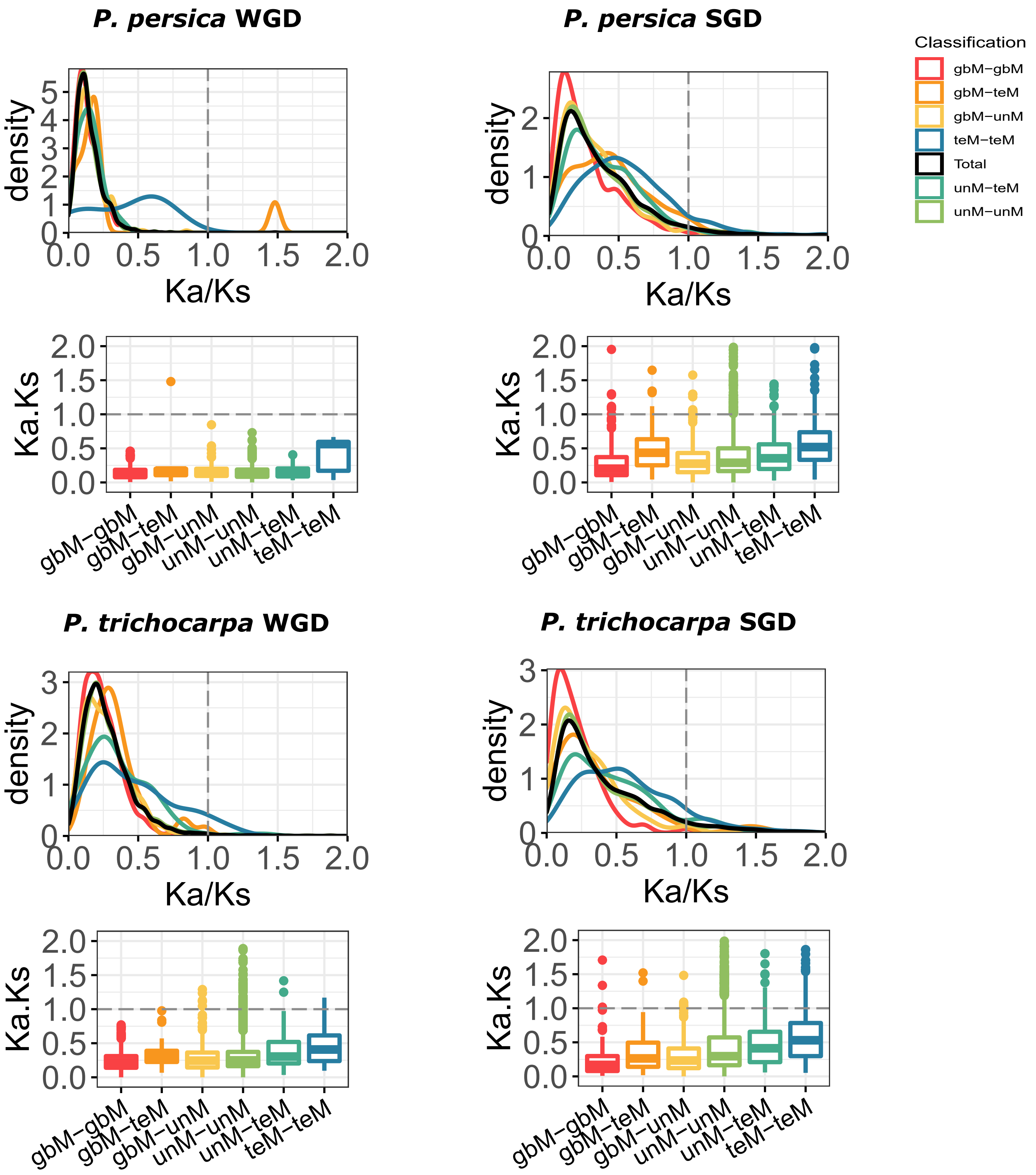

**Figure S10**: Distribution of genic methylation classified genes based on the ratio of nonsynonymous substitution (Ka), with synonymous substitutions (Ks) across different types of gene duplicate pairs.
Whole-genome duplicates - WGD, Single-gene duplicates - SGD (combined data from tandem, proximal, translocated, and dispersed duplicates).

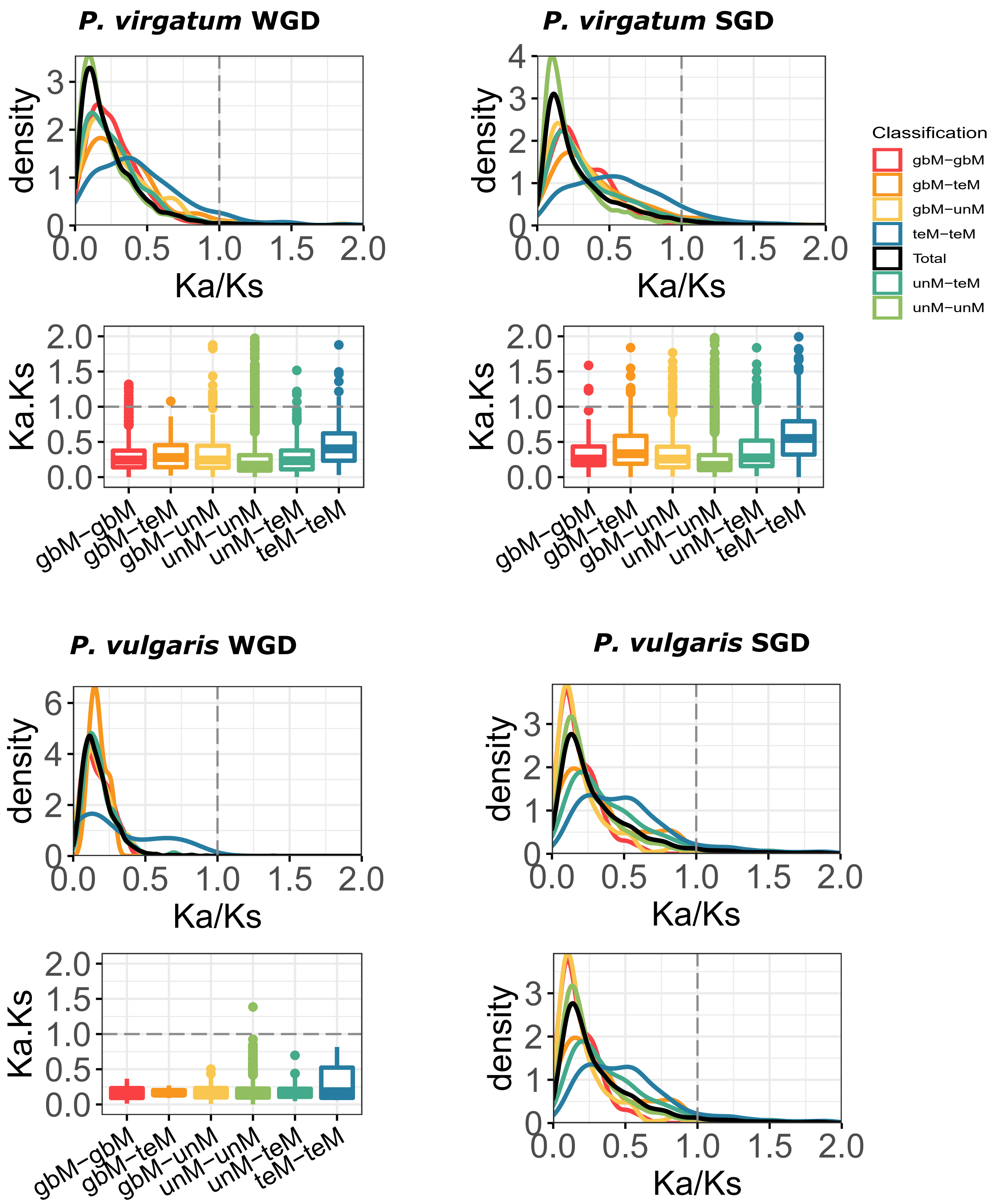

**Figure S10**: Distribution of genic methylation classified genes based on the ratio of nonsynonymous substitution (Ka), with synonymous substitutions (Ks) across different types of gene duplicate pairs.
Whole-genome duplicates - WGD, Single-gene duplicates - SGD (combined data from tandem, proximal, translocated, and dispersed duplicates).

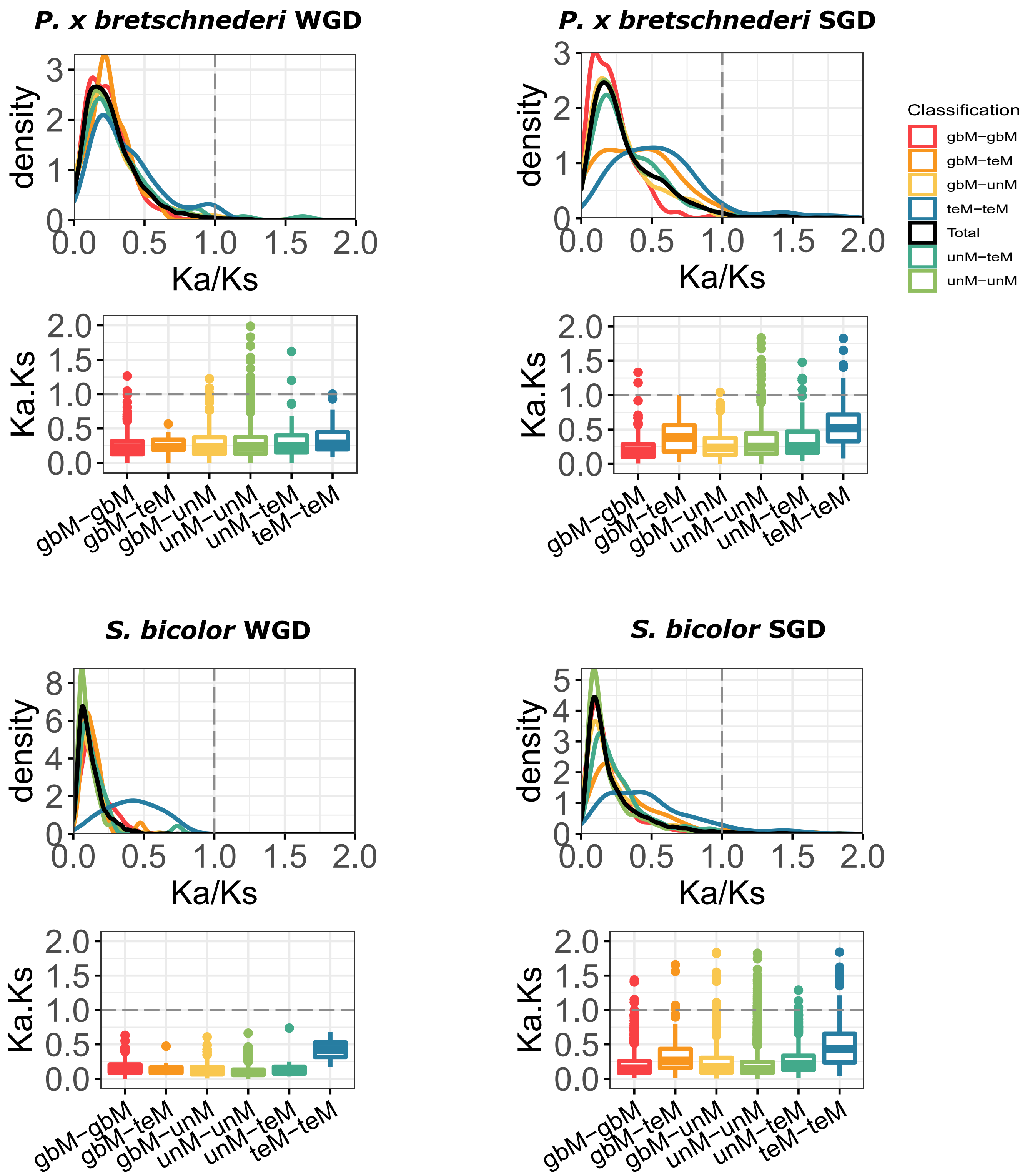

**Figure S10**: Distribution of genic methylation classified genes based on the ratio of nonsynonymous substitution (Ka), with synonymous substitutions (Ks) across different types of gene duplicate pairs.
Whole-genome duplicates - WGD, Single-gene duplicates - SGD (combined data from tandem, proximal, translocated, and dispersed duplicates).

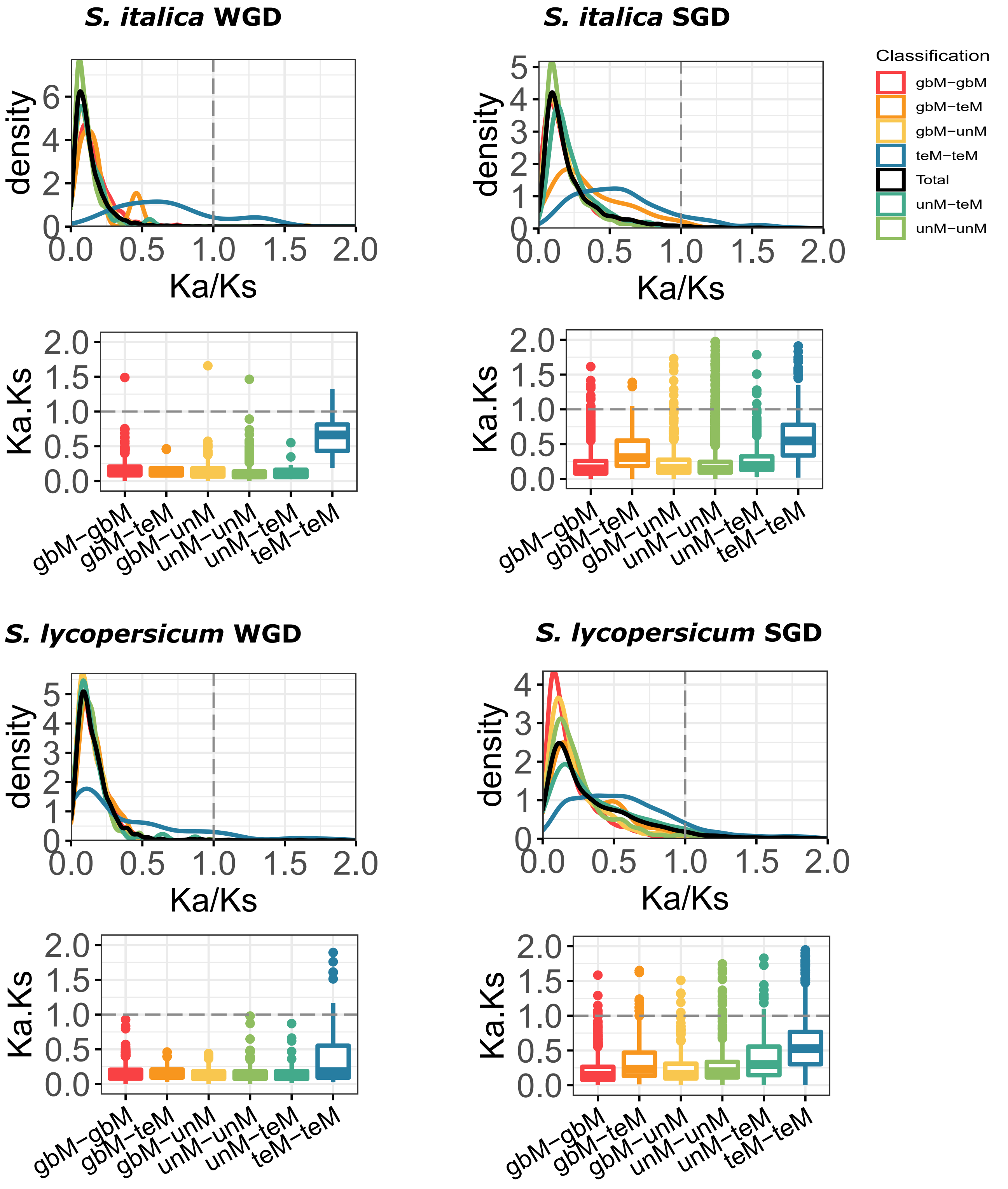

**Figure S10**: Distribution of genic methylation classified genes based on the ratio of nonsynonymous substitution (Ka), with synonymous substitutions (Ks) across different types of gene duplicate pairs.
Whole-genome duplicates - WGD, Single-gene duplicates - SGD (combined data from tandem, proximal, translocated, and dispersed duplicates).

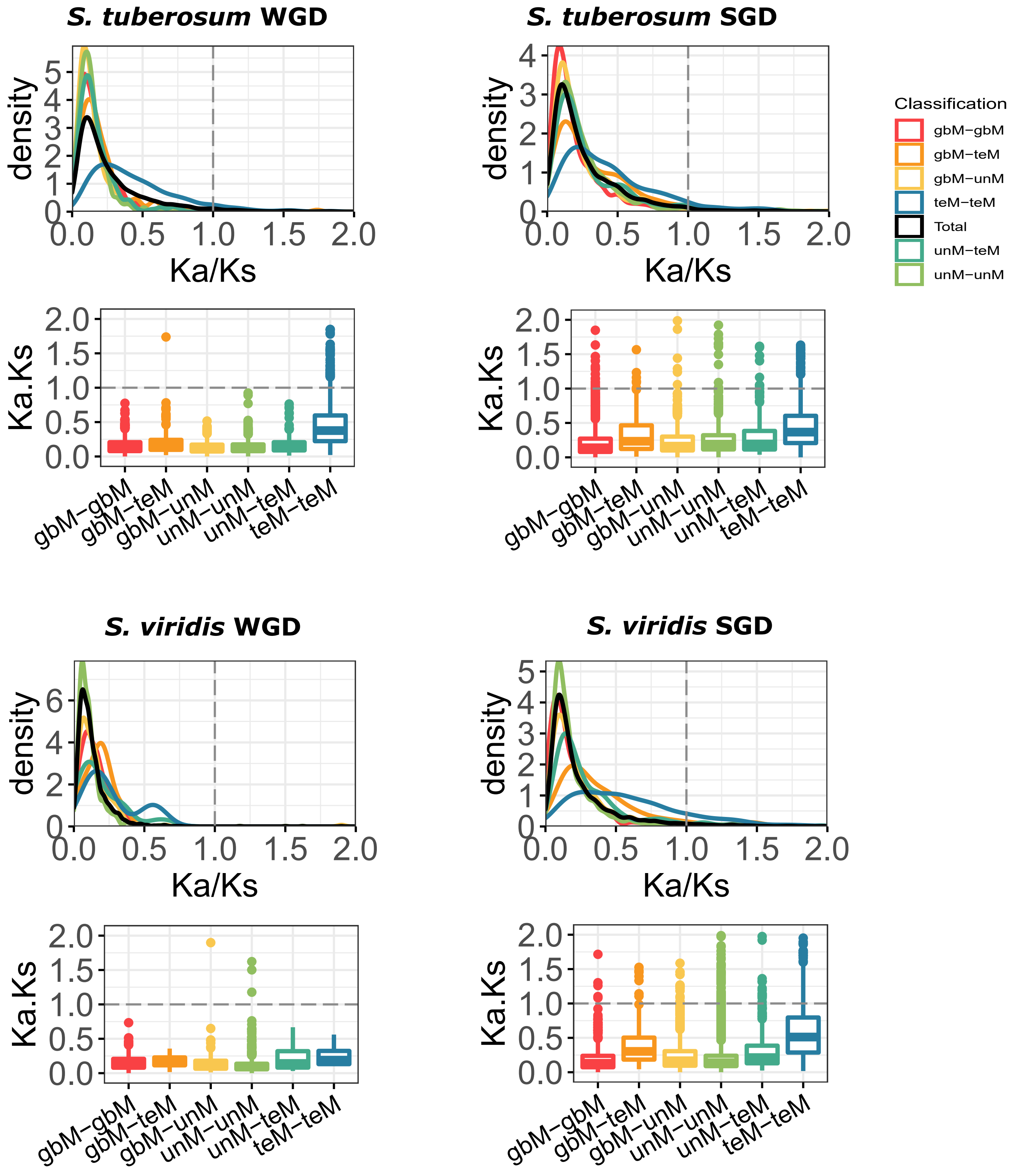

**Figure S10**: Distribution of genic methylation classified genes based on the ratio of nonsynonymous substitution (Ka), with synonymous substitutions (Ks) across different types of gene duplicate pairs.
Whole-genome duplicates - WGD, Single-gene duplicates - SGD (combined data from tandem, proximal, translocated, and dispersed duplicates).

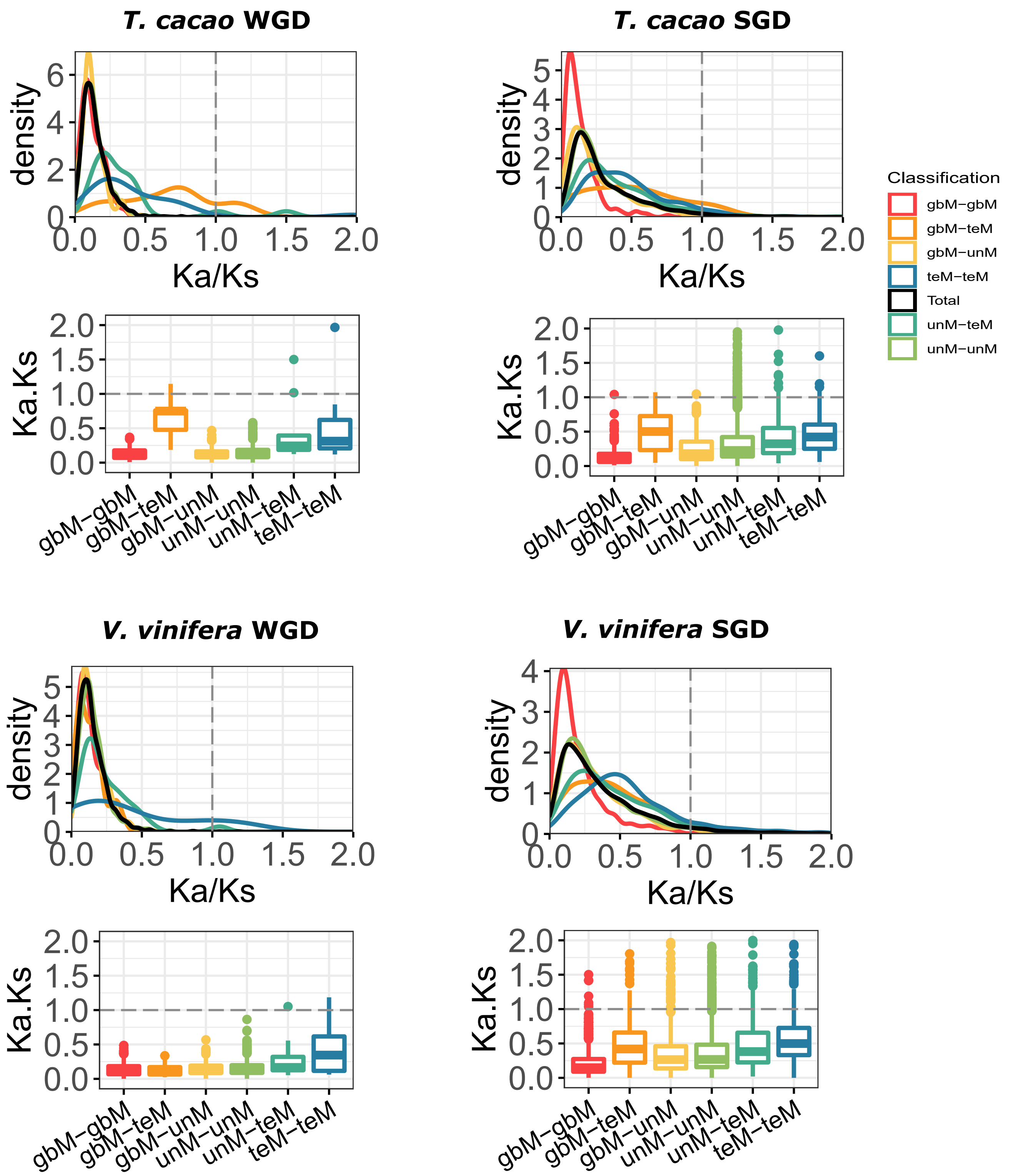

**Figure S10**: Distribution of genic methylation classified genes based on the ratio of nonsynonymous substitution (Ka), with synonymous substitutions (Ks) across different types of gene duplicate pairs.
Whole-genome duplicates - WGD, Single-gene duplicates - SGD (combined data from tandem, proximal, translocated, and dispersed duplicates).

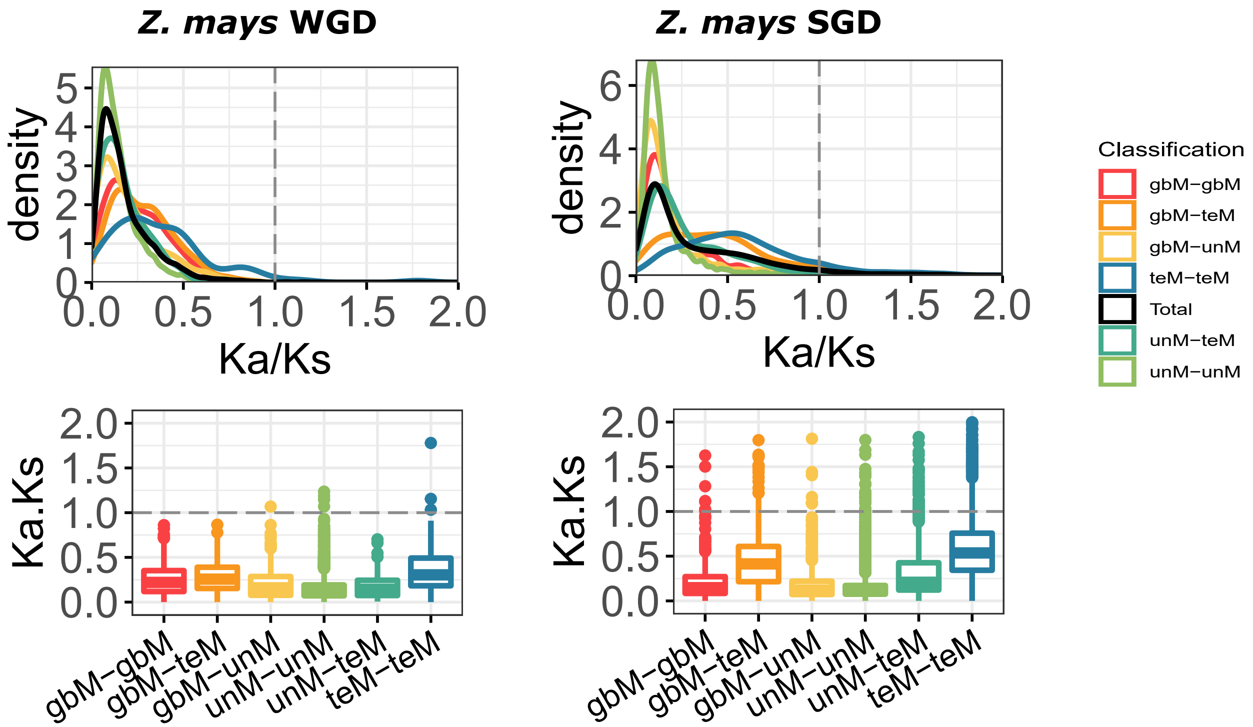

**Figure S11**: Distribution of core: multi-copy and core: single-copy (intermediate) paralogs based on the ratio of nonsynonymous substitution (Ka), with synonymous substitutions (Ks). 'All pairs' represent Ka/Ks ratios of all duplicate gene pairs in the genome, 'Core-MC' represents duplicate pairs among core: multi-copy orthogroup, while 'SC-Int' represent duplicate pairs among the core: single-copy orthogroups.

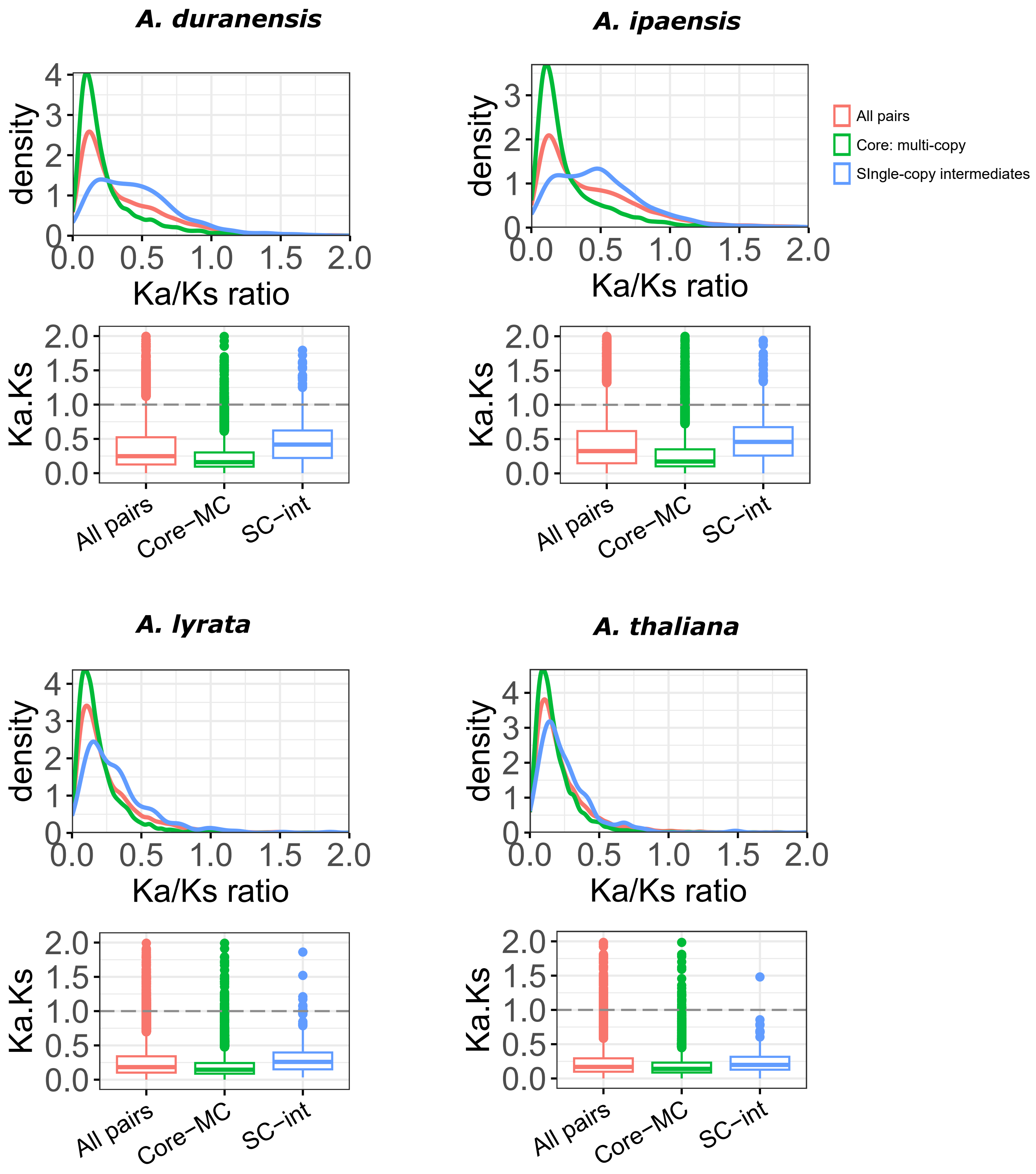

**Figure S11**: Distribution of core: multi-copy and core: single-copy (intermediate) paralogs based on the ratio of nonsynonymous substitution (Ka), with synonymous substitutions (Ks). 'All pairs' represent Ka/Ks ratios of all duplicate gene pairs in the genome, 'Core-MC' represents duplicate pairs among core: multi-copy orthogroup, while 'SC-Int' represent duplicate pairs among the core: single-copy orthogroups.

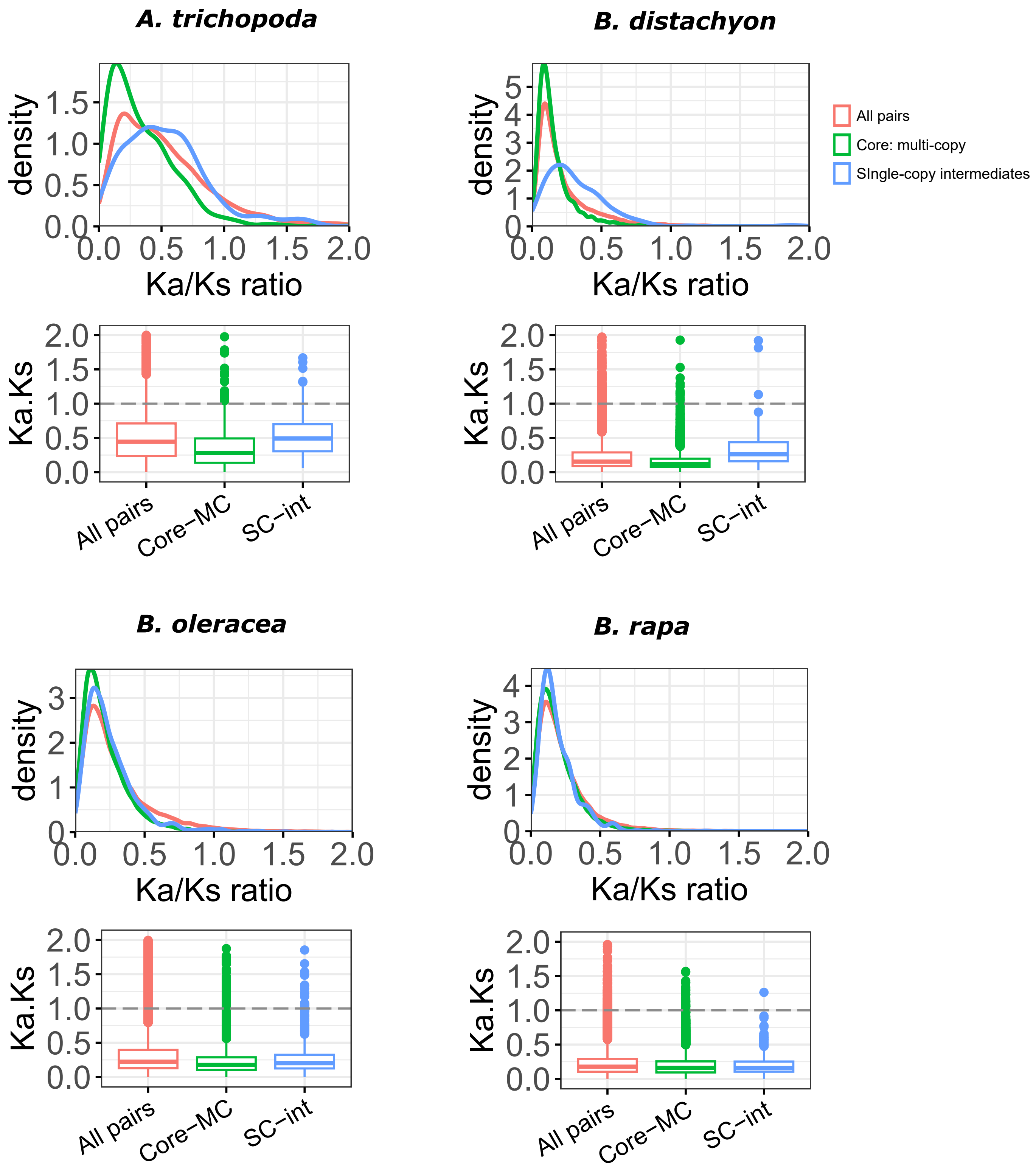

**Figure S11**: Distribution of core: multi-copy and core: single-copy (intermediate) paralogs based on the ratio of nonsynonymous substitution (Ka), with synonymous substitutions (Ks). 'All pairs' represent Ka/Ks ratios of all duplicate gene pairs in the genome, 'Core-MC' represents duplicate pairs among core: multi-copy orthogroup, while 'SC-Int' represent duplicate pairs among the core: single-copy orthogroups.

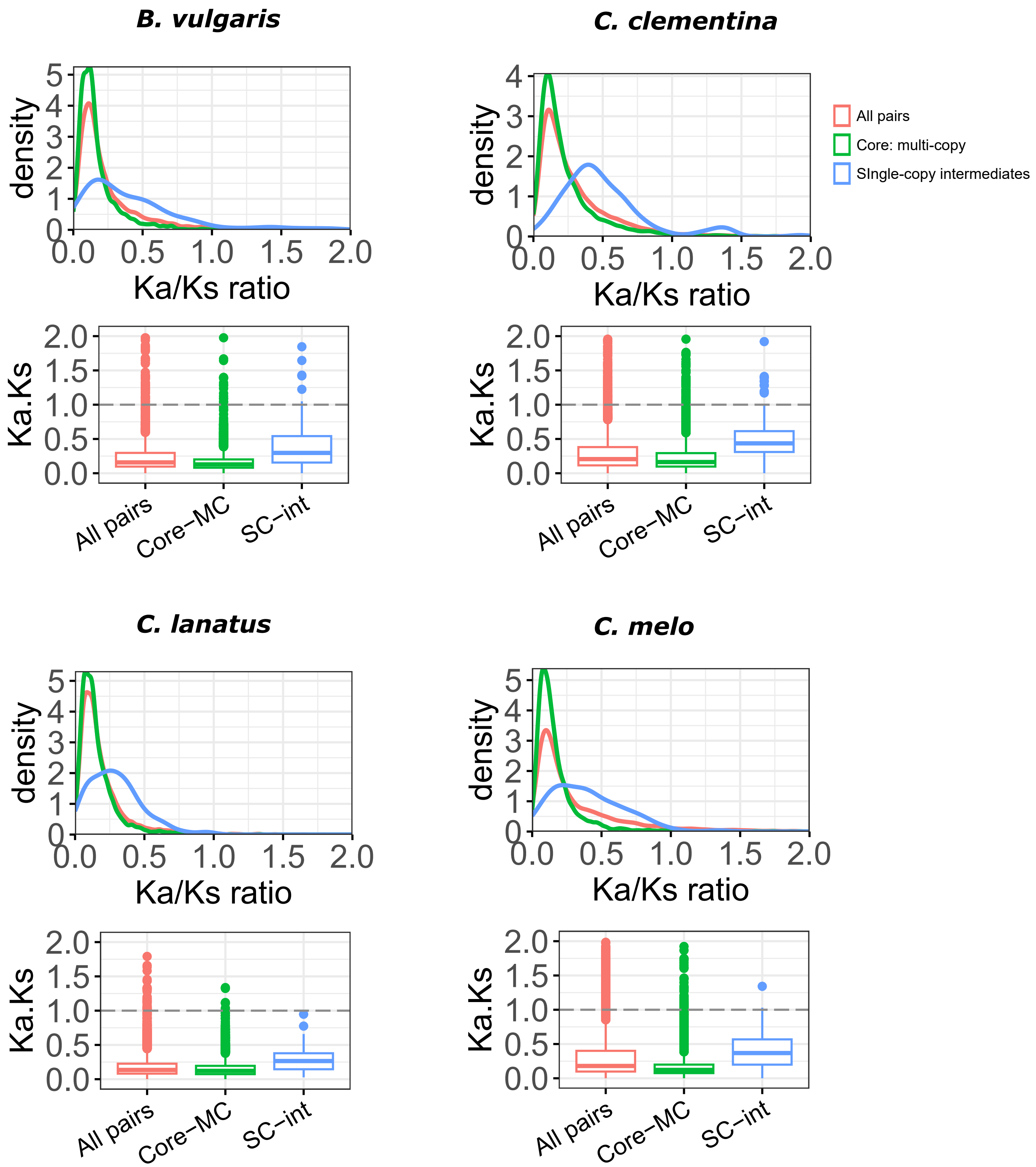

**Figure S11**: Distribution of core: multi-copy and core: single-copy (intermediate) paralogs based on the ratio of nonsynonymous substitution (Ka), with synonymous substitutions (Ks). 'All pairs' represent Ka/Ks ratios of all duplicate gene pairs in the genome, 'Core-MC' represents duplicate pairs among core: multi-copy orthogroup, while 'SC-Int' represent duplicate pairs among the core: single-copy orthogroups.

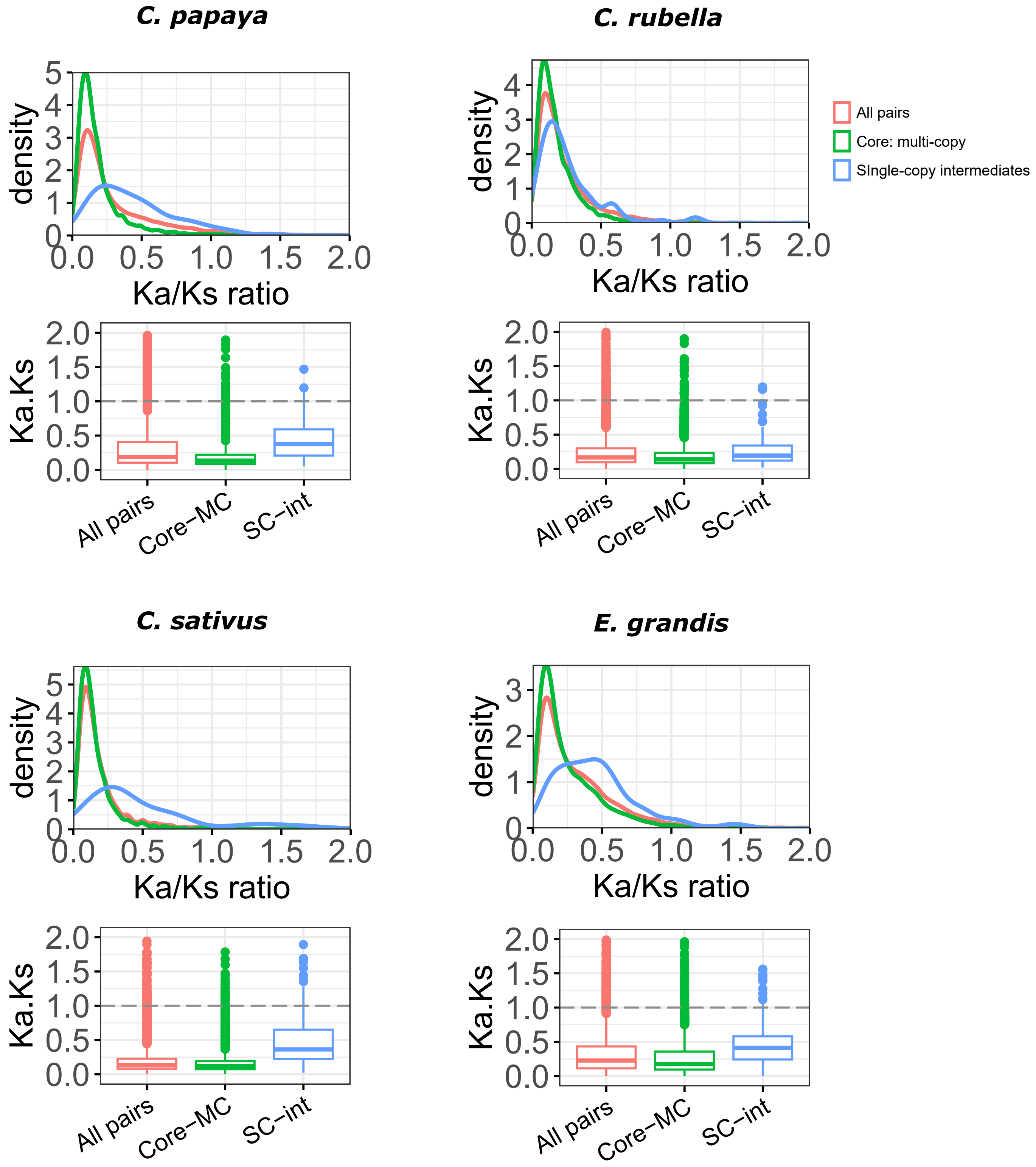

**Figure S11**: Distribution of core: multi-copy and core: single-copy (intermediate) paralogs based on the ratio of nonsynonymous substitution (Ka), with synonymous substitutions (Ks). 'All pairs' represent Ka/Ks ratios of all duplicate gene pairs in the genome, 'Core-MC' represents duplicate pairs among core: multi-copy orthogroup, while 'SC-Int' represent duplicate pairs among the core: single-copy orthogroups.

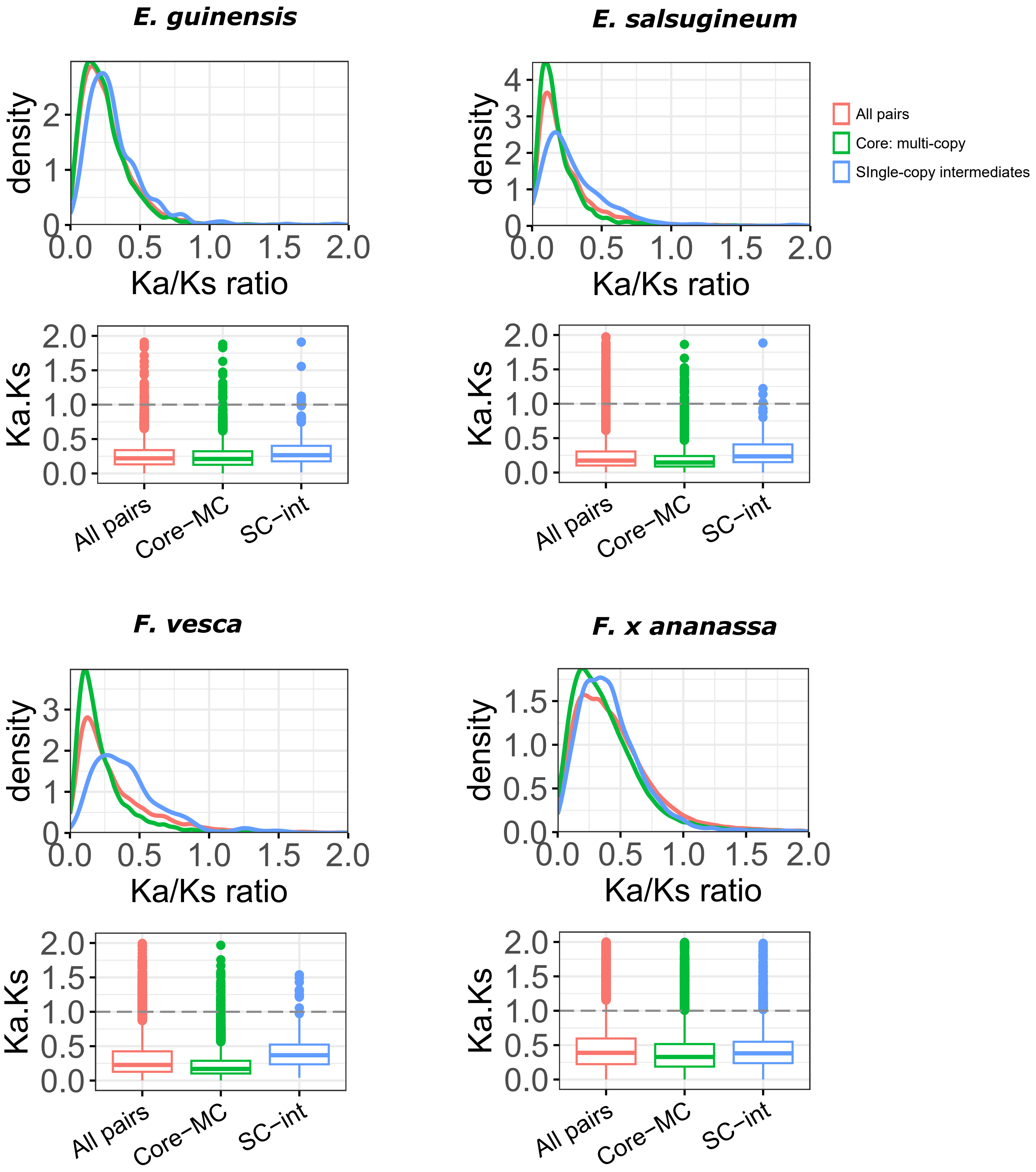

**Figure S11**: Distribution of core: multi-copy and core: single-copy (intermediate) paralogs based on the ratio of nonsynonymous substitution (Ka), with synonymous substitutions (Ks). 'All pairs' represent Ka/Ks ratios of all duplicate gene pairs in the genome, 'Core-MC' represents duplicate pairs among core: multi-copy orthogroup, while 'SC-Int' represent duplicate pairs among the core: single-copy orthogroups.

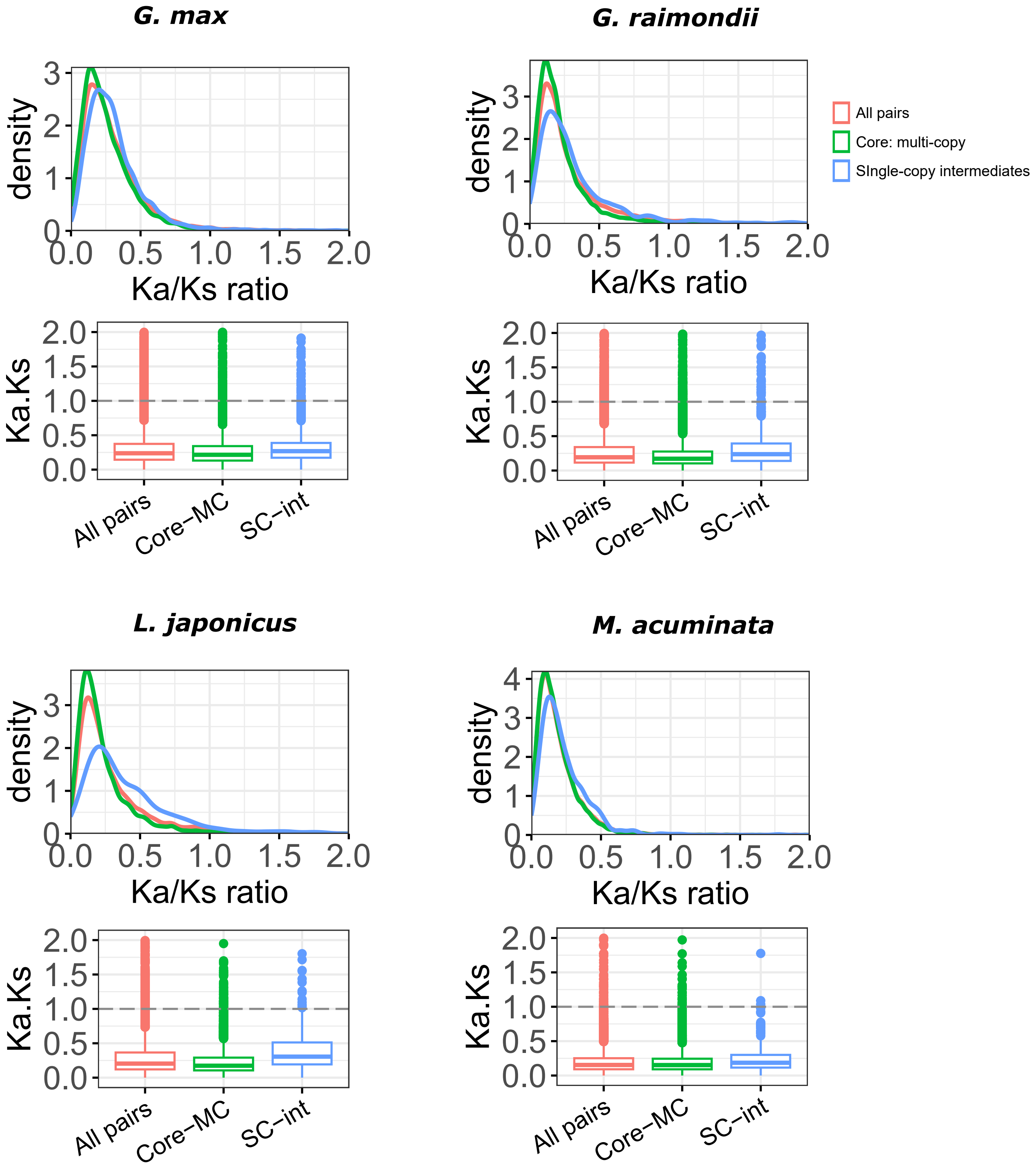

**Figure S11**: Distribution of core: multi-copy and core: single-copy (intermediate) paralogs based on the ratio of nonsynonymous substitution (Ka), with synonymous substitutions (Ks). 'All pairs' represent Ka/Ks ratios of all duplicate gene pairs in the genome, 'Core-MC' represents duplicate pairs among core: multi-copy orthogroup, while 'SC-Int' represent duplicate pairs among the core: single-copy orthogroups.

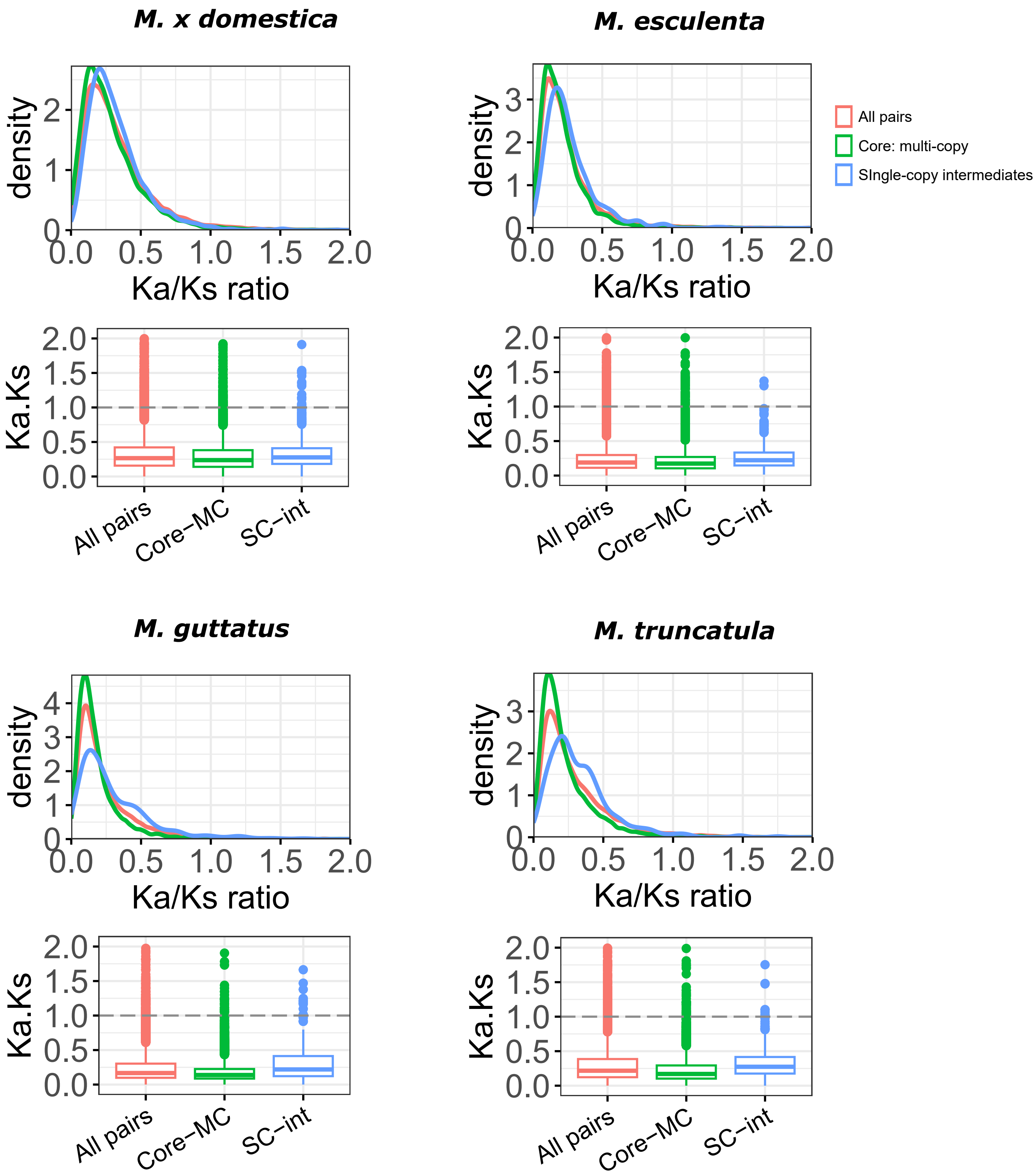

**Figure S11**: Distribution of core: multi-copy and core: single-copy (intermediate) paralogs based on the ratio of nonsynonymous substitution (Ka), with synonymous substitutions (Ks). 'All pairs' represent Ka/Ks ratios of all duplicate gene pairs in the genome, 'Core-MC' represents duplicate pairs among core: multi-copy orthogroup, while 'SC-Int' represent duplicate pairs among the core: single-copy orthogroups.

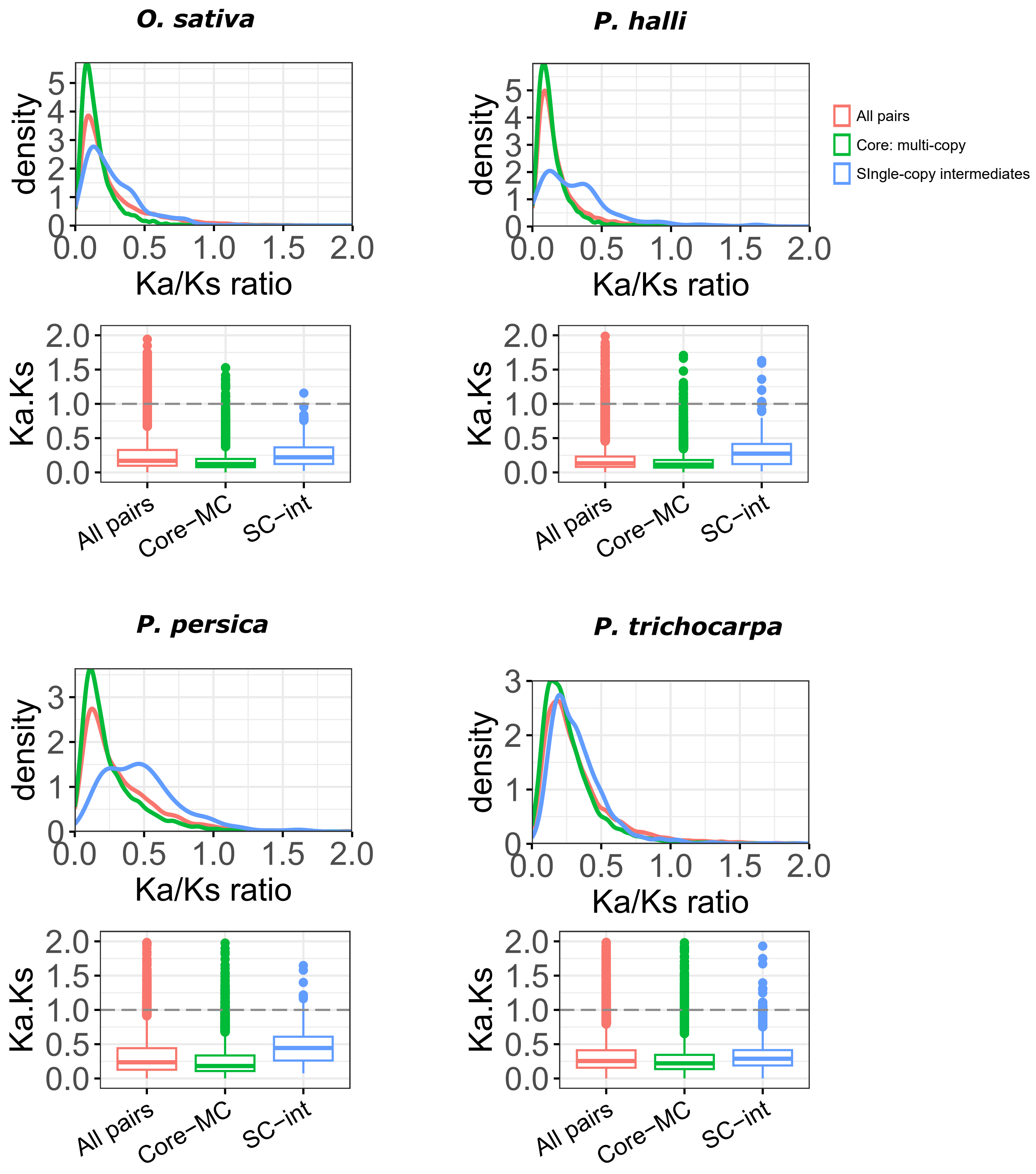

**Figure S11**: Distribution of core: multi-copy and core: single-copy (intermediate) paralogs based on the ratio of nonsynonymous substitution (Ka), with synonymous substitutions (Ks). 'All pairs' represent Ka/Ks ratios of all duplicate gene pairs in the genome, 'Core-MC' represents duplicate pairs among core: multi-copy orthogroup, while 'SC-Int' represent duplicate pairs among the core: single-copy orthogroups.

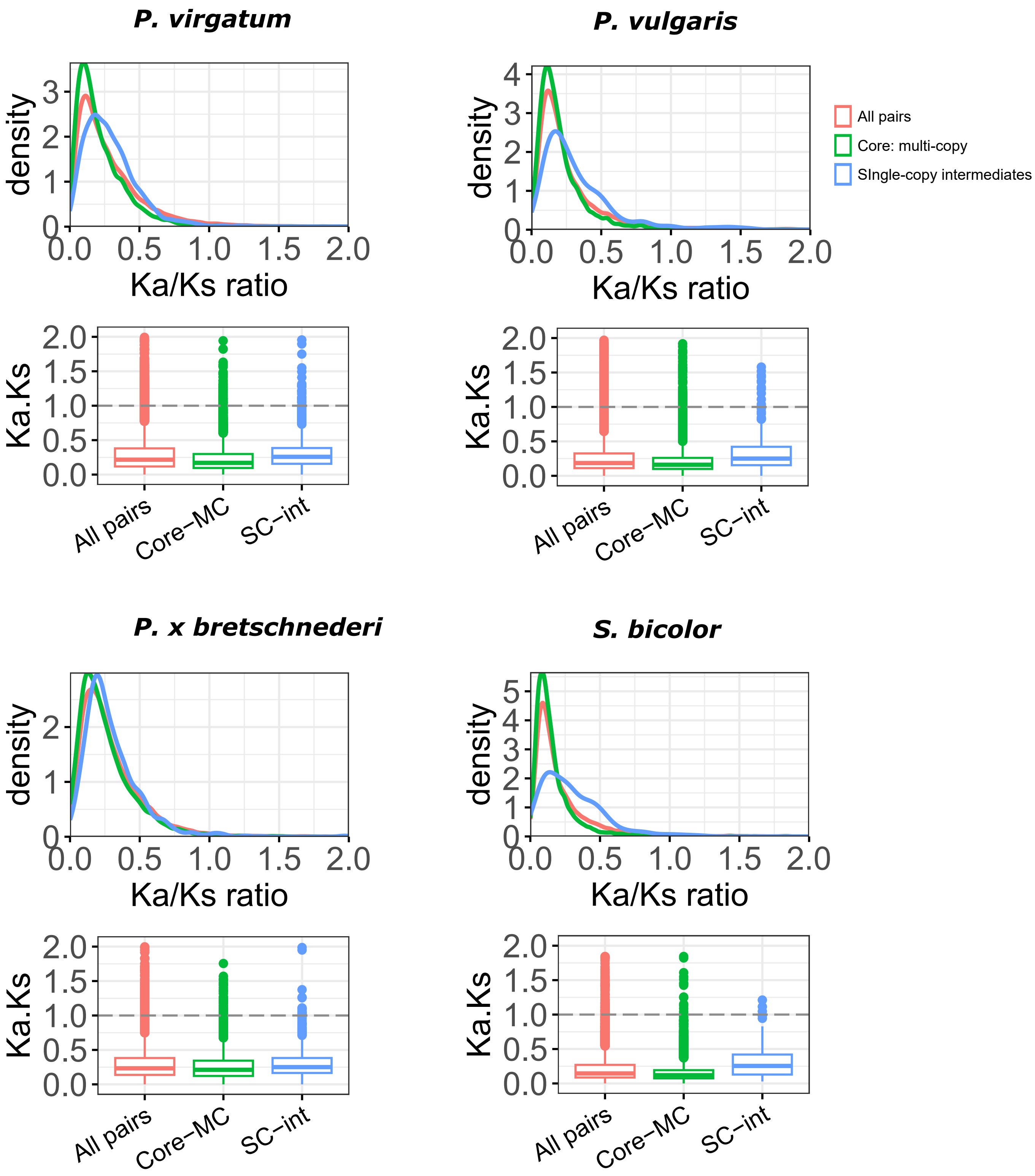

**Figure S11**: Distribution of core: multi-copy and core: single-copy (intermediate) paralogs based on the ratio of nonsynonymous substitution (Ka), with synonymous substitutions (Ks). 'All pairs' represent Ka/Ks ratios of all duplicate gene pairs in the genome, 'Core-MC' represents duplicate pairs among core: multi-copy orthogroup, while 'SC-Int' represent duplicate pairs among the core: single-copy orthogroups.

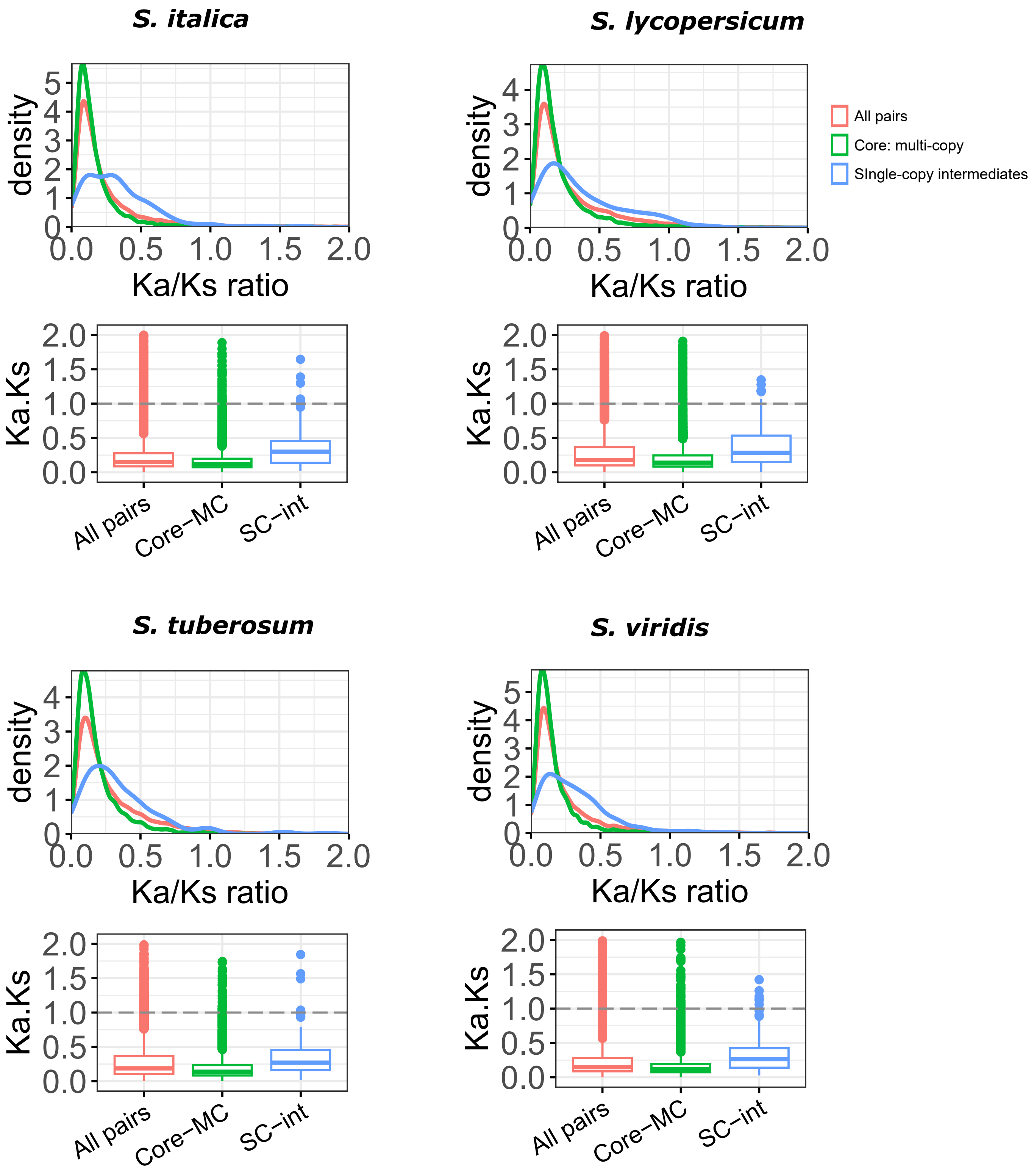

**Figure S11**: Distribution of core: multi-copy and core: single-copy (intermediate) paralogs based on the ratio of nonsynonymous substitution (Ka), with synonymous substitutions (Ks). 'All pairs' represent Ka/Ks ratios of all duplicate gene pairs in the genome, 'Core-MC' represents duplicate pairs among core: multi-copy orthogroup, while 'SC-Int' represent duplicate pairs among the core: single-copy orthogroups.

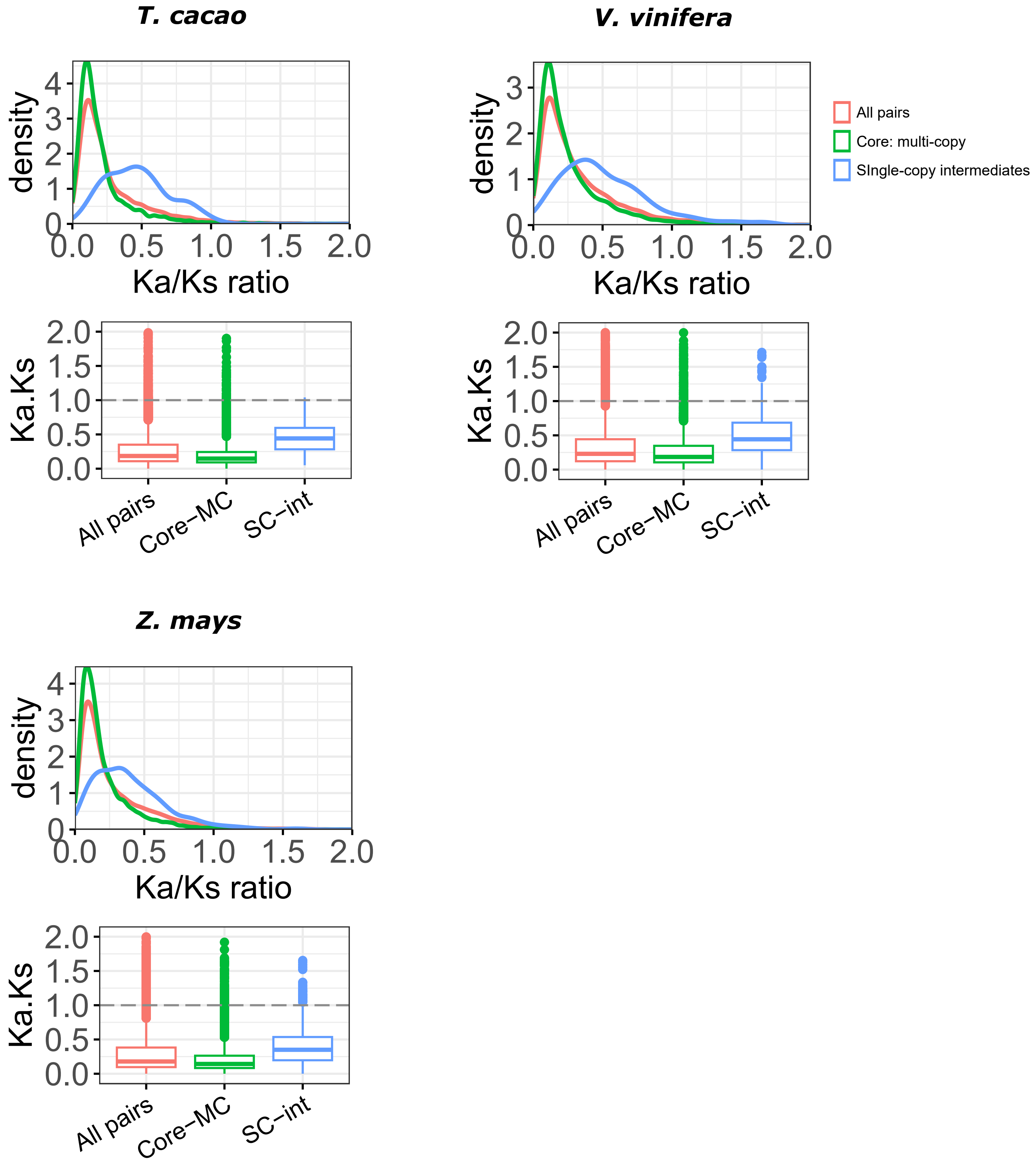

**Figure S12:** Percentage of Total (all genes), gbM, teM, and unM genes with known presence-absence variations. This plot was not restricted to duplicate genes, however the same results were found when limited to duplicates (Table S17). A two-sided Fisher's Exact Test was used to test for depletion or enrichment of PAVs amongst each category of genic methylation.
*FDR corrected p-value < 0.05, **FDR corrected p-value < 0.01,
***FDR corrected p-value < 0.001, NS – Not significantly different.

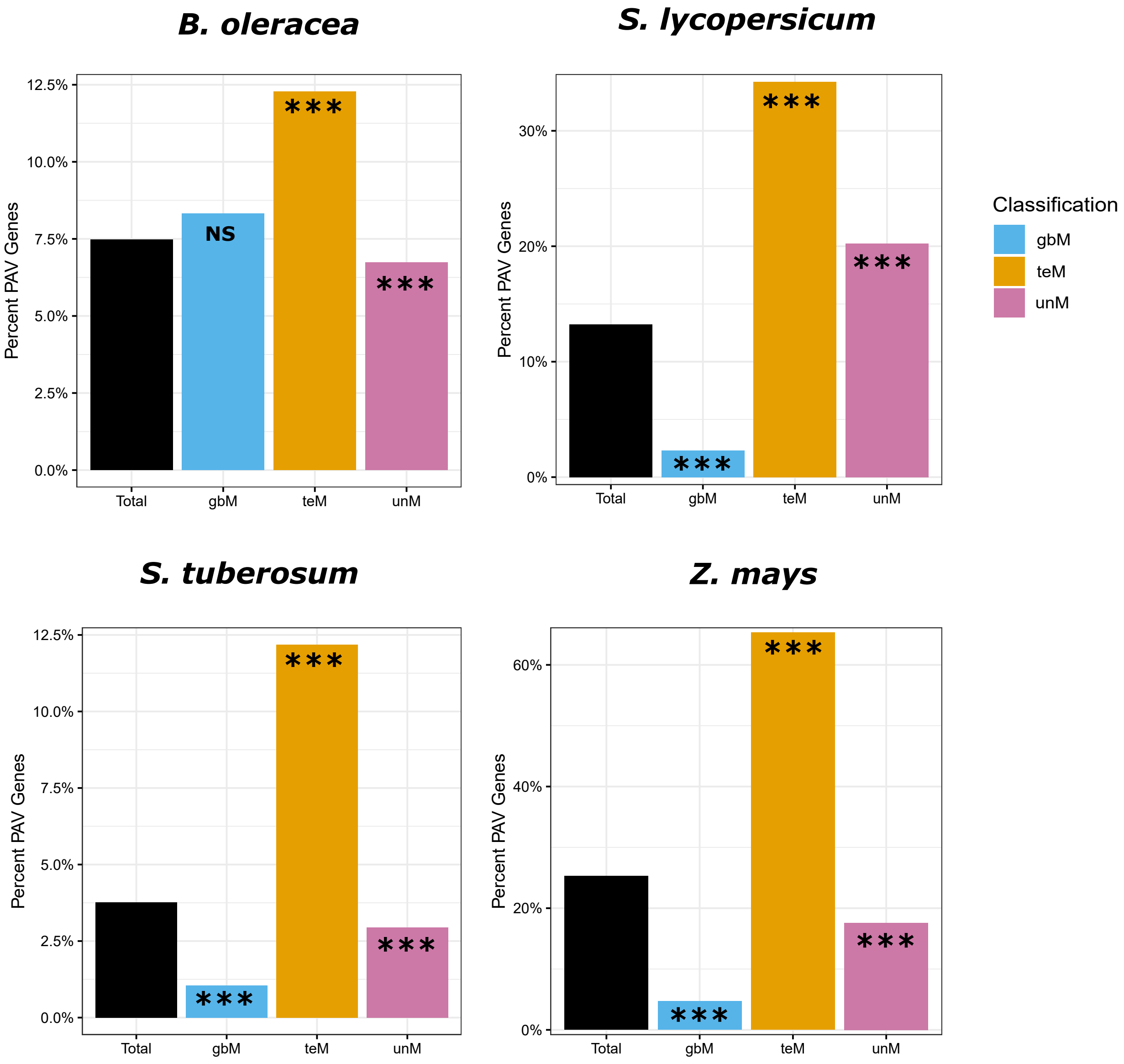

**Figure S13:** Proportion of single-copy singletons and single-copy intermediates in PAV genes in Boleraceae, Slycopersicum, Stuberosum, and Zmays. A proportion test was performed to test for differences in proportion of PAVs among single-copy singletons (SCs) and single-copy intermediates (SCi) compared to Core:multi-copy (Core-MC). '***' represent statistical significant difference at p-value < 0.001, NS – Not significantly different. Proportion of Core:MC, SCs,and SCi were signifcantly different from 'Total'.

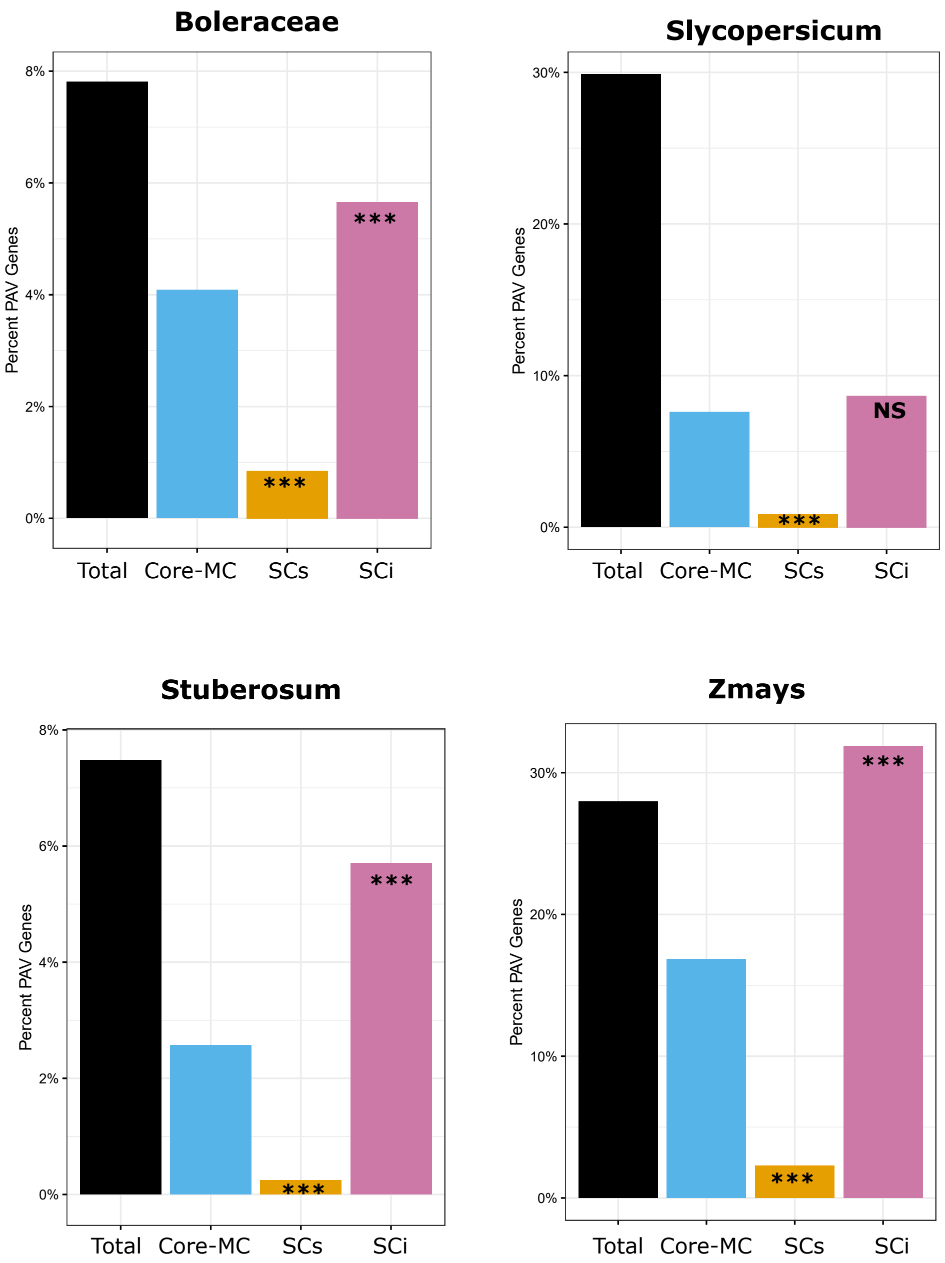

**Figure S14**: Tau specificty of gbM, teM, and unM genes in *G. max, P. vulgaris*, and *S. bicolor*.

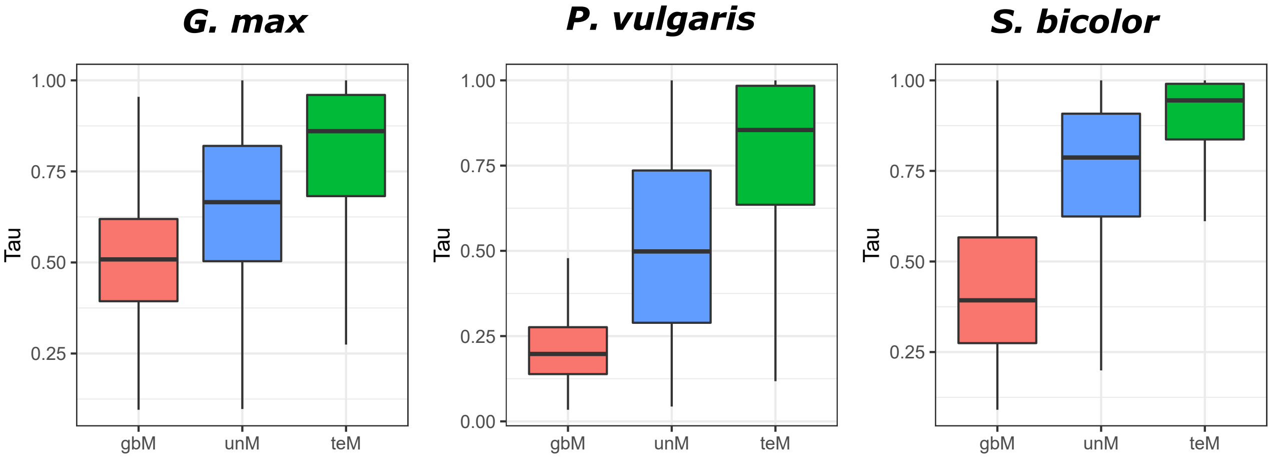

**Figure S15**: Tau specificty of gbM, teM, and unM genes each orthogroup category in *A. thaliana, G. max, P. vulgaris*, and *S. bicolor*.

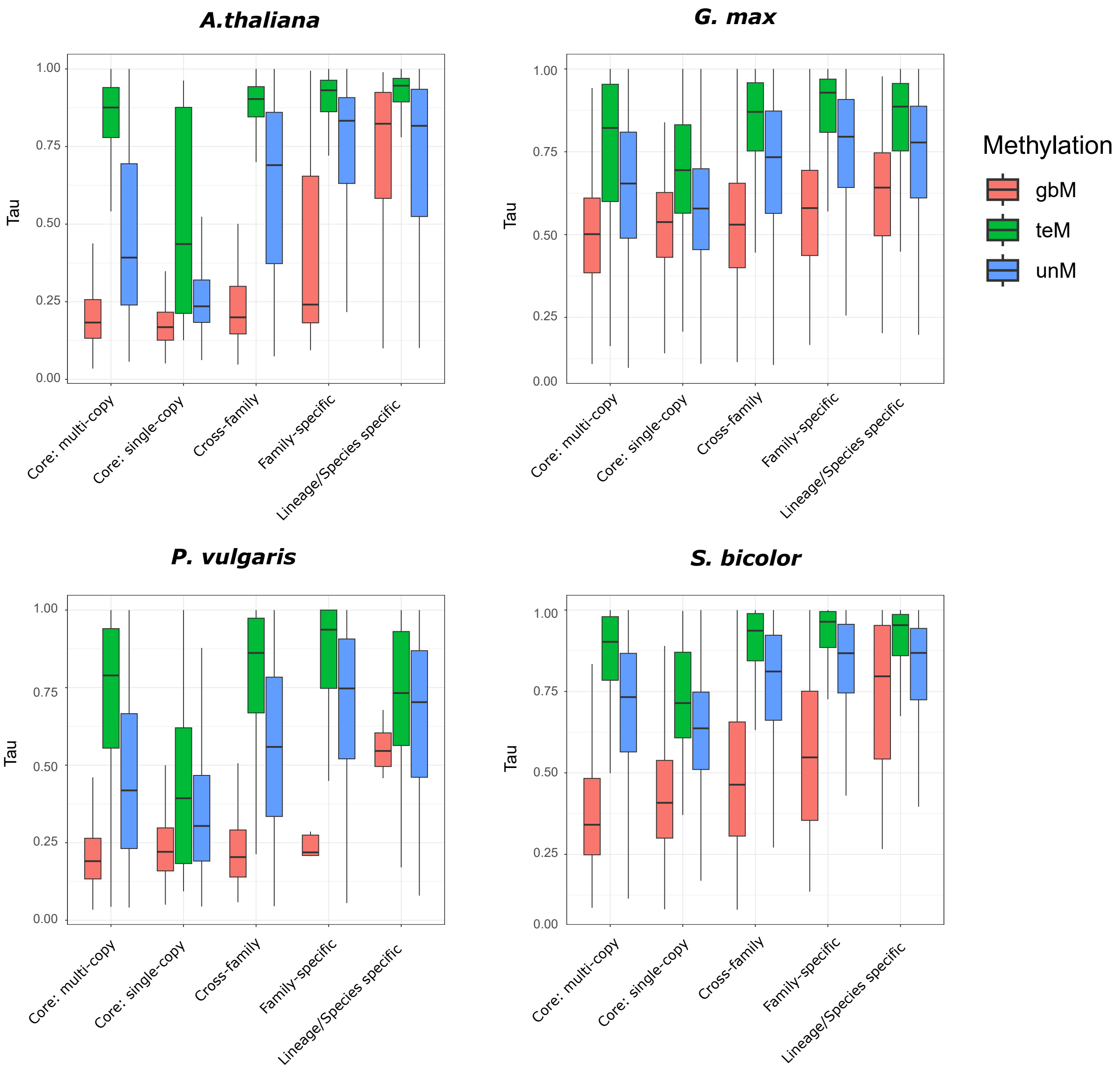

**Figure S16**: Tau specificities of different types of duplicate genes in A. thaliana, G. max, P. vulgaris, and S. bicolor. The distribution of tau for gbM, unM, and teM genes is shown for all duplicates and also broken down based on the type of duplicate gene.

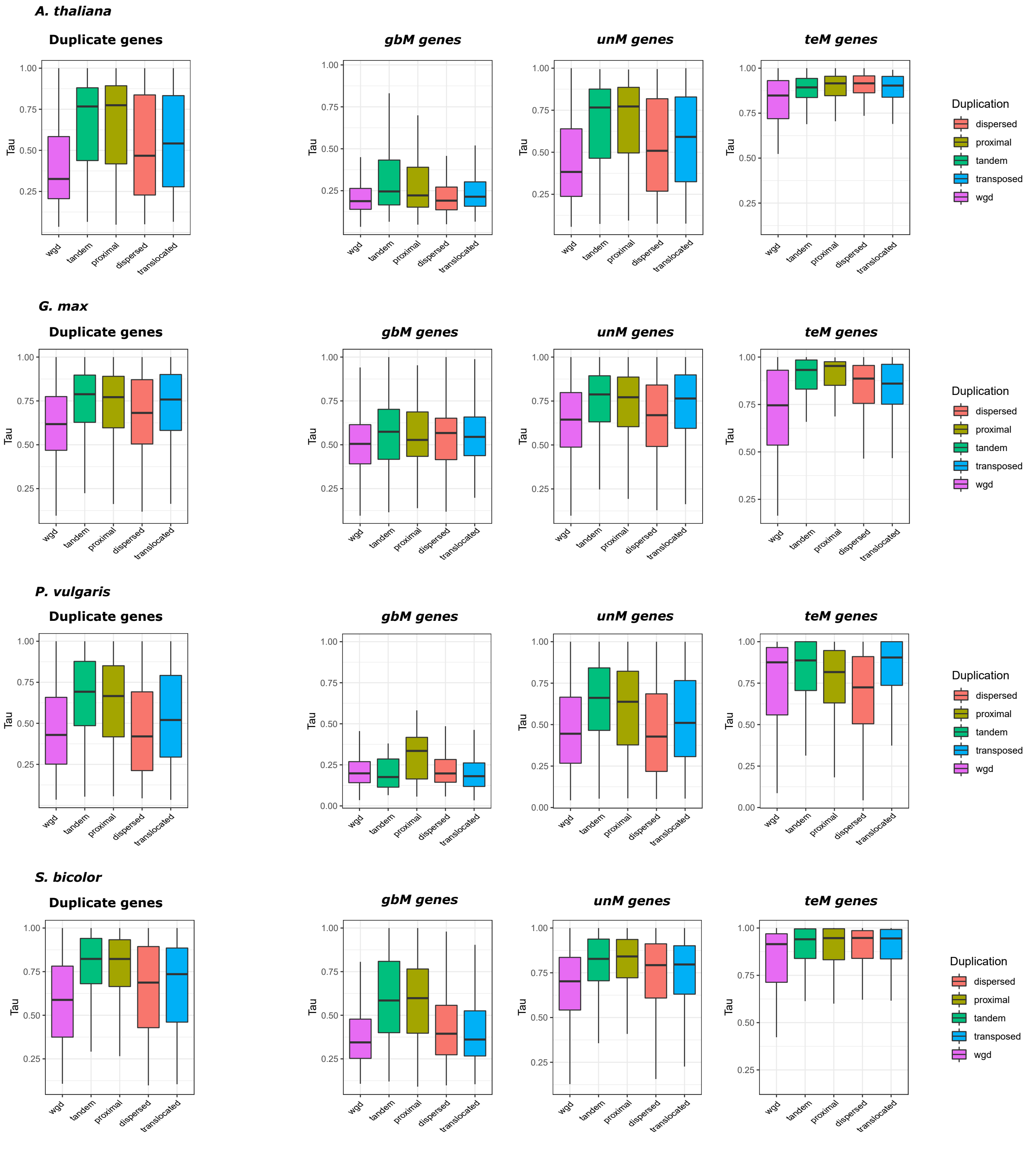

**Figure S17:** Half plots showing gene expression correlations of duplicate pairs based on genic methylation (gbM-gbM, gbM-teM, teM-teM, unM-unM, gbM-unM, and unM-teM) in *A. thaliana, G. max, P. vulgaris,* and *S. bicolor.*

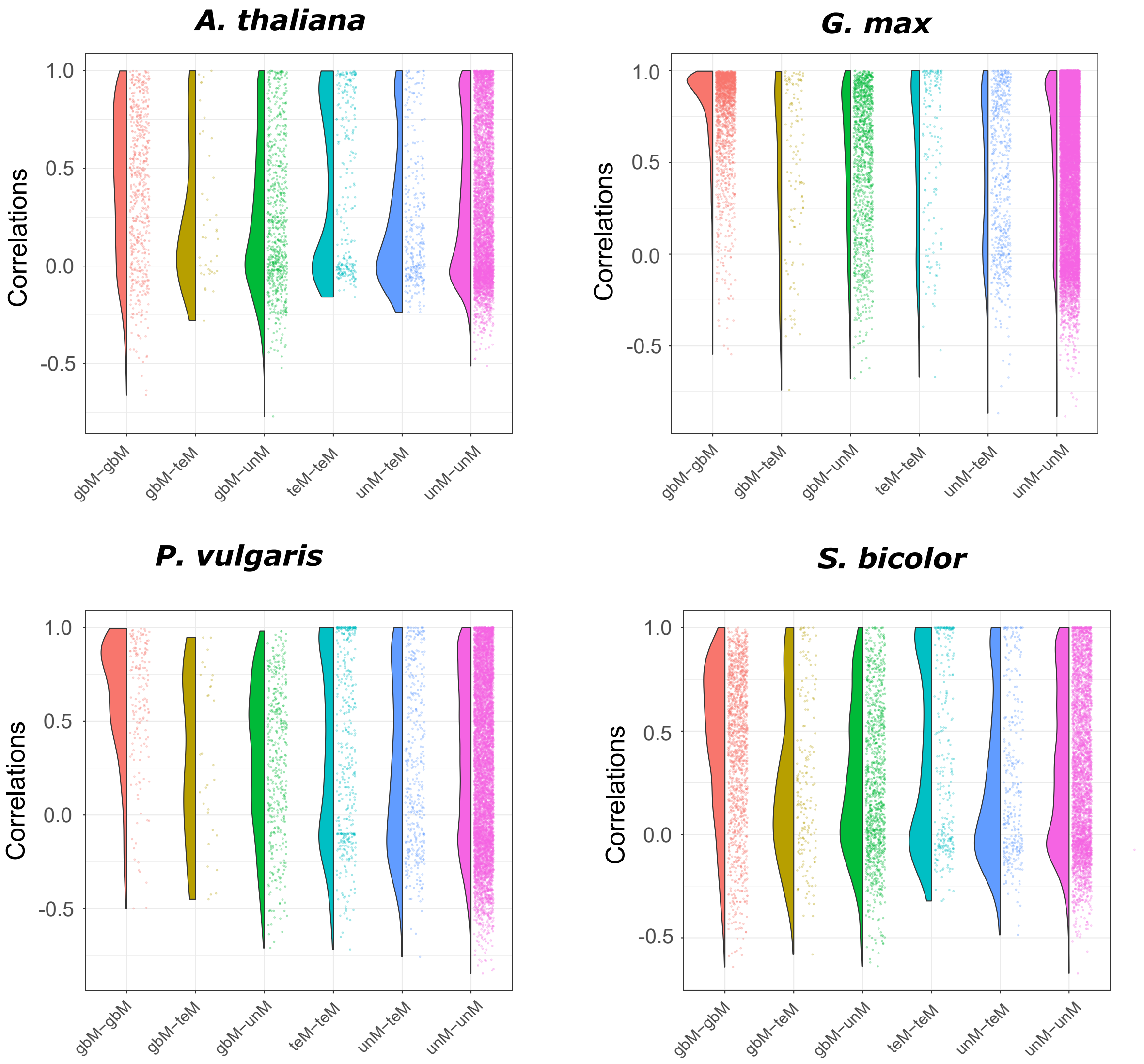

**Figure S18:** Absolute differences in Tau specificity between duplicate pairs in G. max, P. vulgaris, and S. bicolor. Data is broken down based on the genic methylation of the duplicate pairs (gbM-gbM, gbM-teM, teM-teM, unM-unM, gbM-unM, and unM-teM).

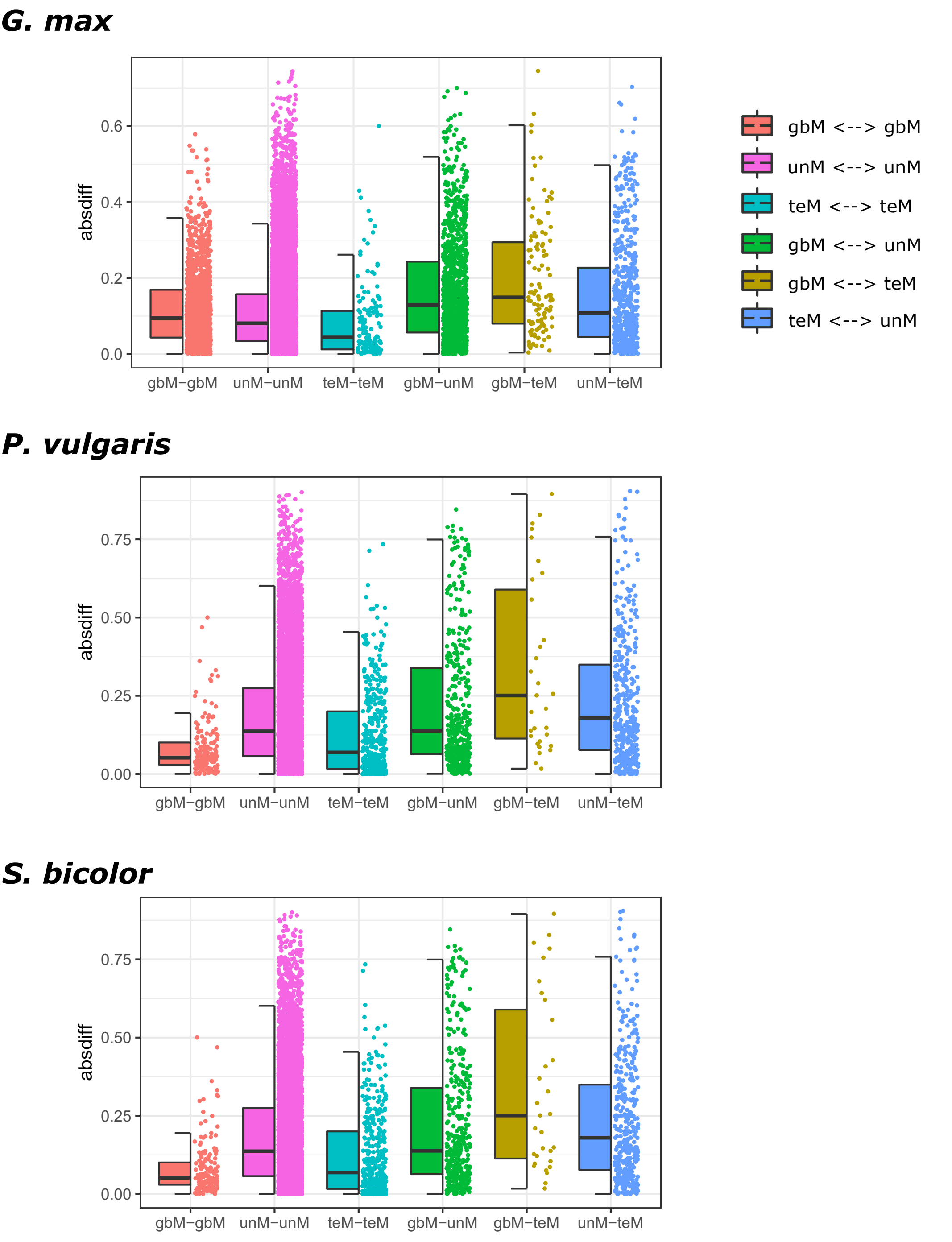

**Figure S19:** Distribution of Tau specificities for gbM, unM, and teM genes separated based on the methylation of their duplicate pair for G. max, P. vulgaris, and S. bicolor. For example, for gbM genes, the tau specificity was plotted for all gbM genes and the gbM paralog in gbM-gbM, gbM-teM, and gbM-unM pairs. For unM genes, the tau of only the unM paralog is shown and similarly for teM genes, only the tau of the teM paralog is shown.

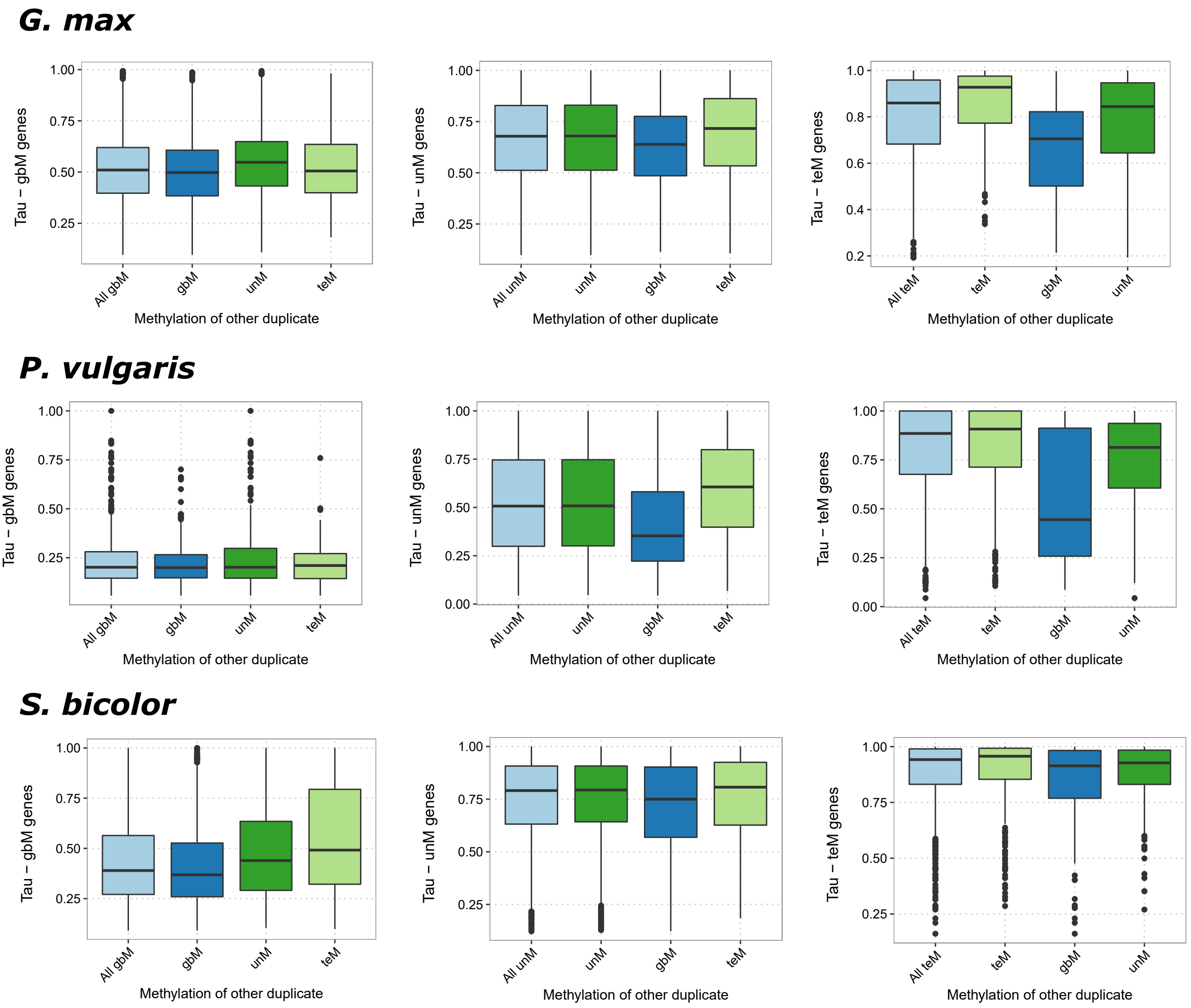

**Figure S20: Proportion of different orthogroup classiifaction and gbM/unM/teM epiallele frequency within *A. thaliana* population.**

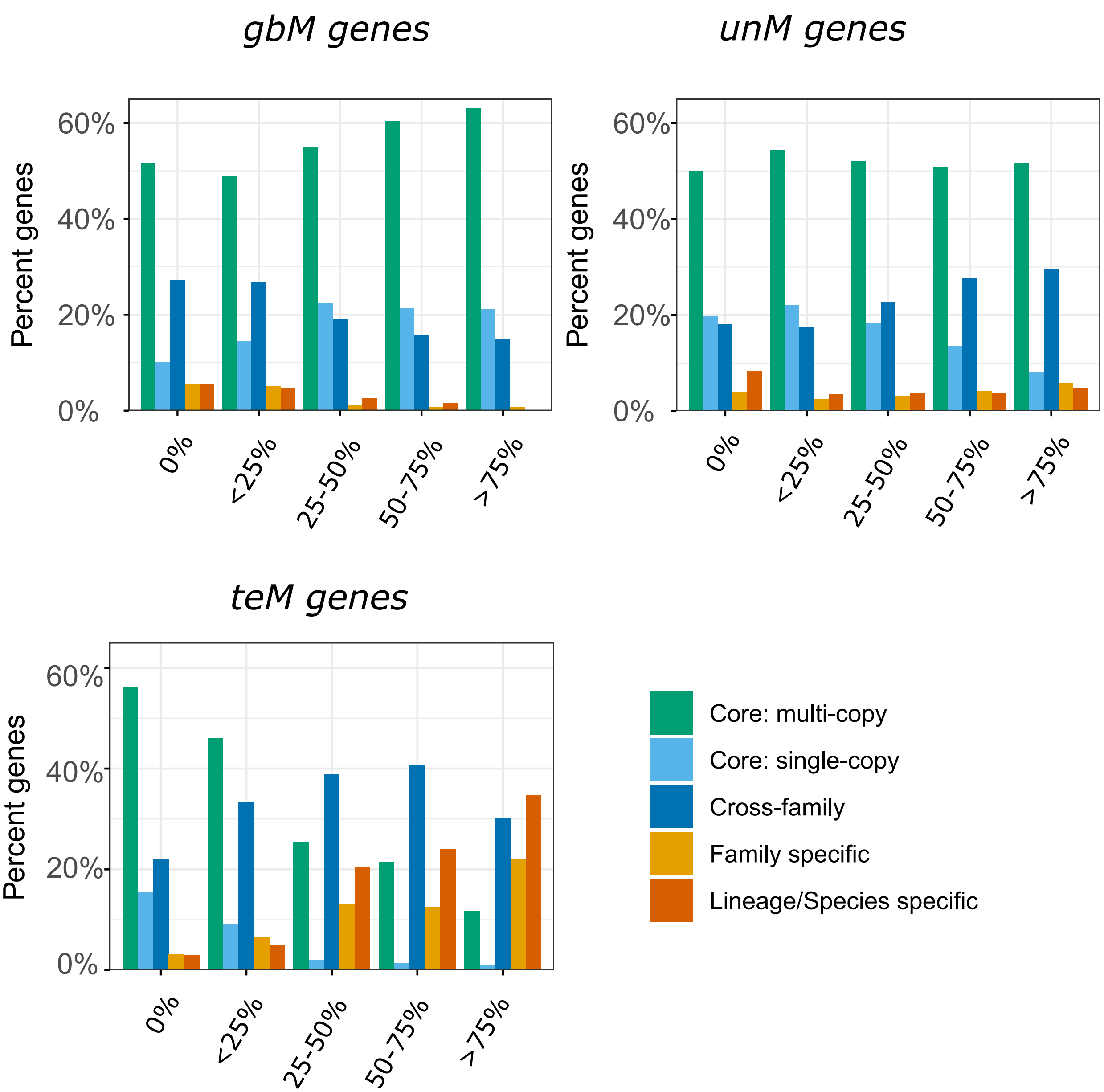
